# LipidQMap - An Open-Source Tool for Quantitative Mass Spectrometry Imaging of Lipids

**DOI:** 10.1101/2025.10.15.682422

**Authors:** Jonas Dehairs, Jakub Idkowiak, Nina Ravoet, Michele Wölk, Marco Giampà, Jan Schwenzfeier, Carla Pallarés-Moratalla, Amna Tahir, Sofia Guedri, Frank Vanderhoydonc, Xander Spotbeen, Vincent de Laat, Klaus Dreisewerd, Gabriele Bergers, Bart Ghesquière, Robert Jirásko, Michal Holčapek, Jens Soltwisch, Maria Fedorova, Johannes V. Swinnen

## Abstract

Mass spectrometry imaging (MSI) is a powerful tool in both basic and clinical research, enabling spatial visualization of biomolecules and drugs in tissue sections. However, factors influencing mass spectrometric response during imaging are increasingly recognized for their impact on apparent molecular distribution. Quantitative mass spectrometry imaging (qMSI) addresses this variability by incorporating analytical standards that undergo the same processes as analytes. While qMSI sample preparation protocols for omics-scale quantitative lipidomics are actively evolving, software solutions for downstream data processing remain scarce. Here, we introduce LipidQMap, the first open-source platform for processing omics-scale qMSI lipidomics data. LipidQMap applies one-point calibration, normalizing annotated lipid signals against class-specific standards on a pixel-by-pixel basis, and generates concentration heat maps in pmol/mm^2^. The software supports centroided data import, recalibration, and lipid identification using a built-in, user-modifiable lipid database. LipidQMap resolves Na/H adduct isobaric overlaps by leveraging sodiated-to-protonated adduct ratios of standards, validated across several MSI platforms using mouse brain sections. Type II isobaric overlaps are corrected using predicted isotopic patterns. Extensive validation demonstrates that LipidQMap is robust across MSI platforms and harmonizes qMSI data, enabling more accurate and reproducible spatial lipid quantification.

## Introduction

Mass spectrometry imaging (MSI) has emerged as a powerful technique for visualizing lipid distributions directly on the intact tissue surface. Among the leading ionization methods are matrix-assisted laser desorption ionization (MALDI), desorption electrospray ionization (DESI), and secondary ion mass spectrometry (SIMS), typically combined with mass spectrometers based on time-of-flight (TOF) or Fourier transform (FT)-based systems ^1–3^. MSI generates extensive lipidomics datasets, often comprising thousands of spectra per tissue section. MSI data share many characteristics with large-scale direct infusion MS (DI-MS), and thus, many analytical challenges encountered in DI-MS lipidomics are equally relevant to MSI. Key issues include matrix effects, where endogenous organic and inorganic sample components affect ionization efficiency and analytical response of desired analytes ^4–7^, lipid ion suppression ^4,8^, in-source fragmentation ^7,9^, and isobaric overlaps from lipids that differ by a single double bond (type II overlap) or from overlapping sodiated and protonated lipid adducts (Na/H adduct overlap) ^7,10,11^. Additionally, MSI-specific factors, such as the chemistry of MALDI matrix and the way of its deposition ^7^ or microextraction efficiency in DESI ^5^, can further impact lipid signal distributions and quantitation.

In bulk lipidomics, the application of standards has become a routine strategy to mitigate these challenges ^12^. Recently, building on protocols for qMSI of drugs or metabolites ^4,13,14^, Vandenbosch et al. adapted this approach for omics-scale MSI lipidomics, paving the way for lipid qMSI ^5^. By uniformly spraying lipid standards onto the tissue surface, they normalized the signals of endogenous lipids within a given class to that of a defined class-specific internal standard (IS). With the growing interest in qMSI, commercially available options, such as the MSI SPLASH^TM^ LIPIDOMIX for Mass Spec Imaging by Avanti Research, have been introduced and can be used for the correction of matrix effects and ion suppression, since co-sprayed lipid standards undergo ionization processes similar to endogenous lipids and yet can be distinguished from endogenous analytes due to the presence of heavy isotopes ^4,5,7^. IS also support the monitoring of in-source fragmentation, and can be used to resolve isobaric overlaps, as demonstrated in shotgun lipidomics ^15^. Importantly, this strategy could be adapted for MSI as well. Taken together, the systematic application of lipid standards in the MSI workflow contributes not only to increased accuracy of lipid quantification but also to intra- and inter-laboratory harmonization.

Despite the availability of various software tools for MSI, including non-commercial (e.g, Cardinal ^16^, ShinyCardinal ^17^, Ion-to-Image (I2I) ^18^, msIQuant ^19^, DataCubeExplorer ^20^, rMSI ^21^, massPix (lipidomics MSI) ^22^) and commercial (e.g, SCiLS Lab, LipoStar MSI ^23^, Pyxis MSI, HDI) solutions, none are specifically tailored to omics-scale qMSI lipidomics. Existing tools often lack integrated support for class-specific standard normalization for each annotated lipid and lipid-specific quantitation workflows, creating a gap between methodological advancements and bioinformatics capabilities.

To address this need, we introduce LipidQMap, an open-source software solution designed for omics-scale qMSI lipidomics. Featuring a user-friendly graphical user interface (GUI) and a permissive license, supporting further community-based developments, LipidQMap enables lipid annotation using user-modifiable lipidomics databases, parallel and vendor-neutral (.imzML) processing of several files, isobaric overlap corrections, as well as one-point calibration-based lipid quantification. By recalculating lipid intensities into concentrations based on one or several user-defined standards, LipidQMap transforms MSI datasets into accurate quantitative lipid maps, facilitating robust and reproducible interpretation of MSI-based spatial lipidomics data.

## Results and discussion

We developed LipidQMap, a comprehensive, open-source software package for the end-to-end processing of lipid mass spectrometry imaging (MSI) data. LipidQMap is designed to guide researchers through the entire analytical workflow within a single, user-friendly graphical interface (GUI), addressing the need for an accessible yet powerful tool for quantitative MSI (qMSI). Its core functionality encompasses all necessary steps to transform raw data into biologically meaningful results, including: **(i)** direct import of standardized .imzML data files, **(ii)** generation of ion images from a user-customizable lipid database, **(iii)** application of correction algorithms to resolve isobaric interferences between sodiated and protonated lipid ions and between lipids of the same class where natural isotopologues from more unsaturated species interfere with more saturated ones, and **(iv)** normalization against internal standards (IS) to produce quantitative lipid maps (in pmol/mm²). The tool was initially validated on healthy mouse brain tissue, which is frequently used as a reference sample for demonstrating novel MSI technologies targeting lipids. To ensure straightforward installation, standalone zero-external-dependency applications are provided for Windows (x64) and macOS (Apple Silicon). For Linux users and other platforms, the software can be run directly from the source code.

### Operating LipidQMap

Upon importing data as platform-independent .imzML files, users define the ion mode (positive or negative), annotation accuracy window (in ppm), and bin size for aligning centroided data collected across every pixel in the entire dataset (for timsTOF, the optimal bin setting is 5 mDa). A dynamic binning algorithm aligns centroided spectra across all pixels, maintaining constant resolution (ppm) across the entire mass range and automatically adjusting the absolute bin width (in mDa) across the entire *m/z* range. This ensures the binning strategy correctly reflects the mass accuracy at different *m/z*. Finally, the intensities of all peaks falling within each bin are averaged to produce the final mean spectrum.

LipidQMap supports lock-mass recalibration using user-selected *m/z* values, with a specific peak assignment tolerance of 30 ppm and a minimum intensity of 10 000 as default parameters optimized for timsTOF datasets. To ensure accurate recalibration, low-abundance signals – such as those with intensities <8000 a.u. on timsTOF instruments – should be avoided for recalibration as these may lead to incorrect signal assignment and compromise data quality. Recalibration parameters may need to be adjusted for datasets acquired using different mass spectrometer platforms.

Lipid annotation is performed using customizable .csv databases containing nine required columns with information provided by the user (**Figure 1A & Materials and Methods**), including lipid identifiers (at lipid species level), lipid class, the neutral formula, types of expected adducts, M-2 isobars, M+Na isobars, amount of IS sprayed on the surface in pmol/mm^2^, and for every lipid, the standard which should be used for the normalization. An example of a basic annotation list is provided in the **Supplementary Table 4**. The annotation lists that are included in the initial software release were generated using Python and organized in a specific order required by the software for data processing. The user can extend or crop them, e.g., by adding/removing lipid annotations, selecting specific ion types for qMSI (e.g., those with higher sensitivity), or defining specific standards (standard ion types) for the normalization of specific signals addressing, e.g., differences in the response of isotopically labeled analogs, double bond number, or different adduct types. It remains essential to preserve the proposed structure of the annotation list.

**Figure 1.**
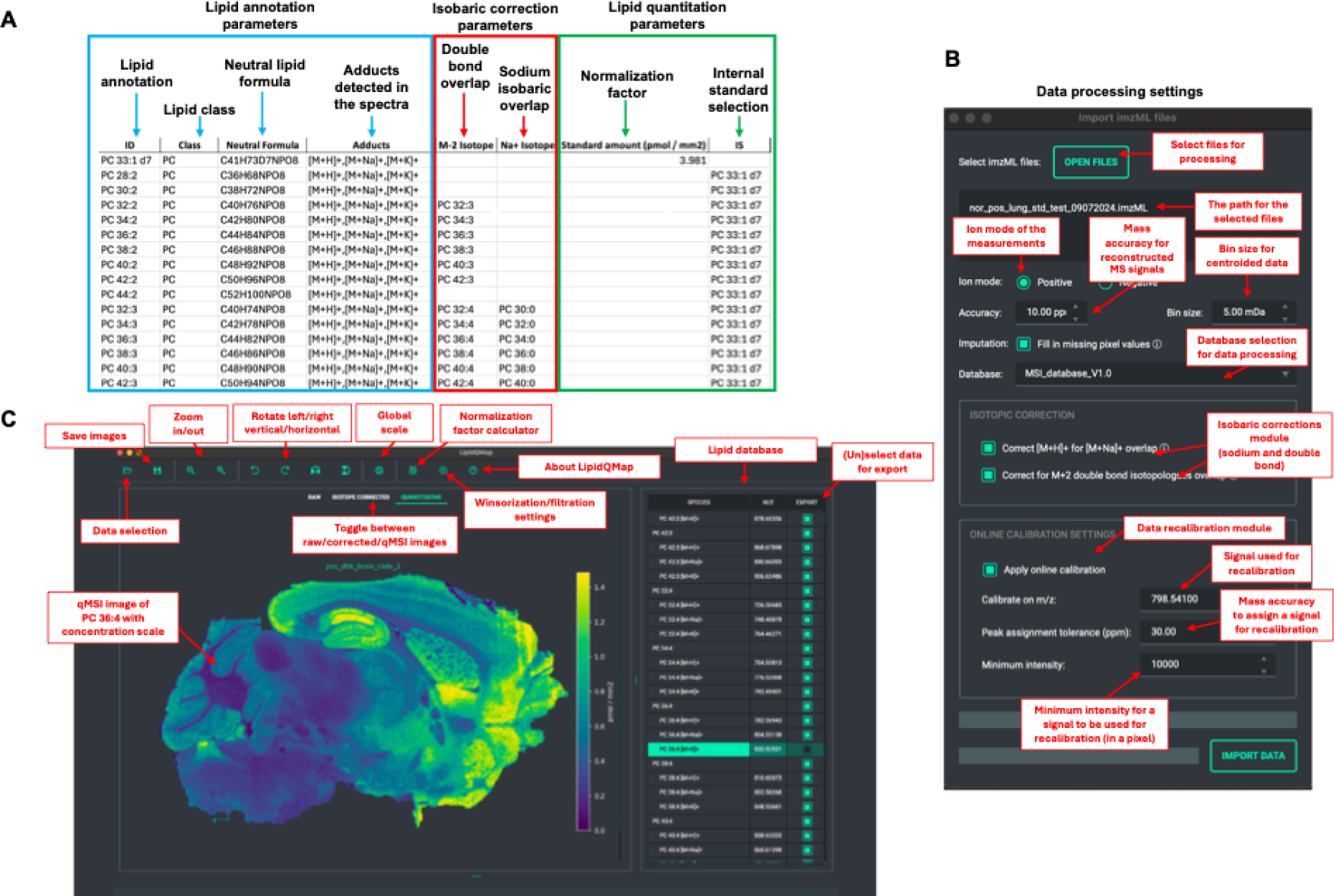
Illustration of LipidQMap database, settings, and user interface: **A**. Illustration of the database and parameter settings in LipidQMap for assigning lipid identifications, calculation of isobaric corrections, and quantitation of lipid concentrations on the tissue surface. **B.** Data processing settings. **C.** Graphical user interface (GUI). The image type can be toggled between Raw, Isotope Corrected, and Quantitative views. The toolbar above the MSI image provides tools for adjusting zoom, orientation, and presentation, e.g., using a global concentration scale across the entire lipid class. The Standard Calculator feature facilitates the computation of normalization factors. Under Settings, users can fine-tune parameters such as winsorization and filtration. The Lipid Database tab on the right allows users to switch between different adducts of lipid species or display the total surface concentration, calculated as the sum of all adducts.

Optional isobaric corrections for Na/H adduct overlaps and Type II overlaps are applied sequentially, improving reliability in datasets with prevalent M+Na adducts, as detailed below.

Quantification of the identified lipids is achieved through one-point calibration, where lipid signals are normalized to the corresponding standard applied to the sample surface. The median value from surrounding pixels replaces missing standard signals. LipidQMap enables imputation of empty pixels in both raw and isobaric-corrected datasets, a feature particularly advantageous for DESI imaging experiments where spray instability can result in systematic row-wise data loss.

To mitigate the influence of outlier pixels, LipidQMap implements winsorization — a statistical technique that replaces extreme values with less extreme percentiles. The degree of winsorization is user-adjustable via the settings (upper software menu). For timsTOF MALDI-imaging, with well-optimized standard application protocols, the ideal winsorization settings preserve 95-99% of the pixels. LipidQMap also enables independent winsorization of raw/isobaric corrected and standard-normalized images.

A screenshot of the data processing interface is shown in **Figure 1B**.

Once the parameters are defined, LipidQMap processes the MSI dataset and presents the annotated lipid signals in a navigable list within the main interface. Users can toggle between raw MSI, isobaric-corrected, and qMSI images, each accompanied by a concentration scale and interactive visualization tools. The software automatically highlights the most relevant identifications based on signal intensity of lipids and spatial completeness, while identifications with substantial missing pixel data remain unchecked by default. Users can refine this selection through adjustable filtering settings.

LipidQMap supports visualization of three image types: (i) raw ion intensity maps, (ii) isobaric-corrected images, and (iii) fully standard-normalized quantitative images (qMSI images). An intensity/concentration scale accompanies each image. Lipids from the same class can also be visualized on a unified concentration scale, facilitating comparative interpretation across the tissue section.

The interface allows users to rotate, mirror, zoom in/out, and arrange images for detailed inspection. To facilitate spectral validation, an average spectrum is generated, allowing for a close examination of peak location (mass accuracy) and surrounding signals. A dedicated species plot view summarizes the average abundance (concentration) of lipid species within each class across the entire section.

All image types—raw MSI, isobaric-corrected, and qMSI images—can be exported to a user-selected directory as one image per identification (with individual scales) or as a panel of selected lipids from the identification lists.

A comprehensive guide to software operation is available on the GitHub repository, along with the software.

The GUI is illustrated in **Figure 1C**.

### Application of LipidQMap for Na/H adduct isobaric correction in MSI

Isobaric overlap resulting from the occurrence of sodium adducts in lipid spectra is a recognized challenge in lipidomics (**Figure 2A** and **2B**). In such cases, the overlapping ion images typically correspond to two lipids of the same class, represented as [X+2:Y+3] [M+H]^+^ and [X:Y] [M+Na]^+^, with a Δ*m/z* of 0.0025. Simulations indicate that separating this mass difference requires a minimum resolution of 600 000. While alternatives involve employing ion mobility separation or applying buffer-based washes before matrix deposition to reduce excessive inorganic salt content, these approaches have notable limitations. Ion mobility imaging substantially increases acquisition time, whereas tissue washing can alter its morphology and composition. To address this, we adapt and extend the strategy proposed by Höring et al. ^15^ for shotgun lipidomics to mass spectrometry imaging data, implementing it on a pixel-wise basis. The correction is characterized in the **Materials and Methods** section. Its effect is firstly illustrated on an MSI dataset of a mouse brain section generated on a timsTOF instrument (**Figure 2**), specifically for PC 36:1 [M+Na]^+^ and PC 38:4 [M+H]^+^ (**Figure 2A**) and PE O-36:4 [M+Na]^+^ and PE O-38:7 (PE P-38:6) [M+H]^+^ (**Figure 2B**).

**Figure 2.**
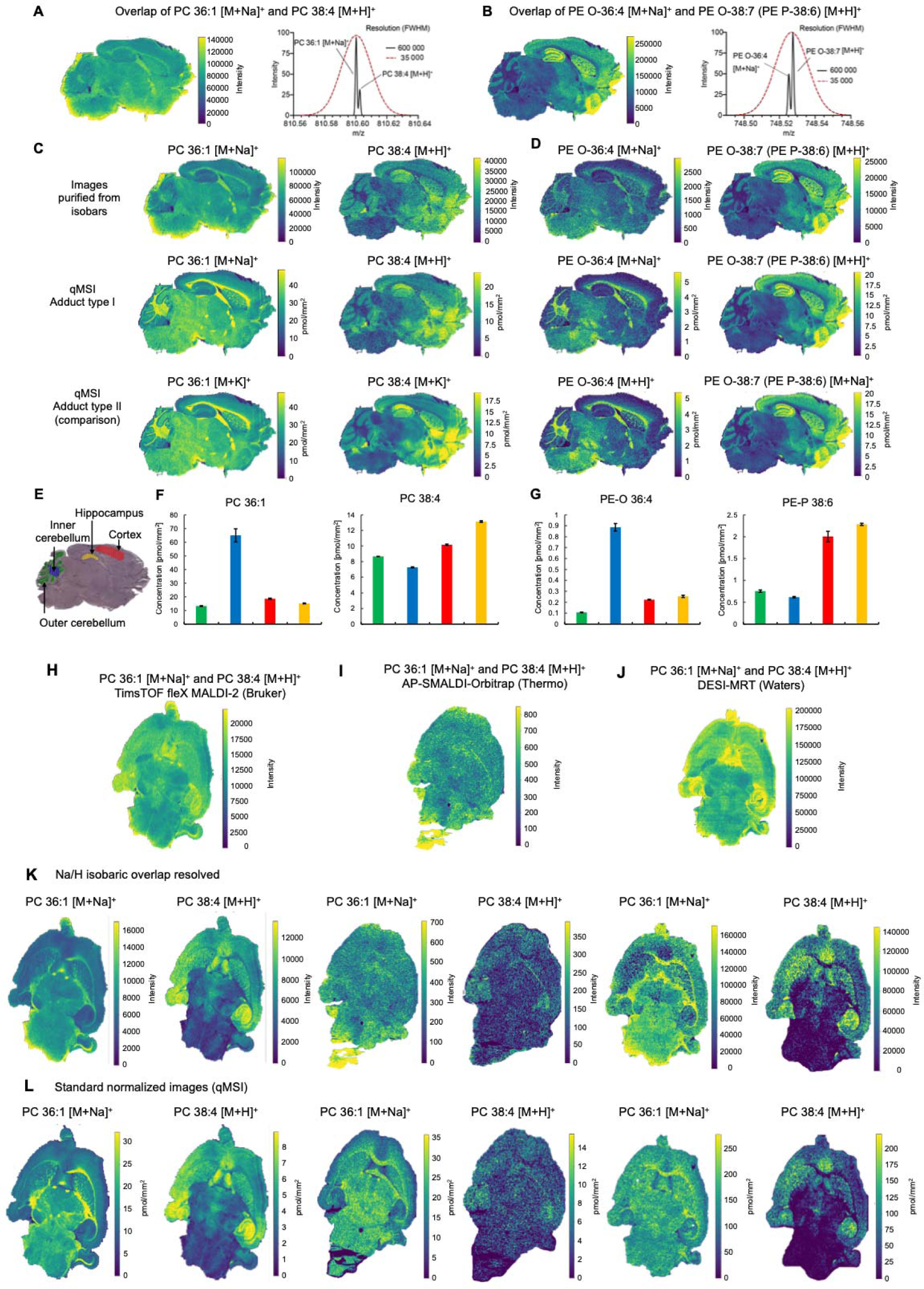
Application of LipidQMap for Na/H adduct isobaric correction in MSI and comparison of Na/H adduct isobaric correction across three mass spectrometers in two different laboratories (KU Leuven, TU Dresden). Sections in panels **A.-C.** were imaged at 30 × 30 µm and **H.-L.** at 50 × 50 µm. **A.** Raw MSI image illustrating the isobaric overlap of PC 36:1 [M+Na]^+^ and PC 38:4 [M+H]^+^ and corresponding MS spectrum of monoisotopic signals at 35 000 (red line) and 600 000 (black line) resolution (simulation). **B.** A raw MSI image showing the overlap of PE O-36:4 [M+Na]^+^ and PE O-38:7 (PE P-38:6) [M+H]^+^ with a comparison of separation capabilities between the same two mass spectrometers (simulation). **C.** MSI images of PC 36:1 [M+Na]^+^ and PC 38:4 [M+H]^+^ after Na/H adduct isobaric correction of raw images. The top row shows images purified from isobars, the middle row after standard normalization (qMSI image of [M+H]^+^ and [M+Na]^+^ adducts), and the bottom row includes an additional qMSI image of other detected adducts for comparison, normalized to PC [D5] 35:1. **D.** MSI images of PE O-36:4 [M+Na]^+^ and PE O-38:7 [M+H]^+^ after Na/H adduct isobaric correction of raw images. The top row shows images before standard normalization, the middle row after standard normalization, and the bottom row includes both protonated and sodiated adducts for comparison. Normalization was performed using PE 33:1[D7]. Isobaric separation was achieved using LipidQMap software, based on the sodium-to-proton adduct ratio of the lipid standard sprayed onto the surface. **E**. Hematoxylin-stained brain section used for laser capture microdissection. A total area of 5 mm^2^ was extracted for each region of interest (ROI) (pooled from multiple sections) for quantitative bulk lipidomics. **F, G.** Bulk lipidomics results (HILIC-MS/MS) for PC 36:1, PC 38:4, PE O-36:4, and PE P-38:6 obtained from the ROI. The standard deviation was obtained from two technical replicates. **H-J.** Raw MSI images showing overlapping signals from two lipid adducts: PC 36:1 [M+Na]^+^ and PC 38:4 [M+H]^+^, acquired on timsTOF fleX MALDI-2, AP-SMALDI Q Exactive^TM^ Orbitrap, DESI-MRT Select Series. **K.** MSI images after Na/H adduct isobaric correction of raw images by applying the sodium-to-proton adduct ratio of the standard sprayed onto the brain section surface. **L.** qMSI images after correction of the isobaric overlap between PC 36:1 and PC 38:4 and subsequent standard normalization (PC [D9] 32:0 for both MALDI platforms, and PC [D5] 35:1 for the DESI-qMSI).

Using the ratio of sodium and proton adducts of the IS sprayed on the brain section surface, including PC [D5] 35:1 and PE 33:1 [D7], the images of PC 36:1 [M+Na]^+^ and PC 38:4 [M+H]^+^ (**Figure 2C**), and PE O-36:4 [M+Na]^+^ and PE O-38:7 (PE P-38:6) [M+H]^+^ (**Figure 2D**) were successfully separated from one another. For comparison, the quantitative potassium adduct images are shown for PC 36:1 and PC 38:4 (**Figure 2C** – bottom row), along with protonated and sodiated images for PE O-36:4 and PE O-38:7 (PE P-38:6) (**Figure 2D** – bottom row). All exhibited a strong resemblance (**Figure 2C** and **2D** – middle and bottom rows) in specific regions of interest (ROIs) in the mouse brain, such as the hippocampus, cortex, cerebellum, and corpus callosum. Interestingly, the range of reported concentrations aligns well between different adducts (**Figure 2C** and **2D** – middle and bottom rows). To validate MSI trends, laser capture microdissection (LCM) and bulk lipidomics were performed on four ROIs: cortex (red), hippocampus (yellow), and two cerebellar areas (green and blue). Hematoxylin staining facilitated precise dissection (**Figure 2E**). Lipids were extracted using the Bligh-Dyer protocol, separated by hydrophilic interaction liquid chromatography (HILIC), and analyzed on a quadrupole-ion trap mass spectrometer (QTRAP) in multiple reaction monitoring (MRM) mode. The resulting bar charts (**Figure 2F-G**) corroborate the MSI-derived trends, including those obtained after Na/H adduct isobaric correction. Notably, comparison of the LCM quantitative bulk results with the qMSI images also revealed similar concentration ranges, despite differences in sample preparation protocols, analytical techniques (LC-MS/MS vs MALDI-MSI), ionization methods (ESI vs MALDI), measurement principles (multiple reaction monitoring vs full MS1 profile registration), and analytical factors such as biological matrix load, ionization suppression, and efficiency. Specifically, qMSI and LCM yielded comparable ranges of 0–40 pmol/mm² vs 0– 80 pmol/mm² for PC 36:1, 0–20 pmol/mm² vs 0–14 pmol/mm² for PC 38:4, 0–5 pmol/mm² vs 0–1 pmol/mm² for PE O-36:4, and 0–20 pmol/mm² vs 0–2.5 pmol/mm² for PE P-38:6.

To further assess the robustness of the correction, we applied it to data acquired from two additional platforms: the Select Series MRT (Waters) with desorption electrospray (DESI) ionization (KU Leuven), and the Q Exactive^TM^ Orbitrap with atmospheric pressure scanning microprobe MALDI (AP-SMALDI) ionization (TU Dresden) (**Figure 2H – L**). Considering the differences in sensitivity of the instruments, the isobaric pair PC 36:1 [M+Na]^+^ and PC 38:4 [M+H]^+^ was selected due to consistent detection of the PC [D9] 32:0 and PC [D5] 35:1 standards across all platforms. A horizontal section from the same brain was used to optimize acquisition settings. Despite differences in raw images, the normalized qMSI images showed consistent trends across platforms and matched bulk lipidomics data from cortex, hippocampus, and cerebellum. Although no significant optimization of the qMSI pipeline has been carried out across laboratories yet, comparable concentration ranges were observed between different mass spectrometry platforms. In particular, timsTOF fleX MALDI-2 and AP-SMALDI Q Exactive^TM^ Orbitrap produced similar results, such as PC 36:1 [M+Na]^+^ ranging from 0 to 30 pmol/mm² and 0 to 35 pmol/mm², and PC 38:4 [M+H]^+^ ranging from 0 to 8.5 pmol/mm² and 0 to 14.5 pmol/mm², respectively. When compared with the DESI-MRT platform - where a more concentrated IS mix was applied to counteract stronger ion suppression, along with notably different sample preparation, ionization, and analytical procedures - the resulting ranges remained consistent with those obtained using MALDI-based qMSI, i.e., 0-250 pmol/mm^2^ for PC 36:1 and 0-200 pmol/mm^2^ for PC 38:4. Altogether, these findings underscore the importance of applying standards in MSI to mitigate variability in analyte responses across different MSI platforms (inter-lab harmonization), as properly applied standards undergo similar ionization processes as endogenous analytes, compensating for differences in ionization efficiency, ion suppression, and microextraction efficiency.

Additionally, preliminary experiments indicate that accurate Na/H adduct isobaric correction requires sensitive detection of protonated adducts, specifically the high abundance of *[IS_i_ + H]^+^* and *I*[X:Y] components, as described in the **Materials and Methods** section. Consequently, the correction performs optimally when protonating matrices are used, such as carboxylic acids or phenols. This is particularly valid for dihydroxybenzoic acid (DHB), which yielded consistent results across all measurements, or dihydroxyacetophenone (DHAP) (effective when *RA^Na^* was below ∼2.5). At this stage, we recommend using DHB. Further research and optimization are necessary to fully leverage the potential of the correction in qMSI using various MALDI matrices.

In conclusion, the Na/H adduct isobaric correction originally developed for shotgun lipidomics was effectively adapted for MSI. It enables resolution and separation of lipid isobars resulting from the natural presence of sodium in tissues, without compromising spatial resolution, acquisition time, or analyte delocalization.

### Correction of type II isobaric overlaps using LipidQMap

In general lipidomics, a commonly observed isobaric overlap for lipids identified at the species level is referred to as a “double bond overlap” or type II overlap. This is mainly due to M+2 isotopologues (up to ∼15% of primary adduct intensity) and, to a lesser extent, M+4 isotopologues (up to ∼0.5% of primary adduct intensity) from more unsaturated lipids co-registered with saturated analogues. This is shown in **Figure 3A** and **3B** for PC 38:4 [M+H]^+^, PC 38:3 [M+H]^+^, and PC 38:2 [M+H]^+^ isotopologues, with abundances matching those observed in the average mouse brain MSI spectrum, assuming no Na/H adduct isobaric overlap from PC 36:1 [M+Na]^+^. Unresolved type II isobaric overlap therefore impacts the raw MSI image of PC 38:3 [M+H]^+^ through the intensity of the contributing PC 38:4 [M+2+H]^+^ isotopologue and likewise affects PC 38:2 [M+H]^+^ intensity via the abundances of PC 38:3 [M+2+H]^+^ and PC 38:4 [M+4+H]^+^ isotopologues.

**Figure 3.**
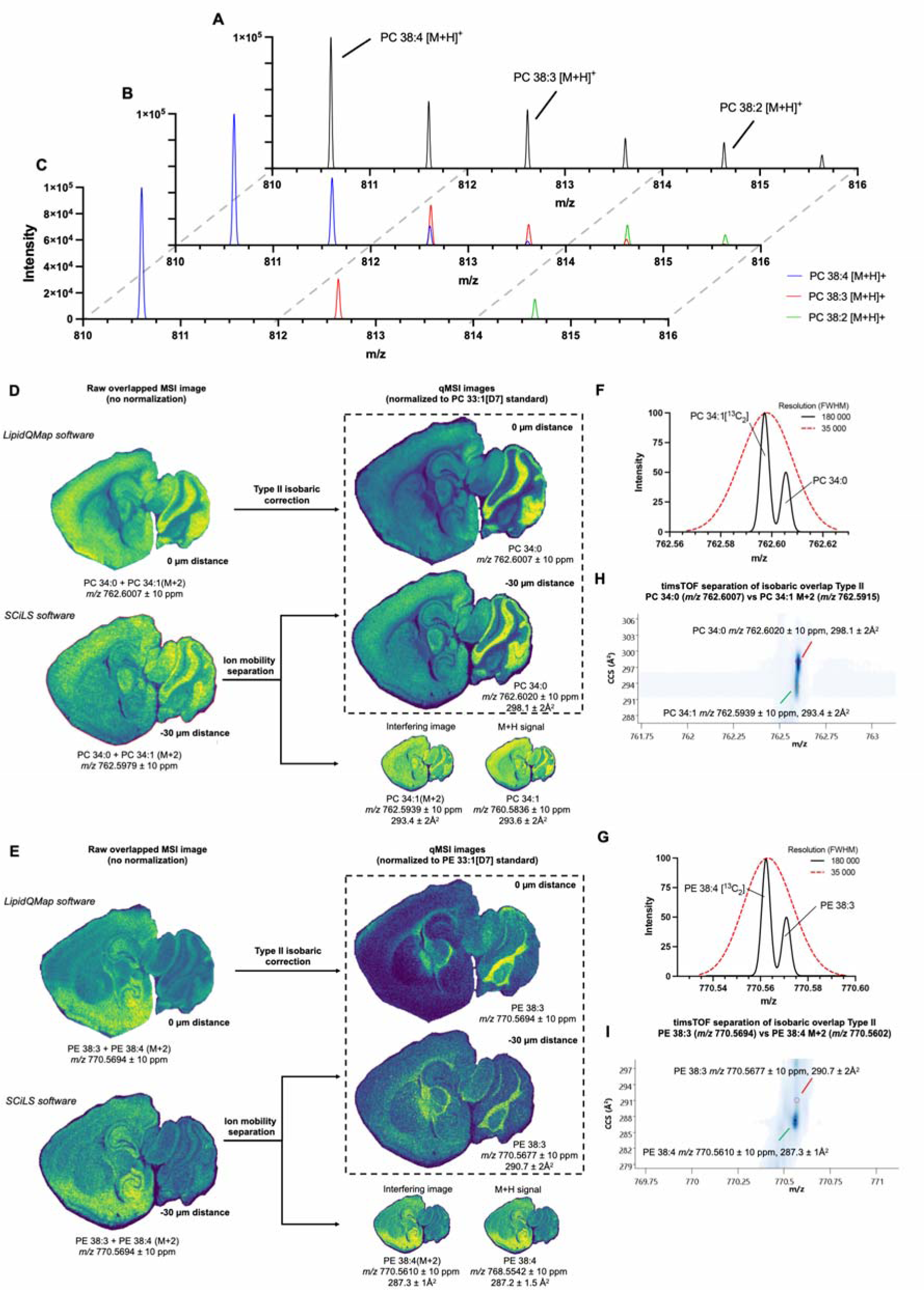
Correction of type II isobaric overlaps using LipidQMap and illustration of type II isobaric overlap correction by LipidQMap versus trapped ion mobility spectrometry (TIMS). Sections in panels **D.-E.** were imaged at 30 × 30 µm. **A.** Raw average MSI spectrum from a mouse brain section, highlighting type II isobaric overlap among PC 38:4 [M+H]^+^, PC 38:3 [M+H]^+^, and PC 38:2 [M+H]^+^. **B.** Contribution of isotopologues to the intensities of PC 38:3 [M+H]^+^ and PC 38:2 [M+H]^+^. **C.** Correction for Type II isobaric overlap using LipidQMap, resulting in refined estimates of individual intensities for PC 38:4 [M+H]^+^, PC 38:3 [M+H]^+^, and PC 38:2 [M+H]^+^. **D, E.** Comparison of qMSI images derived from mathematically corrected MSI data (LipidQMap) normalized against standards PC 33:1 [D7] [M+H]^+^ and PE 33:1 [D7] [M+H]^+^, with standard-normalized images of PC 34:0 [M+H]^+^ and PE 38:3 [M+H]^+^ purified via trapped ion mobility. **F, G.** Type II isobaric overlaps in mass spectra acquired at 35 000 FWHM (timsTOF operational resolution) and 180 000 FWHM for PC 34:1 [M+2+H]^+^ versus PC 34:0 [M+H]^+^ and PE 38:3 [M+H]^+^ versus PE 38:4 [M+2+H]^+^ (simulation). **H, I.** Mobilograms obtained using a timsTOF mass spectrometer with trapped ion mobility (TIMS), showing the resolution of Type II isobars with indications of m/z and collision cross-section (CCS) ranges. The sections were washed with chilled 150 mM ammonium formate and analyzed using MALDI-2 to reduce the influence of other adduct types on the outcome of this experiment.

Based on the simulations, such isobaric overlap can be resolved if 180 000 FWHM resolution is achieved by a mass spectrometer, such as Orbitrap, FT-ICR, or high-resolution Q-TOF mass spectrometers. Alternatively, mass spectrometers unable to achieve such resolving power, e.g., timsTOF, render the advantage of the ion mobility module ^11^. However, imaging runs with ion mobility are more time-consuming than straightforward MALDI or MALDI-2-based imaging. Alternatively, type II isobaric overlap can be resolved through mathematical correction based on the predicted isotope pattern. LipidQMap, after assigning the lipid identification to the measured *m/z*, performs a simple isobaric correction based on the isotopic patterns for M+2 isotopologues, retaining the estimated PC 38:3 [M+H]^+^ and PC 38:2 [M+H]^+^ abundances as presented in **Figure 3C** and explained in more detail in the **Materials and Methods** section.

Type II isobaric overlap correction by LipidQMap is illustrated with two examples: **(i)** PC 34:0 [M+H]^+^ detected at *m/z* 762.6007 overlapping with PC 34:1 [M+2+H]*^+^* at *m/z* 762.5915, and **(ii)** PE 38:3 at *m/z* 770.5692 overlapping with PE 38:4 [M+2+H]^+^ at *m/z* 770.5602. Due to the high abundance of PC 34:1 and PE 38:4, their isotopologues mask the less unsaturated species (**Figure 3D and E**). The timsTOF system without TIMS, offering approximately 35 000 resolution, cannot resolve these isobars (**Figure 3F and G**). Mobilograms showing the successful separation of the isobars by the timsTOF fleX system (using ion mobility) are presented in **Figures 3H** and I.

Comparison of ion-mobility-resolved isobars and mathematically corrected images revealed strong agreement (**Figure 3D** and **E**). The qMSI images of PC 34:1 [M+2+H]^+^ and PE 38:4 [M+2+H]^+^ closely resemble their [M+H]^+^ counterparts.

This result highlights the crucial role of applying such corrections in generating reliable maps of lipid distribution.

### Standard normalization and standard calculator

In the final step of data processing, all corrected MSI images are normalized using selected lipid class standards. Following the recommendations of the Lipidomics Standards Initiative (LSI), it is advisable to include at least one standard per lipid class (see LSI website for reference). In practice, however, multiple standards are often employed to better account for variations in analytical responses due to differences in fatty acyl chain length and degree of unsaturation. Ideally, each standard should yield a unique *m/z* value in the spectra, free from isobaric interference, to ensure accurate lipid quantitation. Using multiple standards further increases the likelihood that at least one will be suitable for normalization across MSI data.

To the best of our knowledge, LipidQMap is the first software to enable large-scale standard normalization of -omics MSI data using multiple user-defined standards, following the LSI recommendations.

As previously mentioned, LipidQMap utilizes a widely adopted one-point calibration approach (presented in the **Materials and Methods** section). LipidQMap includes a built-in calculator to facilitate the computation of the standard normalization factor. The calculator requires input parameters from the standards spraying protocol. The normalization process involves three main steps: **(i)** Sprayed area estimation: the dimensions of the sprayed area (*x* and *y* axes) and any margins, **(ii)** Delivered IS mixture volume estimation: the total volume of standard mix applied to the surface, accounting for syringe pump flow rate, initial equilibration time (if any), drying or pause intervals between layers, and the number of spraying cycles, **(iii)** Standard concentration calculation: the concentration of each standard in the IS mix is determined based on the molecular weight of the standard, stock concentration, volume used, final dilution, and the volume of the target solution. The total amount of standard delivered to the surface (in pmol) is then normalized to the sprayed area (pmol/mm^2^). Additionally, the normalized images can be winsorized independently of the raw intensity data to reduce the impact of extreme concentration values and improve image contrast. LipidQMap supports visualization of individual adducts as well as composite images representing the sum of all adducts (**Figure 4**).

**Figure 4.**
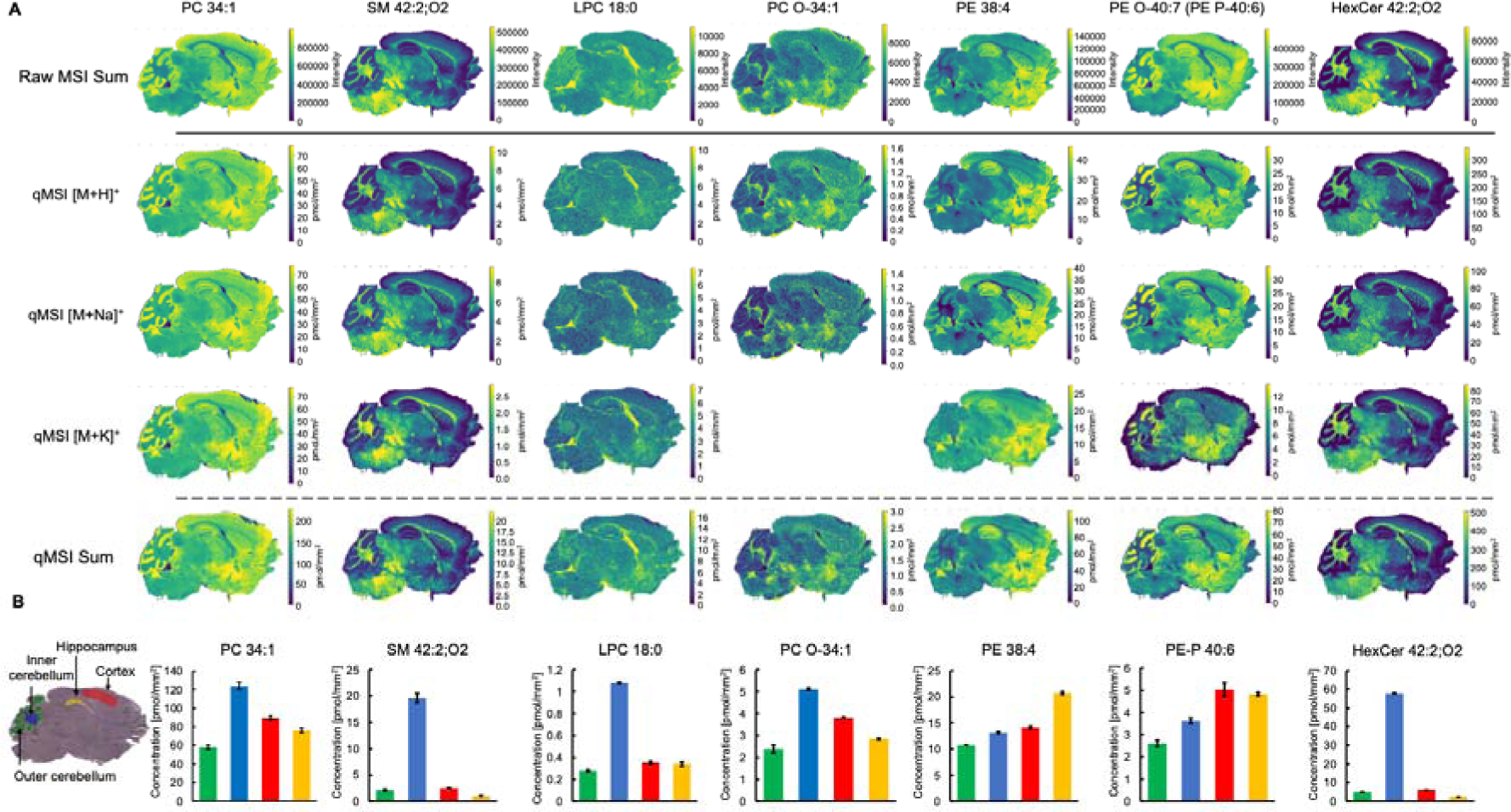
Quantitative mass spectrometry imaging of a healthy mouse brain section. The section was imaged at 30 × 30 µm. **A.** Quantitative MSI images of proton, sodium, and potassium adducts, along with their comparison to raw and quantitative images representing the summed intensities of all adducts. **B.** Bulk lipidomics data obtained from laser capture microdissected regions of interest (shown in the left lower corner). Selected lipid species included PC 34:1, SM 42:2;O2, LPC 18:0, PC O-34:1, PE 38:4, PE O-40:7 (PE P-40:6), and HexCer 42:2;O2.

Inspection of the individual adducts allows the effects of the applied corrections and normalizations to be clearly evaluated. In the raw images, discrepancies in the distribution of specific adducts were evident. These apparent differences between adducts largely disappeared in the resulting qMSI images (**Electronic Supplementary Data**). Notably, individual adducts showed similar concentration ranges. For example, all adducts of PC 34:1 fell within 0-70 pmol/mm^2^, whereas LPC 18:0 exhibited 0-10 pmol/mm^2^ across its adducts; more examples are presented in **Figure 4**.

To validate the trends seen in qMSI images, LCM and bulk lipidomics were performed on four ROIs as discussed above (**Figure 2**). These analyses confirmed the overall distribution pattern of the presented lipid species across the brain. As illustrated in the bar charts (**Figure 4B**), for instance, SM 42:2;O2 and HexCer 42:2;O2 were highly enriched in the inner cerebellum and nearly absent in the other regions. The corresponding qMSI images supported these findings, with predominantly low abundance across remaining areas. Other lipid species exhibited a more uniform distribution across the brain. However, bulk analysis still indicated a higher abundance in the inner cerebellum, as observed for PC 34:1 and PC O-34:1. The qMSI images likewise reflected this trend, albeit with subtle regional differences. Similarly, species such as PE 38:4 and PE O-40:7 were found to be more abundant in the hippocampus and cortex according to bulk measurements, and this trend is once again reproduced in the qMSI data. Finally, apparent discrepancies were observed between the raw and qMSI sums in the cerebellar region when compared with the bulk lipidomics results.

### Application of LipidQMap for imaging data processing of more complex tissue samples: case study of glioblastoma

While a healthy mouse brain is frequently used as a reference sample for demonstrating new MSI technologies targeting lipids, biological studies often involve more complex tissue types, such as malignant tumors with highly heterogeneous characteristics, necrotic regions adjacent to healthy tissue, and areas with immune cell infiltration. Imaging these heterogeneous samples introduces challenges due to factors such as variations in inorganic ion concentrations (e.g., Na^+^, K^+^), pH, differences in biomolecular composition, pathological features (e.g., cell organization and density), and inconsistent analyte microextraction efficiency, whether via MALDI or DESI.

To address these complexities, a series of experiments was conducted with murine NFPP10 glioblastoma, which exhibits typical features of human glioblastoma ^28^, across multiple MSI platforms: timsTOF fleX MALDI-2 (Leuven and Münster), MALDI-Q Exactive^TM^ Orbitrap (Münster), and AP-SMALDI-Q Exactive^TM^ Orbitrap (Dresden). All datasets were processed using LipidQMap. As illustrated in **Figure 5A**, standard normalization resulted in pronounced changes in lipid spatial distribution, particularly within tumor regions. The resulting qMSI images showed strong concordance with quantitative bulk lipidomics data obtained from ROIs (tumor, tumor rim, cortex, corpus callosum), outperforming raw MSI images in terms of biological relevance (**Figure 5A, B**). For instance, qMSI clearly highlighted the downregulation of PC 32:0 in the tumor core, a trend also observed for Cer 42:2;O2, with both findings confirmed by bulk analysis of LCM-extracted ROIs. Furthermore, qMSI revealed a higher abundance of SM 40:1;O2 in healthy mouse brain areas compared with raw images, which is also in agreement with bulk measurement results.

**Figure 5.**
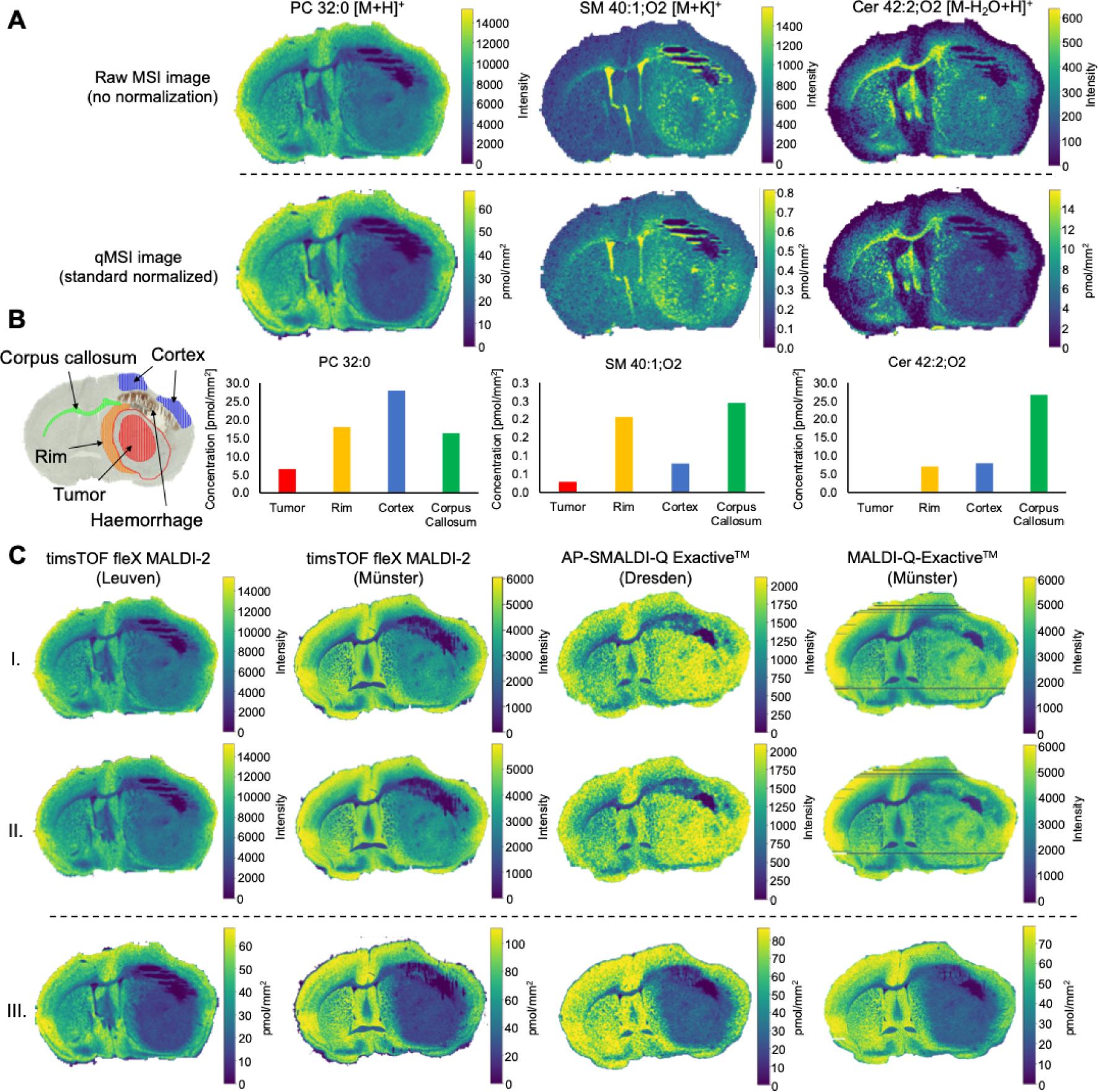
Application of LipidQMap for MSI data processing in a mouse glioblastoma model. Sections in panel **A.** were imaged at 50 × 50 µm, and in panel **C.** (from the left side): at 50 × 50 µm (Leuven), 17 × 17 µm (Münster), 50 × 50 µm (Dresden), and 20 × 20 µm (Münster). The impact of standard normalization on raw MSI images was evaluated using a glioblastoma mouse brain model via four different mass spectrometry platforms: timsTOF fleX MALDI-2 (Leuven, Belgium), timsTOF fleX MALDI-2 (Münster, Germany), MALDI-Q Exactive^TM^ Orbitrap (Münster, Germany), and AP-SMALDI-Q Exactive^TM^ Orbitrap (Dresden, Germany). All datasets were processed using LipidQMap. **A**. Comparison of raw MSI images of three lipid species - PC 32:0 [M+H]^+^, SM 40:1;O2 [M+K]^+^, and Cer 42:2;O2 [M-H_2_O+H]^+^ - before and after normalization using standards PC [D9] 32:0 [M+H]^+^, LPC [D9] 16:0 [M+K]^+^, and Cer 30:1;O2 [M-H_2_O+H]^+^, respectively. **B.** Quantitative bulk HILIC-MS/MS lipidomics results from LCM-extracted ROIs as indicated in the brightfield brain scan: blue – brain cortex, orange – tumor rim, red – tumor (glioblastoma), green – corpus callosum. **C**. Cross-platform comparison of PC 32:0 [M+H]^+^ images in raw (**I.**), isobaric-corrected (**II.**), and standard normalized formats (**III.**). Normalization was performed using PC [D9] 32:0 [M+H]^+^. The qMSI image of PC 32:0 [M+H]^+^ acquired via MALDI-Q Exactive^TM^ Orbitrap (Münster, Germany), contained empty pixel lines, which were corrected using LipidQMap based on the median lipid concentrations of surrounding pixels.

Among the reported lipid species, PC 32:0 [M+H]^+^, detected across all platforms, was used to demonstrate the consistency of qMSI images post-normalization. Whereas raw and isobaric-corrected images showed variability across instruments, especially within the glioblastoma tumor and rim areas, qMSI images were highly comparable between platforms. Moreover, the results also aligned well with quantitative bulk data (**Figure 5C**). The application of standards corrected the variability in response across all the MSI platforms used. While qMSI images generally aligned well with LCM-derived bulk lipidomics data, and the applied correction typically improved concordance with the fully quantitative bulk lipidomics trends, discrepancies for certain lipid species remained (data not shown), underscoring the need for further investigation into analytical chemistry factors affecting lipid response in MSI - a process that tools like LipidQMap can support.

Notably, in **Figure 5C** (last panel on the right, MALDI-Q Exactive^TM^ Orbitrap measurements), the missing pixel lines visible in the raw and isobaric-corrected MSI images were corrected in the qMSI image by LipidQMap. This was achieved by replacing missing pixels with the median lipid concentration of surrounding pixels, preventing division by zero during standard normalization.

Furthermore, Na/H adduct isobaric correction was evaluated to address the isobaric overlap between PC 36:1 [M+Na]^+^ and PC 38:4 [M+H]^+^ across three MSI platforms (violet panel): timsTOF fleX MALDI-2 (Leuven and Münster) and AP-SMALDI-Q Exactive^TM^ Orbitrap (Dresden). The resulting qMSI images were validated against LCM-derived bulk lipidomics (**Figure 6A**). Mathematically deconvoluted qMSI images closely resembled those of other adducts of PC 36:1 (red panels) and PC 38:4 (blue panels) and aligned well with bulk lipidomics data (**Figure 6B**). Moreover, the obtained concentration ranges are highly consistent between the MSI platforms and across all examined adducts, and correspond well with quantitative bulk lipidomics of LCM-extracted ROIs.

**Figure 6.**
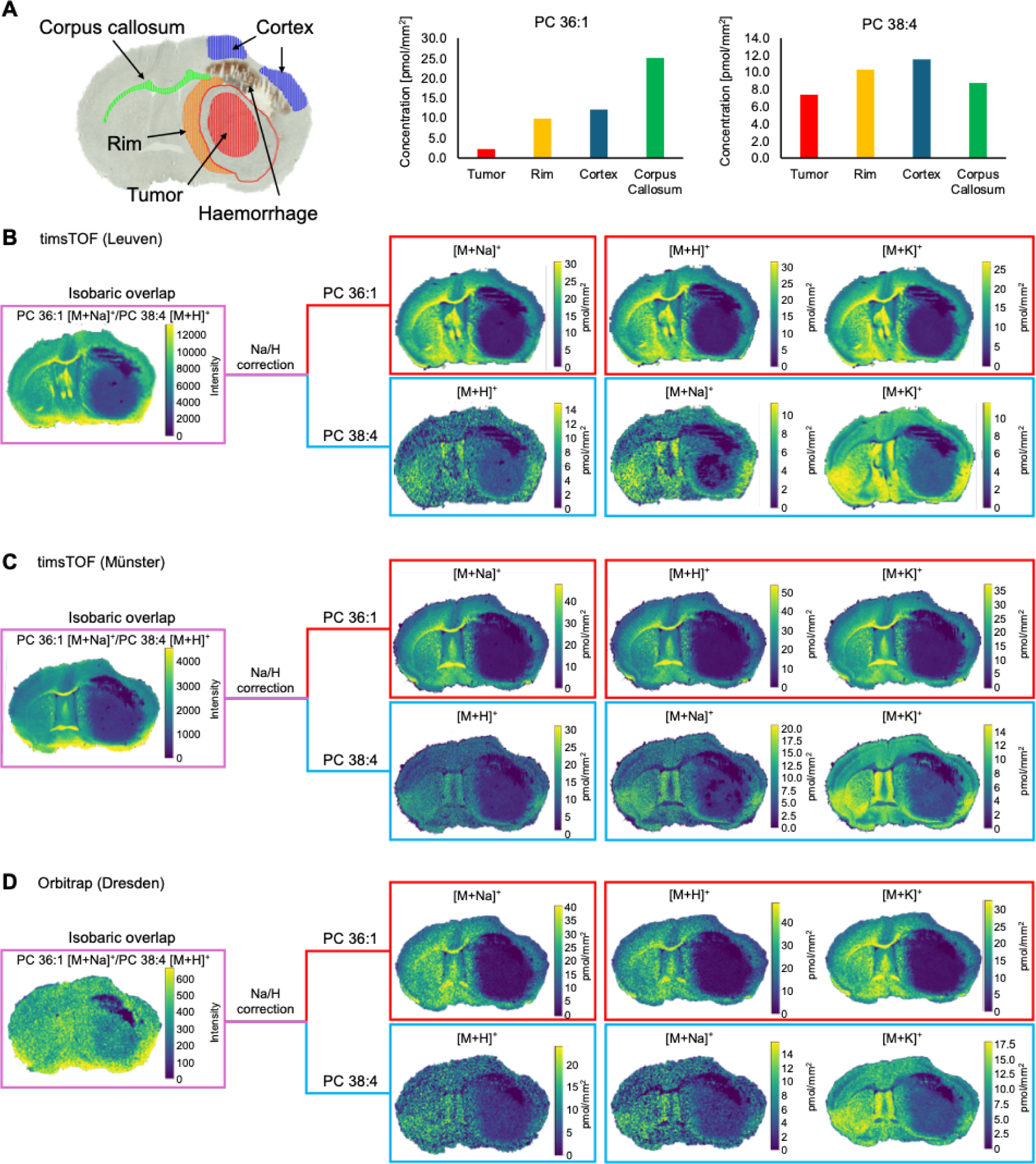
Cross-platform Na/H adduct isobaric correction of the glioblastoma case study using LipidQMap. The application of LipidQMap for Na/H adduct isobaric correction was evaluated using the glioblastoma mouse brain model described in Figure 5 using three mass spectrometer platforms: timsTOF fleX MALDI-2 (Leuven, Belgium), timsTOF fleX MALDI-2 (Münster, Germany), AP-SMALDI-Q Exactive^TM^ Orbitrap (Dresden, Germany). Sections were imaged at 50 × 50 µm (Leuven), 17 × 17 µm (Münster), and 50 × 50 µm (Dresden). To minimize technical variability, standards and matrix (DHAP) were uniformly applied in Leuven using an HTX M5 sprayer on adjacent sections. **A.** The regions of interest (ROIs) were extracted via laser capture microdissection (LCM) from a mouse brain harboring a glioblastoma tumor. The tumor core is highlighted in red with additional ROIs from the cortex (blue), corpus callosum (green), and tumor rim (orange). These regions were subjected to bulk lipidomics to validate MSI-derived lipid distribution trends. **B-D.** Na/H adduct isobaric correction was performed to resolve isobaric overlap between PC 36:1 [M+Na]^+^ and PC 38:4 [M+H]^+^ using the PC [D9] 32:0 standard. The correction was applied via LipidQMap across datasets from Leuven (timsTOF fleX MALDI-2, upper panel), Münster (timsTOF fleX MALDI-2, middle panel), and Dresden (AP-SMALDI-Q Exactive^TM^ Orbitrap, lower panel). Ion images of the mathematically separated signals for PC 36:1 [M+Na]^+^ and PC 38:4 [M+H]^+^ are shown for comparative evaluation.

Overall, LipidQMap demonstrates robust performance in data processing across diverse MSI platforms and vendors. The presented data highlight the importance of standard normalization in MSI of complex tissues, such as glioblastoma, where it significantly influences the observed lipid distribution.

### Conclusions and perspectives

In this report, we introduce LipidQMap, the first open-source, vendor-independent software designed for the processing of omics-scale quantitative mass spectrometry imaging (qMSI) lipidomics data. The software provides a user-friendly interface with integrated guidance to simplify and streamline the workflow.

LipidQMap is capable of processing any centroided lipid imaging data in .imzML format, assuming that such datasets can be treated as large-scale shotgun experiments. The software offers several key functionalities: recalibration of raw data against a selected signal, visualization and winsorization of data, mathematical correction of Na/H adduct isobars based on the ratio of sodium to proton adducts from analytical standards applied to the sample surface, and correction of type II overlaps from M+2 isotopologues of more unsaturated lipid analogs, using predicted isotope patterns. For quantification, LipidQMap performs pixel-by-pixel normalization of deconvoluted and annotated lipid signals using a one-point calibration method with user-selected analytical standards. Final qMSI images can be generated for individual adducts or as a composite sum of all adducts. These images can be individually winsorized and exported for further analysis. Additionally, the software includes a built-in standard calculator to assist in computing normalization factors and supports visualization of lipid classes using a unified concentration scale.

These algorithms and qMSI were validated across five independent MSI platforms in three expert laboratories, using both healthy mouse brain sections and more complex glioblastoma brain tissue characterized by significant pathological heterogeneity. This study also demonstrates that the use of analytical standards significantly improves mass spectrometer response, enhancing the comparability of MSI results with fully quantitative bulk lipidomics. Upcoming developments will extend the type II isobaric correction to M+4 isotopologues as well as dual ion mode imaging qMSI. LipidQMap will also support the export of standard-normalized datasets for downstream analysis in other MSI data processing software platforms such as Cardinal, LipostarMSI, and SCiLS.

## Materials & Methods

### Chemicals

MSI SPLASH^TM^ (#330841), PC 15:0/18:1 [D7] (#791637), PC [D5] 17:0/18:1 (#855681), PC [D5] 17:0/16:1 (#855682), PC [D9] 16:0/16:0 (#860352), PE 15:0/18:1 [D7] (#791638), LPC 18:1 [D7] (#791643), SM d18:1;O_2_/18:1 [D9] (#791649), L-CAR [D9] 16:0 (#870323), UltimateSPLASH^TM^ ONE internal standard mix (#330820), SphingoSPLASH^TM^ I internal standard mix (#330734), GM1 18:1;O_2_/18:0 [D7] (#860120), GM3 18:1;O_2_/18:0 [D5] (#860073), GD1b 18:1;O_2_/18:0 [D7] (#860125) were purchased from Avanti Polar Lipids (Alabama, USA), LPC [D9] 16:0 (#L-1516D), LPC [D9] 18:0 (#L-1518D) from Echelon Biosciences (Utah, USA). Cer 18:1;O_2_/12:0 (#22530) was from Cayman Chemical (Michigan, USA). Norharmane (NOR), 2’,5’-Dihydroxyacetophenone (DHAP), and 2’,5’-dihydroxybenzoic acid (DHB) were purchased from Tokyo Chemical Industry Co. (Tokyo, Japan). Ammonium formate, ammonium acetate, carboxymethylcellulose sodium salt (CMC), 2,6-di-tert-butyl-4-methylphenol, and Dulbecco’s Phosphate Buffered Saline, without calcium, without magnesium (DPBS) were purchased from Sigma-Aldrich (Missouri, USA). Isoflurane (Iso-Vet, 1000 mg/g) was purchased from Piramal Critical Care (Voorschoten, Netherlands), and Mayer’s hematoxylin was from Dako (Glostrup, Denmark). Hydrochloric acid (1 mol/l) was purchased from VWR Chemicals (Pennsylvania, USA). Chloroform was from Avantor (Pennsylvania, USA), and LC/MS-grade methanol, water, and acetonitrile were from BioSolve (Valkenswaard, Netherlands). Acetone was obtained from Thermo Fisher Scientific (Massachusetts, USA).

### Sample preparation for mass spectrometry of lipids

Mass spectrometry imaging of lipids

#### Sample selection

Healthy mouse brain sections were used to illustrate the Na/H adduct and Type II correction in the first chapters of the manuscript, and they were also used to optimize MSI methods with collaborators (AP-SMALDI-Q Exactive^TM^ at TU Dresden and DESI-MRT at VIB-KU Leuven). In contrast, the glioblastoma mouse model was used to compare results across collaborating laboratories, including VIB-KU Leuven, TU Dresden, and the University of Münster.

#### Collection of healthy mouse brain samples

Brain tissue was obtained from mice euthanized under deep isoflurane anesthesia, followed by cervical dislocation with approval from the institutional Ethical Committee for Animal Experimentation at KU Leuven, Belgium (M018/2024), rinsed with DPBS, gently dried using a Kimtech wipe, and then embedded in a 3% CMC medium in a Peel-A-Way cryomold. The sample was frozen on dry ice and stored at -80 °C until further processing.

#### Murine glioblastoma

All animal experiments were performed in accordance with the local Ethical Committee for Animal Experimentation (190/2021) of KU Leuven. NFpp10 glioblastoma cells were obtained from Inder Verma (Salk Institute, CA, US). They were generated from embryonic C57BL/6 neural stem cells infected with shP53, shNF1, and shPTEN containing lentiviral vectors, and a GFP/luciferase reporter. For intracranial tumor transplantation, 7–9-week-old C57BL/6 mice (Charles River Laboratories) were anaesthetized and placed into a stereotactic frame. A small hole was made 2 mm right and 1 mm anterior to the bregma. 20000 NFpp10 GFP/luc positive cells were slowly injected in a volume of 2 μl into the right striatum at a depth of 3.0 mm from the dura. Tumors were grown for 14 days before euthanasia by an injection of Dolethal followed by cold-DPBS perfusion. After cold DPBS perfusion, the brains were carefully extracted, cut at the injection point, and embedded in 3% CMC. Tissues were fresh-frozen in pre-chilled isopentane using liquid nitrogen and stored at -80 °C until further processing.

#### Cryosectioning of brain samples

10 µm sections were obtained using the HM 525NX cryostat (Thermo Fisher Scientific, MA, USA) at -20°C, mounted on IntelliSlides (Bruker Daltonik GmbH) for timsTOF fleX MALDI-2 instruments, standard microscope glasses (Labsolute) for AP-SMALDI- and MALDI-Q Exactive^TM^ Orbitrap, and Superfrost slides (Epredia) for DESI MRT Select Series, and stored at -80°C. For Laser Capture Microdissection (LCM), FrameSlides were used with a 1.4 µm PET membrane (Leica Microsystems, item no. 11505151). After mounting, the slides were dried in a vacuum desiccator for ∼10 minutes, then frozen on dry ice and stored at -80 °C.

#### Deposition of standards and matrices

Frozen tissue sections were dried in a vacuum desiccator for ∼30 min and used for matrix deposition as it is or after washing with aqueous 150 mM NH_4_HCO_2_ solution ^24^, to reduce the interferences of sodium and potassium adducts, and then dried in a vacuum desiccator. Lipid standard mixtures (**Supplementary Table 1**) were sprayed from a syringe onto the tissue surface using a hyphenation of a KD Legato 100 Infuse Single pump (kdScientific) and an HTX M5 sprayer (HTXImaging, HTX Technologies, LLC, Chapel Hill, US) ^5^. After spraying standards, the peristaltic pump was reconnected to the sprayer (Azura P4.1S, Knauer), and matrices were deposited on the sections. DHB (15 mg/ml in CHCl_3_:MeOH, 2:1, *v/v*) was used for healthy mouse brain sections, while DHAP (5 mg/ml in CHCl_3_:MeOH, 2:1, *v/v*) was used for glioblastoma-containing tissues. All sprayer settings for standard and matrix delivery are summarized in the **Supplementary Table 2**. All samples were uniformly prepared at KU Leuven (Leuven, Belgium), including the application of standards and matrix, and subsequently transferred to collaborating laboratories within 24 h after sample preparation.

#### Quantitative bulk lipidomics

Laser capture microdissection (LCM) and lipid extraction protocol After cryosectioning, tissue slides were dried in a vacuum desiccator, stained in an aqueous hematoxylin solution for 3 minutes, washed with tap water, and then dried under the vacuum. Selected regions were excised using a Leica Laser Microdissection (LMD) 6000B microscope equipped with a diode-pumped, solid-state laser (wavelength - 355 nm, average pulse energy of 70 µJ, repetition rate of 80 Hz). Regions of interest (ROIs) were selected to obtain a total of 5 mm² across five consecutive sections and captured in 0.2 ml PCR tube caps (Greiner). The material was stored at -80 °C until further processing. Before lipid extraction, samples were thawed on ice and spun at 5000×g for 20 minutes at 4°C to collect tissue fractions at the bottom of the tube. 200 µl of methanol was added, and the samples were sonicated for 30 minutes in the ultrasonic bath. The homogenates were transferred to 12 ml Pyrex glass tubes, rinsing the PCR tube three times with 200 µl of fresh methanol portions (amounting to a total volume of methanol 800 µl). Next, 100 µl of HCl 1M, 700 µl of LC-MS grade water, 825 µl of chloroform, 5 µl of antioxidant (2,6-di-tert-butyl-4-methylphenol, BHT, Sigma Aldrich) (10 mg/100 µl in 100% ethanol), 3 µl of lipid standards (UltimateSPLASH™ ONE internal standard mix and of SphingoSPLASH™ I internal standard mix) were added. The tubes were vortexed twice for 30 seconds and then spun at 6000×g at 4°C for 10 minutes. 600 µl of the chloroform phase was recovered with a glass Pasteur pipette and transferred to a 4 ml glass tube to be evaporated in a Savant Speedvac spd111v (Thermo Fisher Scientific, Massachusetts, USA). Dried lipids were reconstituted in 60 µL of ethanol (vortexed twice for 30 seconds, then for 1 minute) and transferred to amber vials with glass insert tubes for measurements.

### Mass spectrometry measurements

Mass spectrometry imaging of lipids

timsTOF fleX MALDI-2 at KU Leuven

MALDI-MSI imaging was performed on a timsTOF fleX MALDI-2 mass spectrometer (Bruker Daltonik, Bremen, Germany). The detailed settings for the trapped ion mobility and mass spectrometer used in every experiment presented in the manuscript are summarized in the **Supplementary Table 3.**

#### AP-SMALDI at TU Dresden

The MALDI-MSI imaging was performed on a Q Exactive^TM^ Plus Hybrid Quadrupole-Orbitrap mass spectrometer (Thermo Fisher Scientific, Bremen, Germany) equipped with an atmospheric pressure (AP)-SMALDI5 AF source (Transmit GmbH, Giessen, Germany). Instrument control was achieved using Q Exactive^TM^ Plus Tune 2.13 with a Leveraged license (Thermo Fisher Scientific, Bremen, Germany). SMALDI Control V1.4 (Transmit GmbH) was used for acquisition and stage movement. The data was acquired in positive ion mode, using a full MS scan. The *m/z* range was set to 350-1300 with a mass resolution of 140 000 at *m/z* 200. The automatic gain control target was set to 1 × 10^6^, with a maximum ion injection time of 500 ms. Microscans were set to 1, and a spray voltage of 3 kV (transfer capillary voltage) was applied. The capillary temperature was maintained at 250°C, and the S-lens RF level was set to 35. MALDI imaging was performed in 2D spot mode with a pixel size of 50 × 50 µm and acquired in 2D topography. A pulsed laser with 50 shots at 100 Hz per pixel was applied. The attenuator was set to 37° for measurements of healthy brain sections and 37.5° for measurements of the sections derived from the glioblastoma mice. Raw data files were converted to .imzML format using Raw2IMZML V1.8r3 (Transmit GmbH, Giessen, Germany).

#### DESI-MSI at KU Leuven

A Waters ACQUITY M-Class µBSM pump was used to continuously deliver a methanol:isopropanol (9:1, *v/v*) mixture at a flow rate of 2 µl/min to the XS DESI source coupled with the MRT Select Series Q-TOF mass spectrometer (Waters, Massachusetts, USA). Initial calibration was performed using polyalanine at a concentration of 5 mg/ml in a methanol:water (95:5, *v/v*) mixture deposited on a Superfrost slide. PC 34:1 detected at *m/z* 798.5410 was used as a lock mass.

Instrumental parameters were set as follows: capillary voltage, 0.57 kV; cone voltage, 116 V; ion source temperature, 95°C; API gas pressure, 0.05 psi; and heated transfer line temperature, 420°C. MRT voltage settings were optimized for enhanced detection of lipids with *m/z* values above 400 obtaining a single-pixel mass resolution of 162 000 at *m/z* 798.5410. Ion guidance was achieved using a StepWave RF set at 200 V and ion guide RF at 700 V. The trap RF was maintained at 400 V. Transfer RF was set to 457 V with a gain of 5. The Travelling Wave (TW) velocity was configured at 300 m/s with a pulse height of 0.2 V. Entrance and exit voltages for the ion guide were 2.0 and 0.0 V, respectively. The DC offset was set to -4.0 V. Quadrupole resolution settings included LM resolution at 4.9 and HM resolution at 15.0. The MS profile was set to Manual at *m/z* 400, 800, and 1200, dwell time (%scan time) 10 and 20, ramp time 20 and 50. Collision energy was applied at 4 V, with a collision gas flow rate of 1.2 ml/min. The detector voltage was maintained at 1.90 kV. The Enhanced Frequent Encoding (EFP) was set to Automatic. The mass range was set at 50-2400. The scan time was set to 0.2 seconds per pixel, with a pixel resolution of 50 × 50 µm, resulting in an XY-stage rate of 250 µm/s.

Spray stability was assessed through multiple tests, including solvent stability, ink measurement, and a 30-minute acquisition on test tissue, followed by evaluation of the TIC heatmap. The ink intensity exceeded 5 × 10^6^, passing the sensitivity control.

#### timsTOF fleX MALDI-2 at UK Münster

MALDI-MSI imaging was performed on a timsTOF fleX MALDI-2 mass spectrometer (Bruker Daltonik, Bremen, Germany), using a 13 × 13 µm beam scan range, resulting in a 17 × 17 µm pixel size. General instrumental parameters were identical to measurements using the timsTOF fleX MALDI-2 at KU Leuven; deviations in settings are further specified in the **Supplementary Table 3.**

#### MALDI-Q Exactive^TM^ Orbitrap at Münster

The MALDI-MSI imaging was performed on a Q Exactive^TM^ Plus Hybrid Quadrupole-Orbitrap mass spectrometer (Thermo Fisher Scientific, Bremen, Germany), similar to that described in ^25^, equipped with a dual ion funnel source (MALDI/ESI injector; Spectroglyph, Kennewick, WA). The source was operated at an N2 buffer gas pressure of 5.5 mbar. The stage was controlled using the MALDI Injector software (Spectroglyph). The data was acquired in positive ion mode, using a full MS scan. The *m/z* range was set to 300-1200 with a mass resolution (FWHM) of 140 000 at *m/z* 200. The maximum injection time was set to 400 ms. MALDI imaging was performed with a pixel size of 20 x 20 µm using a frequency-tripled (349 nm) Nd:YLF laser (Explorer ICT-349-120-E, Spectra-Physics, Stahnsdorf, Germany) with a focal beam diameter of 10 µm. The laser pulse repetition rate was 100 Hz, and the laser pulse energy exciting the laser was 1.7 µJ. The .imzML file was created by combining the raw file with the position file produced, using the image inside (spectroglyph).

### Quantitative bulk lipidomics

#### HILIC-MS/MS

Lipids extracted from LCM-derived material were analyzed using hydrophilic interaction chromatography (HILIC) on a Nexera X2 UHPLC system (Shimadzu) coupled online with a hybrid triple quadrupole/linear ion trap mass spectrometer (6500+ QTRAP system; AB SCIEX). Chromatographic separation was performed on a XBridge amide HILIC column (150 mm × 4.6 mm, 3.5 μm; Waters) maintained at 35°C using mobile phases A (1 mM ammonium acetate in water-acetonitrile 5:95 (*v/v*)) and B (1 mM ammonium acetate in water-acetonitrile 50:50 (*v/v*)) in the following gradient: 0-60 min: 0 % B → 6 % B; 6-10 min: 6 % B → 25 % B; 10-11 min: 25 % B → 98 % B; 11-130 min: 98 % B → 100 % B; 13-190 min: 100 % B; 19-240 min: 0 % B at a flow rate of 0.70 ml/min which was increased to 1.50 mL/min from 13 minutes onwards. The instrument parameters were as follows: Curtain Gas = 35 psi; Collision Gas = 8 a.u. (medium); IonSpray Voltage = 5500 V and -4500 V; Temperature = 550°C; Ion Source Gas 1 = 50 psi; Ion Source Gas 2 = 60 psi; Declustering Potential = 60 V and -80 V; Entrance Potential = 10 V and -10 V; Collision Cell Exit Potential = 15 V and -15 V. Lipid detection was performed using scheduled multiple reaction monitoring (MRM), with transitions based on neutral losses or the typical product ions. SM, CE, Cer, DCer, HexCer, and Hex2Cer were measured in positive ion mode by detection a product ion at *m/z* 184.1, 369.4, 264.4, 266.4, 264.4, or 264.4, respectively. TG, DG, and MG were measured in positive ion mode by monitoring product ions corresponding to the neutral loss for one of the fatty acyl moieties. PC, LPC, PE, LPE, PG, PI, and PS were measured in negative ion mode by detecting the fragmentation ions of the corresponding fatty acyls. The following fatty acyl moieties were taken into account for the lipidomic analysis: 14:0, 14:1, 16:0, 16:1, 16:2, 18:0, 18:1, 18:2, 18:3, 20:0, 20:1, 20:2, 20:3, 20:4, 20:5, 22:0, 22:1, 22:2, 22:4, 22:5 and 22:6 except for TGs which considered: 16:0, 16:1, 18:0, 18:1, 18:2, 18:3, 20:3, 20:4, 20:5, 22:2, 22:3, 22:4, 22:5, 22:6.

### MSI data processing in LipidQMap software

All raw data acquired on the timsTOF fleX MALDI-2 systems were converted into .imzML files using SCiLS Lab software (v2025b Pro). The centroided data were processed with a standard bin size of 5 mDa at 1000 *m/z*, and the *m/z* range was automatically determined from the measurements. Raw DESI MRT Select Series data (.RAW) were converted to the .imzML format by using HDI version 1.7 (Waters).

All data were uploaded as .imzML files into LipidQMap. Mass accuracy was set to 10 ppm, and the bin size for aligning centroided data was 5 mDa. Type II isobaric overlap correction was applied to all measurements, except for the DESI MRT data, considering MRT resolution of 200 000. Na/H adduct isobaric correction was applied in all cases except the Type II isobaric correction experiments (sections were washed). The lock-mass-based recalibration was applied to the raw centroided data using the PC 34:1 [M+K]^+^ adduct (or [M+H]^+^ adduct for the section washed with ammonium formate and DESI-qMSI), detected in the majority of pixels, with a peak assignment tolerance of 30 ppm and a minimum intensity of 10 000 to perform recalibration. Processed data were subjected to Gaussian filtering using a winsorizing percentile of 95% to 98% for raw, isotope-corrected, and qMSI images.

Peak integration from LC-MS/MS MRM data was performed using MultiQuant^TM^ software (v3.0.3). Lipid signals were corrected for isotopic contributions (calculated using Python Molmass v2019.1.1) and quantified based on IS signals.

#### Lipid shorthand annotation

Lipid structures were annotated according to the shorthand notation proposed on the LIPID MAPS classification system ^26,27^.

### Development of LipidQMap software in Python

The LipidQMap software was developed in Python (version 3.12) to provide a comprehensive, cross-platform GUI for the processing and quantitative analysis of MSI data. The application’s front-end was constructed using the Qt framework via its Python binding, PySide6 (version 6.7). The core data processing and numerical computations rely on a suite of scientific libraries, including NumPy, pandas for database processing, and molmass for calculating monoisotopic masses from chemical formulas. The core functionality of LipidQMap, including online calibration, deisotoping, and quantification algorithms, was independently developed without relying on third-party implementations. To ensure correctness, all code parts related to data analysis were thoroughly unit-tested and validated against manual calculations.

Data input and management

LipidQMap processes MSI data stored in the standardized .imzML format. To handle data import, we utilized the pyimzml library, initially developed by the team of Theodore Alexandrov [https://github.com/alexandrovteam/pyimzML]. In contrast to the original pyimzML implementation, which reads binary data files during each call to getspectrum, LipidQMap reads all spectral data upon file opening, subsequently closing the connection to the data file. When applied to a 5-gigabyte MSI benchmark dataset of a mouse brain imaged at 30 µm (containing about 59 000 spectra, with an average of 7400 *m/z* features), this optimization resulted in a two-fold speedup in extracting 2500 ion images. Further performance gains were achieved by replacing Python’s standard library bisect module for binary search with NumPy’s searchsorted function for ion image extraction. This allowed single vectorized operations across all required *m/z* values, eliminating the need for iterative loops and yielding a 10-fold speed increase on our benchmark dataset. Combined with the initial optimization, a 160-fold speed improvement was observed. The integration of the Numba just-in-time (JIT) compiler and the substitution of NumPy functions with Numba-compatible alternatives provided an additional 5-fold speedup, culminating in an overall 820-fold increase in data processing speed, which reduced computation time from hours to seconds.

A user-editable lipid database, stored as a Microsoft Excel (.xlsx) file, serves as the primary source for lipid species annotation (**Supplementary Table 4**). At runtime, this file is parsed using the pandas library. Each row in the database corresponds to a lipid species and must contain the following columns: ID, Class, Neutral formula, Adducts, M-2 Isotope, Na^+^ Isotope, Is standard (is a lipid a standard), IS (ID of the IS that should be used to quantify this species), and a standard amount (pmol/mm^2^). The software uses the molmass library to calculate the precise monoisotopic mass for each species and its specified adducts.

#### Data processing pipeline

The core functionality of LipidQMap is structured as a sequential data processing pipeline, which performs:

##### 1. Mass recalibration

An optional mass recalibration step can be applied to each spectrum. This process identifies a user-specified reference *m/z* within a defined tolerance (in parts per million, ppm) and a minimum intensity. If the peak is found, the *m/z* axis of the entire spectrum is shifted to align the reference peak with its theoretical *m/z* value, correcting for systematic mass drift.

##### 2. Ion image generation

For each lipid species defined in the database, a two-dimensional ion image is generated. The intensity for each pixel is determined by extracting the maximum signal within a user-defined mass tolerance (ppm) centered on the calculated *m/z* of the target lipid. This extraction is performed by a Numba-accelerated function for optimal processing speed.

##### 3. Isotopic correction

LipidQMap implements two isotopic correction algorithms to address common isobaric interferences: correction for X:Y [M+Na]^+^ on X+2:Y+3 [M+H]^+^ overlap (X number of carbon atoms and Y number of DB in their acyl chains) and Type II isobaric overlap correction.

##### 3a. Na/H adducts isobaric correction algorithm

To address Na/H adduct isobaric overlap, LipidQMap adapts the correction method developed by Höring et al. ^15^, initially designed for processing shotgun lipidomics spectra, thereby extending its applicability to MSI in a pixel-wise manner across the dataset. It relies on the ratio (calculated per lipid class) of sodiated to protonated signals (*RA^Na^*) of *n* standards, which can be presented as:

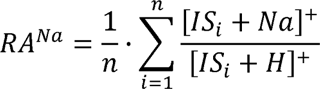

The *[IS_i_+Na]^+^* and *[IS_i_+H]^+^* correspond to the intensities of sodiated and protonated adducts of lipid class standards, respectively. The following formula presents the correction:

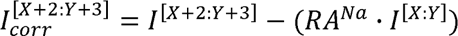

The *I*[X+2:Y+3] is the intensity of overlapped isobaric signals, and *I*[X:Y] is the intensity of the protonated lipid signal ^15^. Information on the necessary correction is provided in the lipid identification databases, containing the adduct formula responsible for the isobaric overlap. This correction precedes the Type II isobaric overlap correction.

### 3b. Type II isobaric correction algorithm

In the current release, LipidQMap applies a standard Type II correction algorithm based on the predicted isotopic distribution of signals, incorporating adjustments for M+2 isotopologues in the protonated ions. Type II isotopic overlap, where species [X:Y] overlap with the M+2 isotopologue of species [X:Y+1], was corrected using the following algorithm:

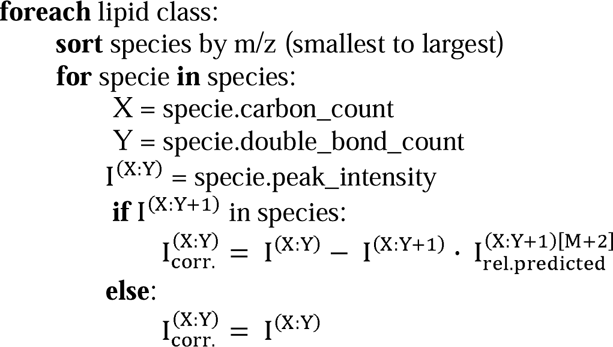

Where *I* is peak intensity and *I rel.predicted*. is the predicted relative intensity of the respective isotopologue, based on the natural isotopic distribution.

#### 4. Lipid quantitation

LipidQMap utilizes a widely adopted one-point calibration approach. This involves relating the signal intensity of the analyte to that of a selected lipid class standard within each individual spectrum (i.e. per pixel). This relationship is defined by the following formula:

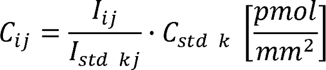

Where *C_ij_* is the concentration of the *i*-th lipid in the *j*-th pixel, *I_ij_* is the intensity of the *i*-th lipid in the *j*-th pixel, *I_std kj_* is the intensity of the *k*-th standard in the *j*-th pixel, and *C_std k_* is the concentration of the *k*-th lipid standard applied to the tissue surface. The delivered amount of standard is normalized against the area sprayed, based on the information provided by the sprayer, and ultimately expressed as pmol/mm^2^.

#### 5. Post-processing and visualization

To handle pixels with no detectable signal (i.e., NaN values), an optional median imputation can be performed, where each missing value is replaced by the median of its 3×3 neighboring pixels. To mitigate the effect of high-intensity outliers, pixel intensities in the final images can be winsorized. The user can set an upper percentile threshold, above which all intensity values are capped at the value corresponding to that percentile.

##### Integrated standard concentration calculator

LipidQMap includes an integrated standard concentration calculator. This tool provides a guided workflow for determining the surface concentration (in pmol/mm²) of an IS applied to a sample surface via automated spraying, a critical normalization factor for quantitative analysis. The calculator directly interfaces with the user’s lipid databases, allowing for the calculated value to be saved for the selected standard. The solution was tested using the HTX M5 sprayer.

The calculation is based on a multi-step model that accounts for key experimental parameters, divided into four logical stages within the user interface:

### Sprayed Area Calculation

The total surface area (*A*, in mm²) is determined from user-provided sprayer coordinates (*x_left_*, *x_right_*, *y_bottom_*, *y_top_*) and an additional margin (mm), according to the formula:

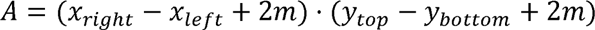

### Effective Sprayed Volume Calculation

The calculator computes the net volume of solution delivered to the target area (*V_delivered_*, in ml). This is derived from the total volume consumed from the syringe (*V_total_*), the syringe pump flow rate (*F_syringe_*, in ml/min), the duration of initial system equilibration (*t_equilibration_*, in min), the drying time between spray cycles (*t_drying_*, in min), and the number of drying cycles (*n_cycles_*). The volume lost during non-spraying periods is subtracted from the total volume used:

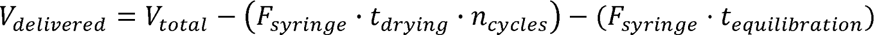

### Spraying Solution Concentration Calculation

The molar concentration of the IS in the final spraying mixture (*C_mix_*, in pmol/ml) is calculated from the initial stock concentration (*C_stock_*, in mg/ml), a working dilution factor (*DF_working_*), the volume of the working stock used (*V_working_*, in µl), the final mixture volume (*V_final_mix_*, in ml), and the molecular weight of the standard (MW, in g/mol or µg/µmol):

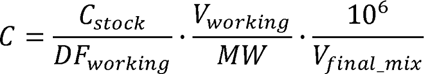

### Final Surface Concentration Calculation

The final surface concentration (*C_surface_*, in pmol/mm²) is computed by combining the values from the previous steps, representing the total amount of IS delivered per unit area:

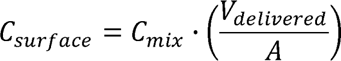

Upon calculation, the user can apply this value to any standard listed in the selected database. This action populates the IS amount (pmol/mm^2^) field for the chosen species. Saving these changes updates the underlying database Excel file directly, ensuring that the new, accurately determined normalization factor is persistently stored and available for subsequent quantitative analyses within LipidQMap. This integrated approach minimizes manual calculation errors and streamlines the workflow from experimental setup to data processing.

## Code availability

LipidQMap is an open-source software. The installation wizard with instruction files can be accessed at: https://github.com/swinnenteam/LipidQMap.

## Data availability

Access to raw mass spectrometry imaging data: https://doi.org/10.6084/m9.figshare.30359338.v1

## Supplementary Tables

Supplementary Table 1: Composition of internal standard mixtures. Supplementary Table 2: HTX Sprayer settings. Supplementary Table 3: timsTOF fleX MALDI-2 mass spectrometers’ settings. Supplementary Table 4: LipidQMap lipid database example.

## Supplementary Data

Additional data of qMSI analyses processed by LipidQMap and validated by quantitative bulk lipidomics.

## Acknowledgments

J.D. is supported by a KU Leuven Core Facility Incubation and Translational grant. J.I. is supported by the Leuven Future Fund LISCO-BIOMED, and C.P.M. by the FWO 11F1423N and KOTK13897 grants. M.G. is supported by a HORIZON TMA MSCA Postdoctoral Fellowship (European Fellowship, 101152131). This work was supported by an FWO-SBO grant LIPOMACS (S001623N), a Focus Group grant from the Leuven Cancer Institute, the KU Leuven Opening the Future Campaign, and the KU Leuven Fund Dieter de Cauderlier for Research on Brain Tumor. Work in the Fedorova lab is supported by “Sonderzuweisung zur Unterstützung profilbestimmender Struktureinheiten” by the SMWK to TUD, TG70 by Sächsische Aufbaubank and SMWK, the measure is co-financed with tax funds on the basis of the budget passed by the Saxon state parliament (to M.F.), Deutsche Forschungsgemeinschaft (FE 1236/5-1, FE 1236/8-1 to M.F.), and Bundesministerium für Bildung und Forschung (031L0315A, DEEP_HCC and 01EJ2205A, FERROPath to M.F.). K.D. and Jens S. acknowledge German Research Foundation (DFG, project no. 326945247 to K.D. and J.S.) and Bruker Daltonics for financial and technical support.

## Author Contributions Statement

J.I. and J.D. had the original idea for this project. J.I., J.D., N.R., and J.V.S. prepared the original draft of the manuscript. J.I., J.D., and N.R. prepared the figures. J.D. developed the software. J.I., N.R., J.D., M.W., J.S., and Jens S. tested and validated the software. N.R., C.P.M., F.V., A.T., S.G., and V.D.L. collected and prepared the biological material. N.R., J.I., F.V., A.T., S.G., and X.S. prepared the samples for mass spectrometry analysis. N.R., J.I., J.D., M.W., M.G., J.S., Jens S., A.T., S.G., X.S., performed the mass spectrometry measurements. J.I., N.R., J.D., M.W., M.G., J.S., and Jens S. performed the data processing. The supervision was performed by J.V.S., M.F., Jens S., M.H., R.J., B.G., G.B., and K.D. All coauthors contributed to the manuscript review and editing and approved the final version of the manuscript. J.V.S. and J.I. were responsible for project administration.

