## Supplementary Data for "LipidQMap - An Open-Source Tool for Quantitative Mass Spectrometry Imaging of Lipids"

**Electronic supplementary material to the preprint:** Quantitative mass spectrometry imaging (qMSI) analysis of a sagittal healthy mouse brain section in positive ion mode with corresponding laser-capture microdissection-based quantitative bulk lipidomics.

### Sodium correction – PC example: 1

#### PC 35:1[$D_5$] / PC 32:0[$D_9$] normalization

LCM
(bulk via HILIC-MS/MS
MRM transitions based on fatty acyl losses)

Overlap of PC 34:1 $[M+Na]^+$ and PC 36:4 $[M+H]^+$

Raw image:

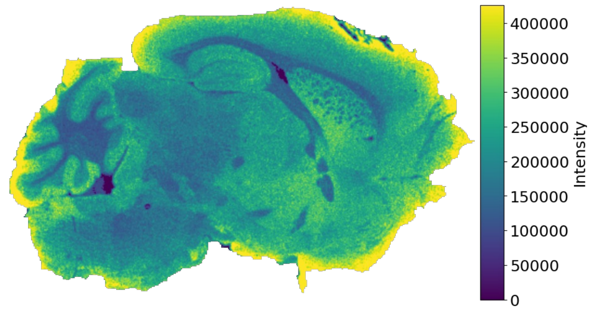


LCM ROIs:

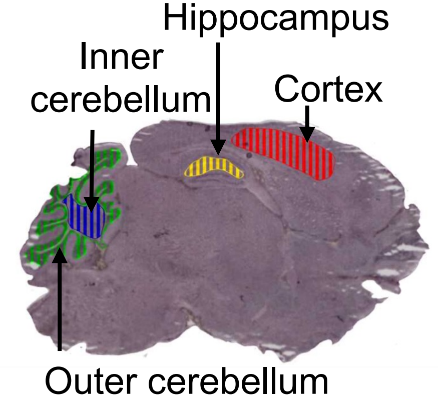


**LipidQMap**

**SCiLS**

Deisotoped image

qMSI image

TIMS image (Normalized to standard)

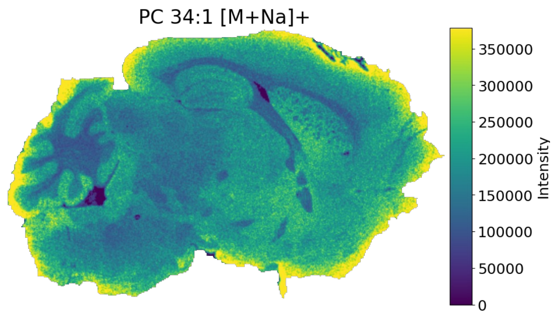


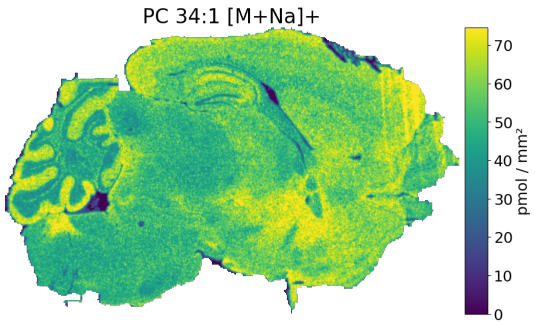


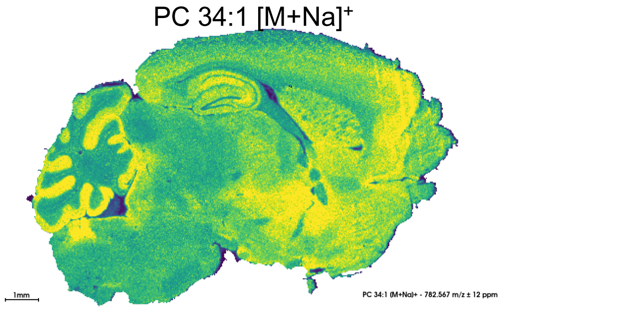


m/z 782.5670±12ppm 292.5±2.5Å

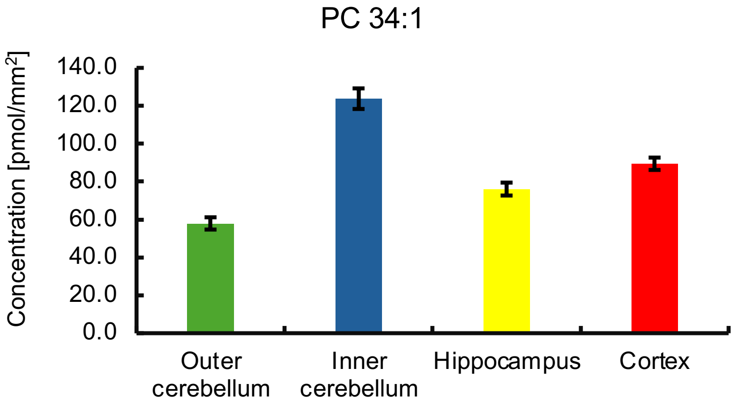


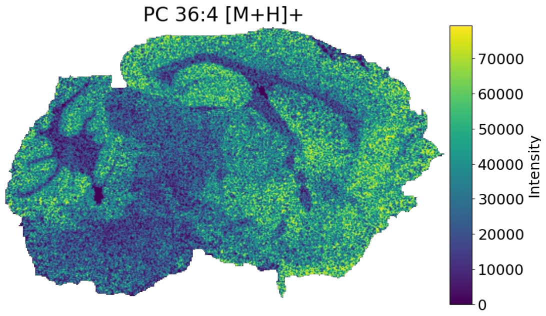


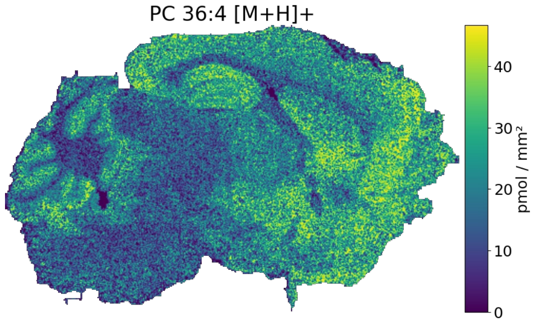


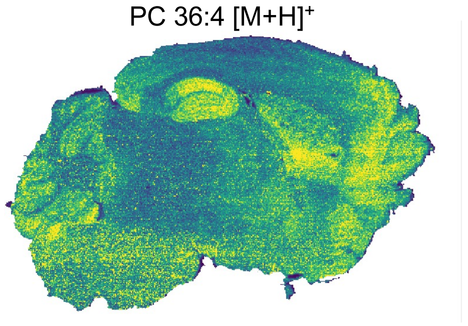


m/z 782.5691±10ppm 289.1±2.0Å²

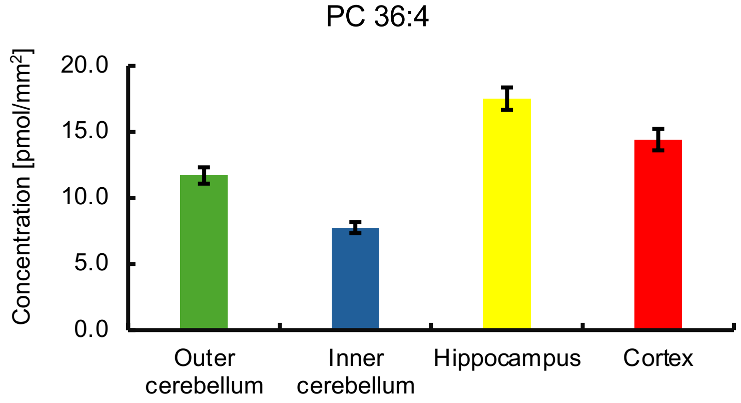

### Sodium correction – PC example: 2

#### PC 35:1[$D_5$] normalization

Overlap of PC 36:1 $[M+Na]^+$ and PC 38:4 $[M+H]^+$

Raw image:

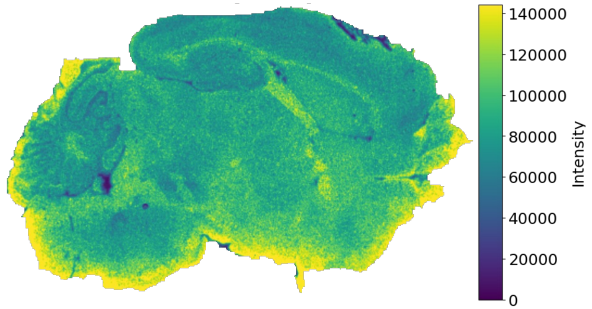


LCM
(bulk via HILIC-MS/MS
MRM transitions based on fatty acyl losses)

LCM ROIs:

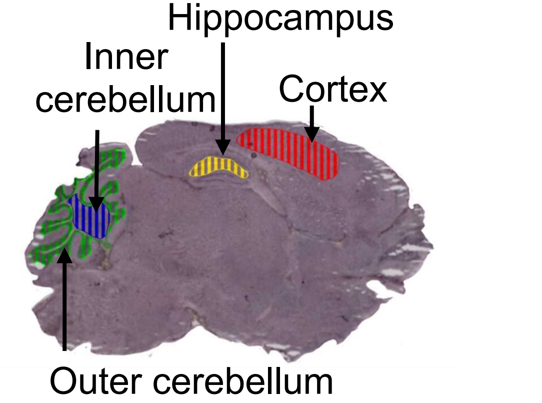


**LipidQMap**

Deisotoped image

qMSI image

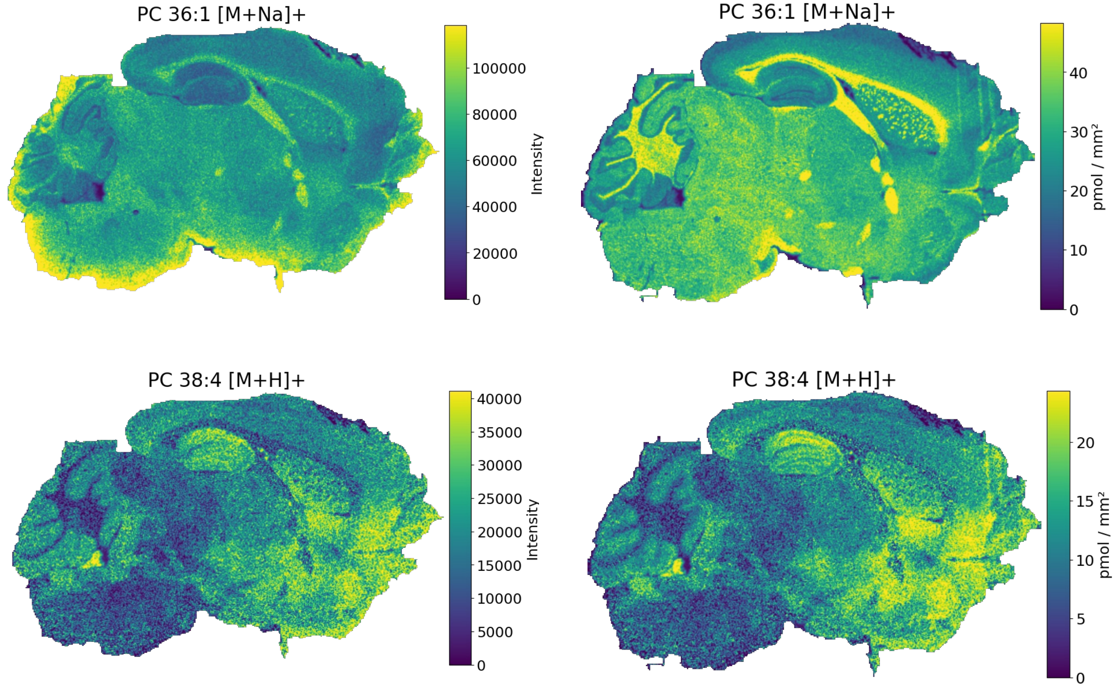


**SCiLS**

TIMS image (Normalized to standard)

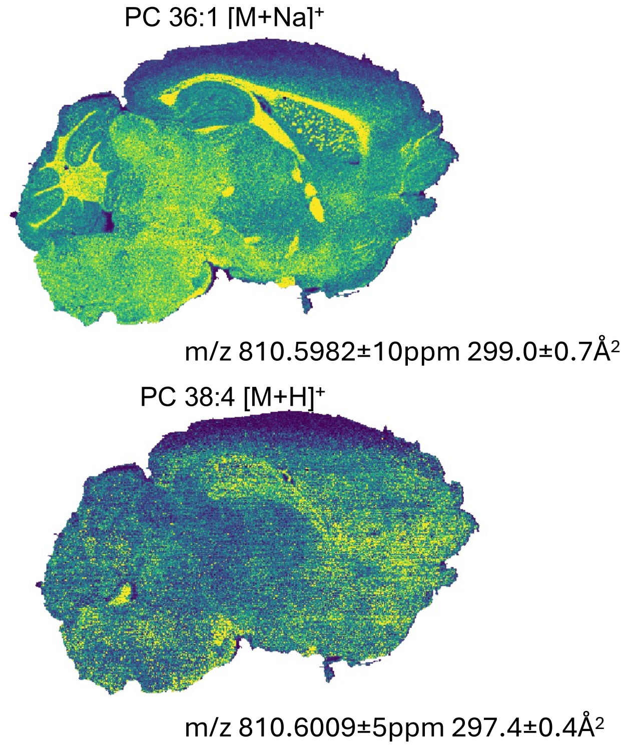


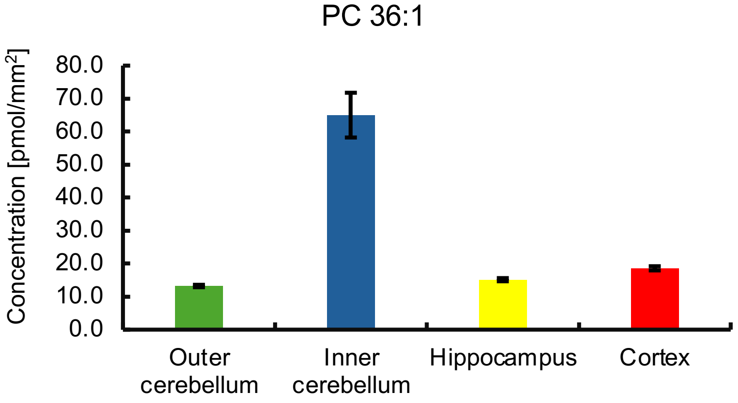


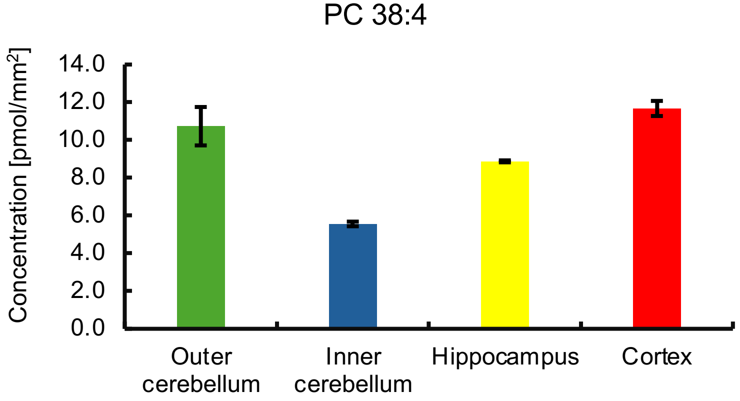

### Sodium correction – PE example 1

#### PE 33:1[$D_7$] normalization

Overlap of PE 36:1 $[M+Na]^+$ and PE 38:4 $[M+H]^+$

Raw image:

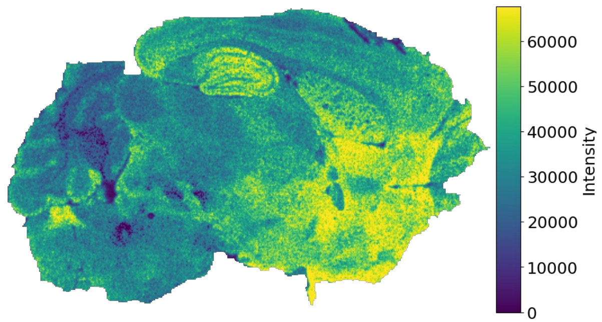


LCM
(bulk via HILIC-MS/MS
MRM transitions based on fatty acyl losses)

LCM ROIs:

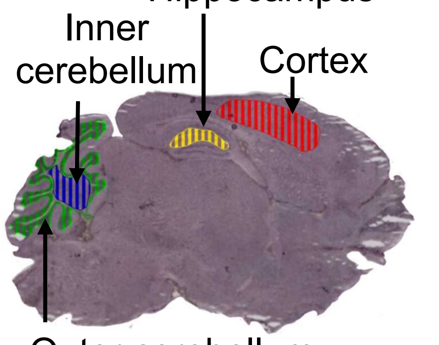


**LipidQMap**

**SCiLS**

Deisotoped image

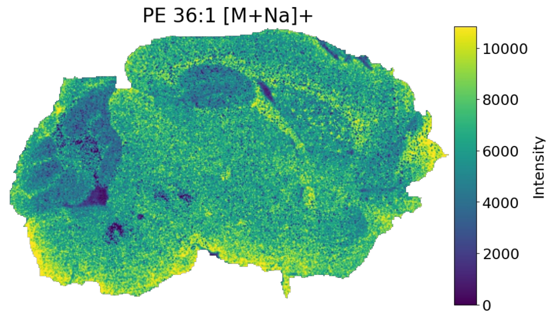


qMSI image

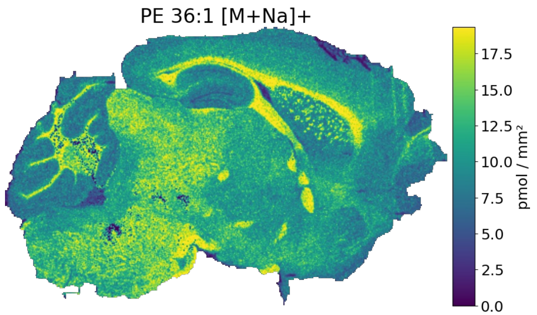


TIMS image (Normalized to standard)

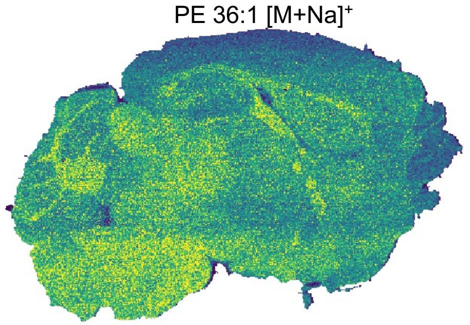


m/z 768.5521±10ppm 288.9±1.0Å²

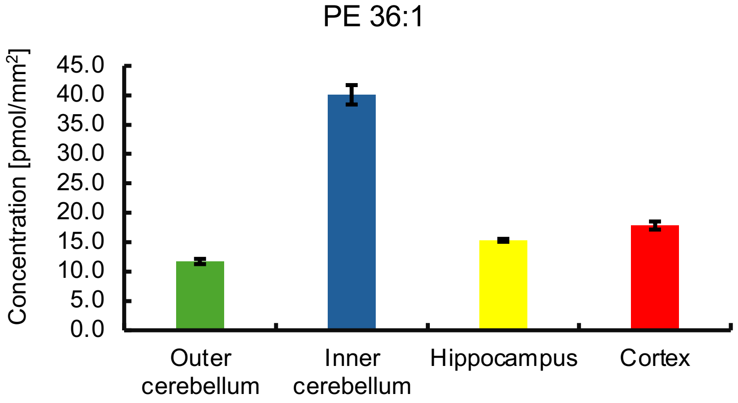


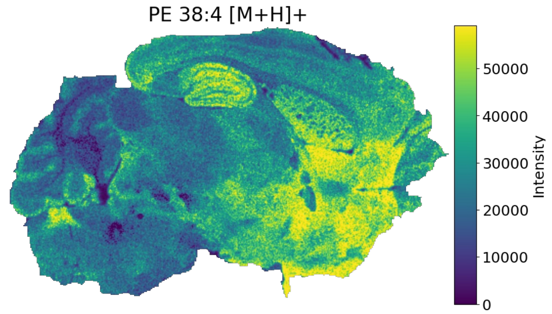


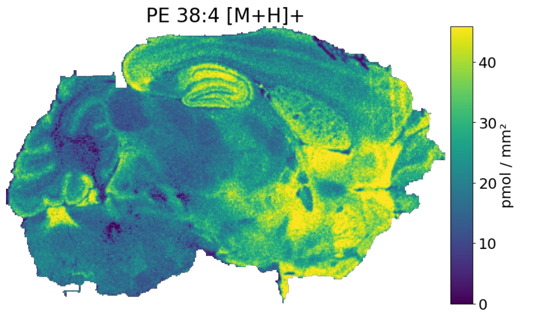


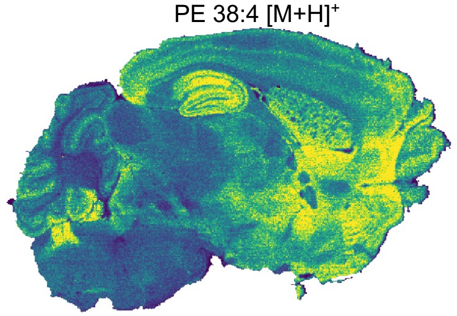


m/z 768.5545±10ppm 284.4±2.0Å²

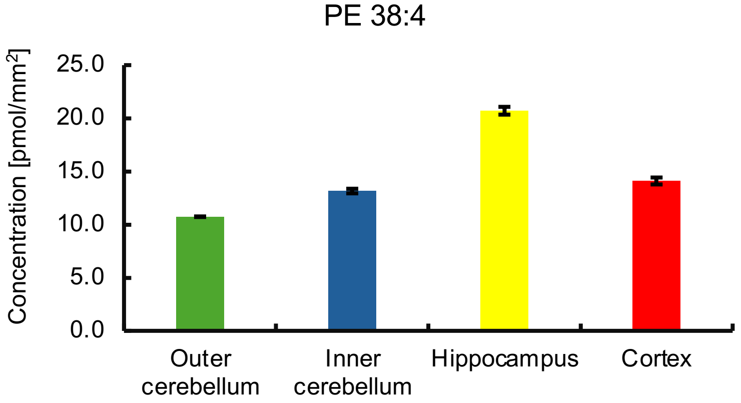

### Sodium correction – PE example 2

#### PE 33:1[$D_7$] normalization

Overlap of PE 38:1 $[M+Na]^+$ and PE 40:4 $[M+H]^+$

Raw image:

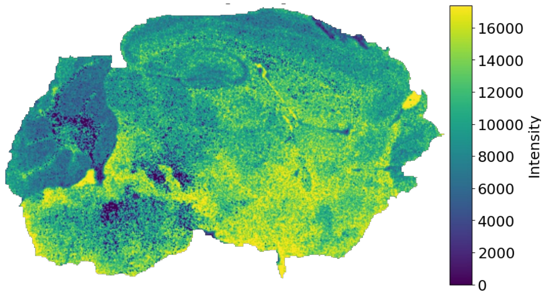


**LipidQMap**

Deisotoped image

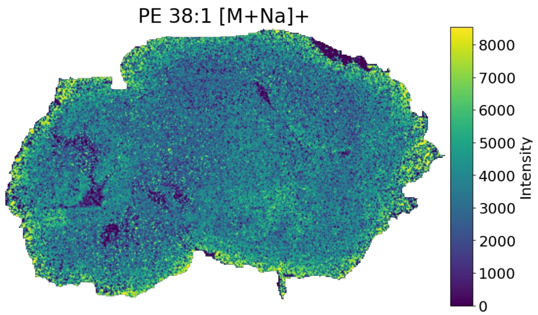


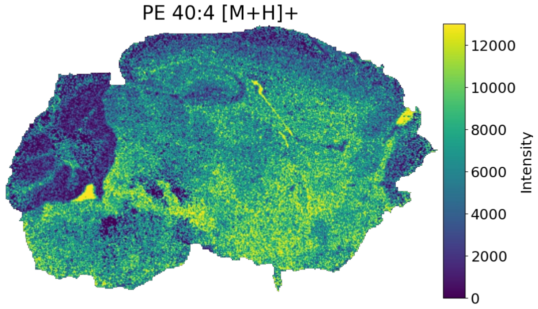


qMSI image

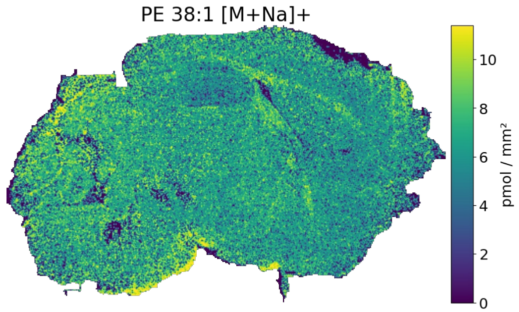


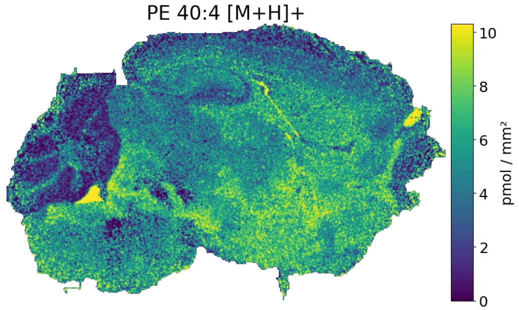


LCM
(bulk via HILIC-MS/MS
MRM transitions based on fatty acyl losses)

LCM ROIs:

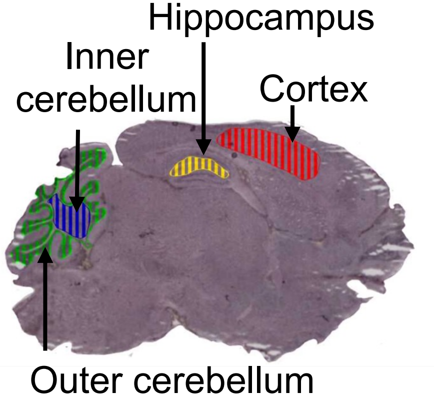


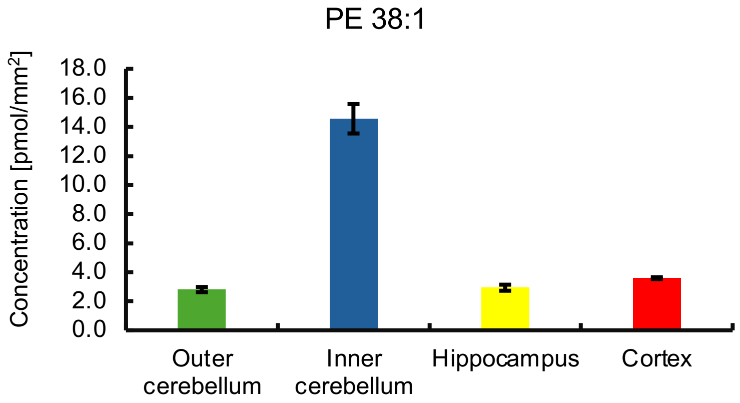


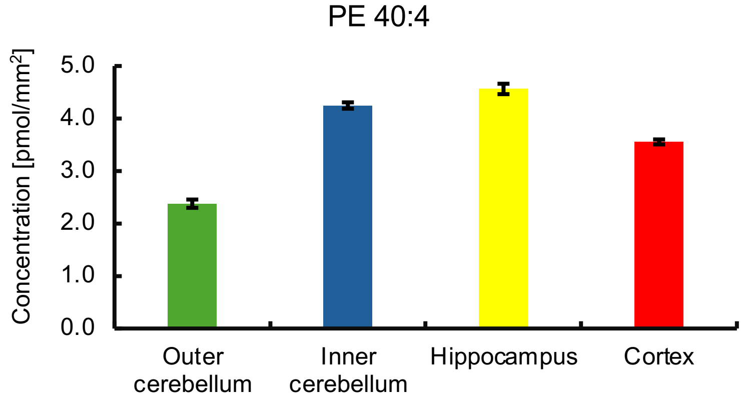

### Sodium correction – PE O-/PE P- example 1

#### PE 33:1[$D_7$] normalization

Overlap of PE O-36:4 / PE P-36:3 $[M+Na]^+$ and PE O-38:7 / PE P-38:6 $[M+H]^+$

Raw image:

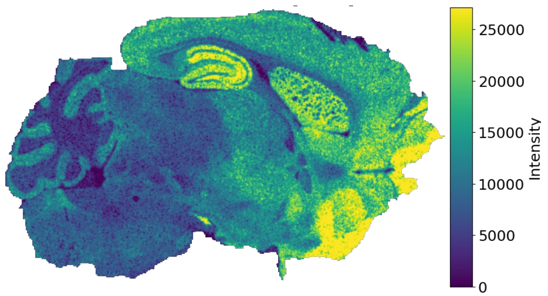


**LipidQMap**

Deisotoped image | qMSI image

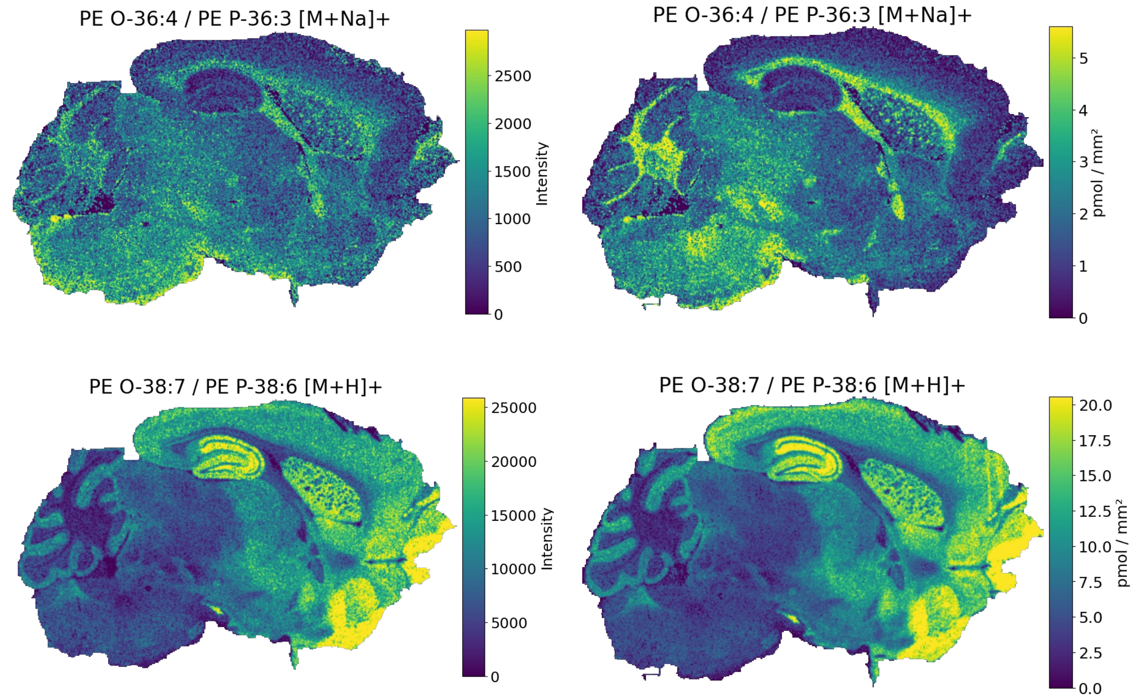


LCM
(bulk via HILIC-MS/MS
MRM transitions based on fatty acyl losses)

LCM ROIs:

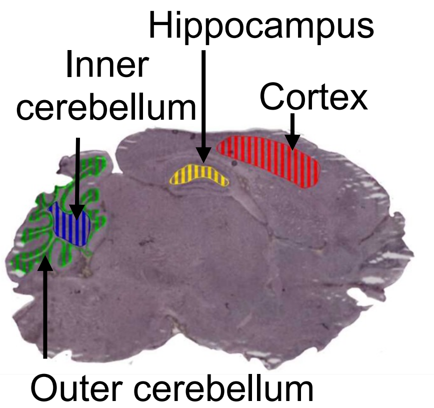


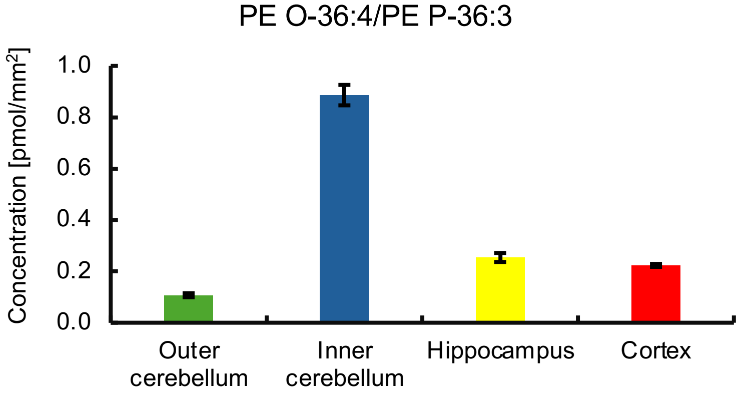


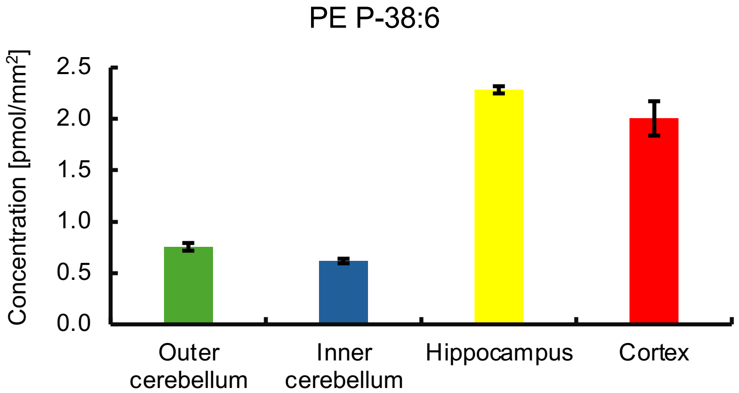

### Sodium correction – PE O-/PE P- example 2

#### PE 33:1[$D_7$] normalization

Overlap of PE O-38:2 / PE P-38:1 $[M+Na]^+$ and PE O-40:5 / PE P-40:4 $[M+H]^+$

Raw image:

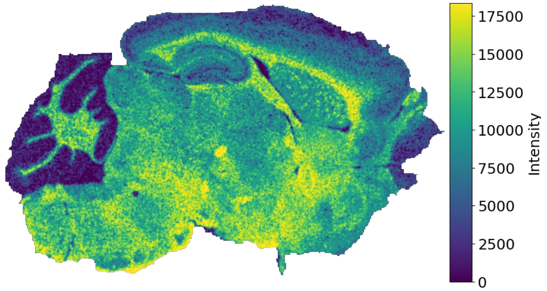


LCM
(bulk via HILIC-MS/MS
MRM transitions based on fatty acyl losses)

LCM ROIs:

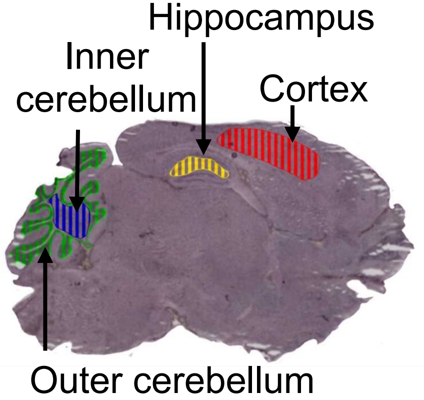


**LipidQMap**

Deisotoped image

qMSI image

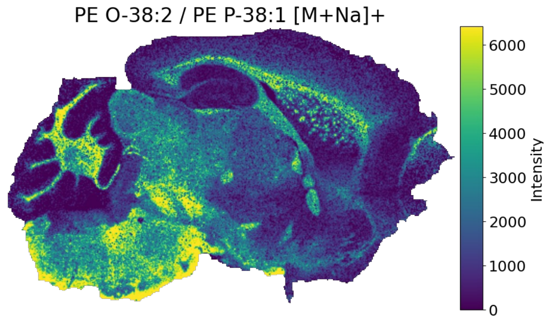


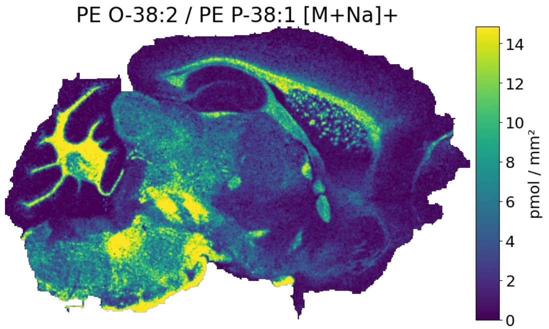


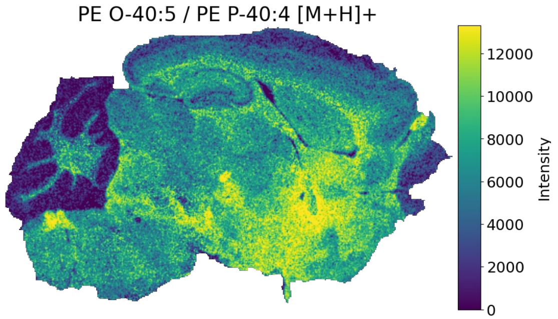


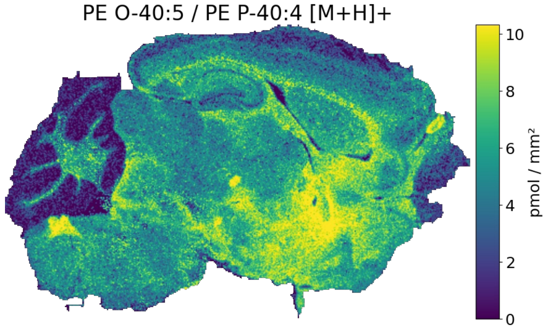


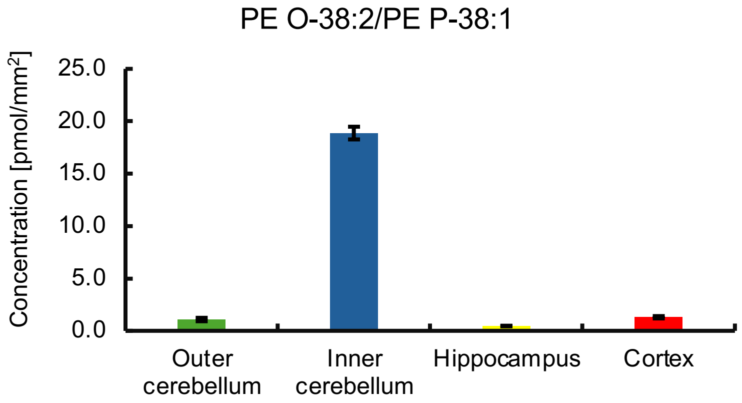


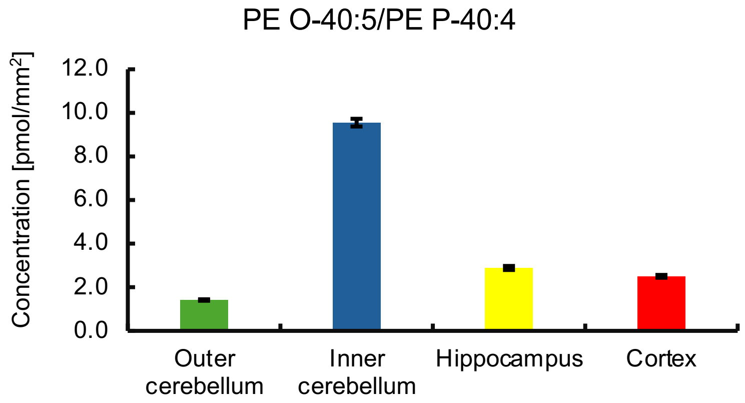

### Sodium correction – PE O-/PE P- example 3

#### PE 33:1[$D_7$] normalization

Overlap of PE O-34:1 / PE P-34:0 $[M+Na]^+$ and PE O-36:4 / PE P-36:3 $[M+H]^+$

Raw image:

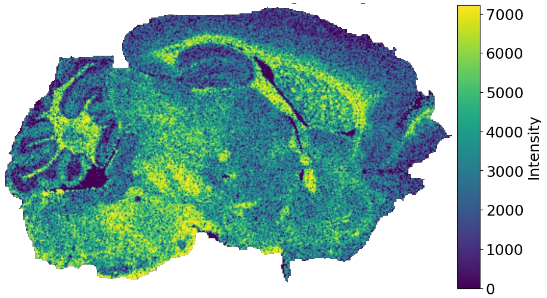


LCM
(bulk via HILIC-MS/MS
MRM transitions based on fatty acyl losses)

LCM ROIs:

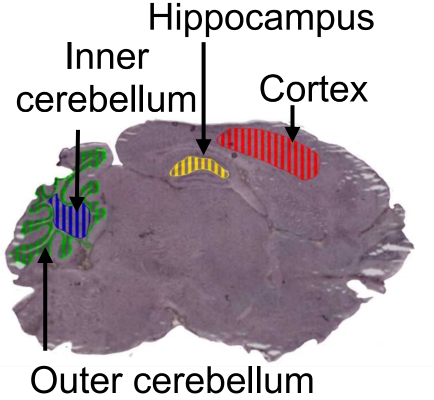


**LipidQMap**

Deisotoped image

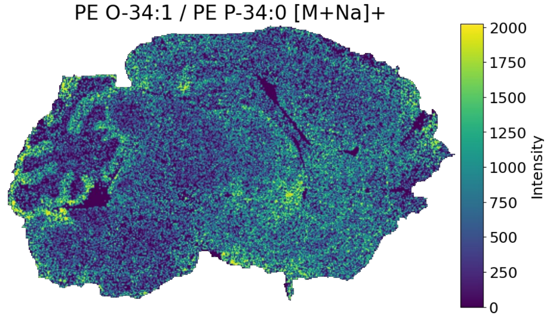


qMSI image

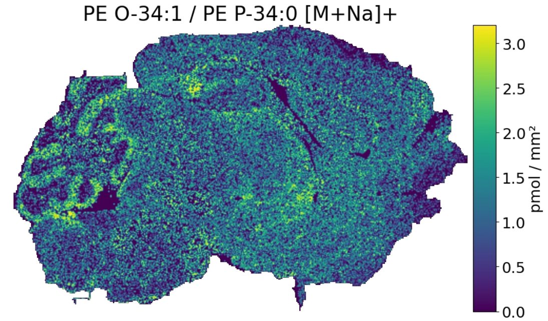


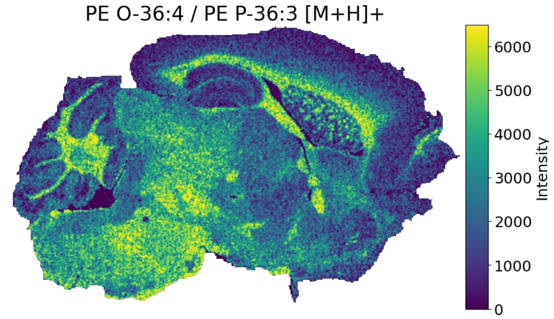


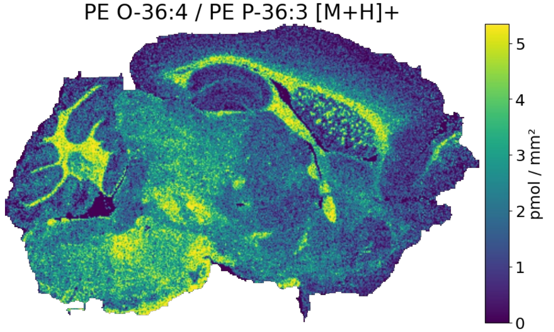


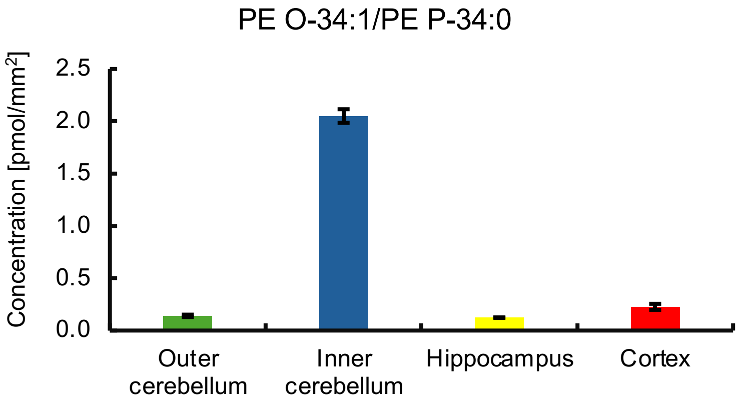


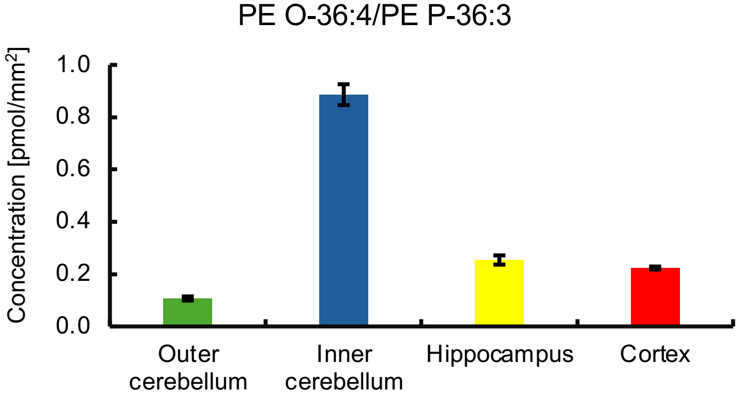

### Sodium correction – PE O-/PE P- example 3

#### PE 33:1[$D_7$] normalization

Overlap of PE O-36:2 / PE P-36:1 $[M+Na]^+$ and PE O-38:5 / PE P-38:4 $[M+H]^+$

Raw image:

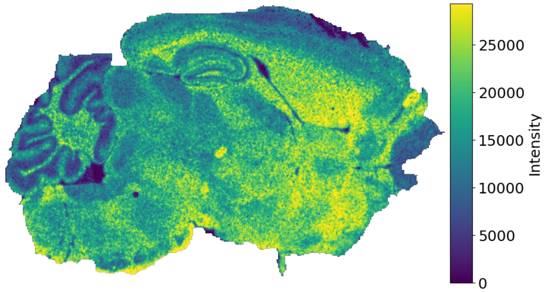


**LipidQMap**

Deisotoped image

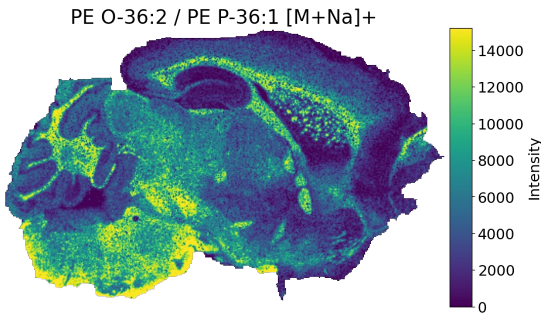


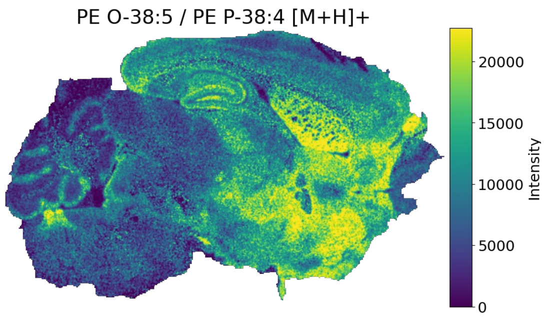


qMSI image

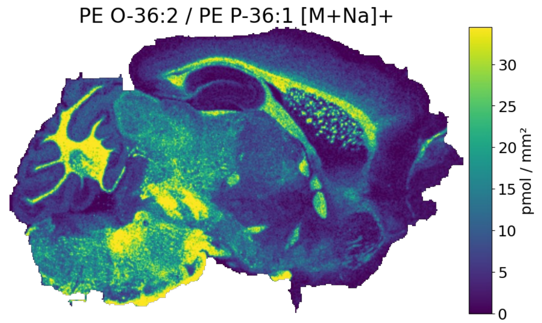


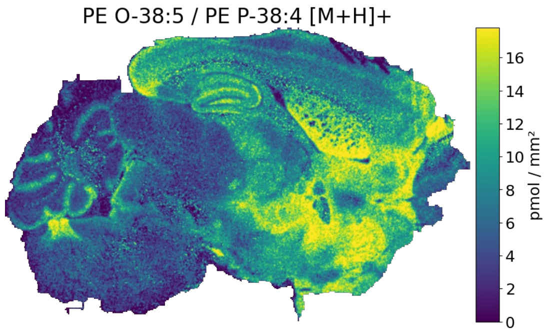


LCM
(bulk via HILIC-MS/MS
MRM transitions based on fatty acyl losses)

LCM ROIs:

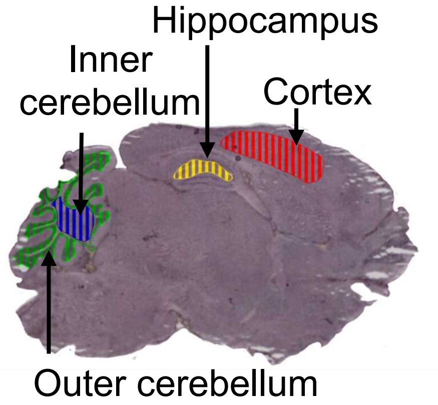


PE O-36:2/PE P-36:1

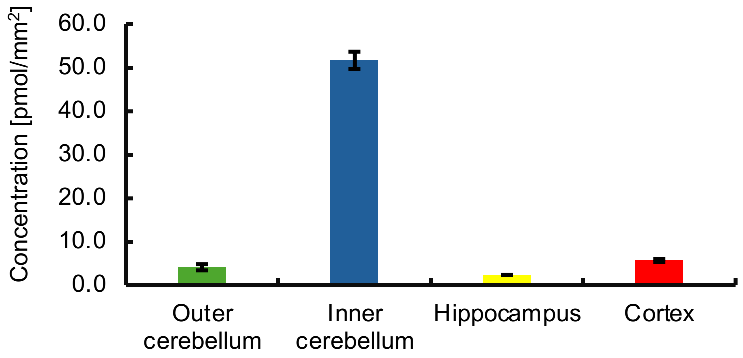


PE O-38:5/PE P-38:4

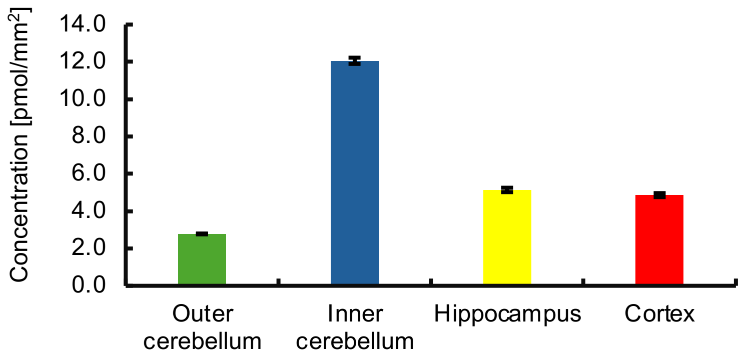

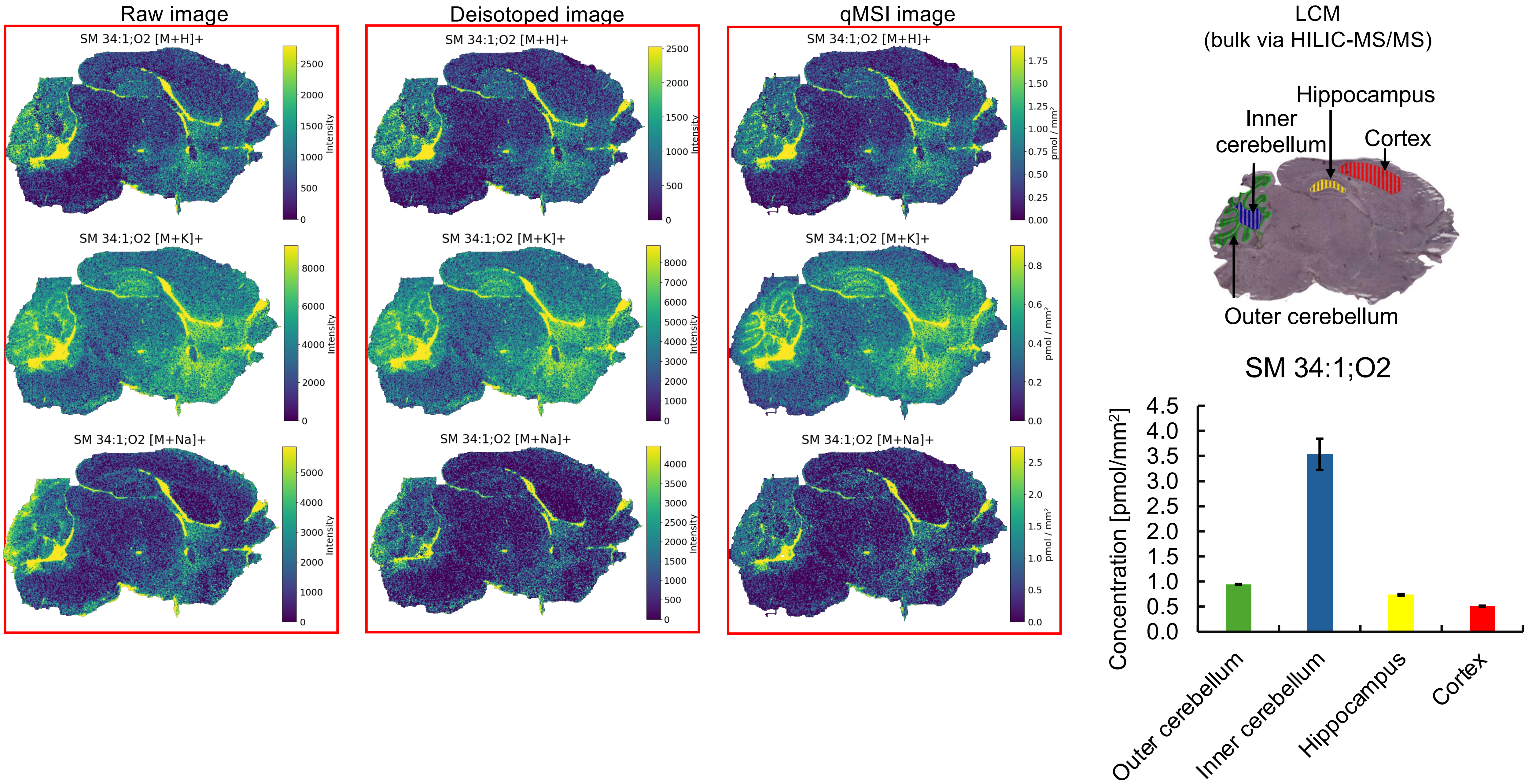


* Missing pixels for **[M+H]⁺, [M+Na]⁺** adducts (profile to centroid transformation).

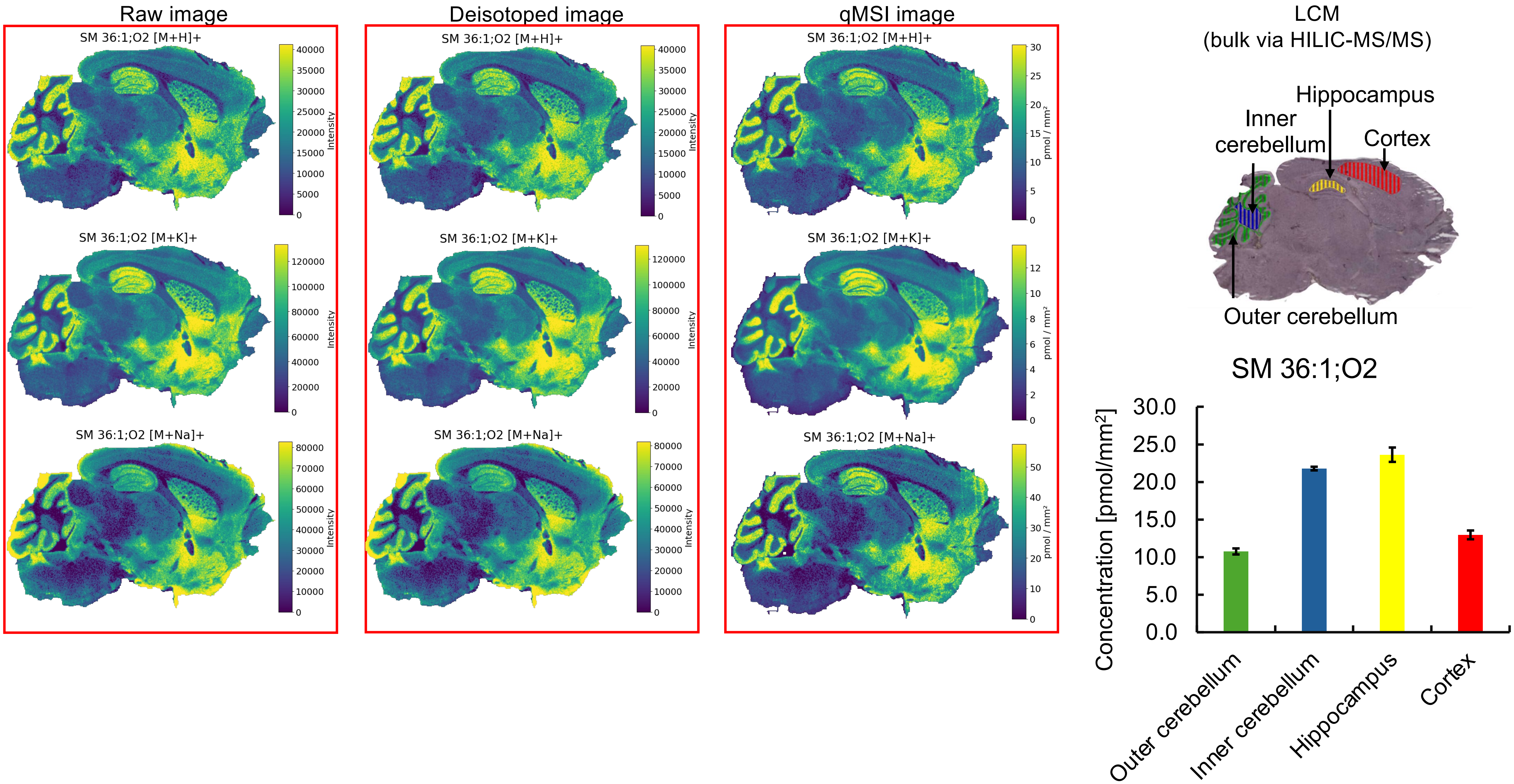


* Missing pixels for **[M+H]⁺, [M+Na]⁺** adducts (profile to centroid transformation).

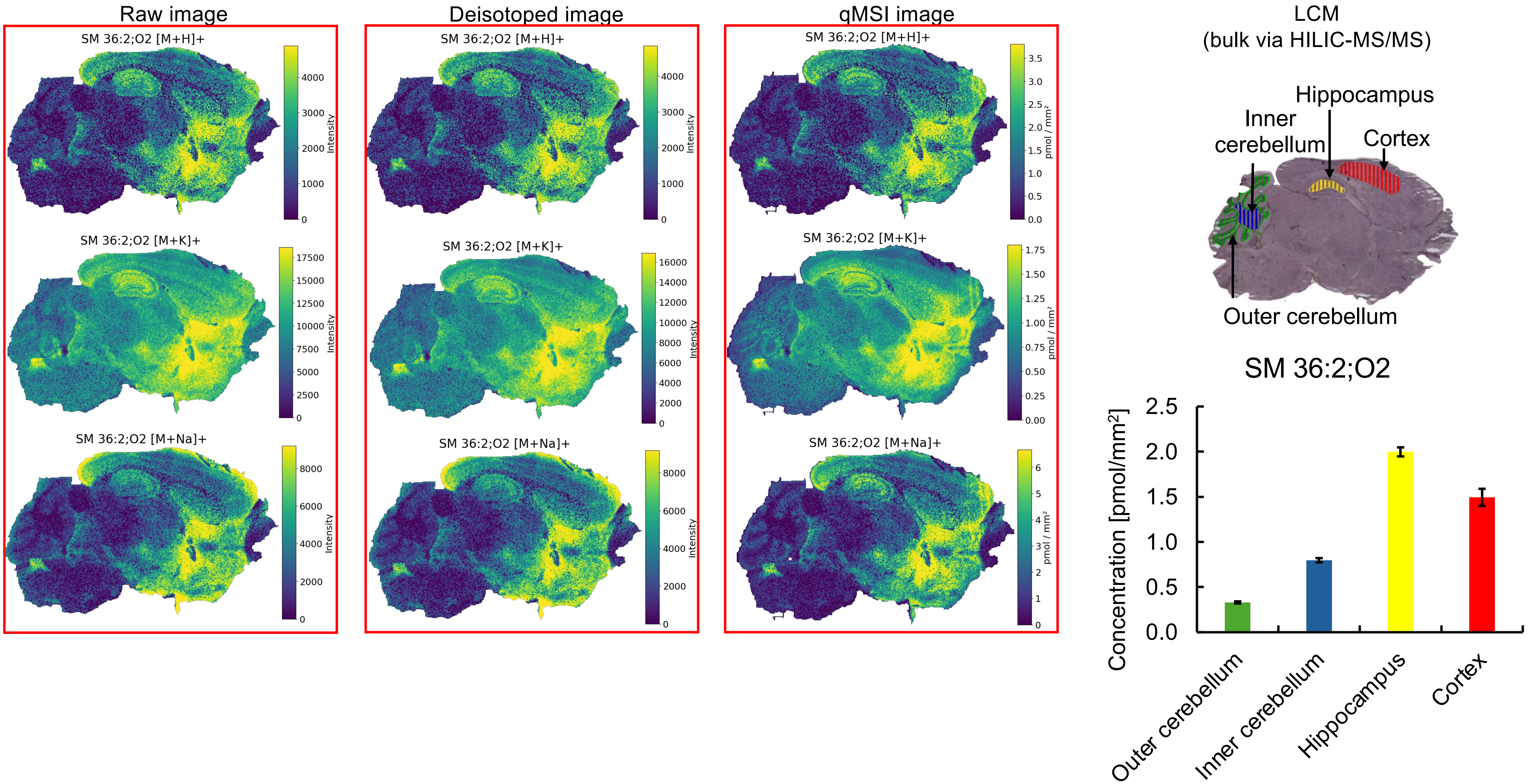


* Missing pixels for **[M+H]⁺, [M+Na]⁺** adducts (profile to centroid transformation), e.g. cerebellum.

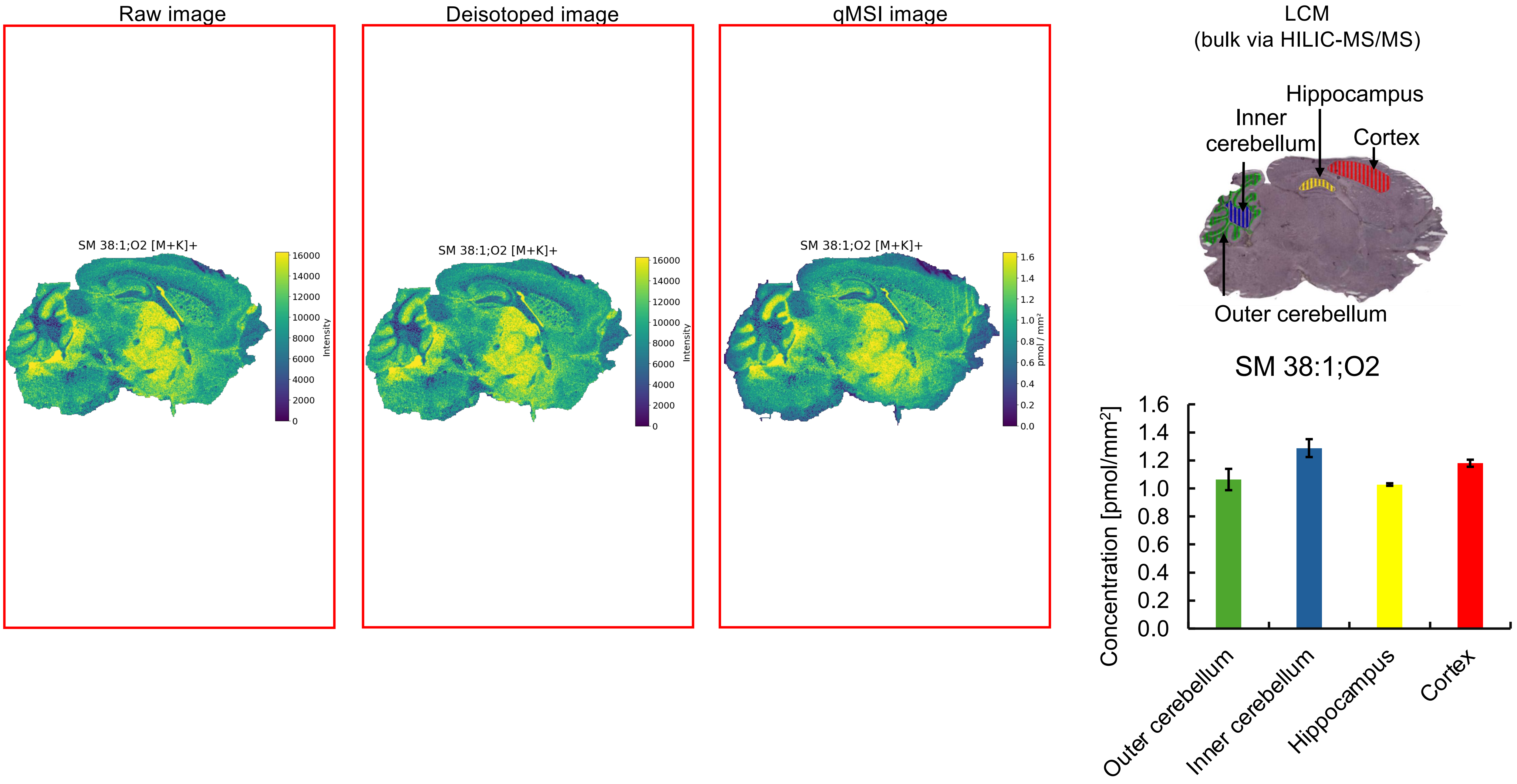


* Missing pixels for **[M+K]$^+$** adducts (profile to centroid transformation).

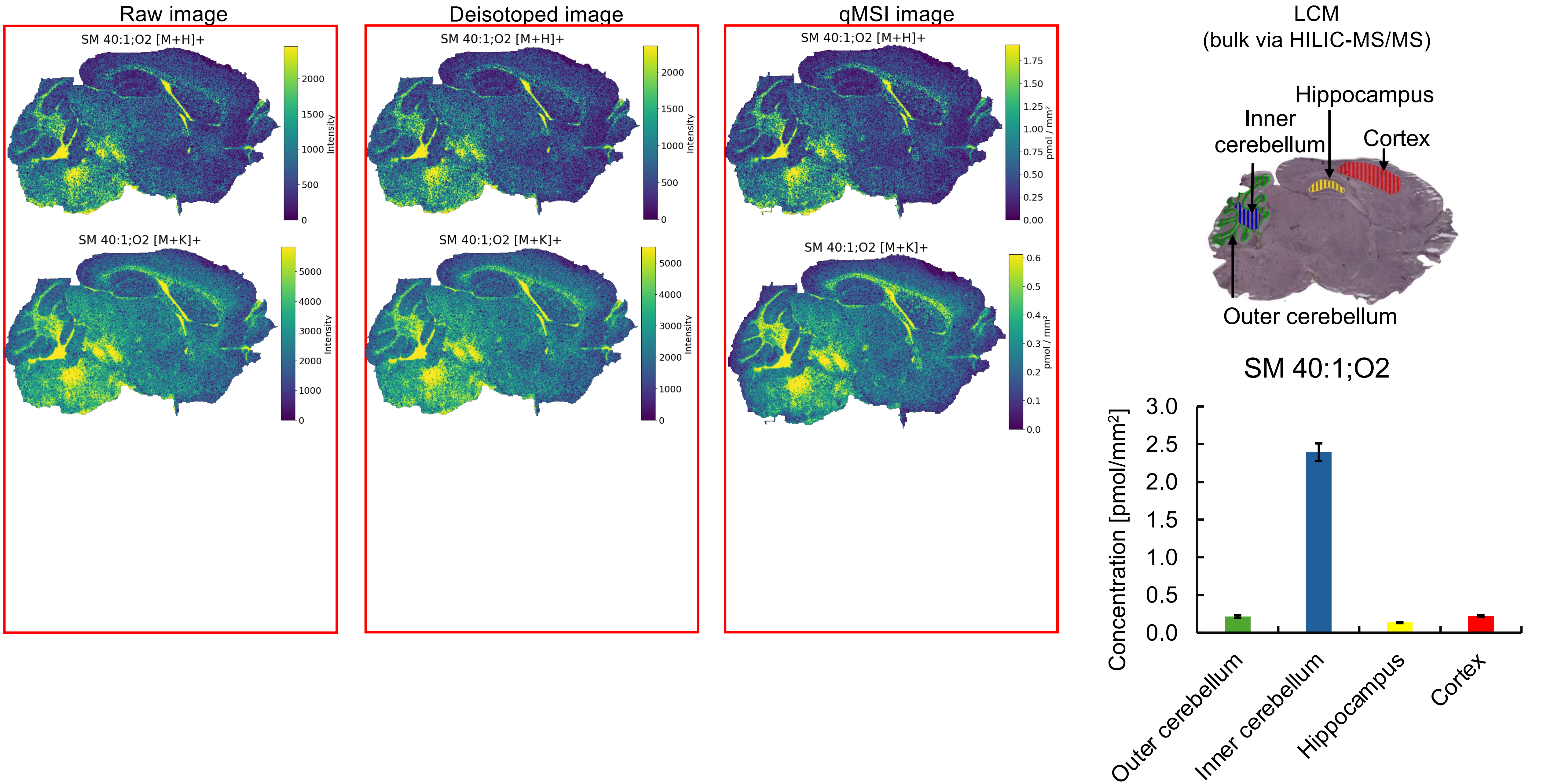

Raw image
Deisotoped image
qMSI image
SM 40:1;O2 [M+H]+
SM 40:1;O2 [M+K]+
Intensity
pmol / mm²
LCM
(bulk via HILIC-MS/MS)
Hippocampus
Inner cerebellum
Cortex
Outer cerebellum
SM 40:1;O2
Concentration [pmol/mm²]
Outer cerebellum
Inner cerebellum
Hippocampus
Cortex

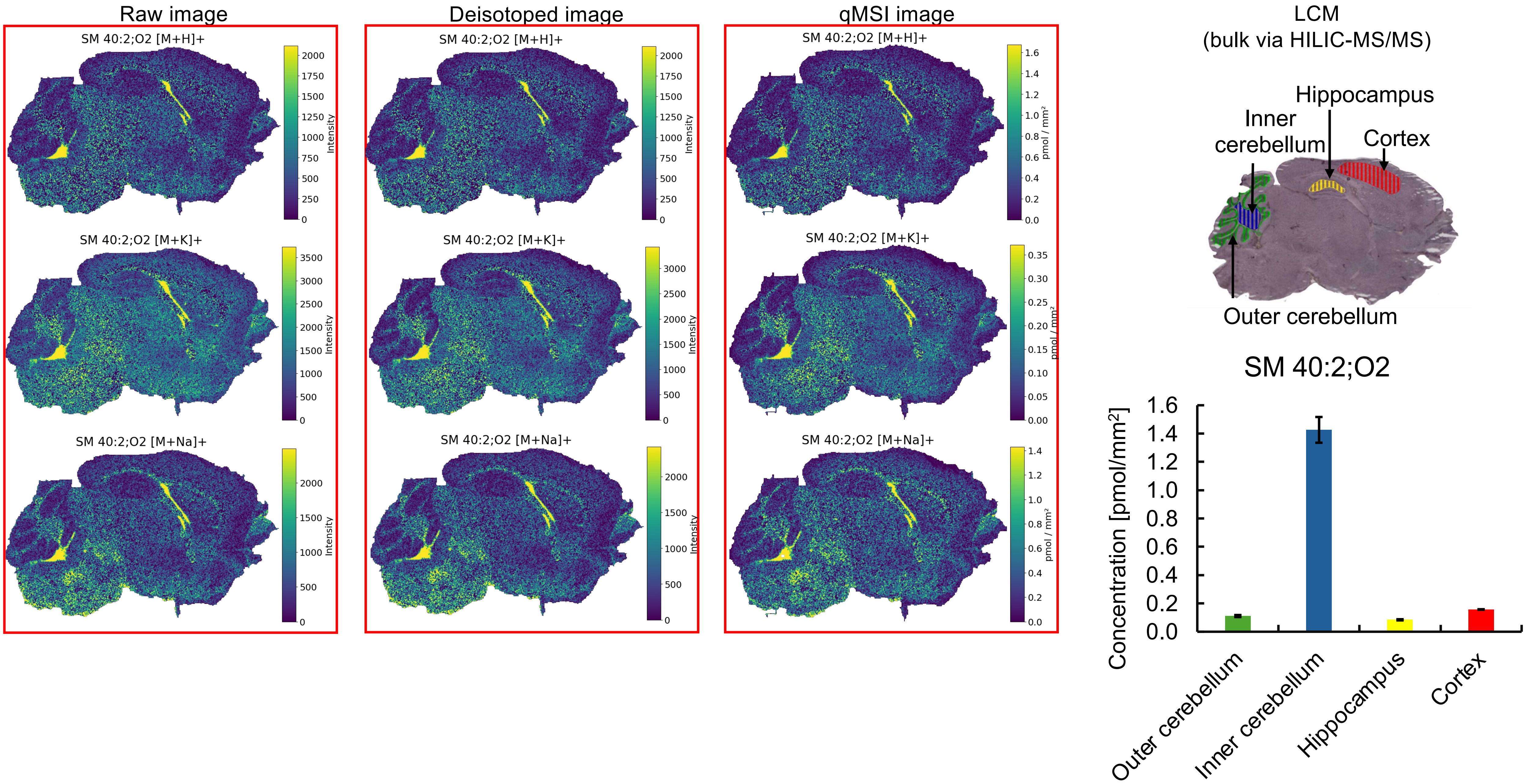

Raw image
Deisotoped image
qMSI image
LCM
(bulk via HILIC-MS/MS)
SM 40:2;O2 [M+H]+
SM 40:2;O2 [M+K]+
SM 40:2;O2 [M+Na]+
Intensity
pmol / mm²
Hippocampus
Inner cerebellum
Cortex
Outer cerebellum
SM 40:2;O2
Concentration [pmol/mm²]
Outer cerebellum
Inner cerebellum
Hippocampus
Cortex

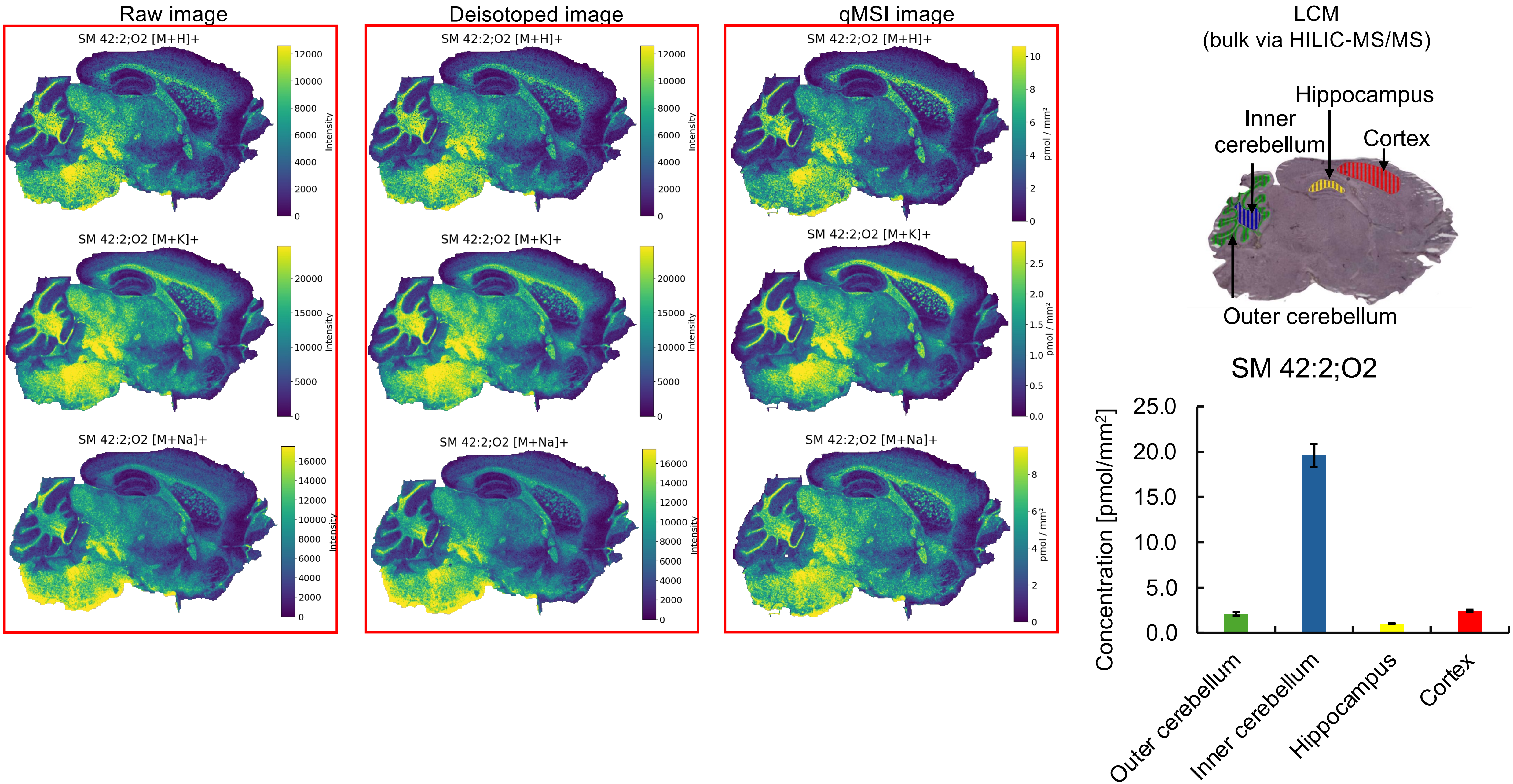

Raw image
Deisotoped image
qMSI image
SM 42:2;O2 [M+H]+
SM 42:2;O2 [M+K]+
SM 42:2;O2 [M+Na]+
Intensity
pmol / mm²
LCM
(bulk via HILIC-MS/MS)
Hippocampus
Inner cerebellum
Cortex
Outer cerebellum
SM 42:2;O2
Concentration [pmol/mm²]
25.0
20.0
15.0
10.0
5.0
0.0
Outer cerebellum
Inner cerebellum
Hippocampus
Cortex

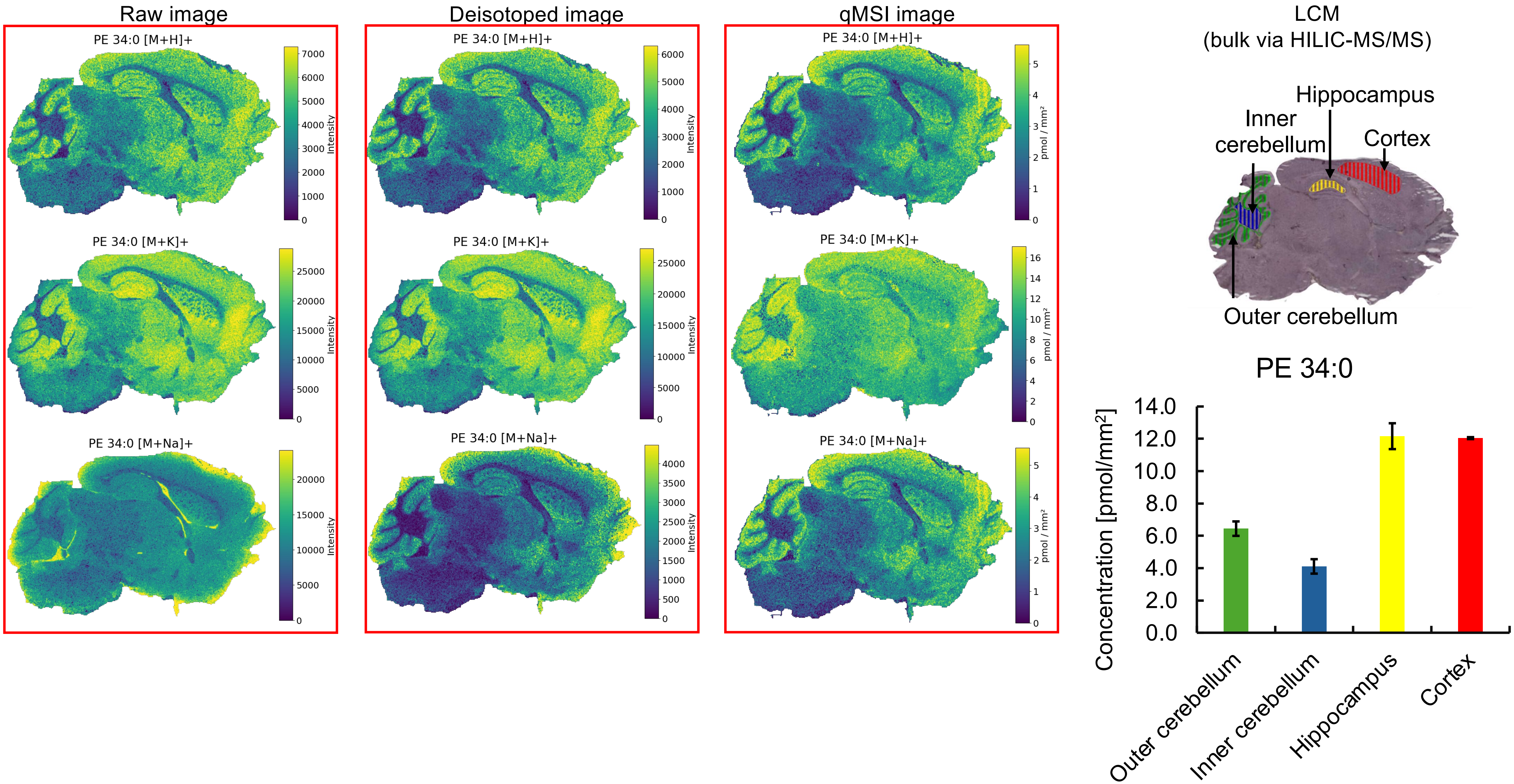


* Missing pixels for **[M+H]⁺, [M+Na]⁺** adducts (profile to centroid transformation).

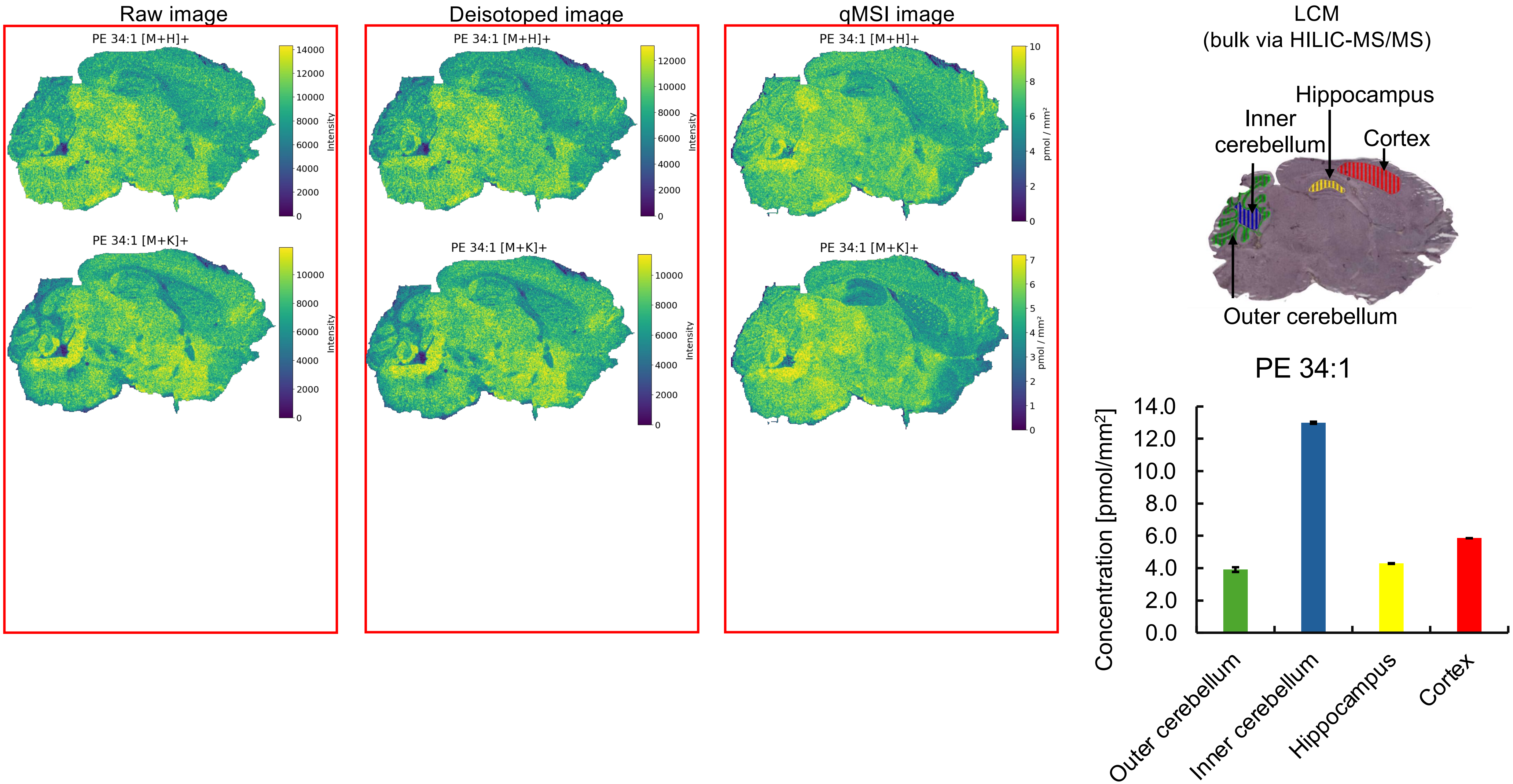

Raw image
PE 34:1 [M+H]+
PE 34:1 [M+K]+
Intensity
Deisotoped image
PE 34:1 [M+H]+
PE 34:1 [M+K]+
Intensity
qMSI image
PE 34:1 [M+H]+
PE 34:1 [M+K]+
pmol / mm²
LCM
(bulk via HILIC-MS/MS)
Hippocampus
Inner cerebellum
Cortex
Outer cerebellum
PE 34:1
Concentration [pmol/mm²]
Outer cerebellum
Inner cerebellum
Hippocampus
Cortex

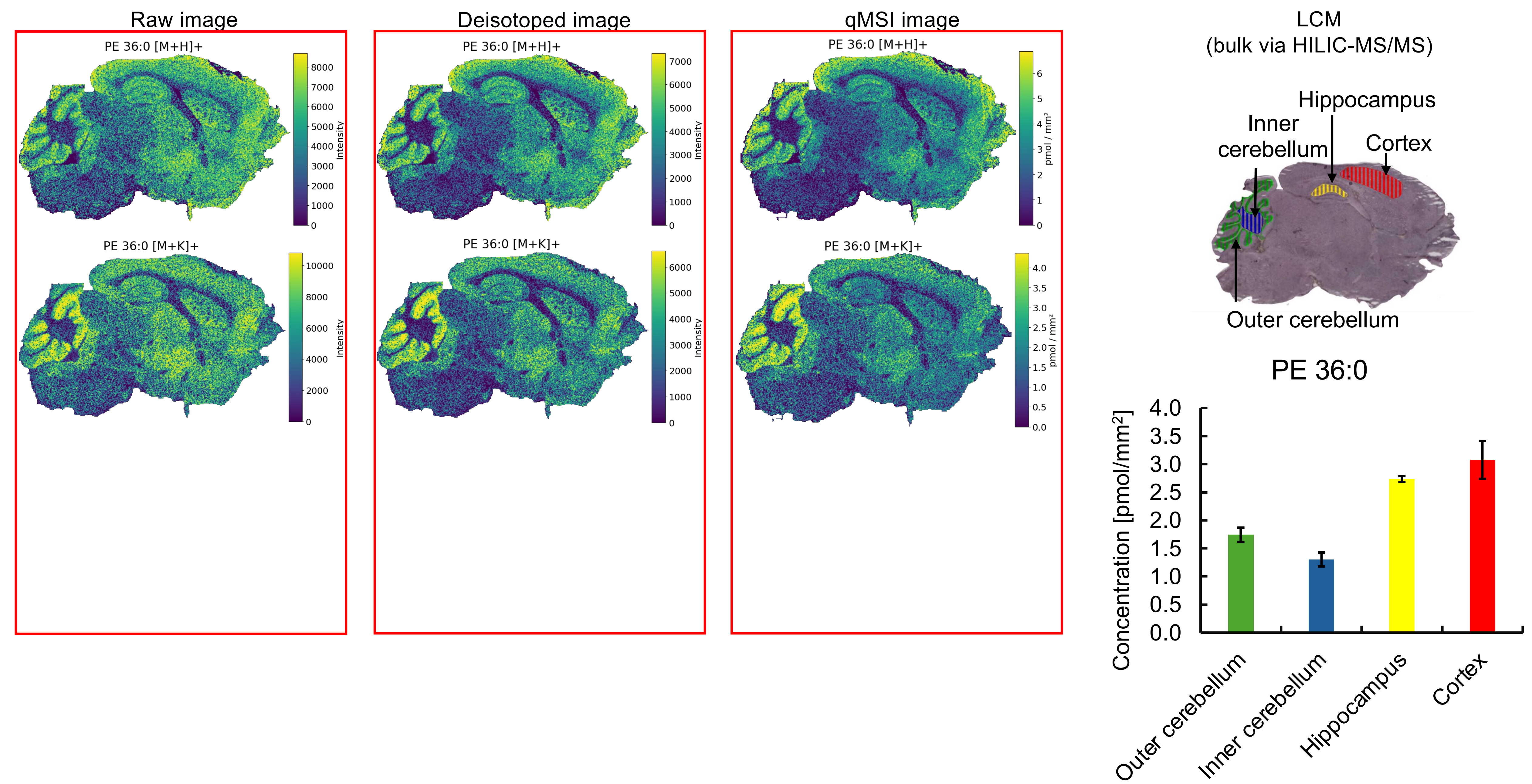


* Missing pixels for **[M+H]⁺, [M+K]⁺** adducts (profile to centroid transformation).

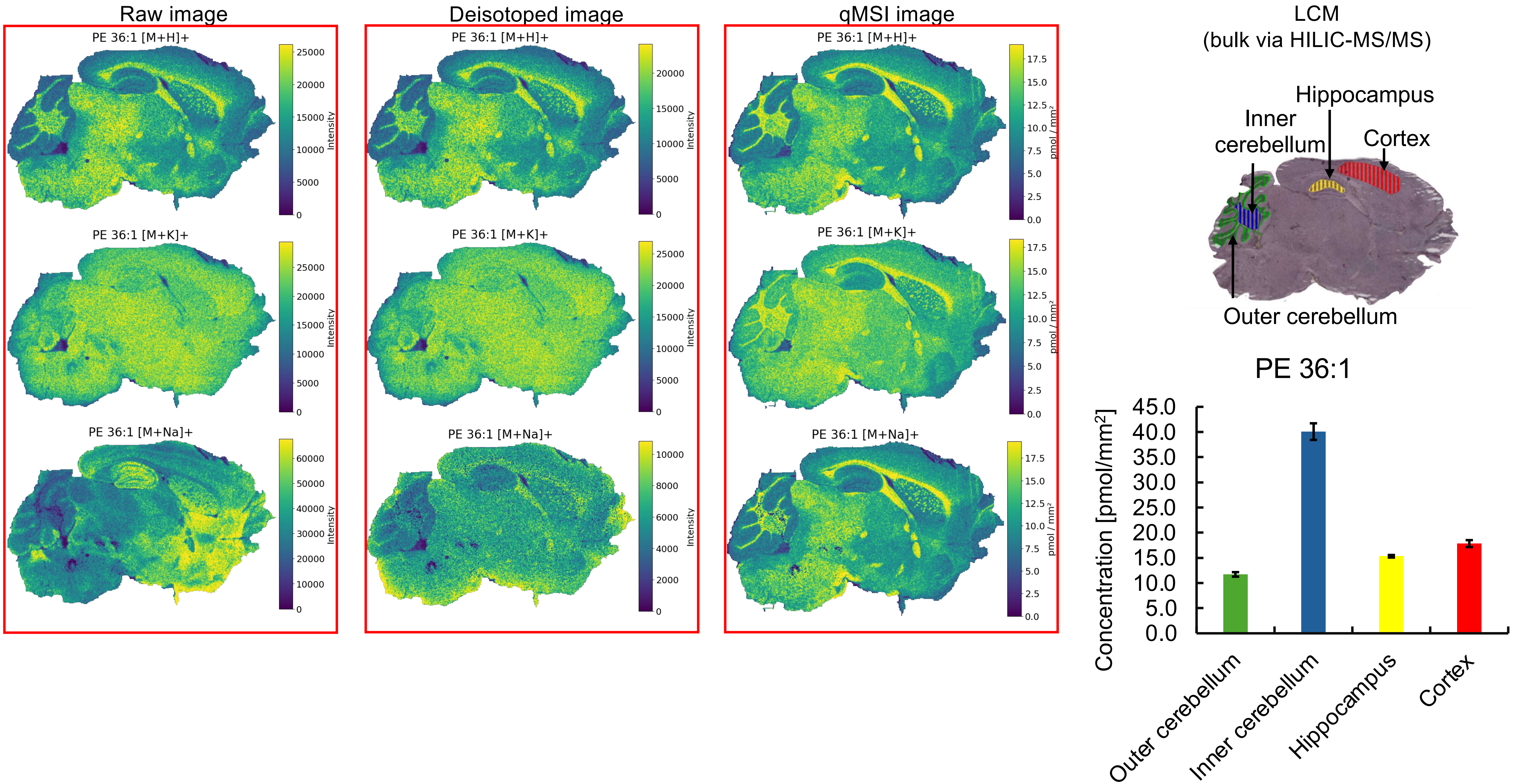

Raw image
Deisotoped image
qMSI image
PE 36:1 [M+H]+
PE 36:1 [M+K]+
PE 36:1 [M+Na]+
Intensity
pmol / mm²
LCM
(bulk via HILIC-MS/MS)
Hippocampus
Inner cerebellum
Cortex
Outer cerebellum
PE 36:1
Concentration [pmol/mm²]
Outer cerebellum
Inner cerebellum
Hippocampus
Cortex

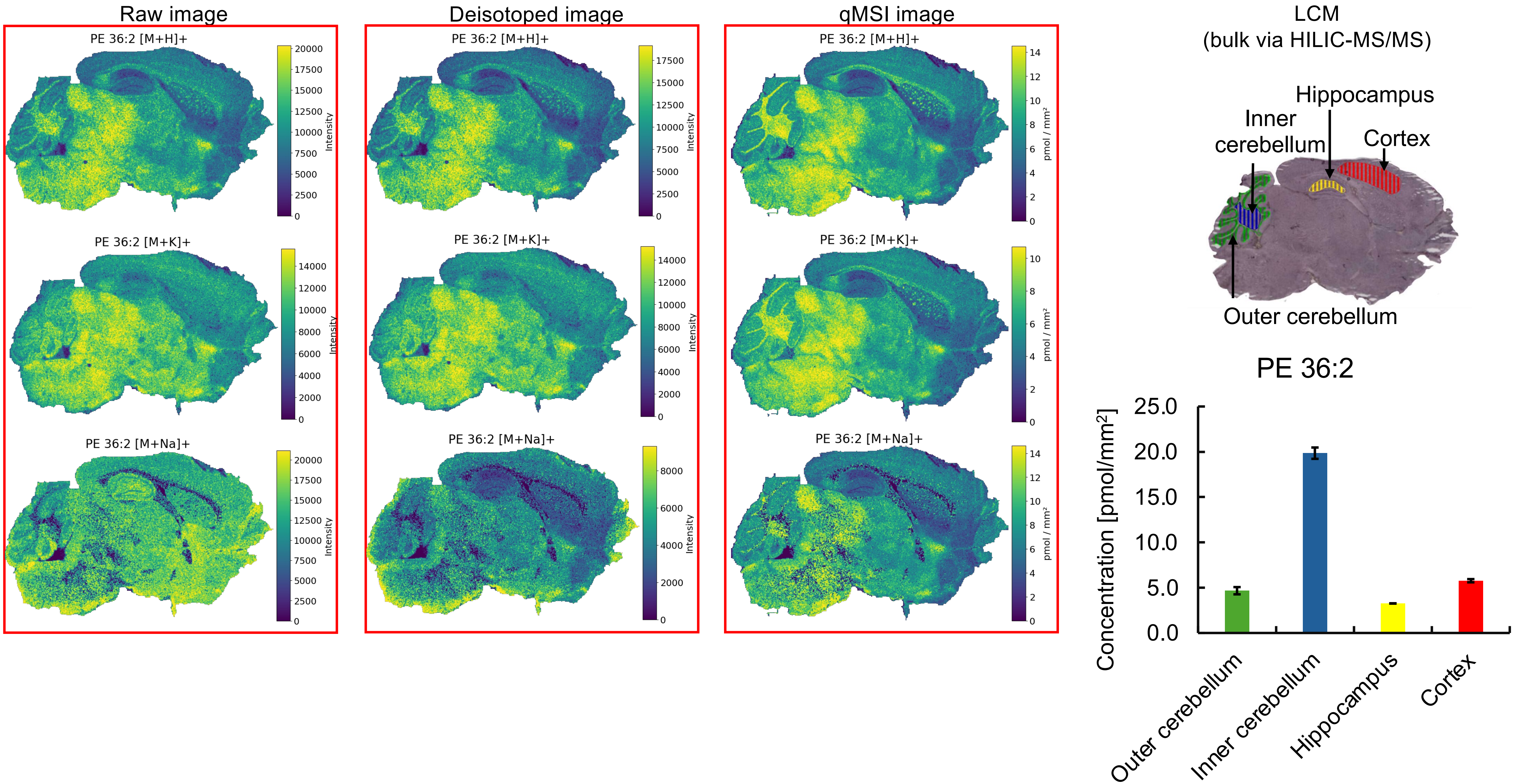

Raw image
Deisotoped image
qMSI image
PE 36:2 [M+H]+
PE 36:2 [M+K]+
PE 36:2 [M+Na]+
Intensity
pmol / mm²
LCM
(bulk via HILIC-MS/MS)
Hippocampus
Inner cerebellum
Cortex
Outer cerebellum
PE 36:2
Concentration [pmol/mm²]
25.0
20.0
15.0
10.0
5.0
0.0
Outer cerebellum
Inner cerebellum
Hippocampus
Cortex

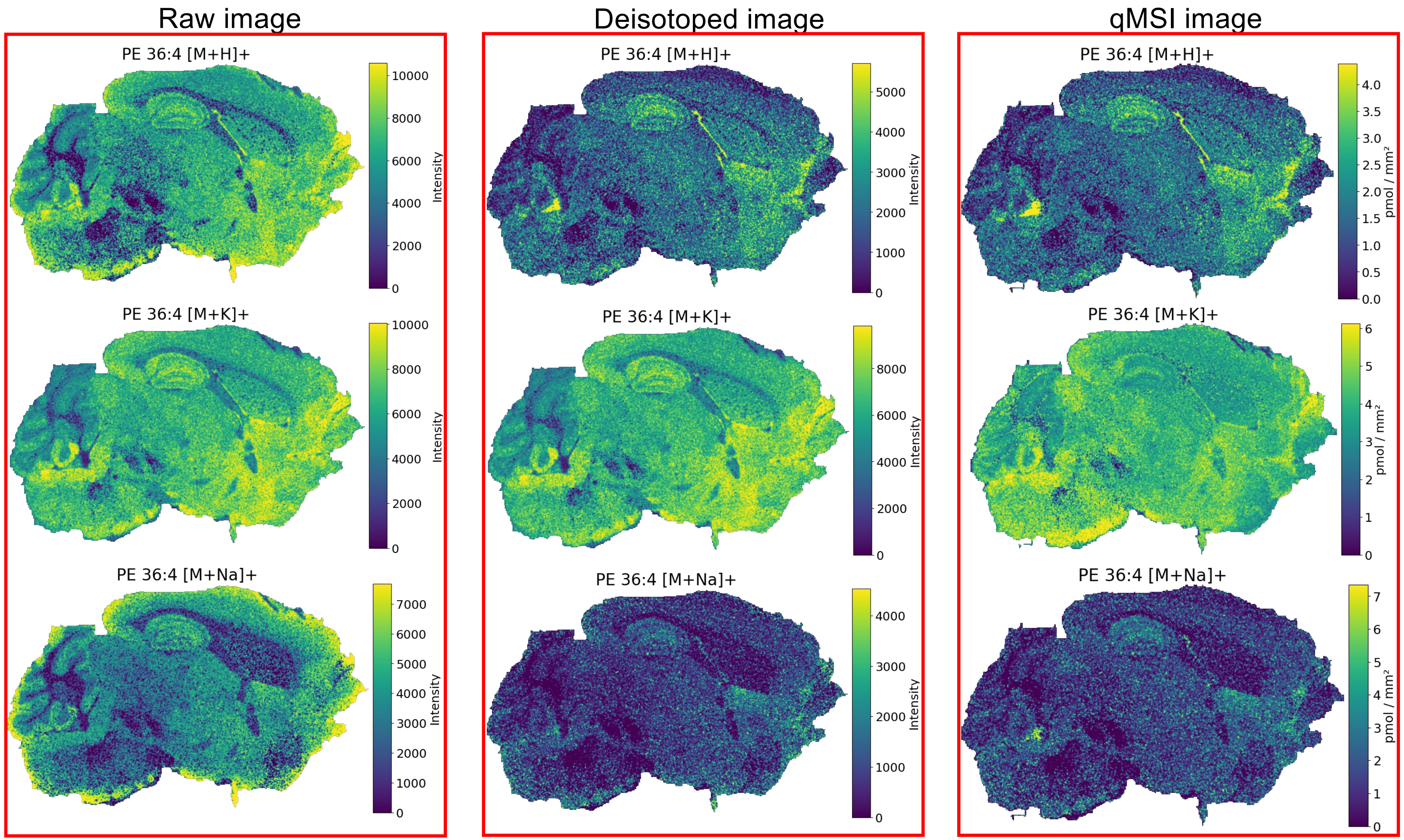


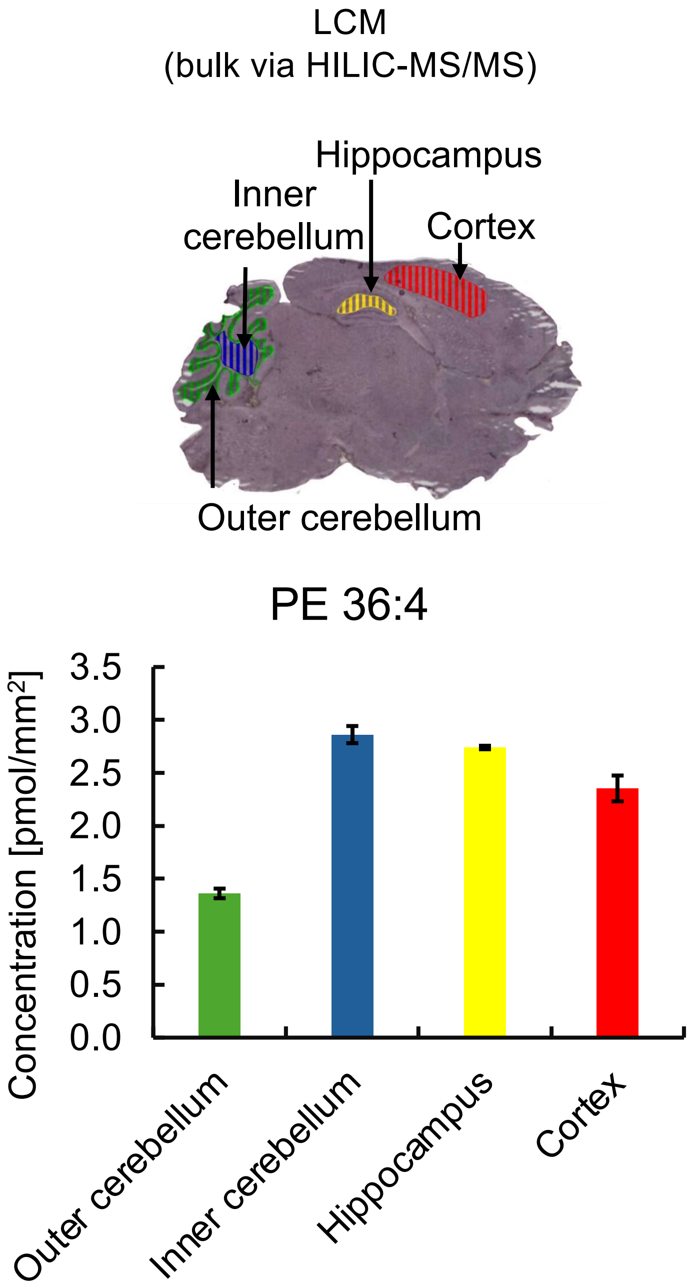


* Missing pixels for **[M+H]⁺, [M+K]⁺,[M+Na]⁺** adducts (profile to centroid transformation).

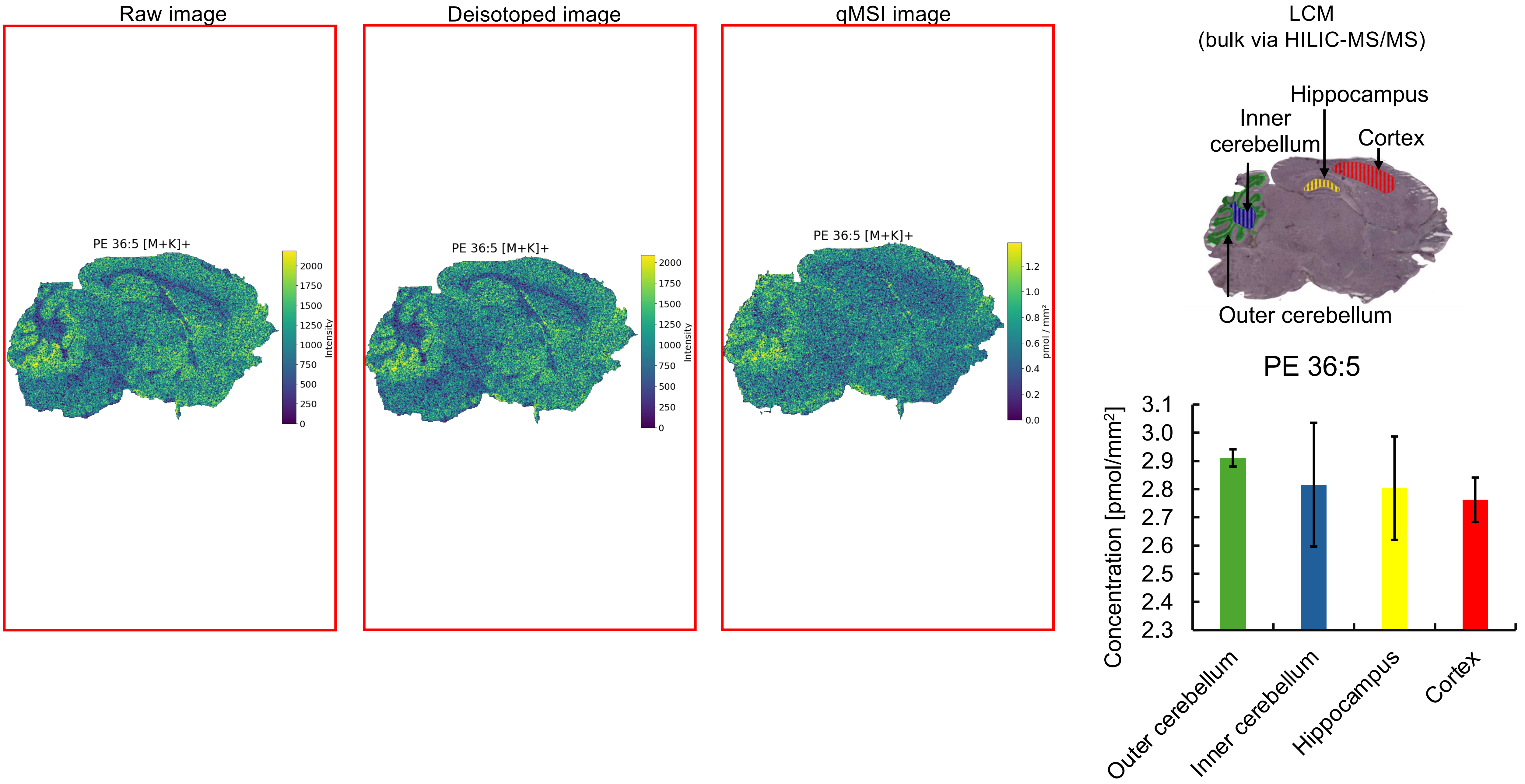

Raw image
PE 36:5 [M+K]+
Intensity
Deisotoped image
PE 36:5 [M+K]+
Intensity
qMSI image
PE 36:5 [M+K]+
pmol / mm²
LCM
(bulk via HILIC-MS/MS)
Hippocampus
Inner cerebellum
Cortex
Outer cerebellum
PE 36:5
Concentration [pmol/mm²]
Outer cerebellum
Inner cerebellum
Hippocampus
Cortex

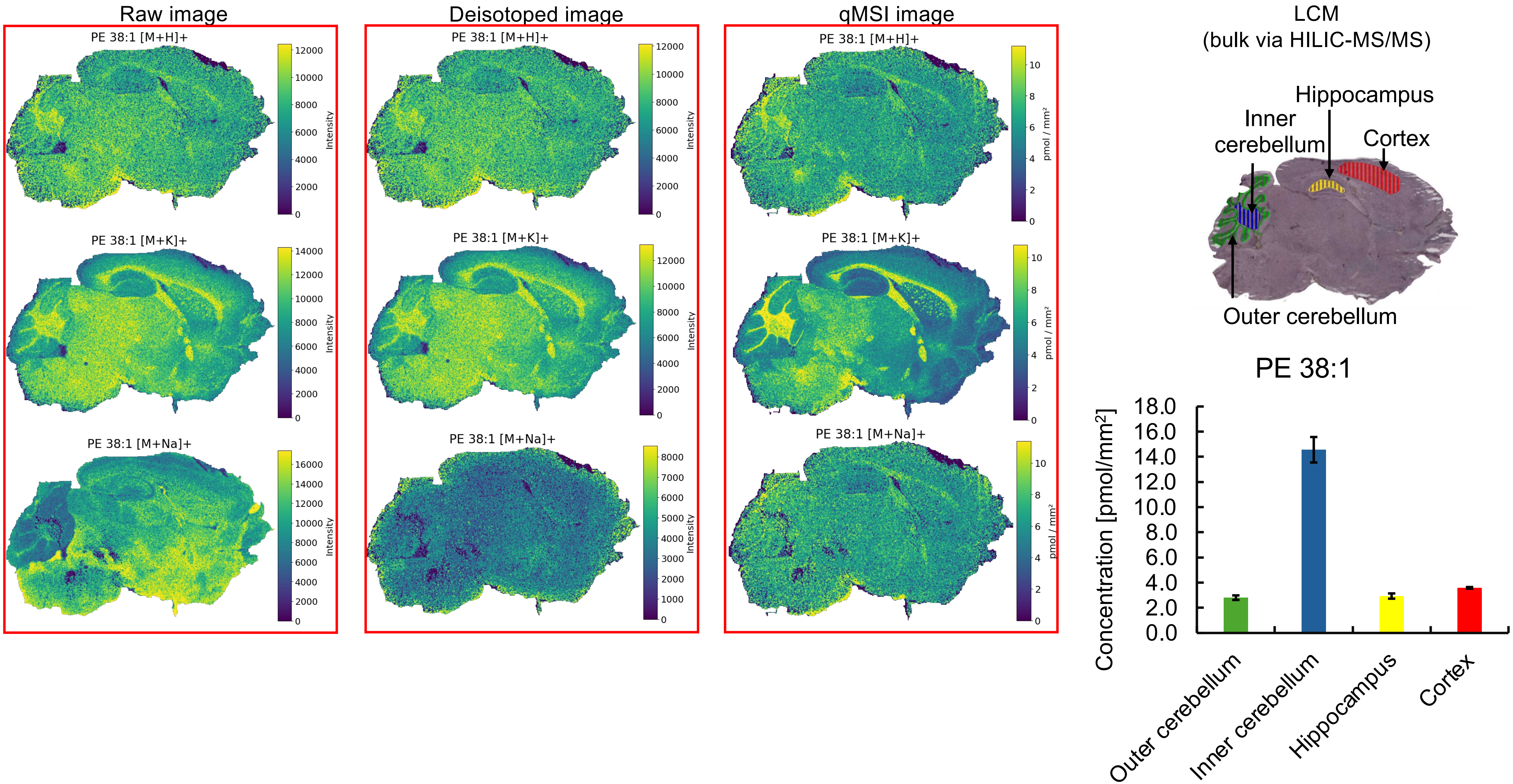
Raw image
Deisotoped image
qMSI image
PE 38:1 [M+H]+
PE 38:1 [M+K]+
PE 38:1 [M+Na]+
Intensity
pmol / mm²
LCM
(bulk via HILIC-MS/MS)
Hippocampus
Inner cerebellum
Cortex
Outer cerebellum
PE 38:1
Concentration [pmol/mm²]
0.0
2.0
4.0
6.0
8.0
10.0
12.0
14.0
16.0
18.0
Outer cerebellum
Inner cerebellum
Hippocampus
Cortex

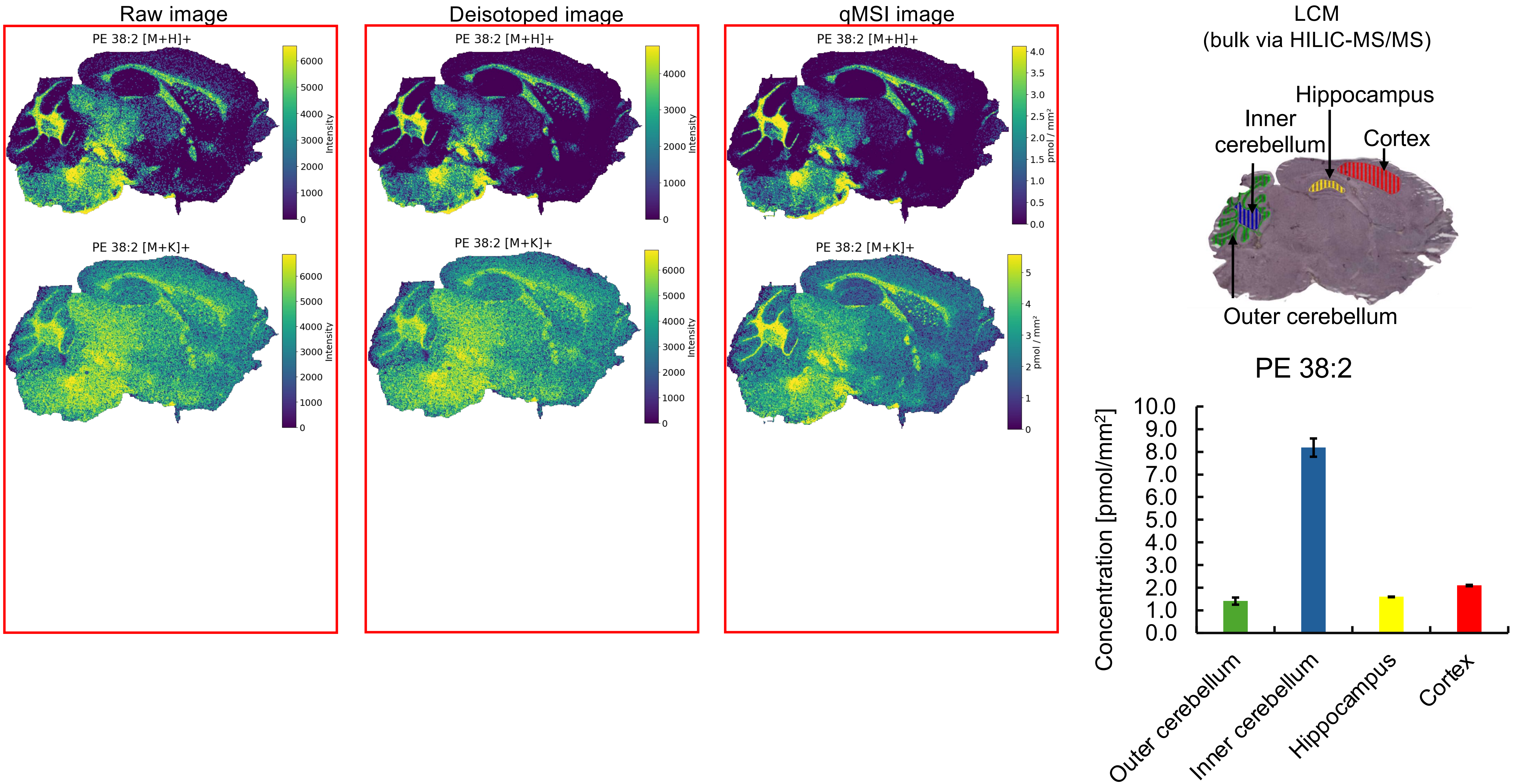


* Missing pixels for **[M+H]$^+$** adducts (profile to centroid transformation).

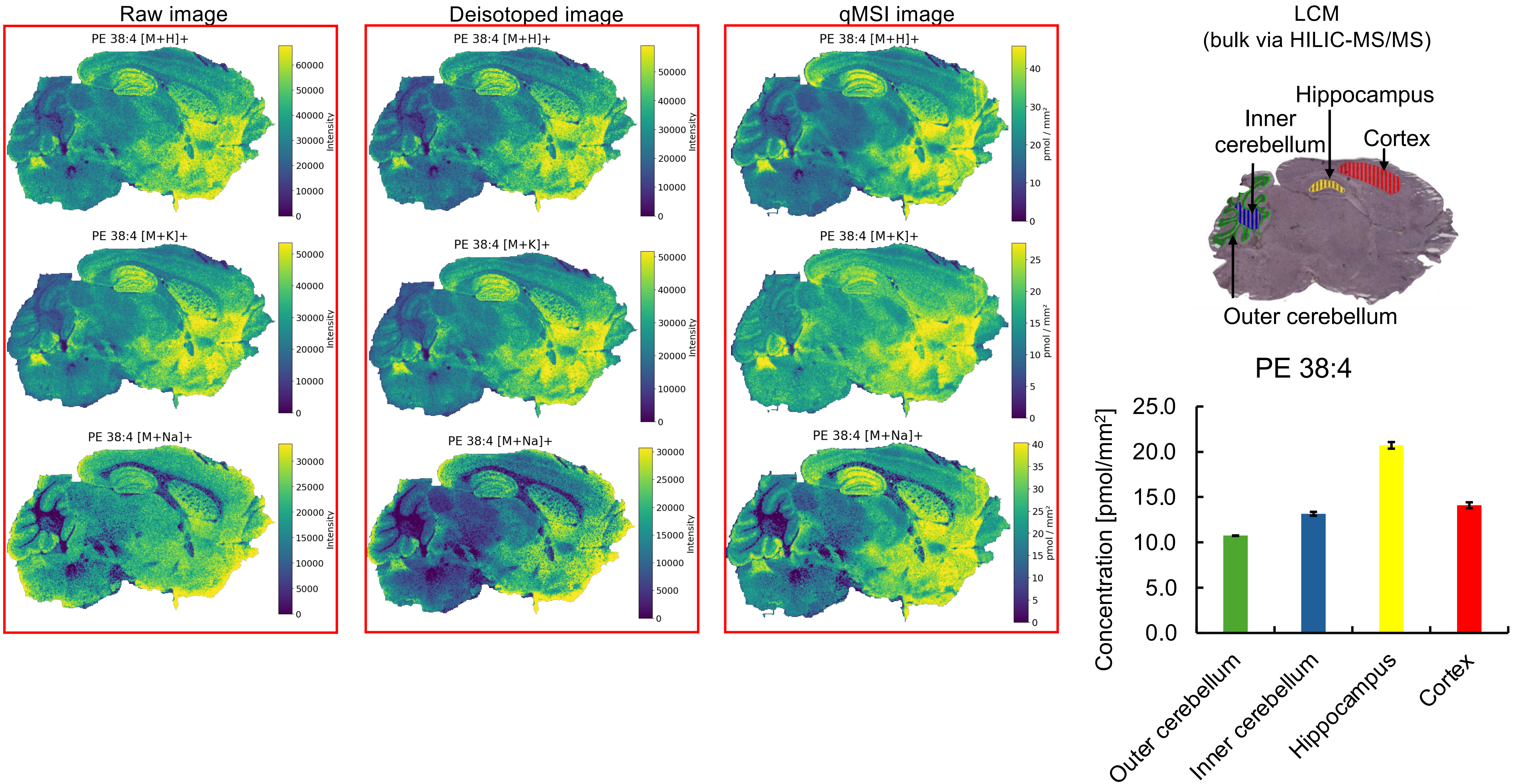
Raw image
Deisotoped image
qMSI image
LCM
(bulk via HILIC-MS/MS)
PE 38:4 [M+H]+
PE 38:4 [M+K]+
PE 38:4 [M+Na]+
Intensity
pmol / mm²
Hippocampus
Inner cerebellum
Cortex
Outer cerebellum
PE 38:4
Concentration [pmol/mm²]
25.0
20.0
15.0
10.0
5.0
0.0
Outer cerebellum
Inner cerebellum
Hippocampus
Cortex

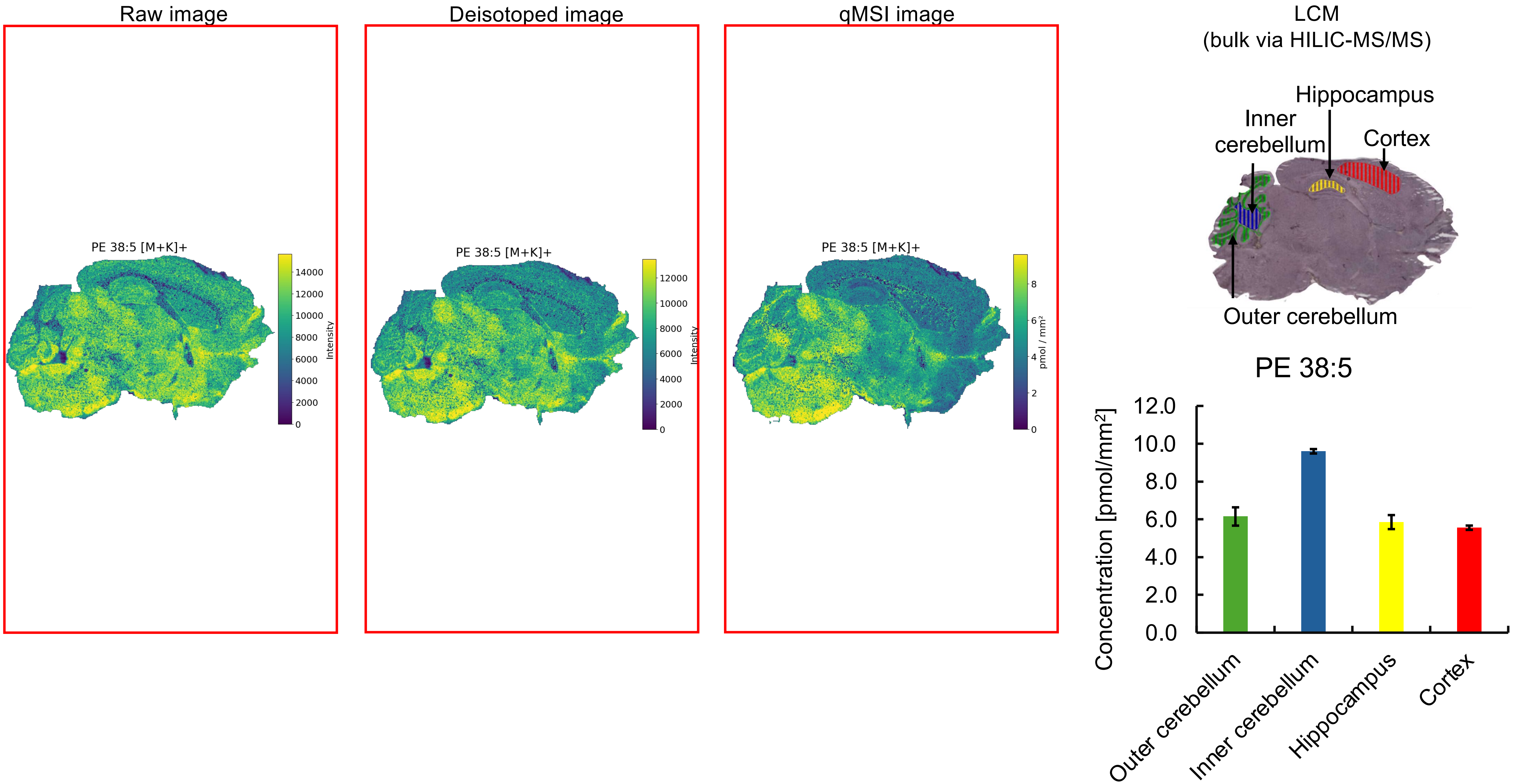


* Missing pixels for **[M+K]⁺** adducts (profile to centroid transformation).

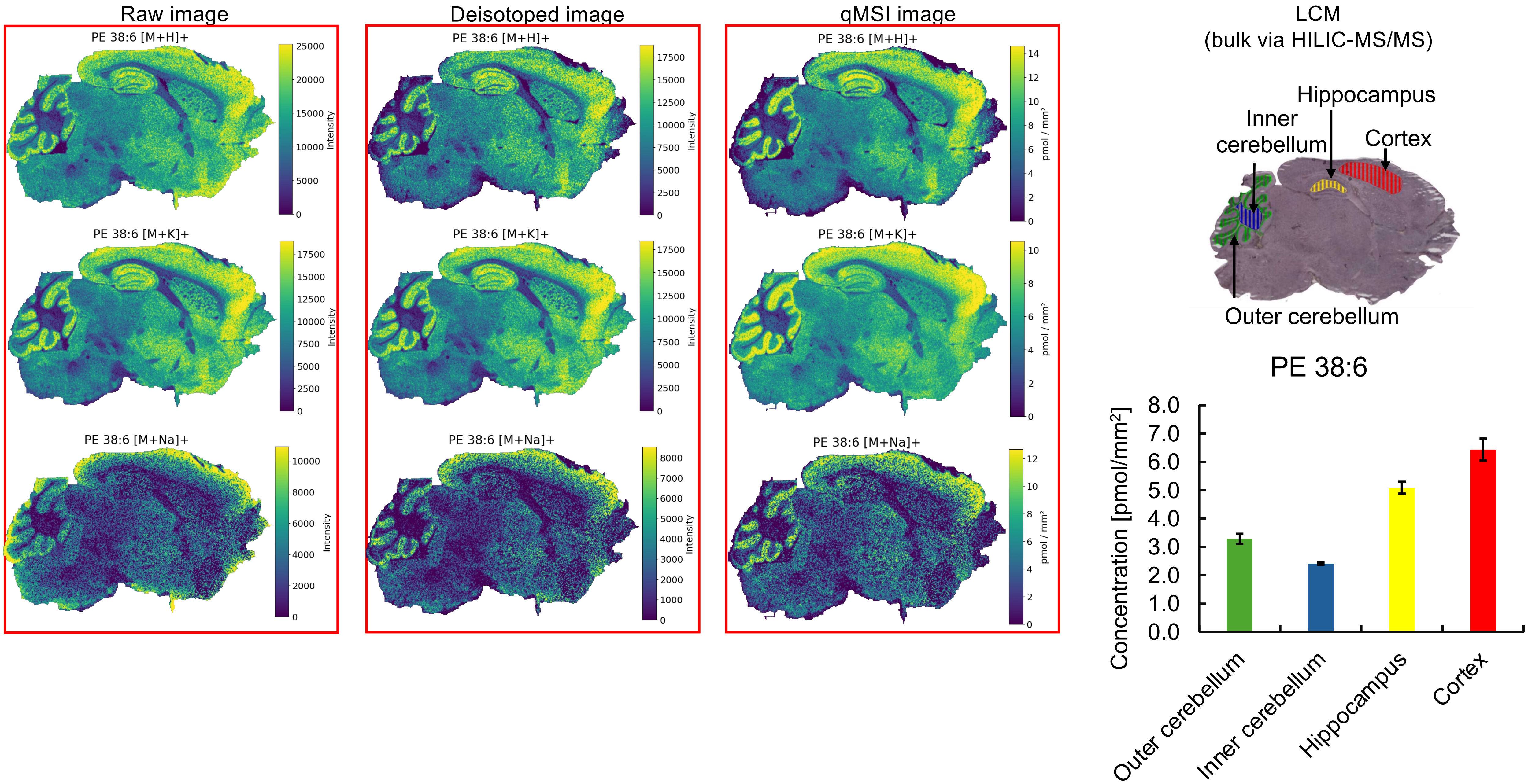


* Missing pixels for **[M+H]⁺**, **[M+K]⁺**, **[M+Na]⁺** adducts (profile to centroid transformation).

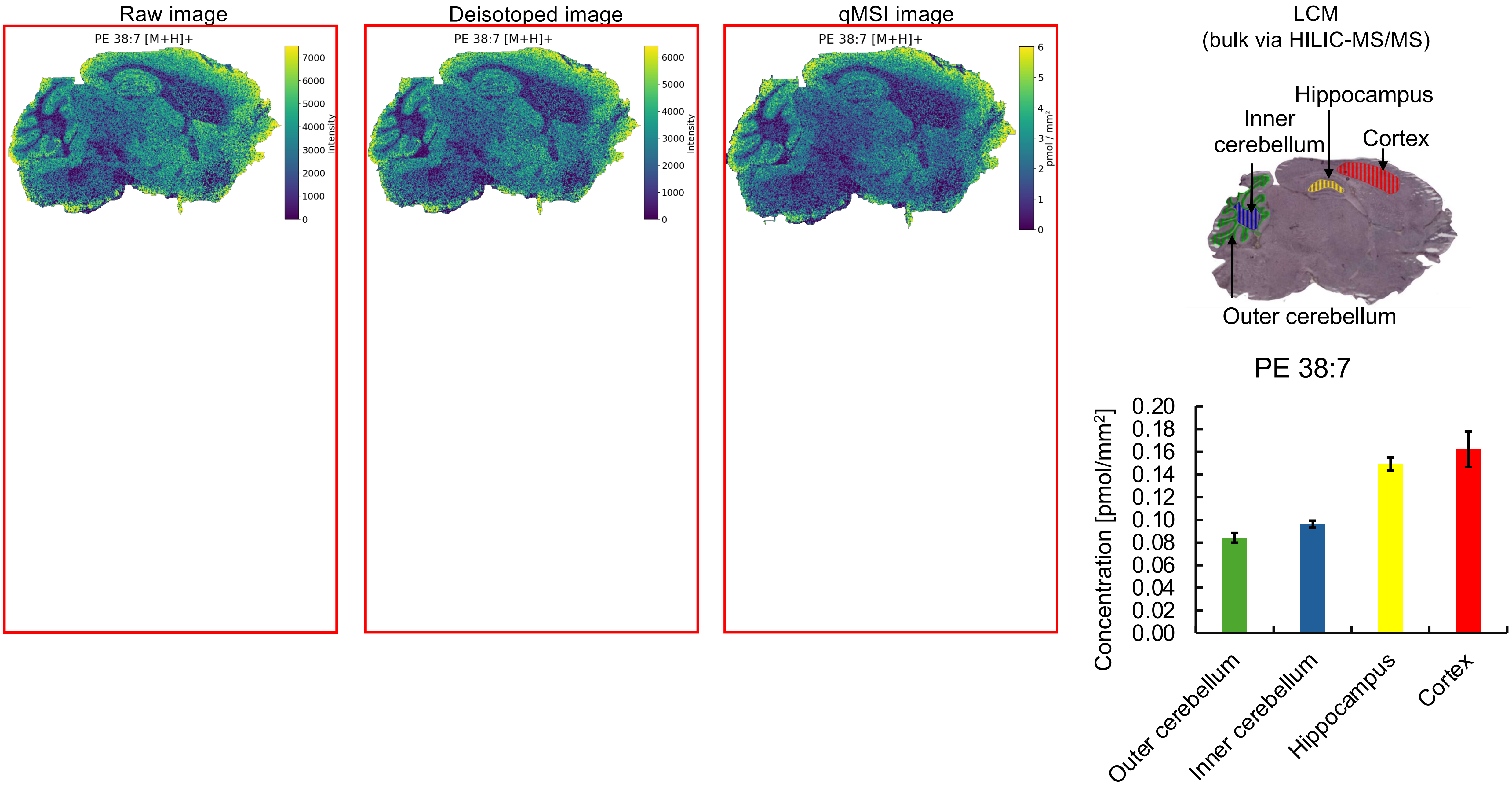


* Missing pixels for **[M+H]$^+$** adducts (profile to centroid transformation).

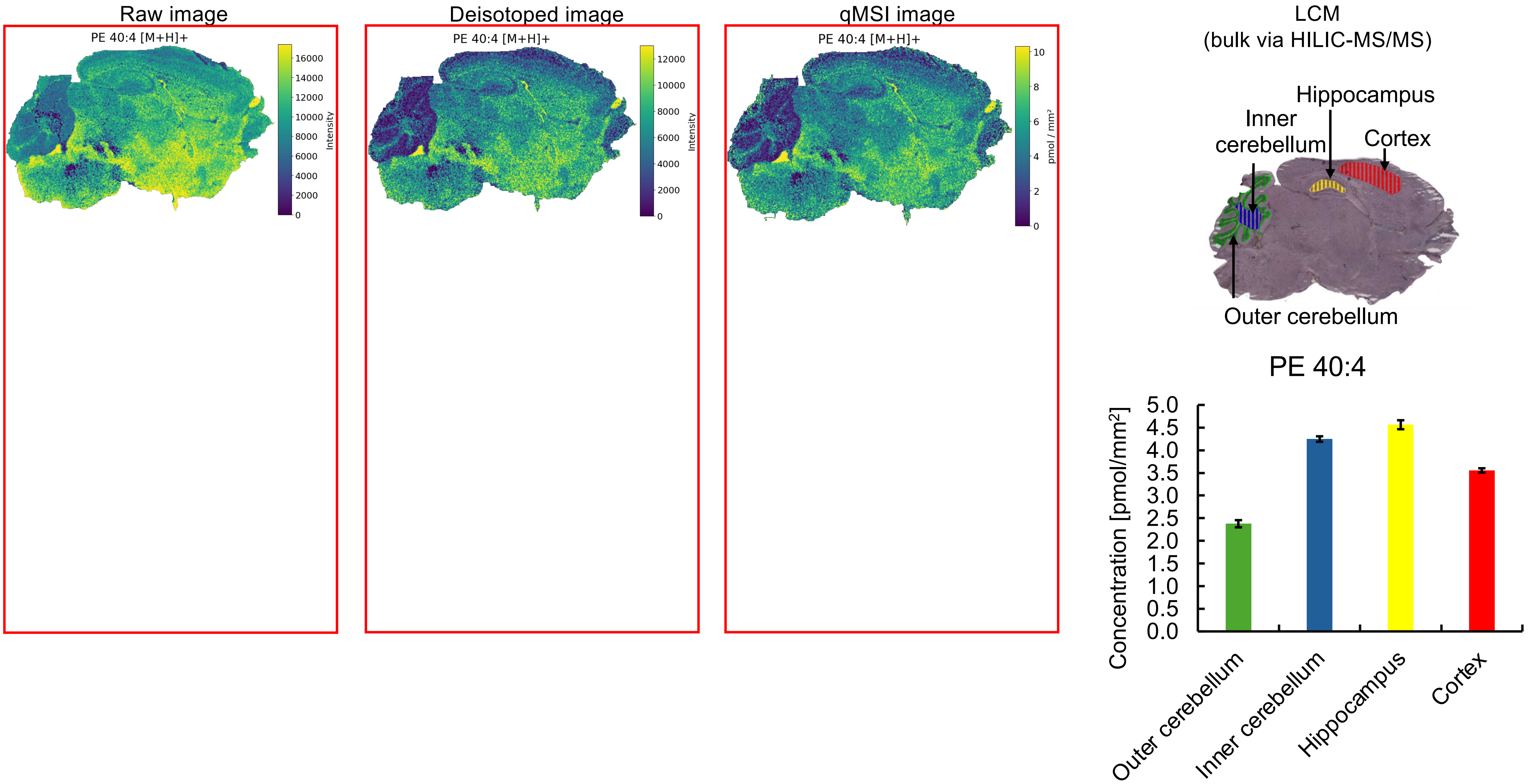


* Missing pixels for **[M+H]⁺** adducts (profile to centroid transformation).

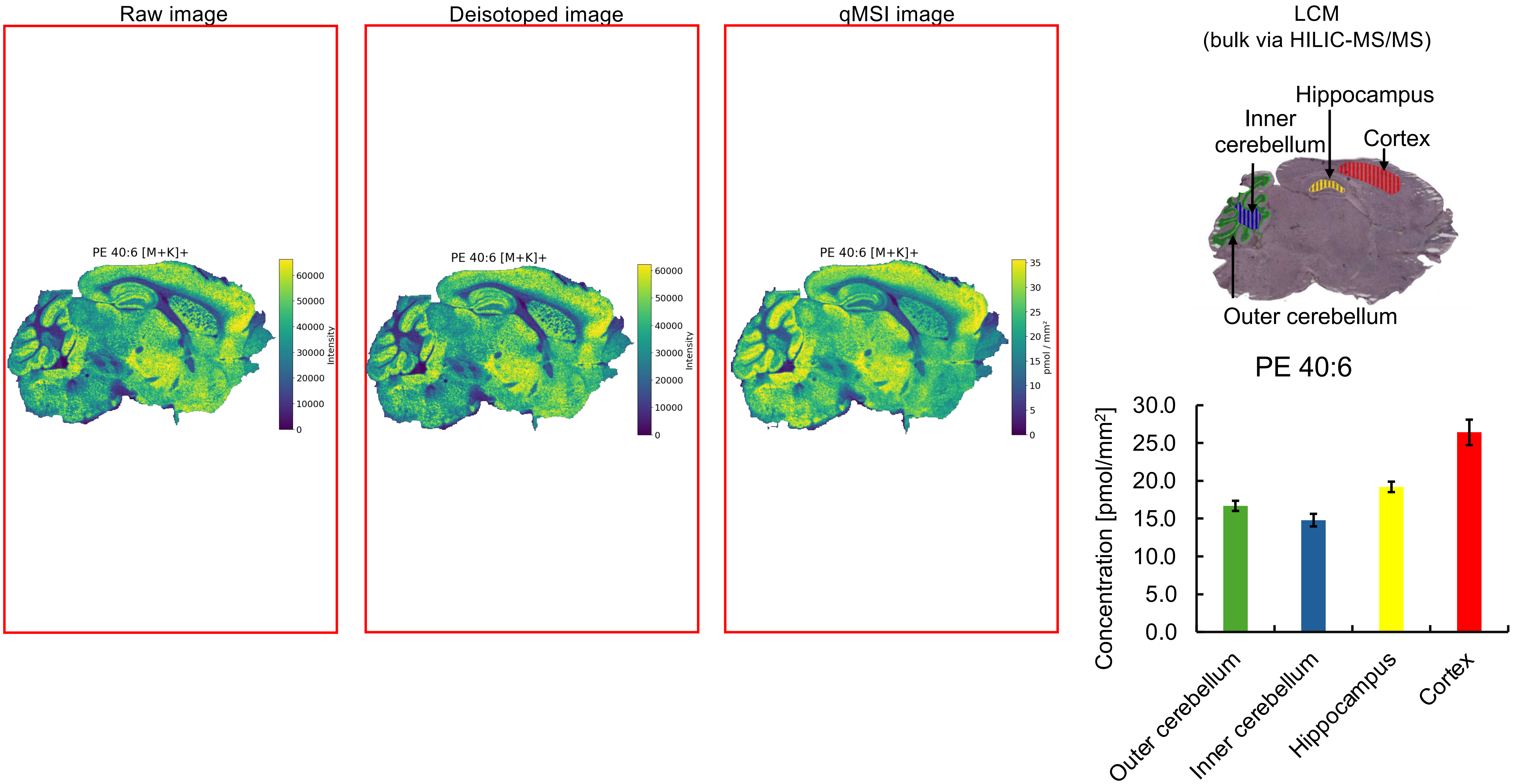

Raw image
Deisotoped image
qMSI image
PE 40:6 [M+K]+
Intensity
pmol / mm²
LCM
(bulk via HILIC-MS/MS)
Hippocampus
Inner cerebellum
Cortex
Outer cerebellum
PE 40:6
Concentration [pmol/mm²]
Outer cerebellum
Inner cerebellum
Hippocampus
Cortex

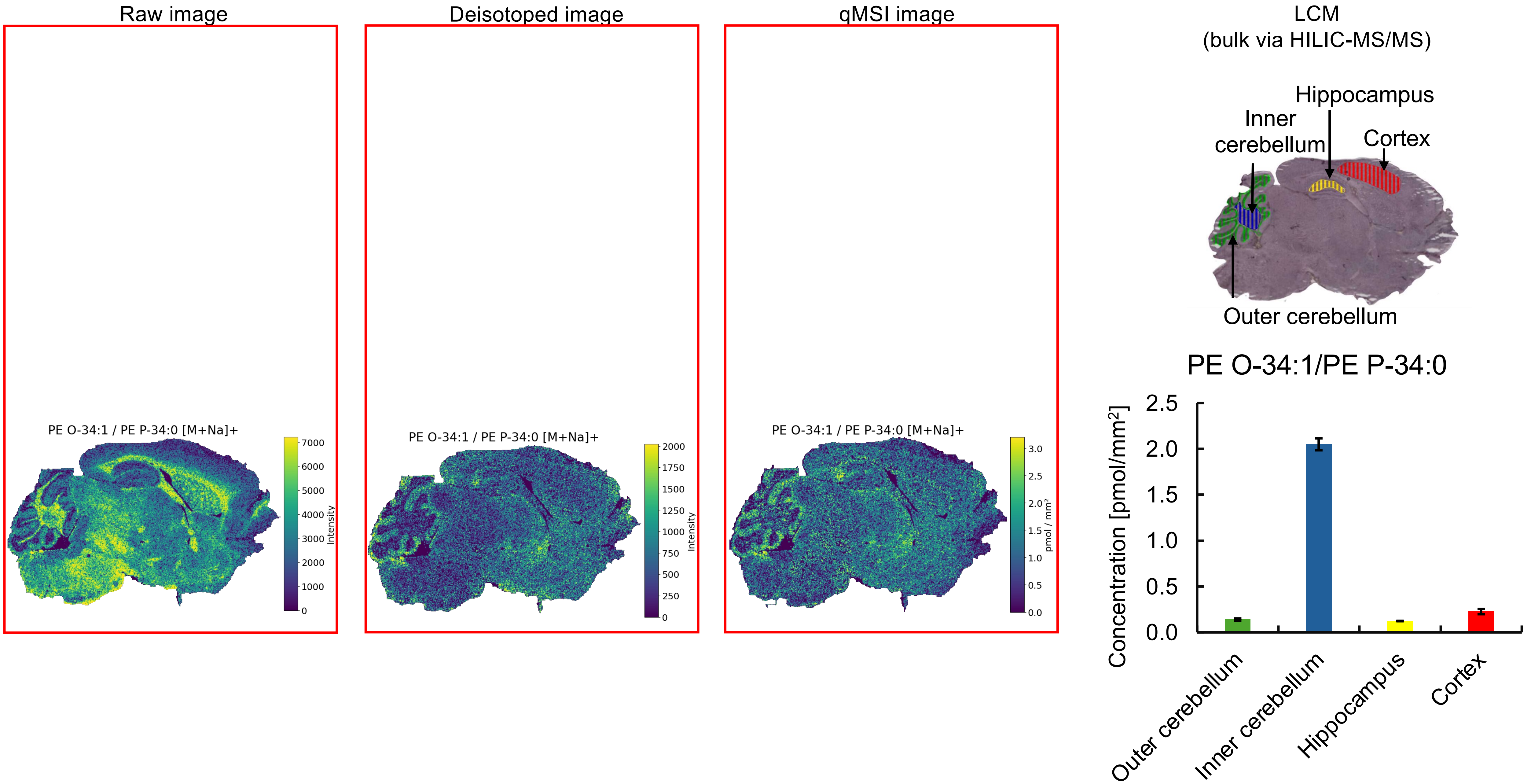

Raw image
Deisotoped image
qMSI image
LCM
(bulk via HILIC-MS/MS)
PE O-34:1 / PE P-34:0 [M+Na]+
Intensity
0
1000
2000
3000
4000
5000
6000
7000
PE O-34:1 / PE P-34:0 [M+Na]+
Intensity
0
250
500
750
1000
1250
1500
1750
2000
PE O-34:1 / PE P-34:0 [M+Na]+
pmol / mm²
0.0
0.5
1.0
1.5
2.0
2.5
3.0
Hippocampus
Inner cerebellum
Cortex
Outer cerebellum
PE O-34:1/PE P-34:0
Concentration [pmol/mm²]
0.0
0.5
1.0
1.5
2.0
2.5
Outer cerebellum
Inner cerebellum
Hippocampus
Cortex

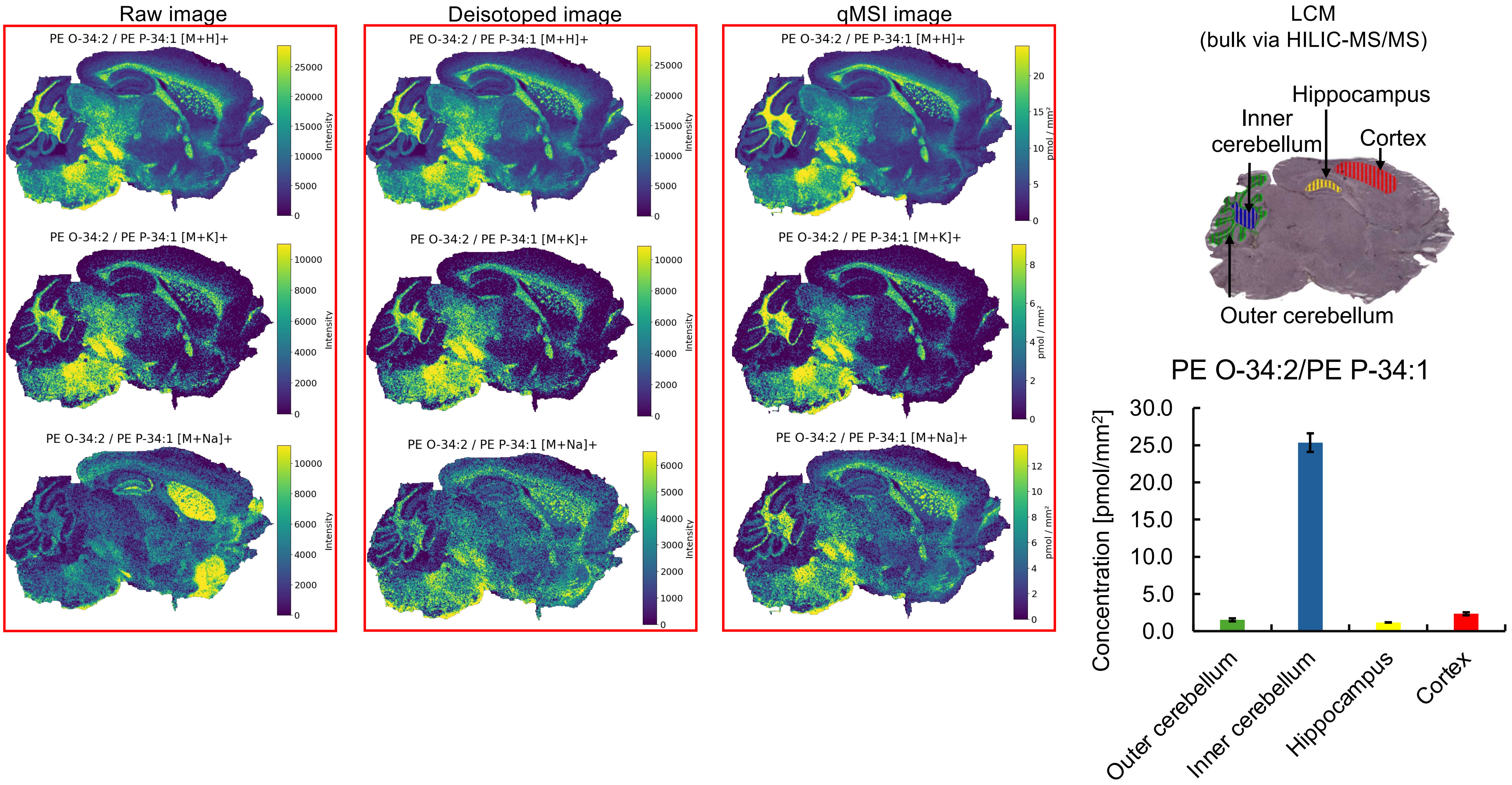


* Missing pixels for **[M+K]⁺** adducts (profile to centroid transformation).

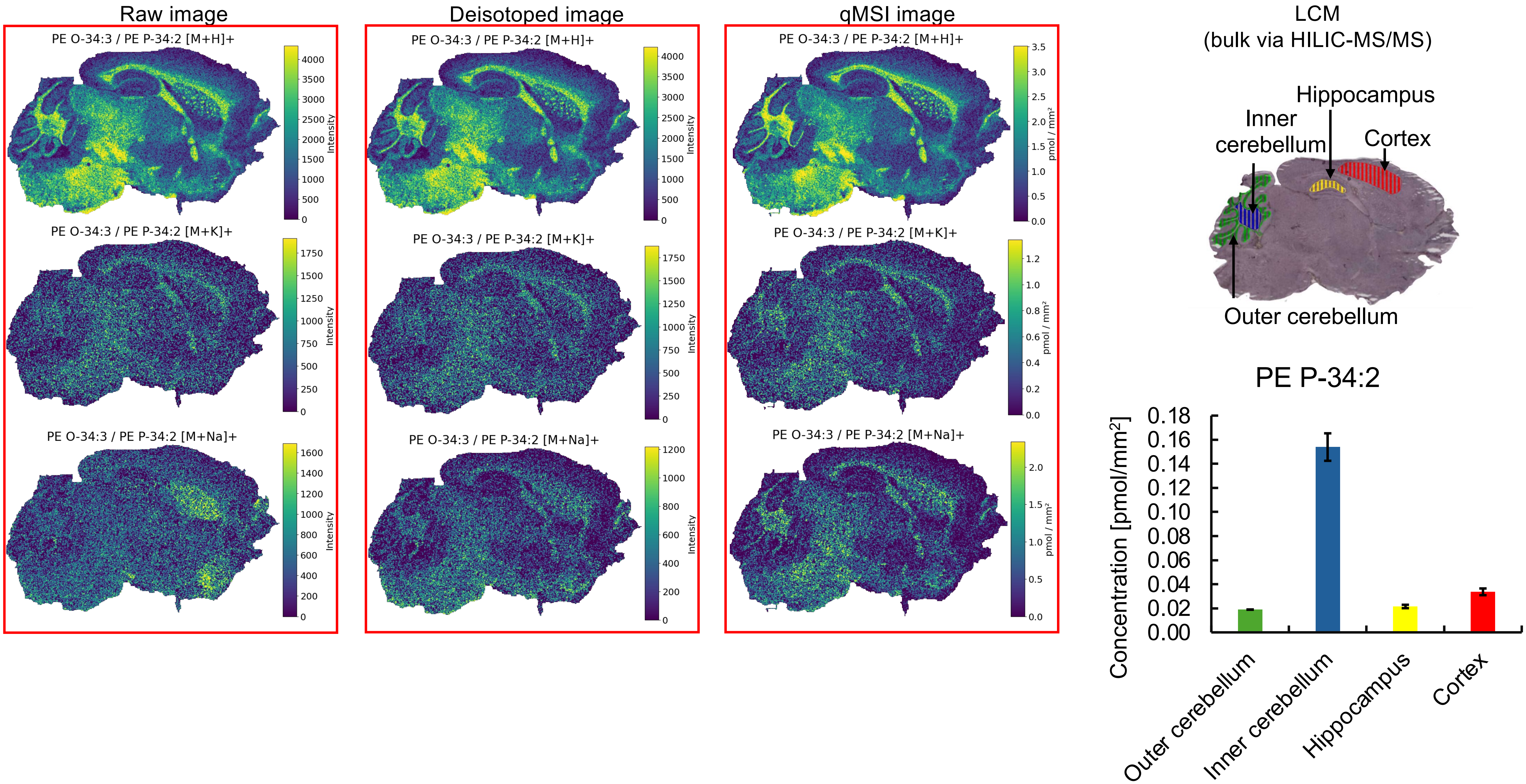


No bulk data for PE O-34:3.

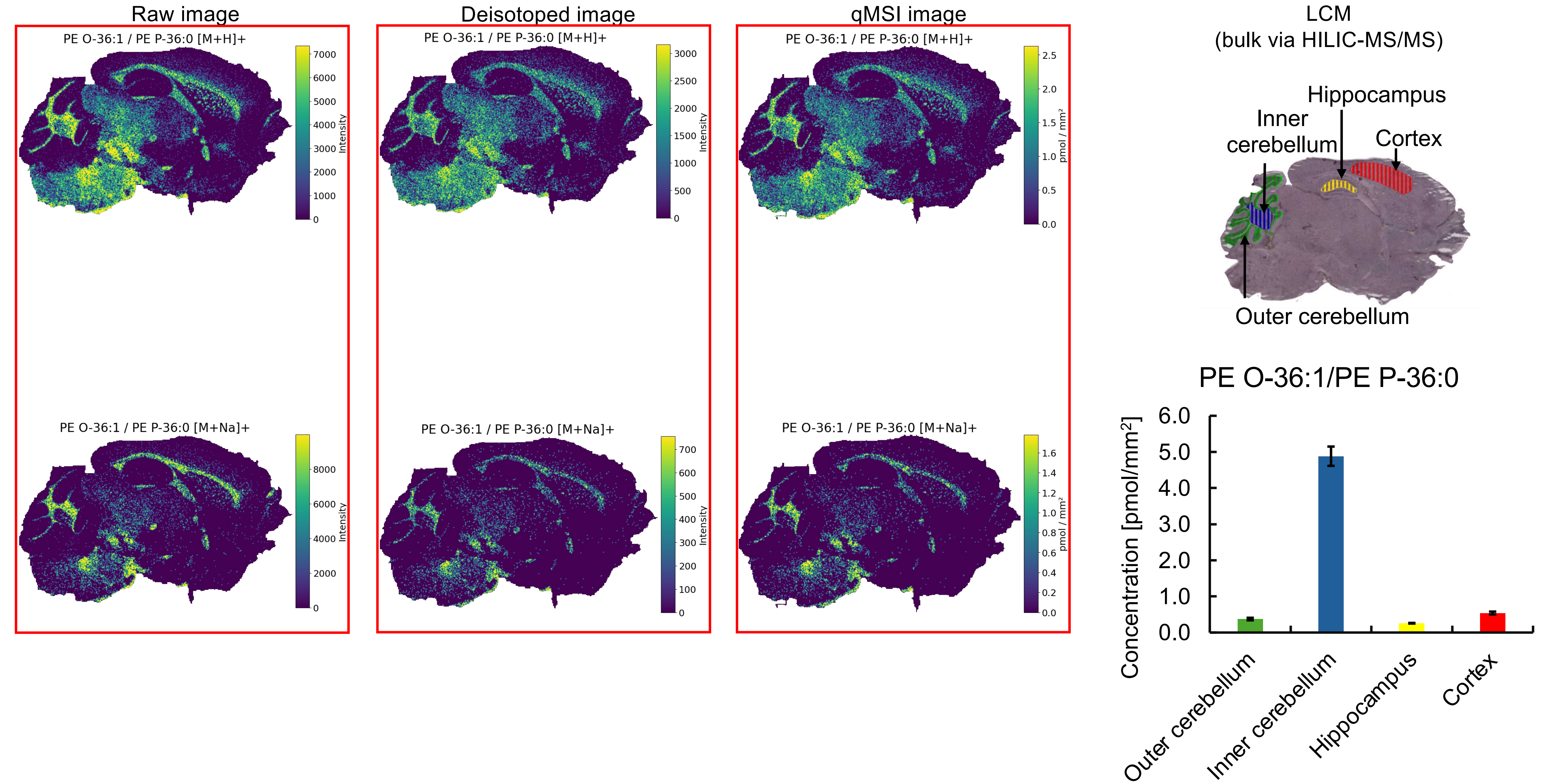


* Missing pixels for **[M+H]⁺, [M+Na]⁺** adducts (profile to centroid transformation).

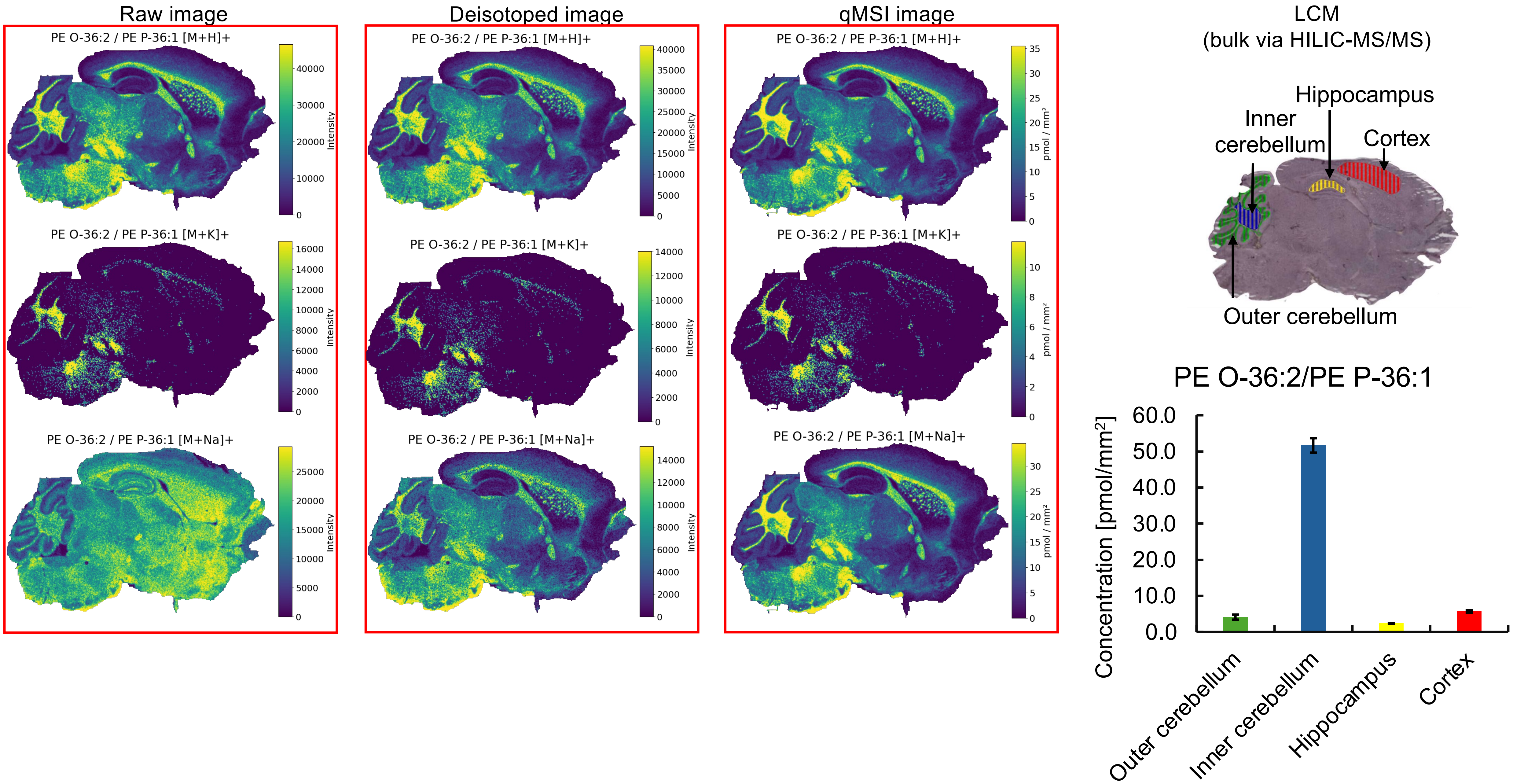


* Missing pixels for **[M+K]⁺** adducts (profile to centroid transformation).

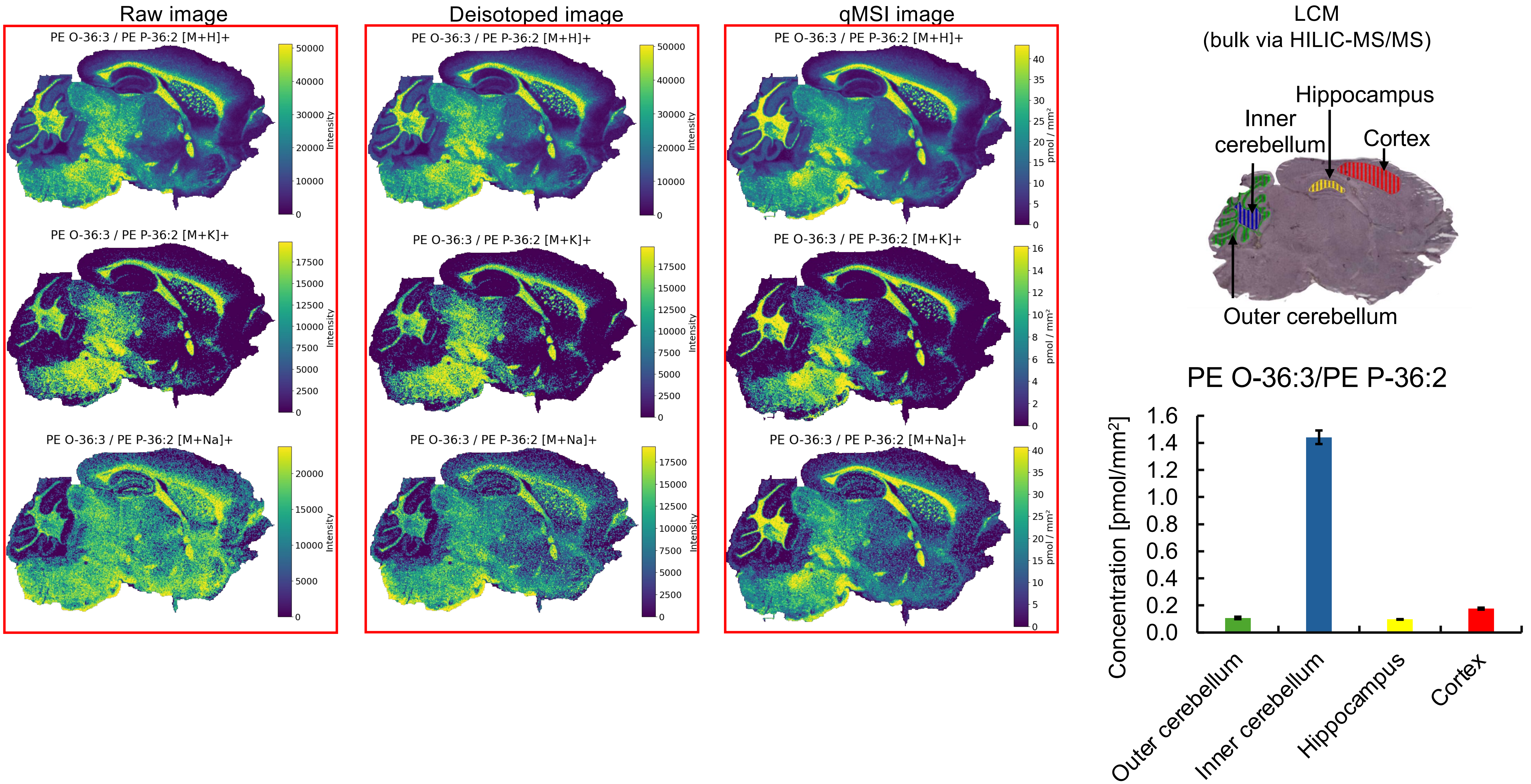


* Missing pixels for **[M+K]⁺**,**[M+Na]⁺** adducts (profile to centroid transformation).

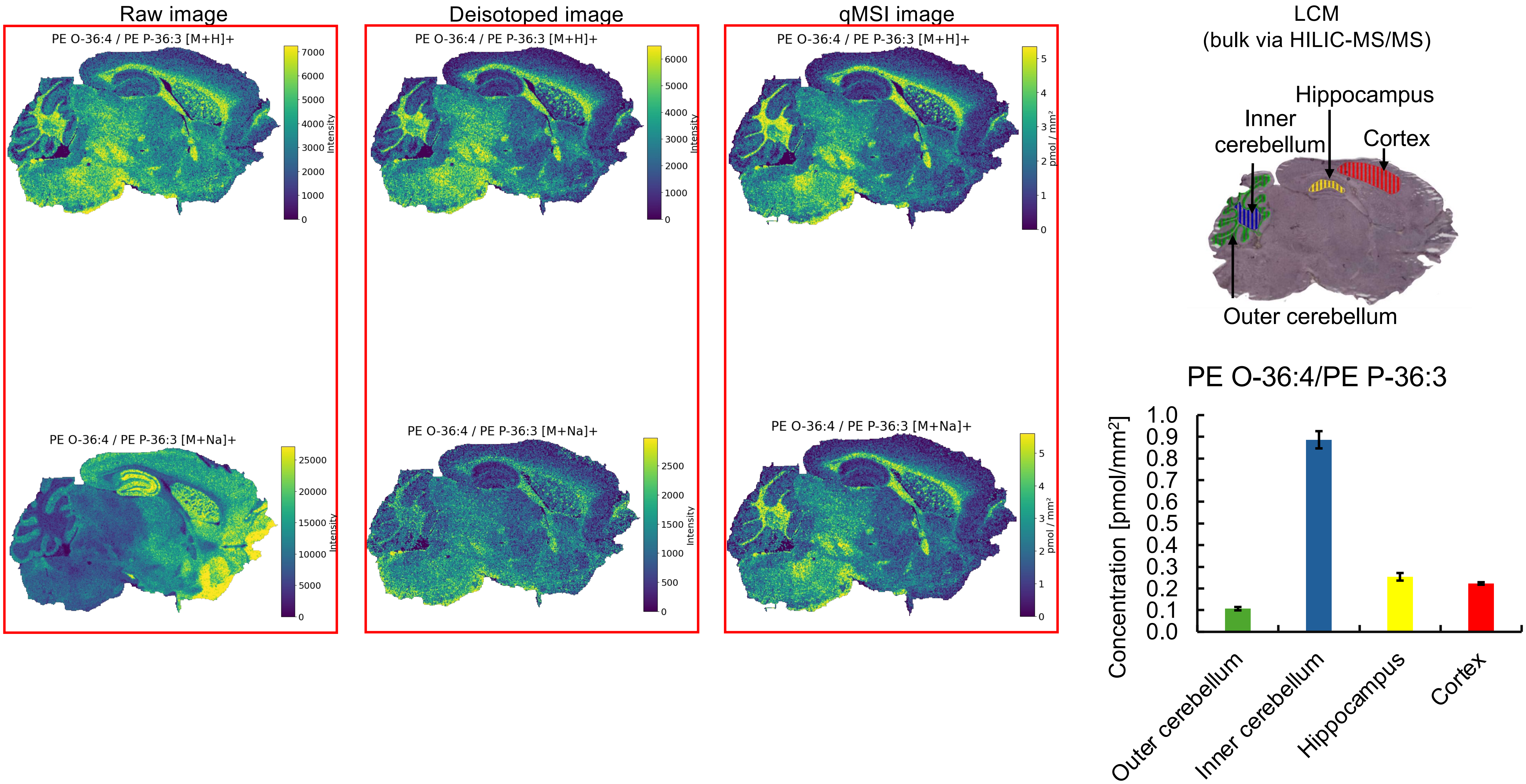

Raw image
Deisotoped image
qMSI image
LCM
(bulk via HILIC-MS/MS)
PE O-36:4 / PE P-36:3 [M+H]+
PE O-36:4 / PE P-36:3 [M+Na]+
Intensity
pmol / mm²
Hippocampus
Inner cerebellum
Cortex
Outer cerebellum
PE O-36:4/PE P-36:3
Concentration [pmol/mm²]
Outer cerebellum
Inner cerebellum
Hippocampus
Cortex

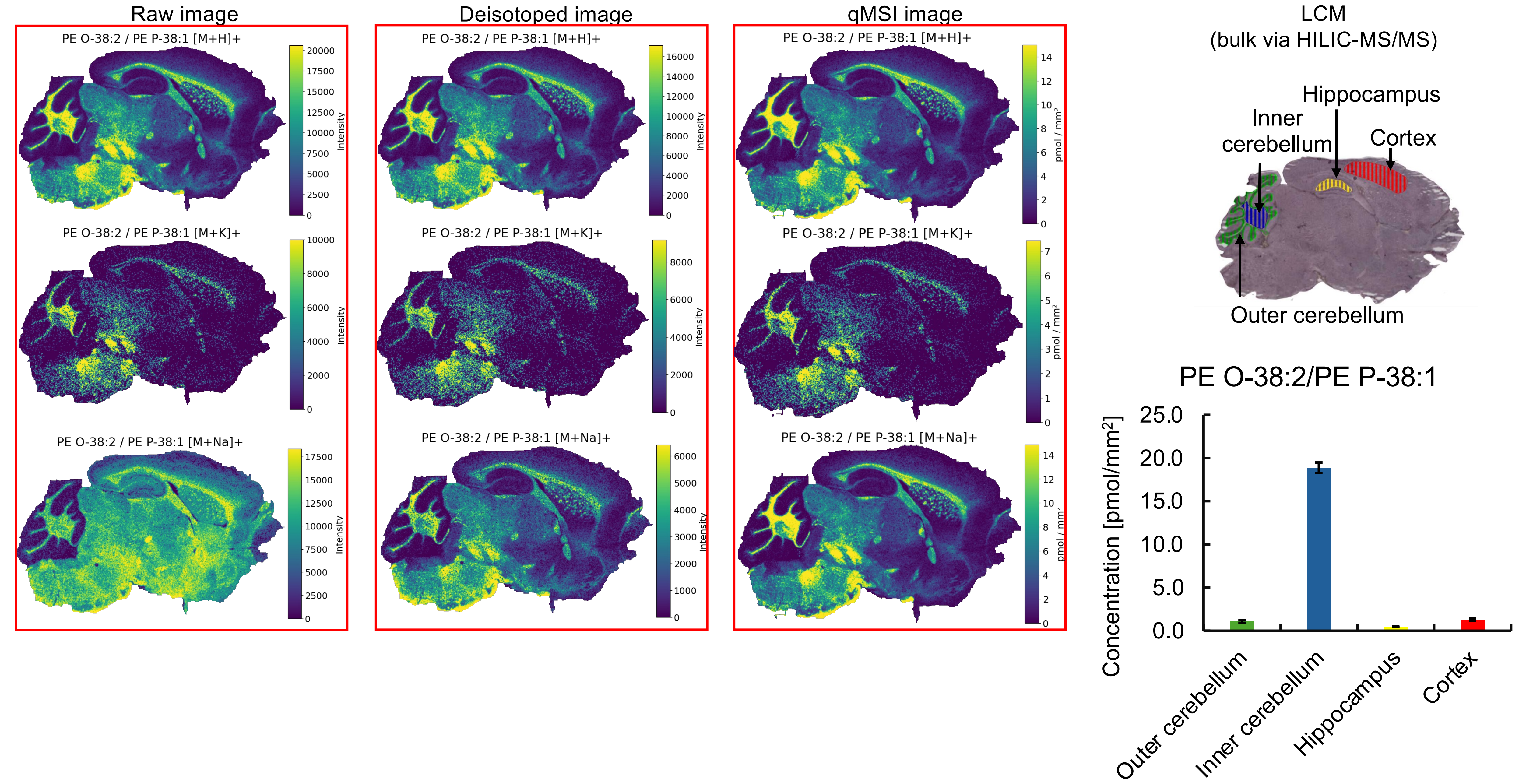


* Missing pixels for **[M+H]⁺, [M+K]⁺,[M+Na]⁺** adducts (profile to centroid transformation).

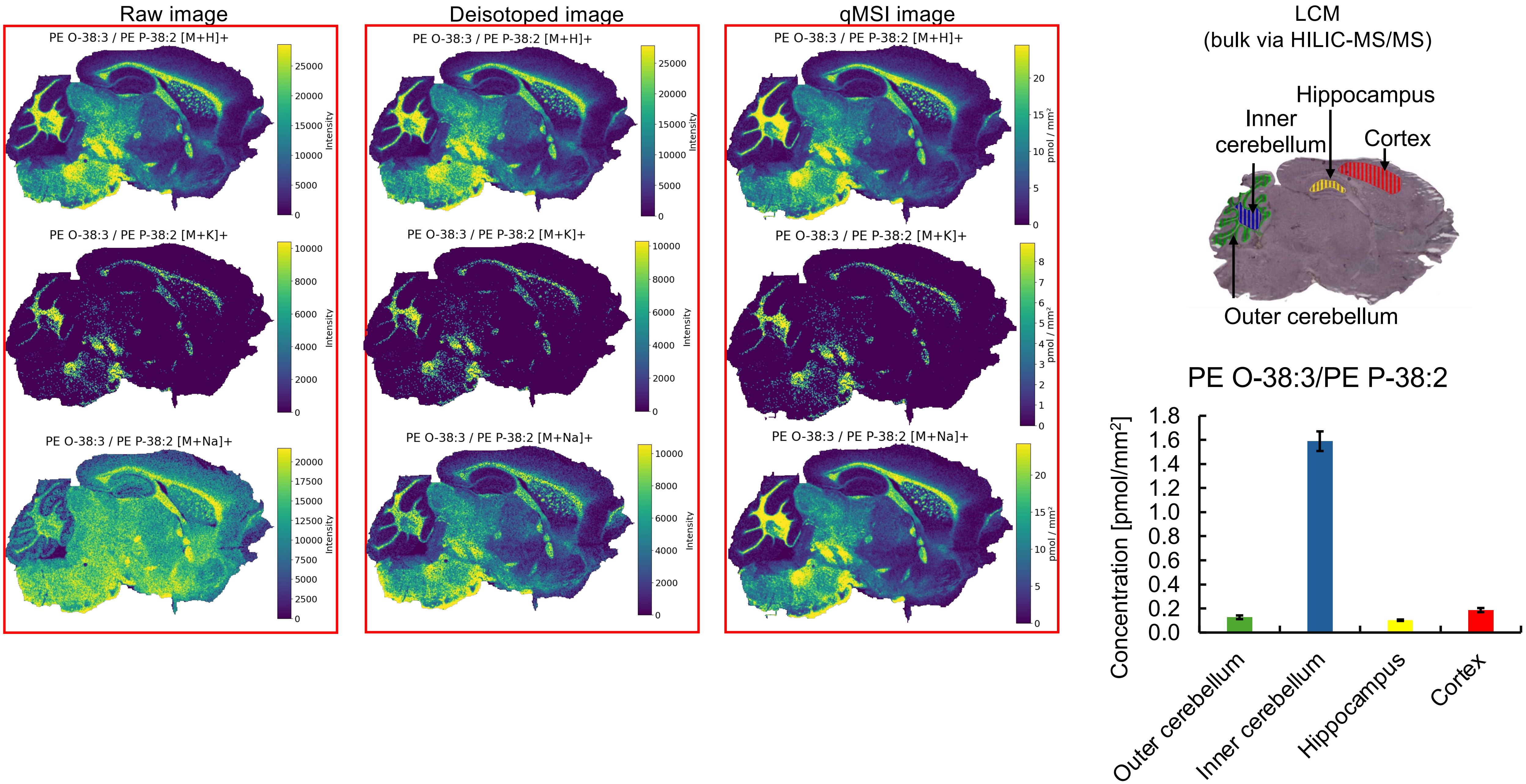


* Missing pixels for **[M+K]⁺** adducts (profile to centroid transformation).

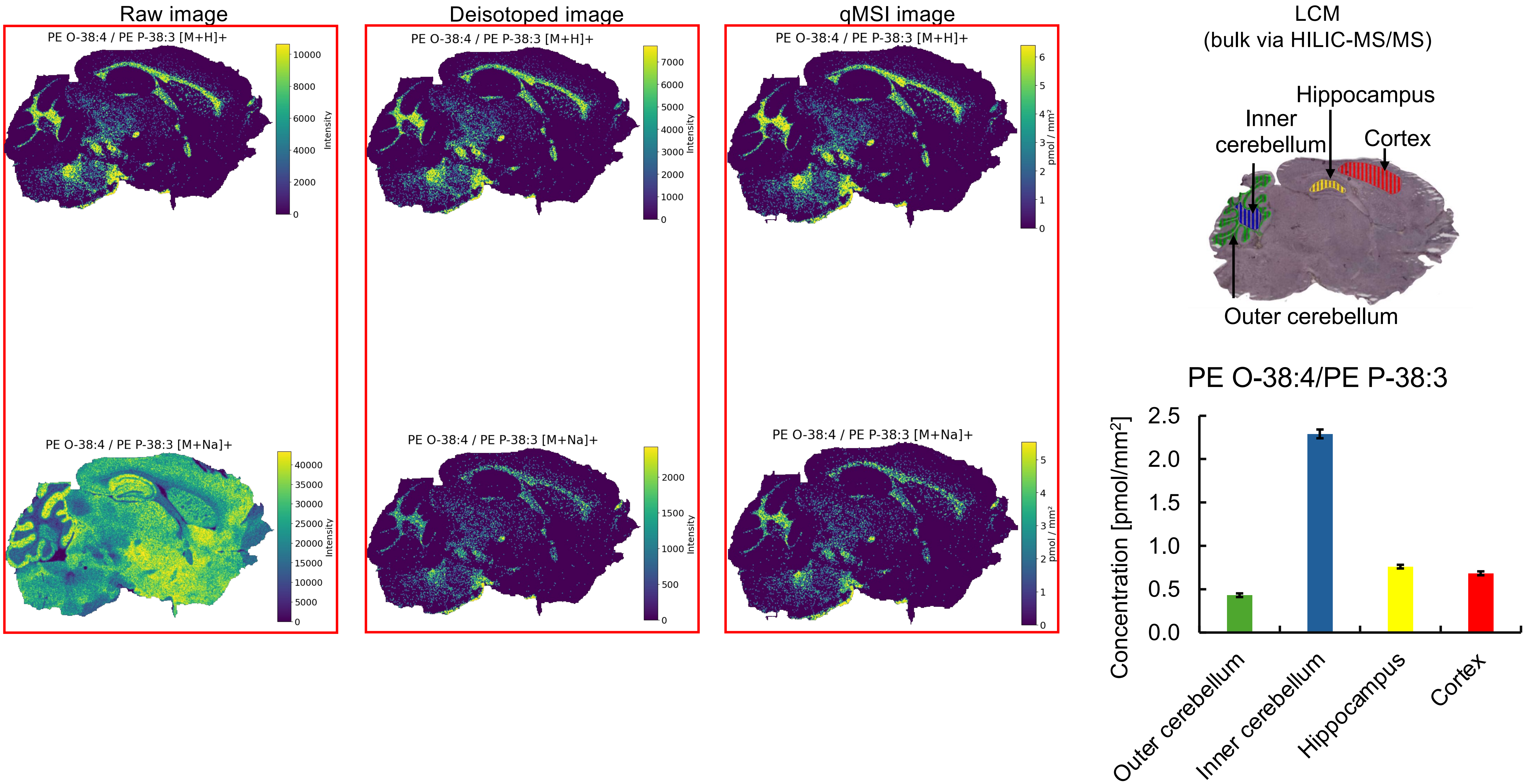


* Missing pixels for **[M+H]$^+$, [M+Na]$^+$** adducts (profile to centroid transformation).

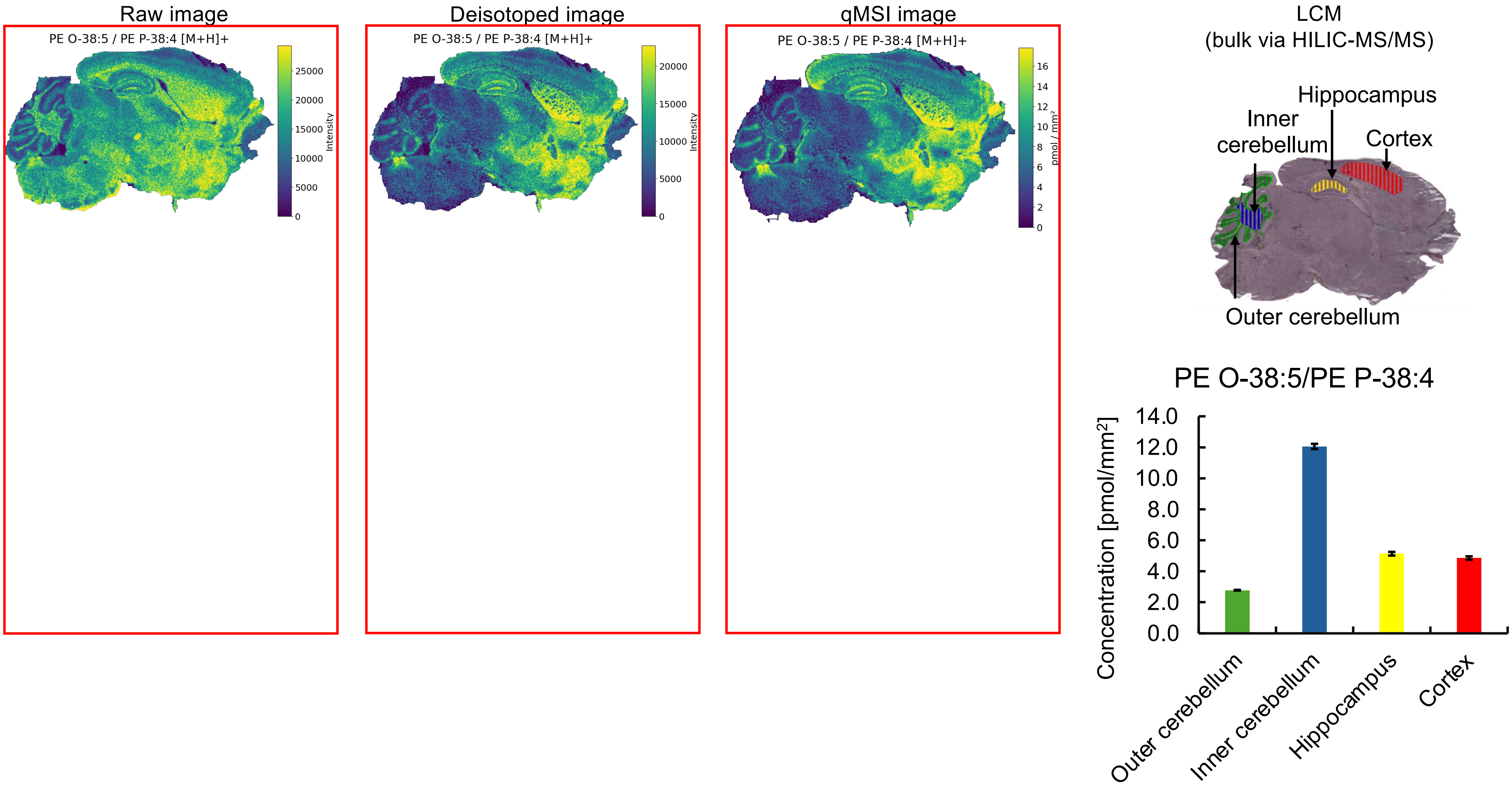


* Missing pixels for **[M+H]⁺**, **[M+K]⁺**, **[M+Na]⁺** adducts (profile to centroid transformation).

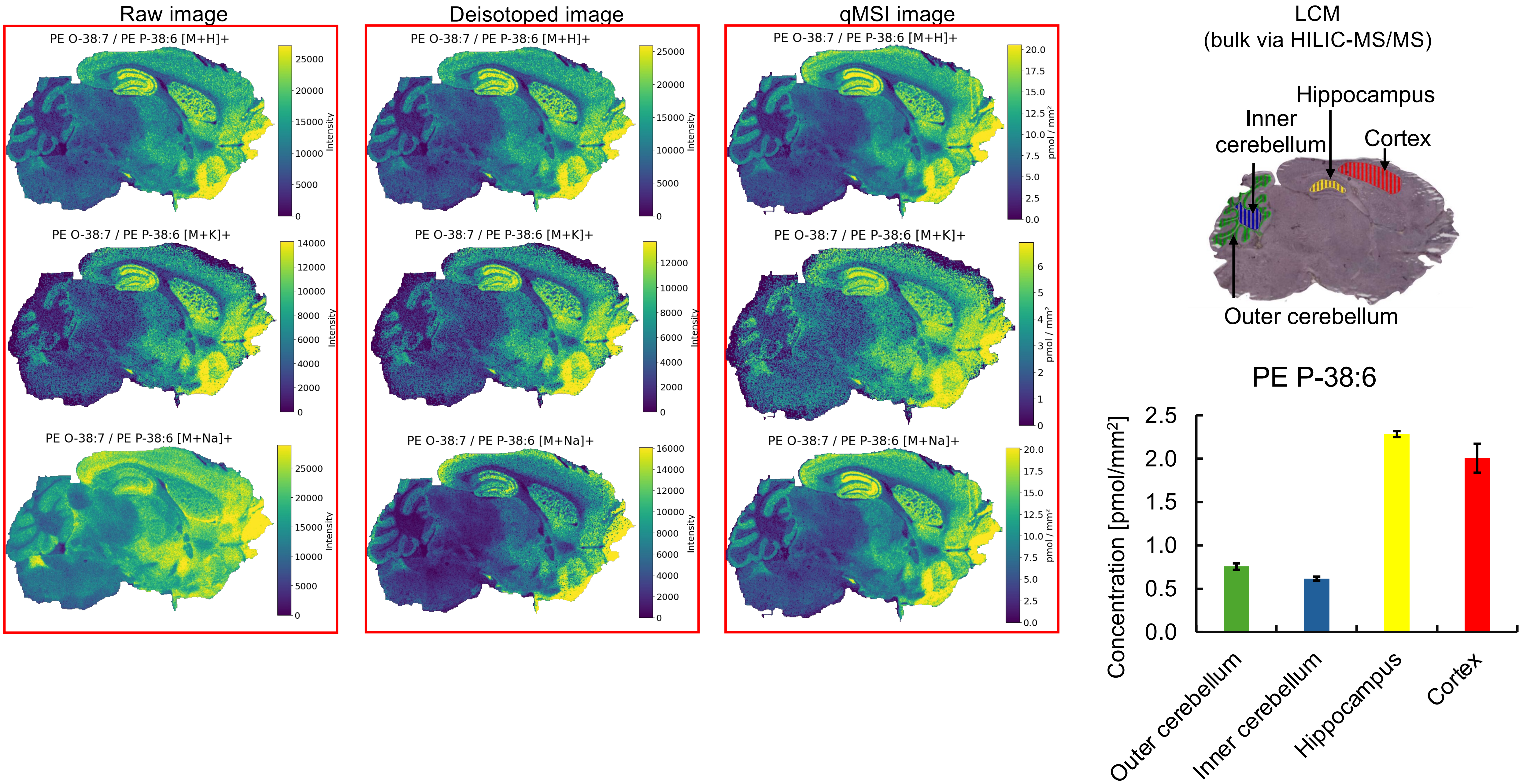


No bulk data for PE O-38:7.

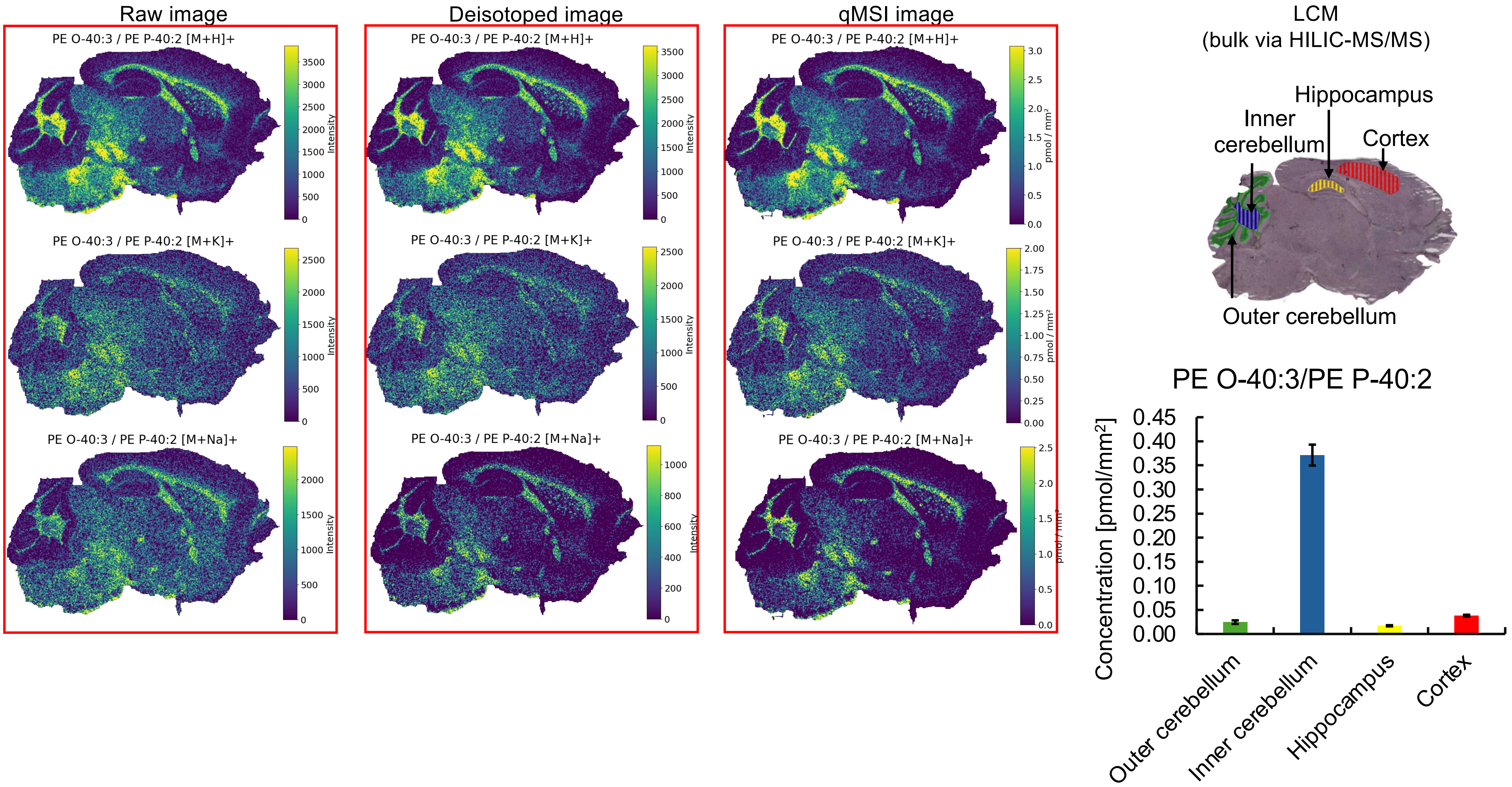


* Missing pixels for **[M+H]⁺, [M+K]⁺,[M+Na]⁺** adducts (profile to centroid transformation).

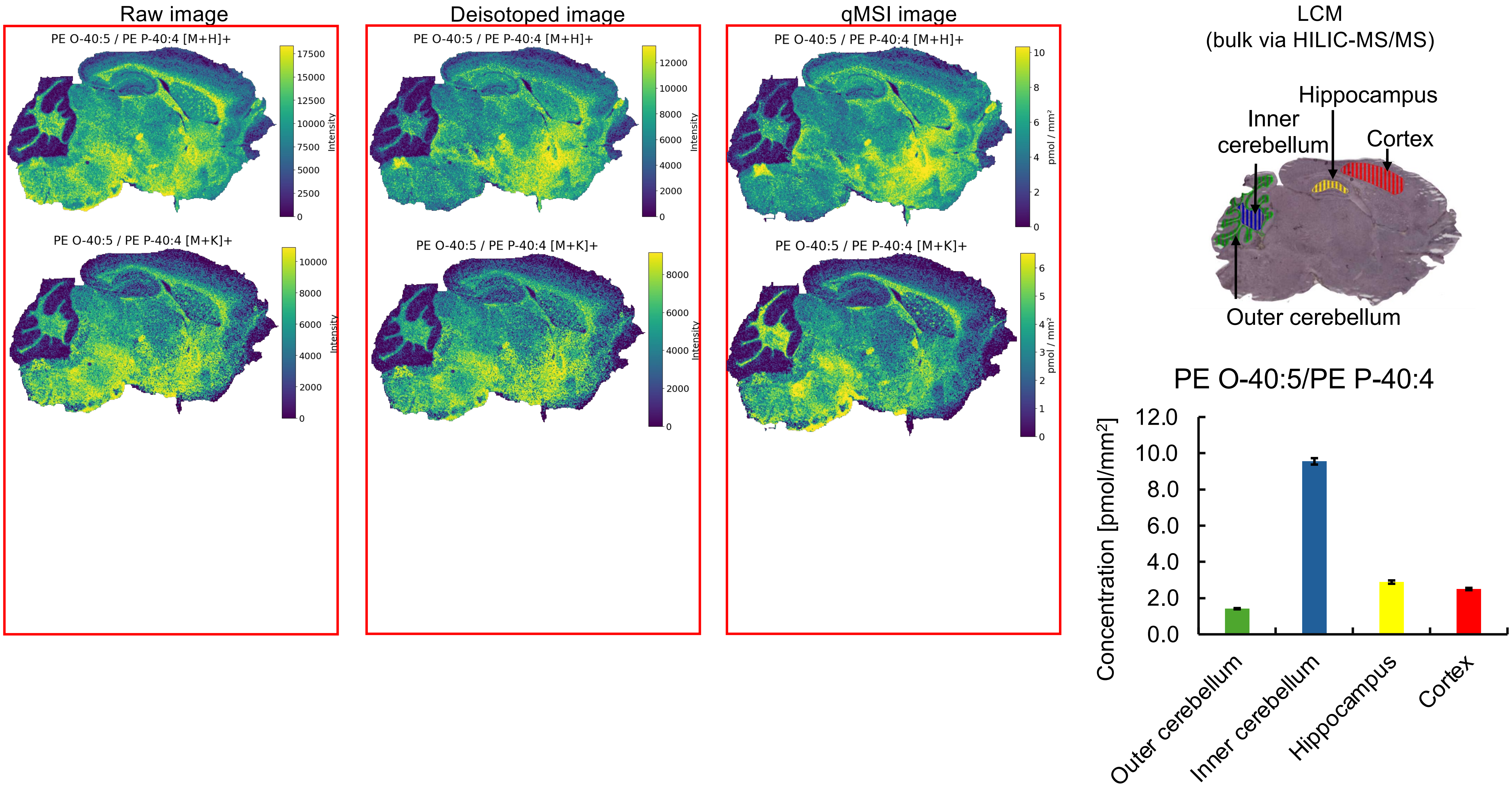

Raw image
Deisotoped image
qMSI image
LCM
(bulk via HILIC-MS/MS)
PE O-40:5 / PE P-40:4 [M+H]+
PE O-40:5 / PE P-40:4 [M+K]+
Intensity
pmol / mm²
Hippocampus
Inner cerebellum
Cortex
Outer cerebellum
PE O-40:5/PE P-40:4
Concentration [pmol/mm²]
Outer cerebellum
Inner cerebellum
Hippocampus
Cortex

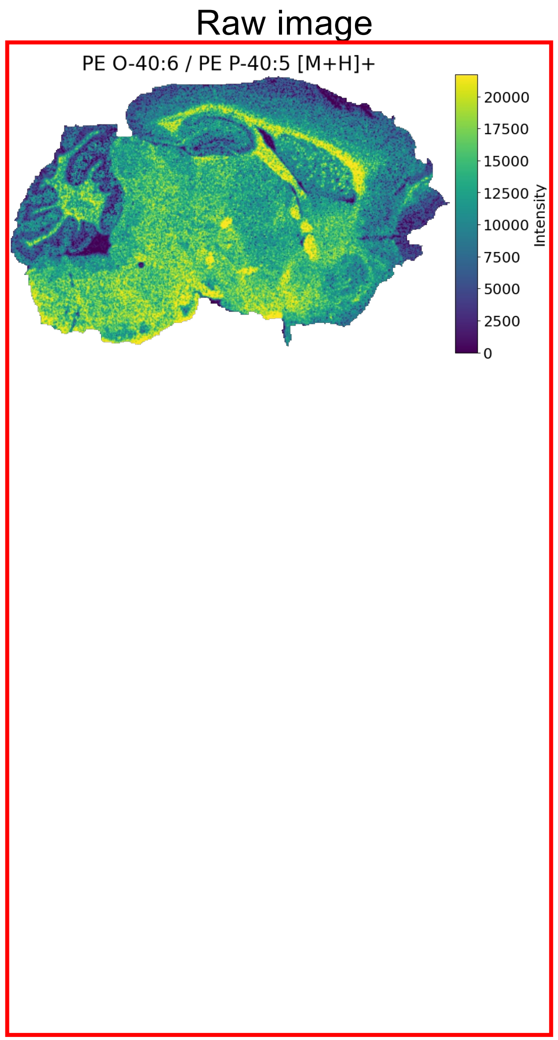


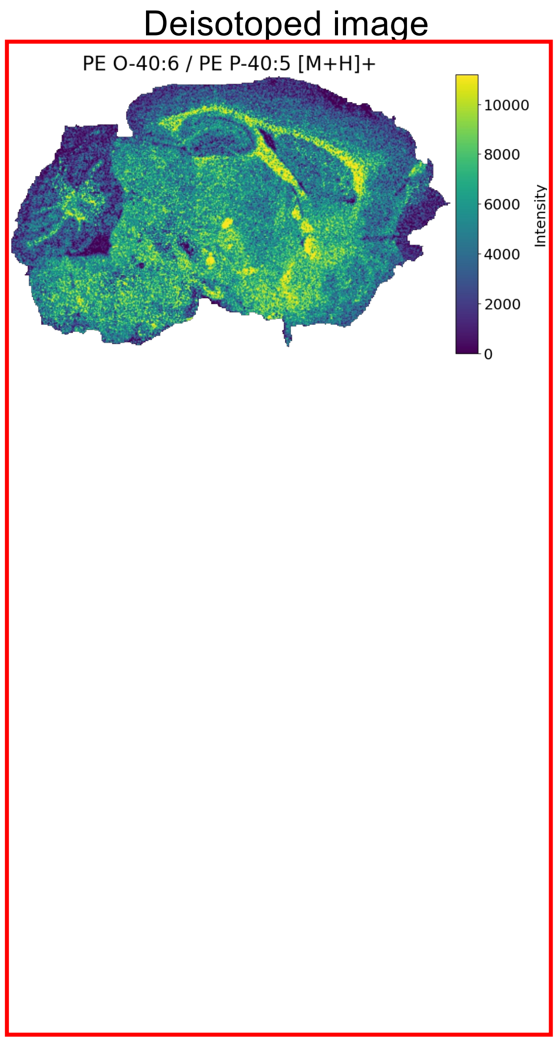


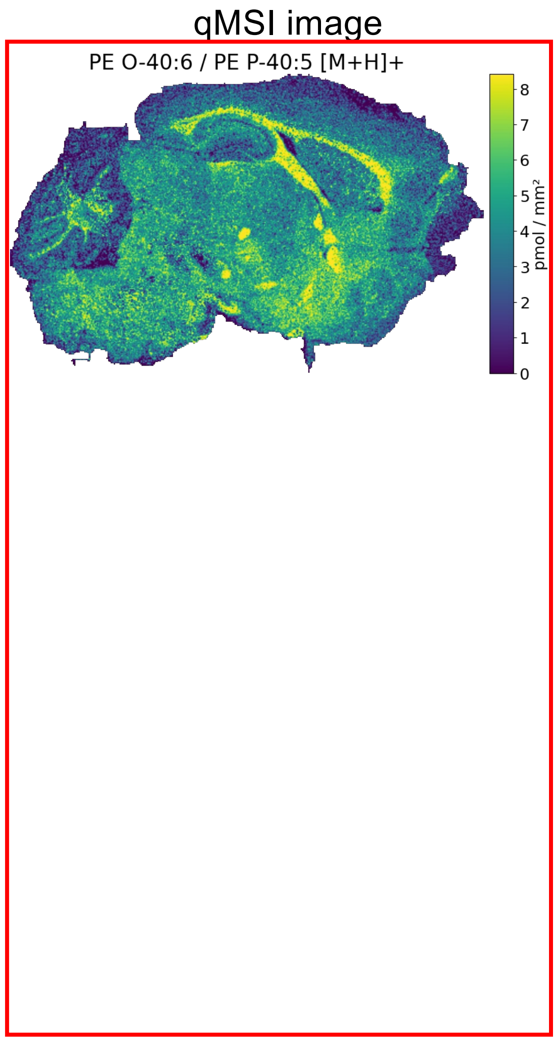


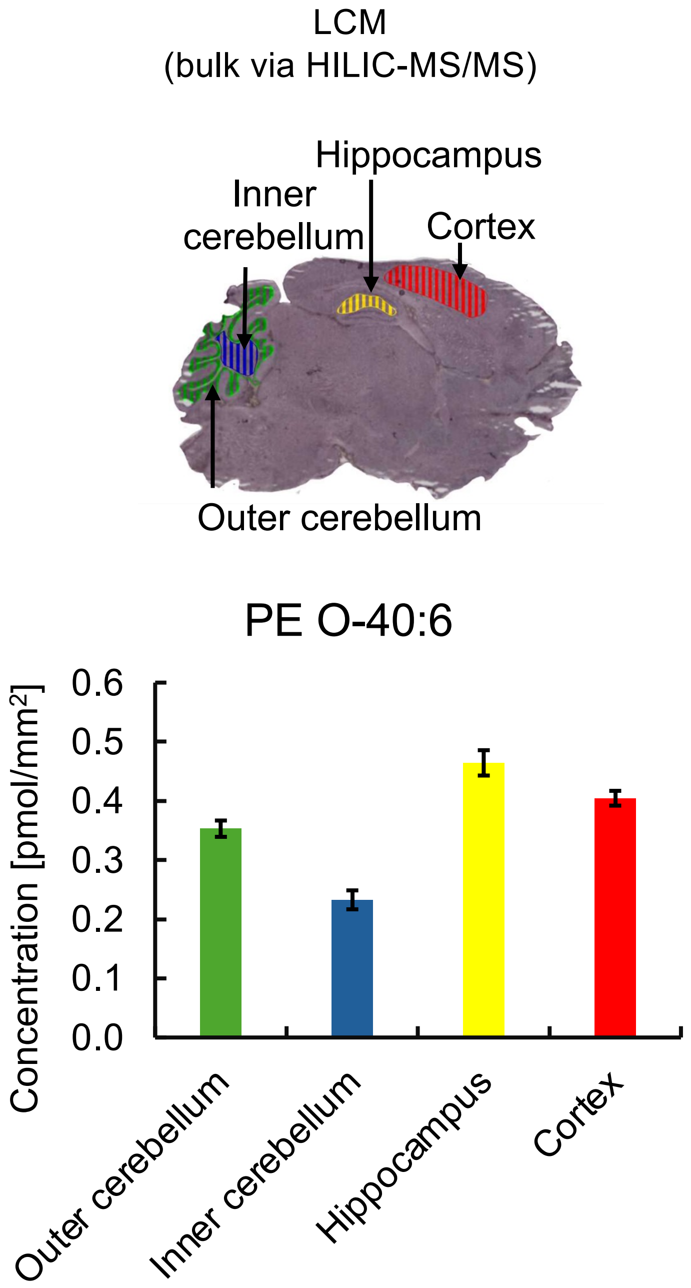


Bulk data for PE P-40:5 are <LOD (HILIC-MS/MS method).

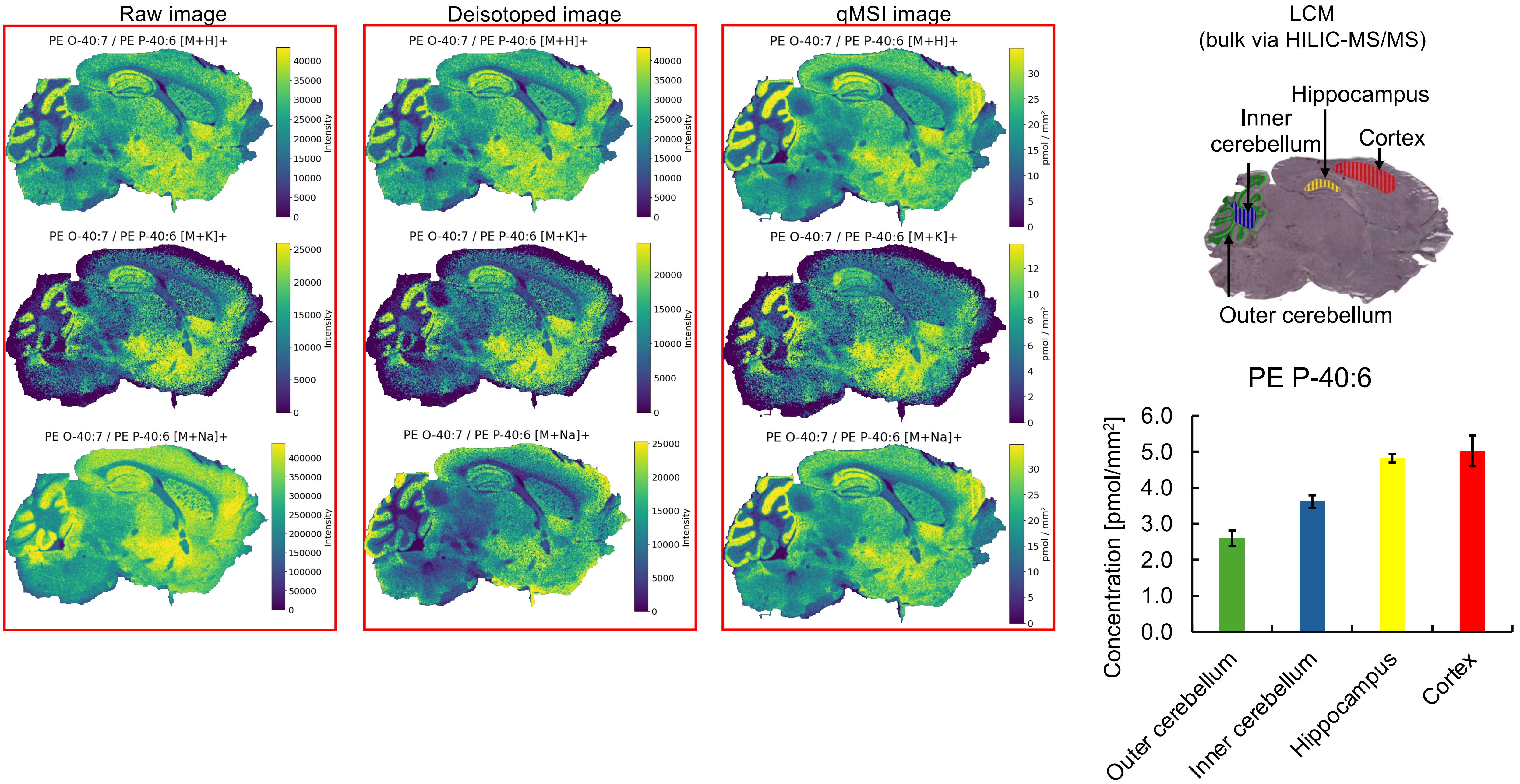


* Missing pixels for **[M+K]$^+$** adducts (profile to centroid transformation).

No bulk data for PE O-40:7.

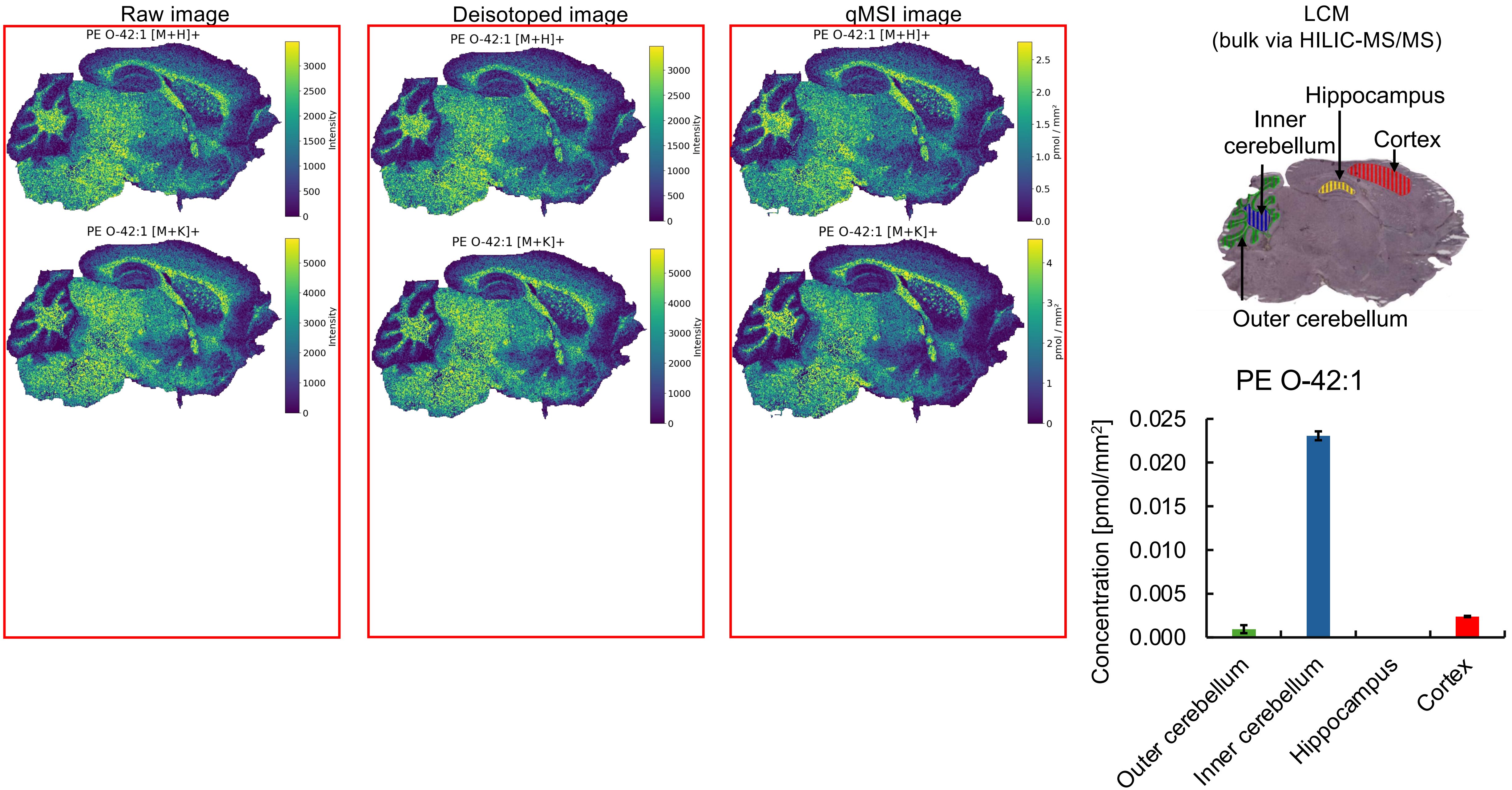

Raw image
PE O-42:1 [M+H]+
Intensity
PE O-42:1 [M+K]+
Deisotoped image
qMSI image
pmol / mm²
LCM
(bulk via HILIC-MS/MS)
Hippocampus
Inner cerebellum
Cortex
Outer cerebellum
PE O-42:1
Concentration [pmol/mm²]
0.025
0.020
0.015
0.010
0.005
0.000
Outer cerebellum
Inner cerebellum
Hippocampus
Cortex

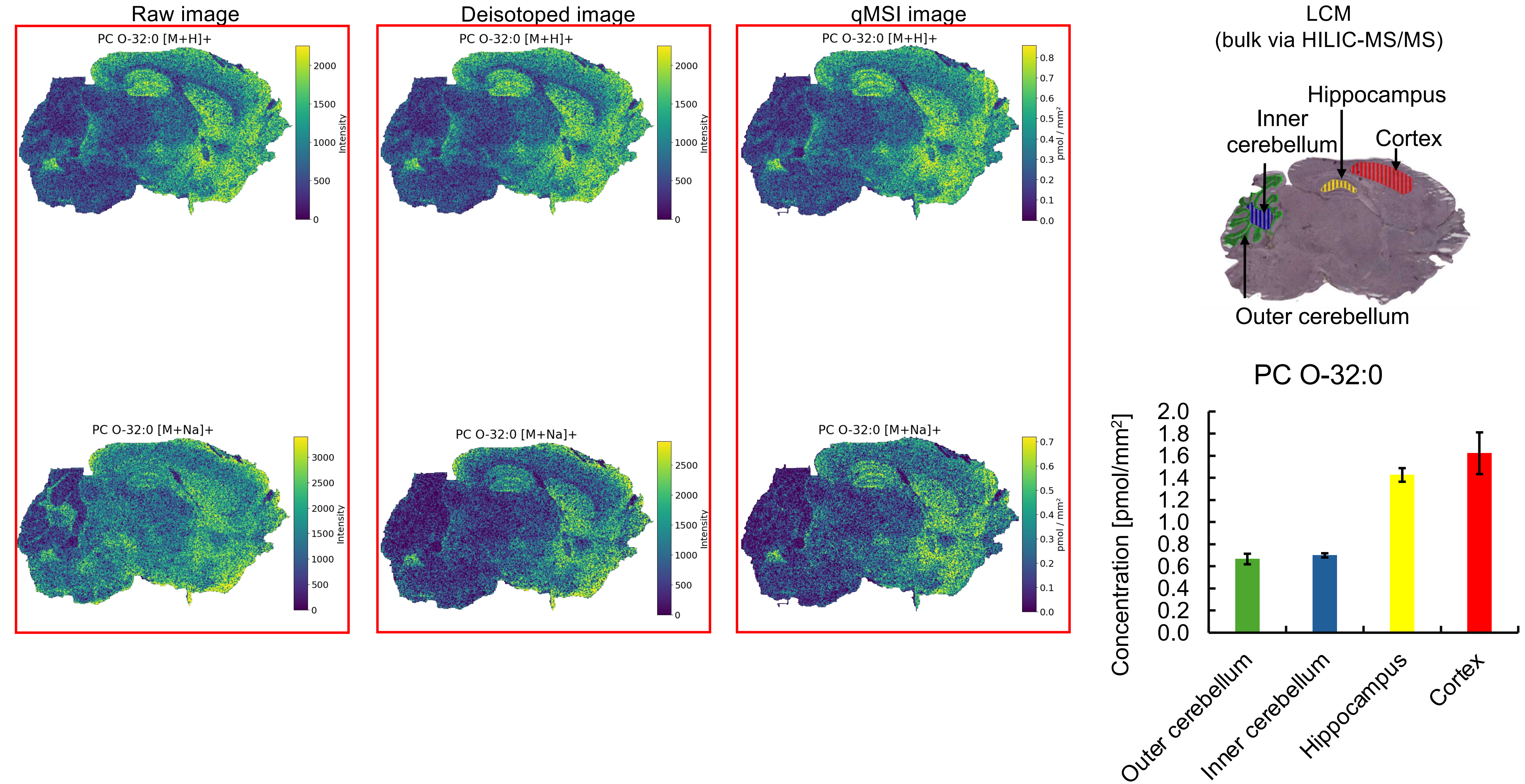

Raw image
Deisotoped image
qMSI image
PC O-32:0 [M+H]+
PC O-32:0 [M+Na]+
Intensity
pmol / mm²
LCM
(bulk via HILIC-MS/MS)
Hippocampus
Inner cerebellum
Cortex
Outer cerebellum
PC O-32:0
Concentration [pmol/mm²]
Outer cerebellum
Inner cerebellum
Hippocampus
Cortex

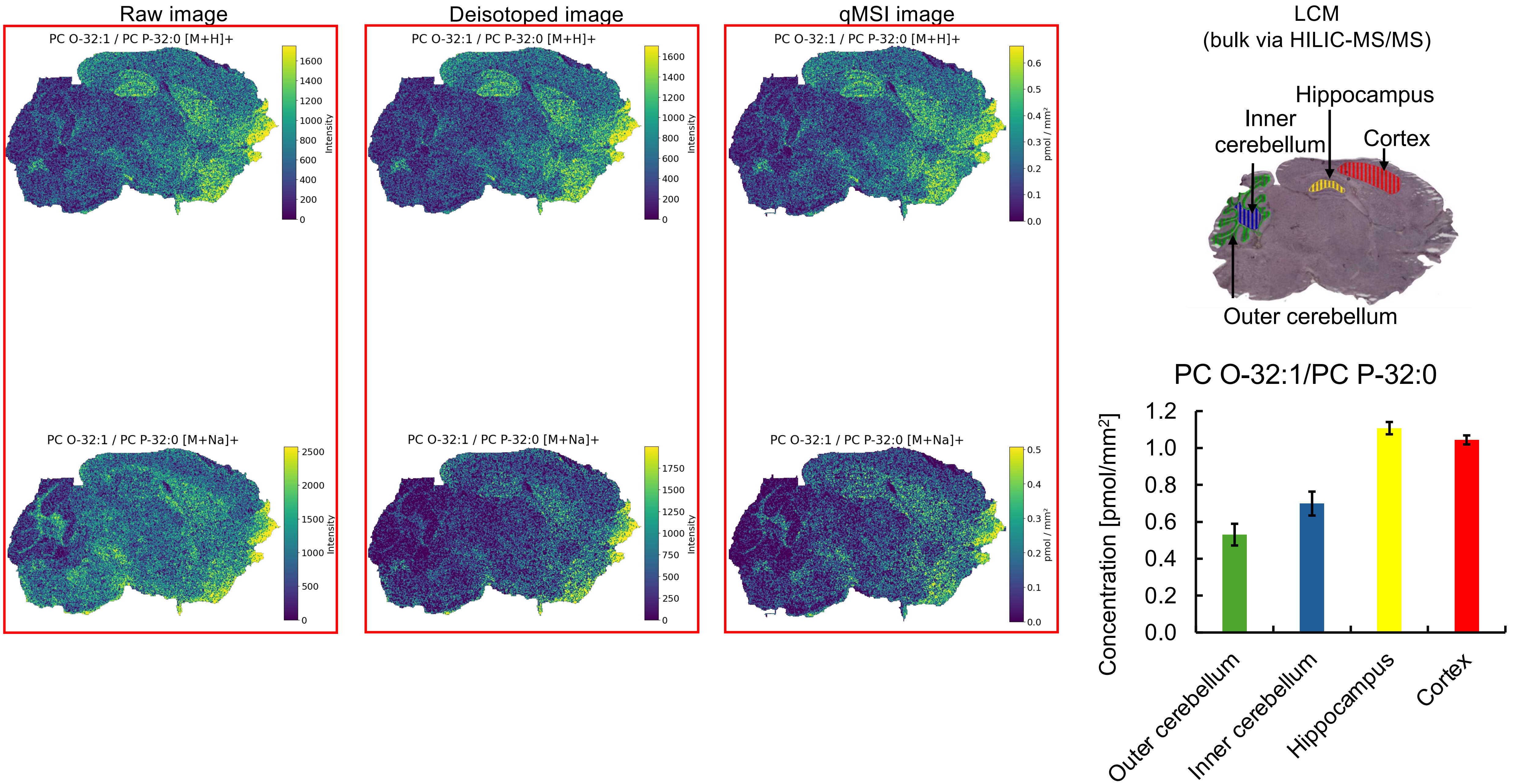

Raw image
Deisotoped image
qMSI image
PC O-32:1 / PC P-32:0 [M+H]+
PC O-32:1 / PC P-32:0 [M+Na]+
Intensity
pmol / mm²
LCM
(bulk via HILIC-MS/MS)
Hippocampus
Inner cerebellum
Cortex
Outer cerebellum
PC O-32:1/PC P-32:0
Concentration [pmol/mm²]
Outer cerebellum
Inner cerebellum
Hippocampus
Cortex

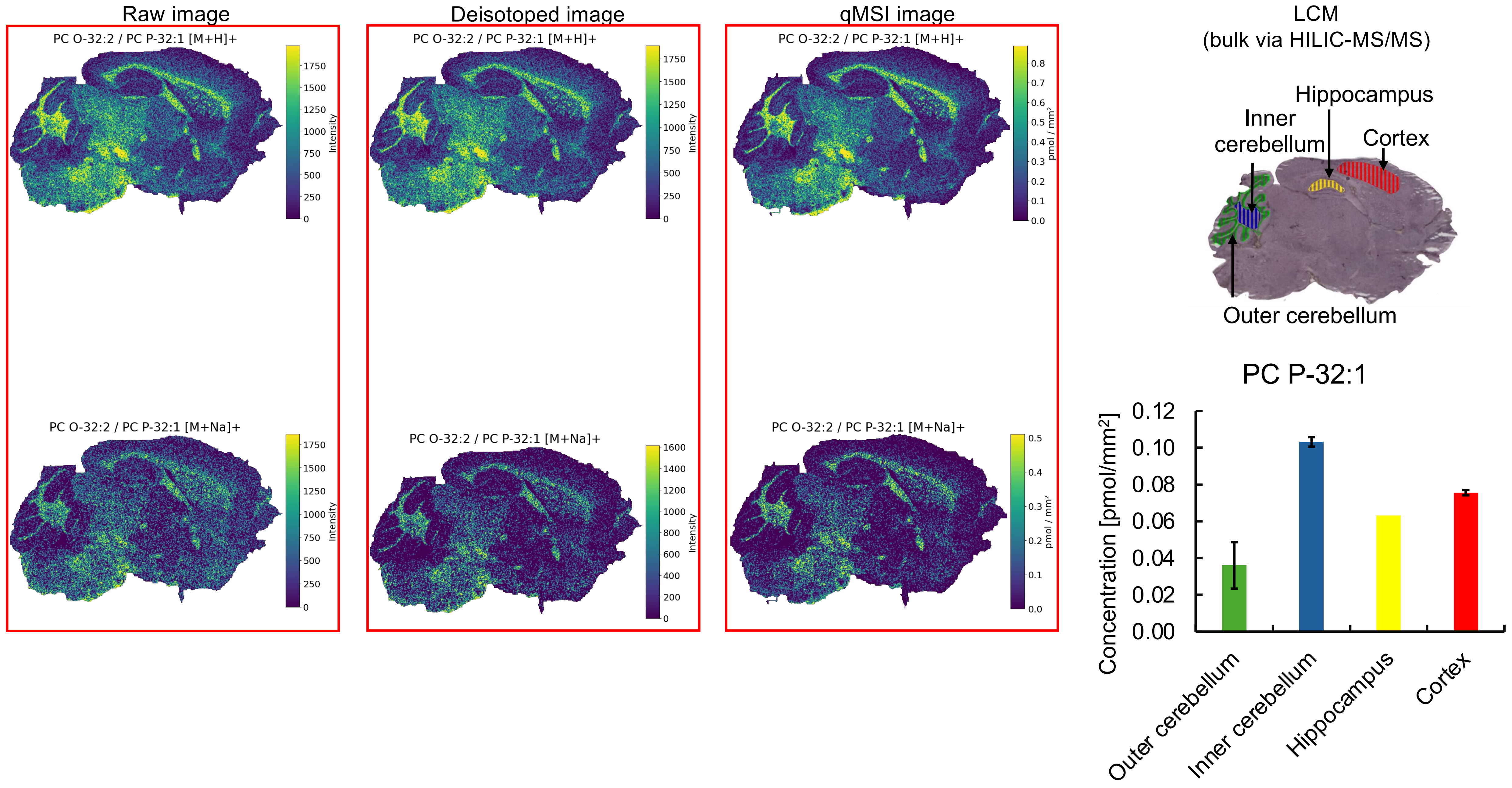


No bulk data for PC O-32:2.

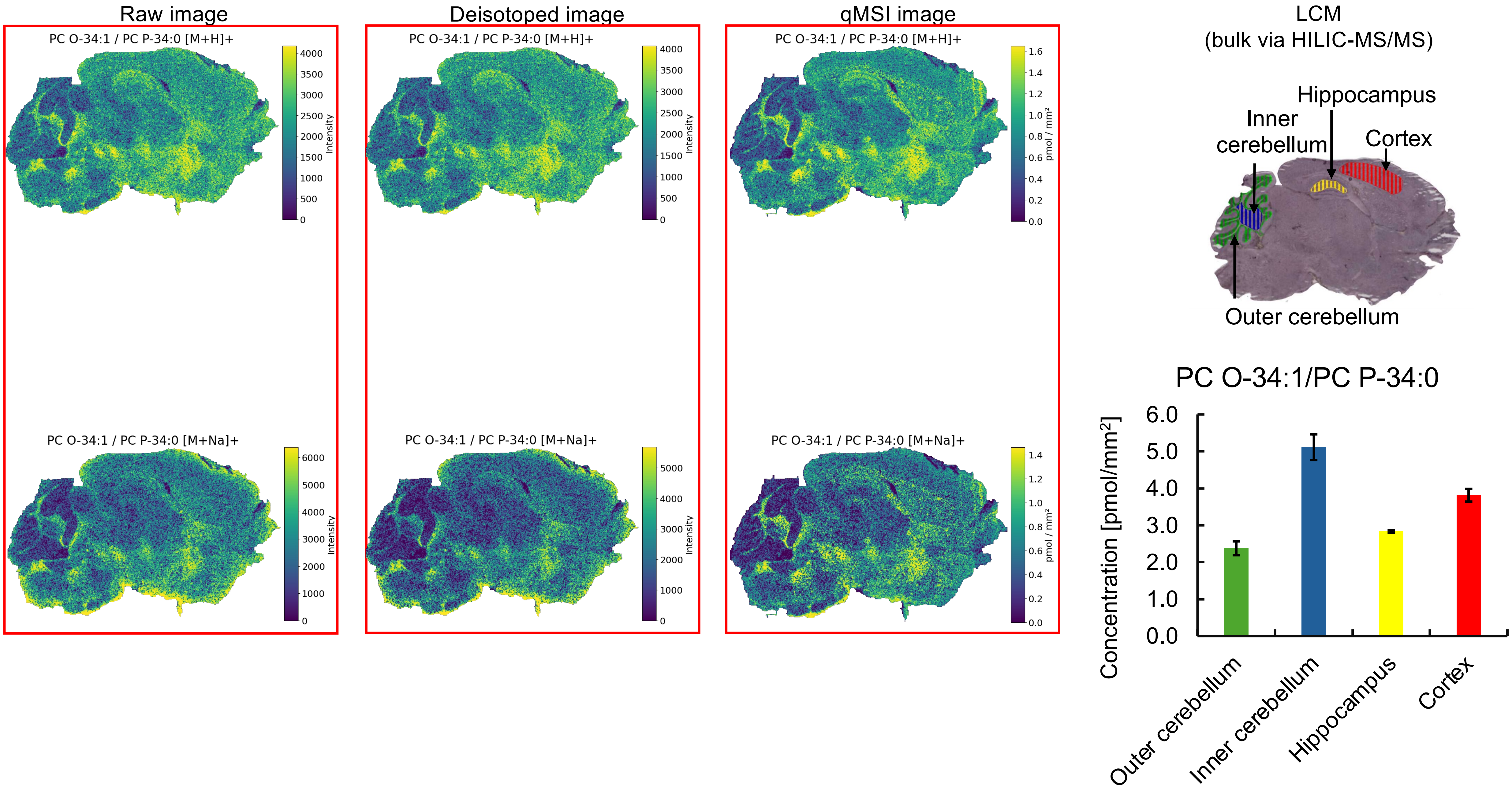

Raw image
Deisotoped image
qMSI image
PC O-34:1 / PC P-34:0 [M+H]+
PC O-34:1 / PC P-34:0 [M+Na]+
Intensity
pmol / mm²
LCM
(bulk via HILIC-MS/MS)
Hippocampus
Inner cerebellum
Cortex
Outer cerebellum
PC O-34:1/PC P-34:0
Concentration [pmol/mm²]
Outer cerebellum
Inner cerebellum
Hippocampus
Cortex

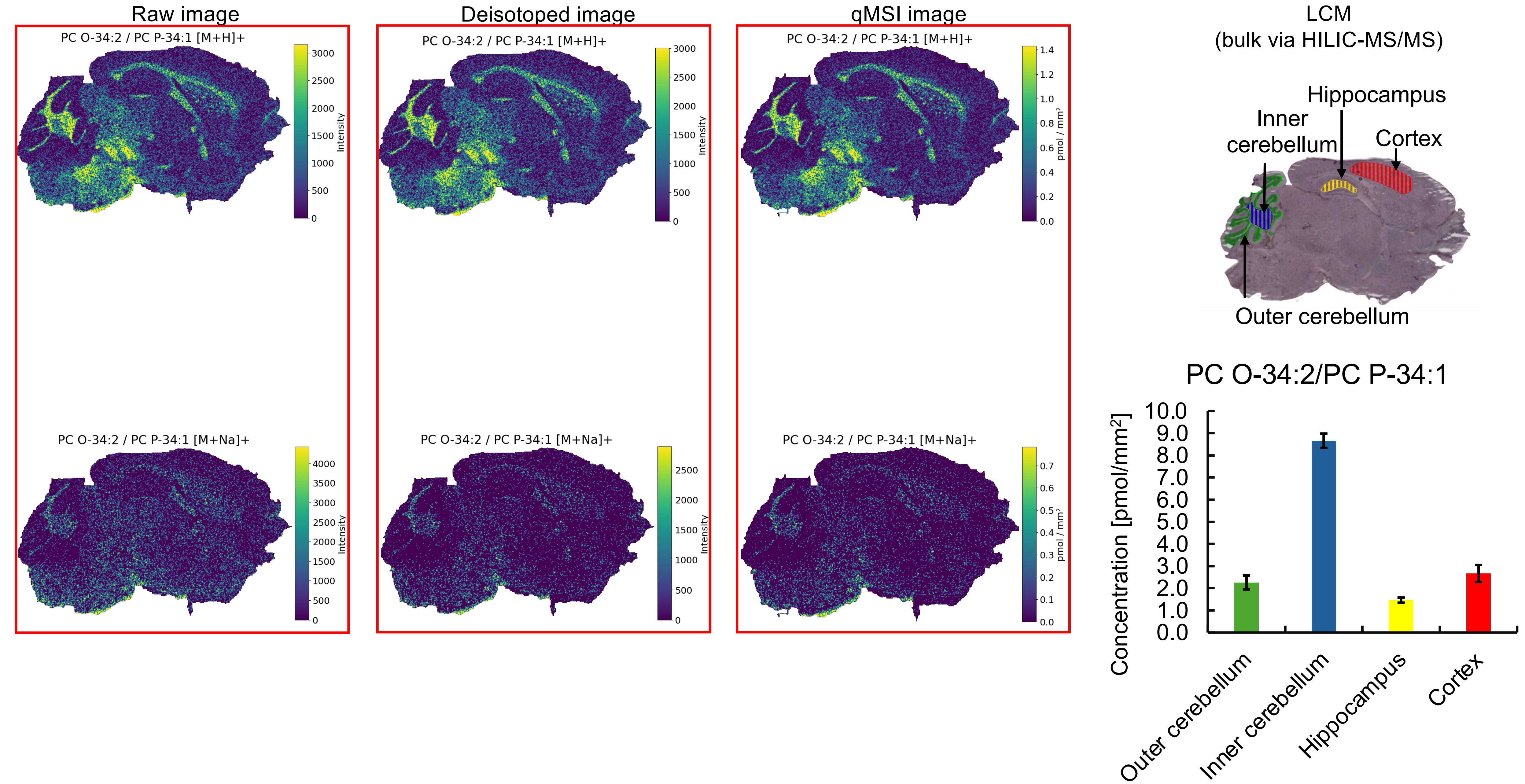

Raw image
Deisotoped image
qMSI image
PC O-34:2 / PC P-34:1 [M+H]+
PC O-34:2 / PC P-34:1 [M+Na]+
Intensity
pmol / mm²
LCM
(bulk via HILIC-MS/MS)
Hippocampus
Inner cerebellum
Cortex
Outer cerebellum
PC O-34:2/PC P-34:1
Concentration [pmol/mm²]
Outer cerebellum
Inner cerebellum
Hippocampus
Cortex

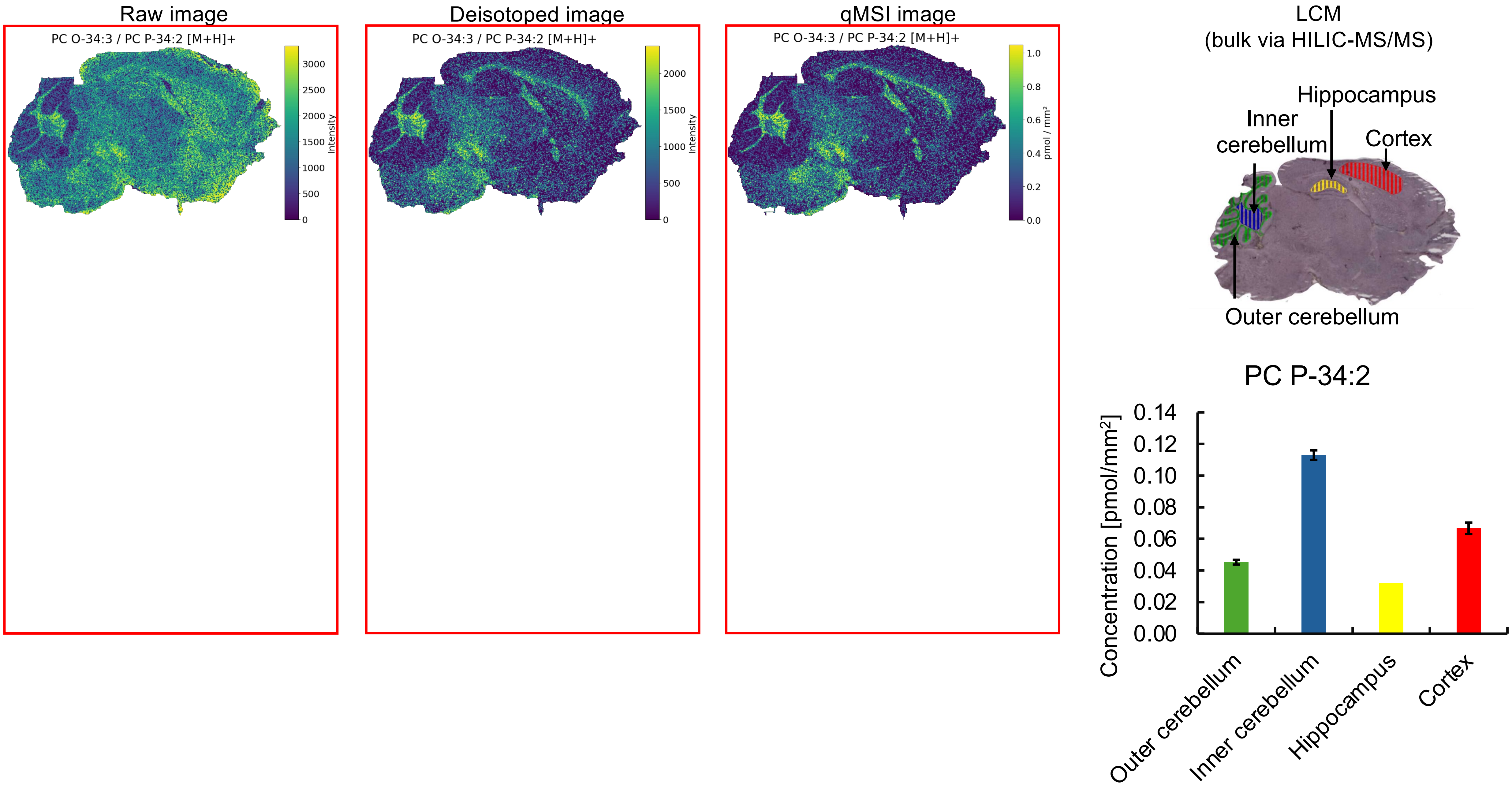


No bulk data for PC O-34:3.

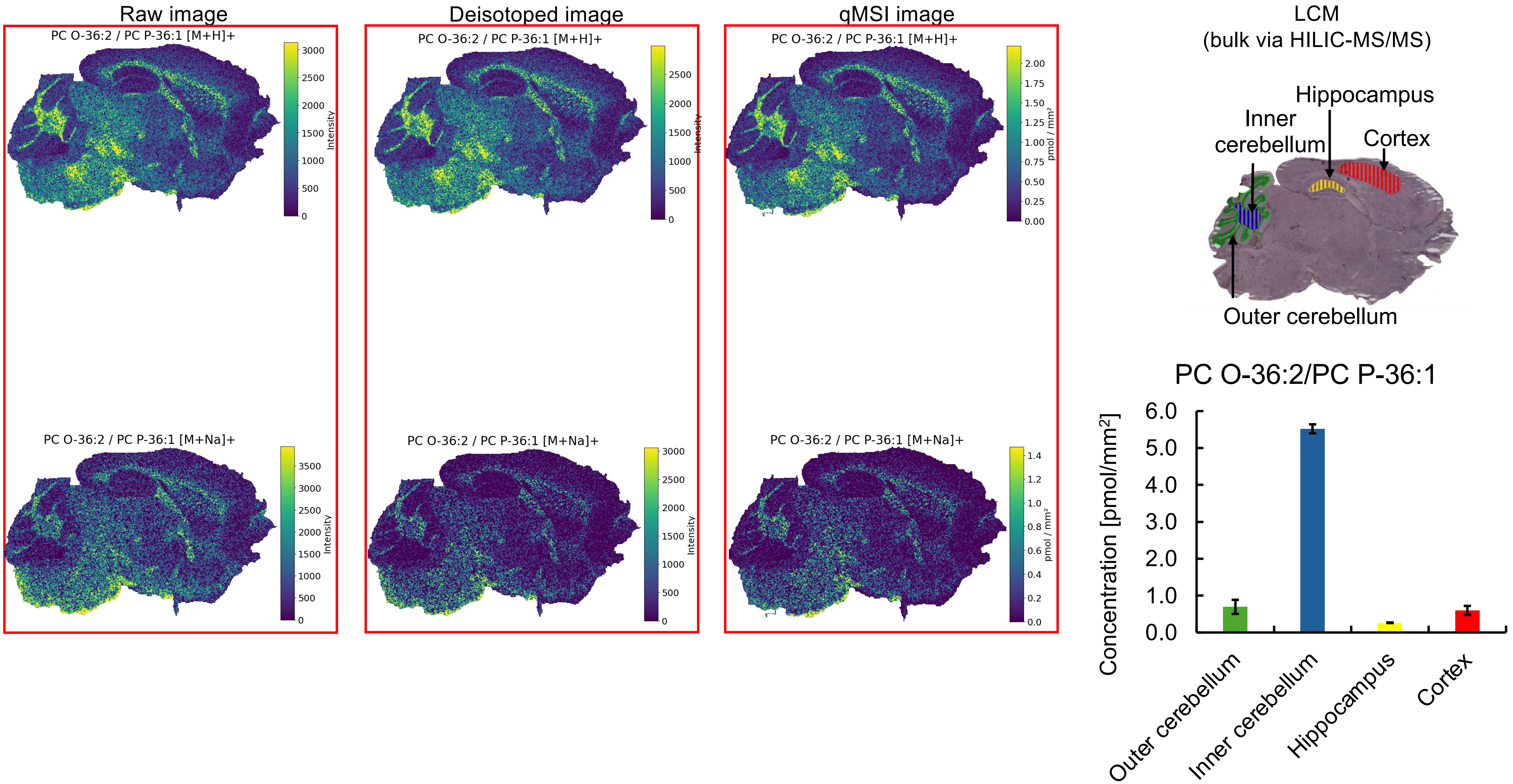
Raw image
Deisotoped image
qMSI image
PC O-36:2 / PC P-36:1 [M+H]+
PC O-36:2 / PC P-36:1 [M+Na]+
Intensity
pmol / mm²
LCM
(bulk via HILIC-MS/MS)
Hippocampus
Inner cerebellum
Cortex
Outer cerebellum
PC O-36:2/PC P-36:1
Concentration [pmol/mm²]
Outer cerebellum
Inner cerebellum
Hippocampus
Cortex

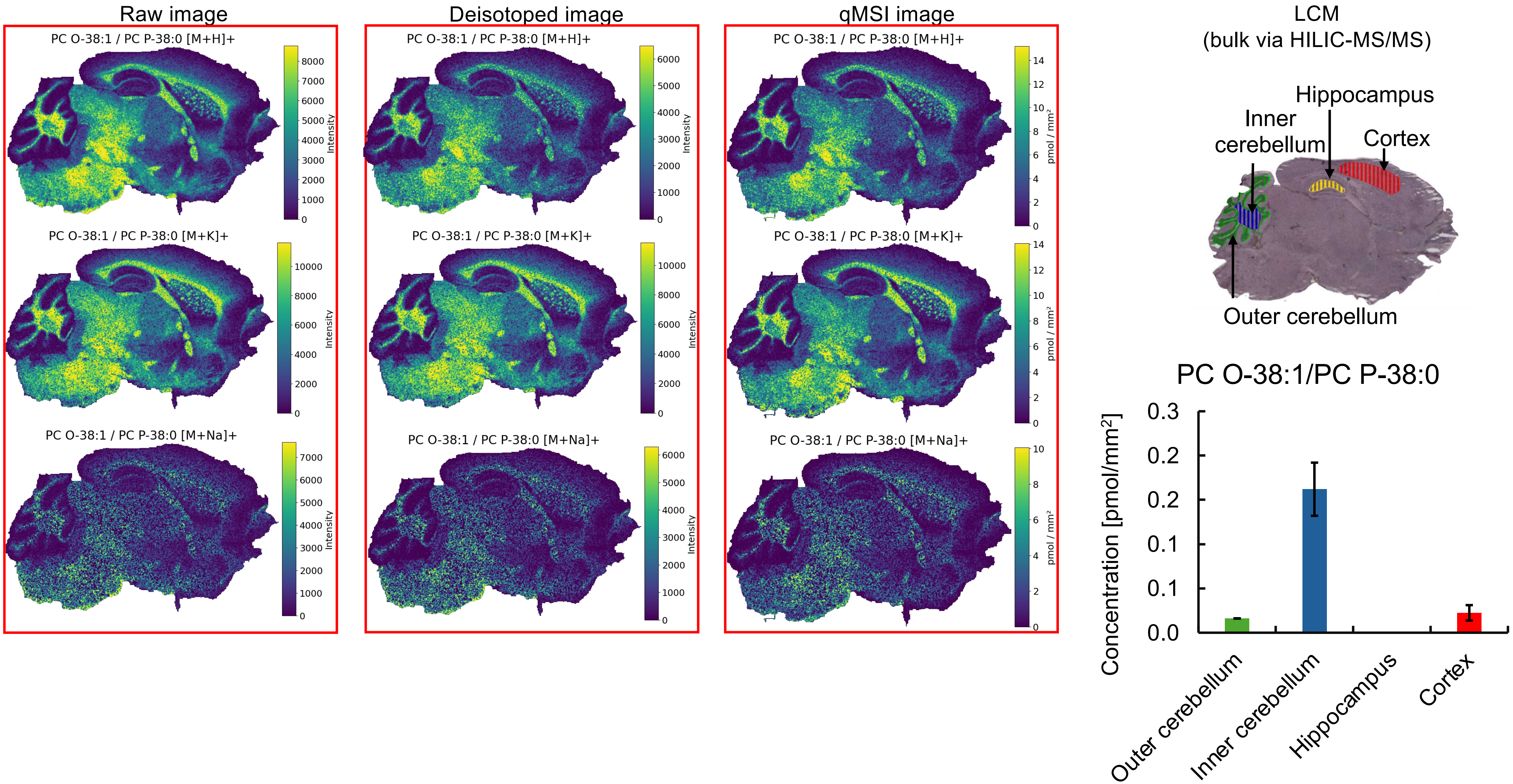
Raw image
Deisotoped image
qMSI image
LCM
(bulk via HILIC-MS/MS)
PC O-38:1 / PC P-38:0 [M+H]+
PC O-38:1 / PC P-38:0 [M+K]+
PC O-38:1 / PC P-38:0 [M+Na]+
Intensity
pmol / mm²
Hippocampus
Inner cerebellum
Cortex
Outer cerebellum
PC O-38:1/PC P-38:0
Concentration [pmol/mm²]
0.0
0.1
0.1
0.2
0.2
0.3
Outer cerebellum
Inner cerebellum
Hippocampus
Cortex

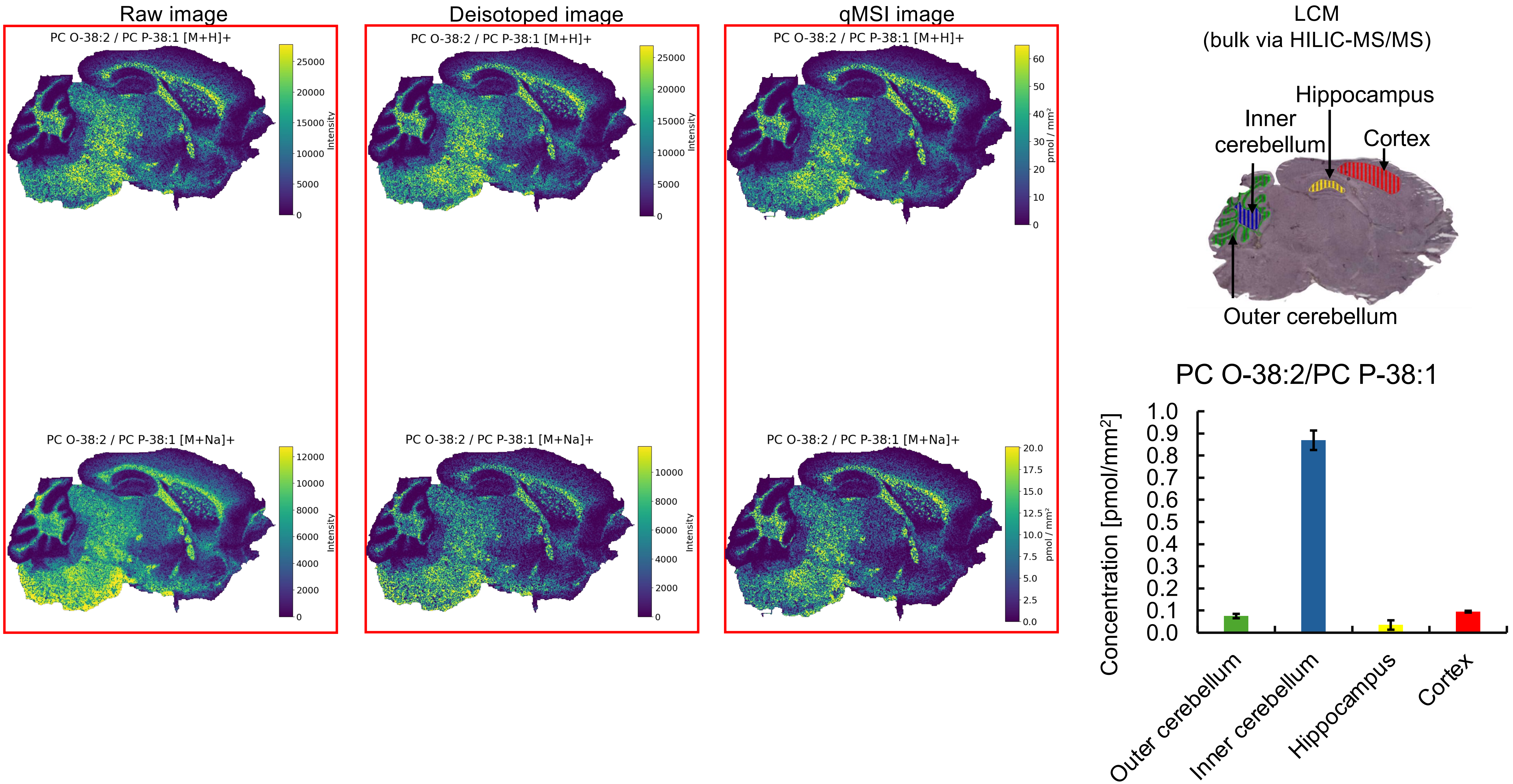

Raw image
Deisotoped image
qMSI image
LCM
(bulk via HILIC-MS/MS)
PC O-38:2 / PC P-38:1 [M+H]+
PC O-38:2 / PC P-38:1 [M+Na]+
Intensity
pmol / mm²
Hippocampus
Inner cerebellum
Cortex
Outer cerebellum
PC O-38:2/PC P-38:1
Concentration [pmol/mm²]
Outer cerebellum
Inner cerebellum
Hippocampus
Cortex

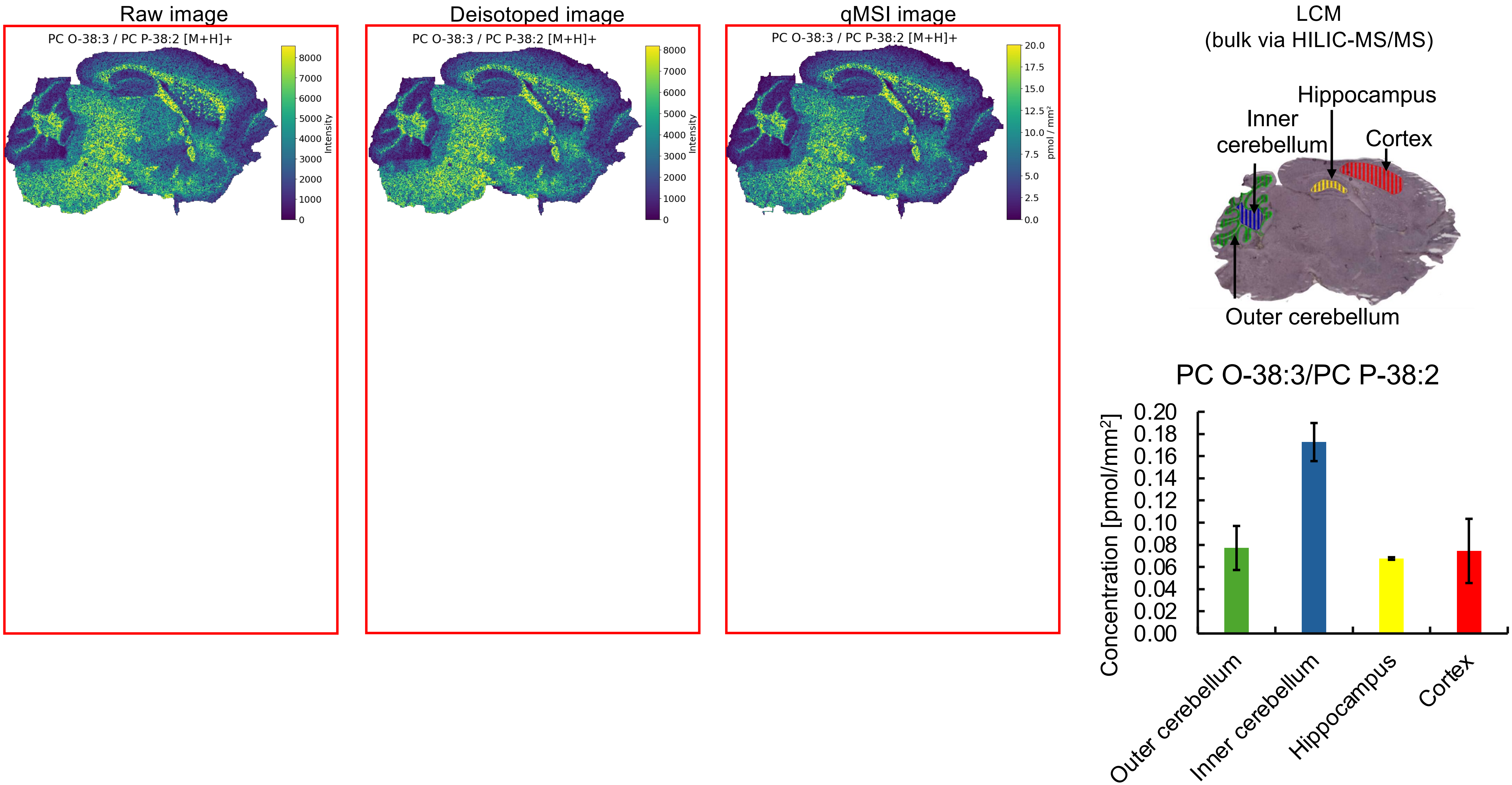

Raw image
PC O-38:3 / PC P-38:2 [M+H]+
Intensity
Deisotoped image
PC O-38:3 / PC P-38:2 [M+H]+
Intensity
qMSI image
PC O-38:3 / PC P-38:2 [M+H]+
pmol / mm²
LCM
(bulk via HILIC-MS/MS)
Hippocampus
Inner cerebellum
Cortex
Outer cerebellum
PC O-38:3/PC P-38:2
Concentration [pmol/mm²]
Outer cerebellum
Inner cerebellum
Hippocampus
Cortex

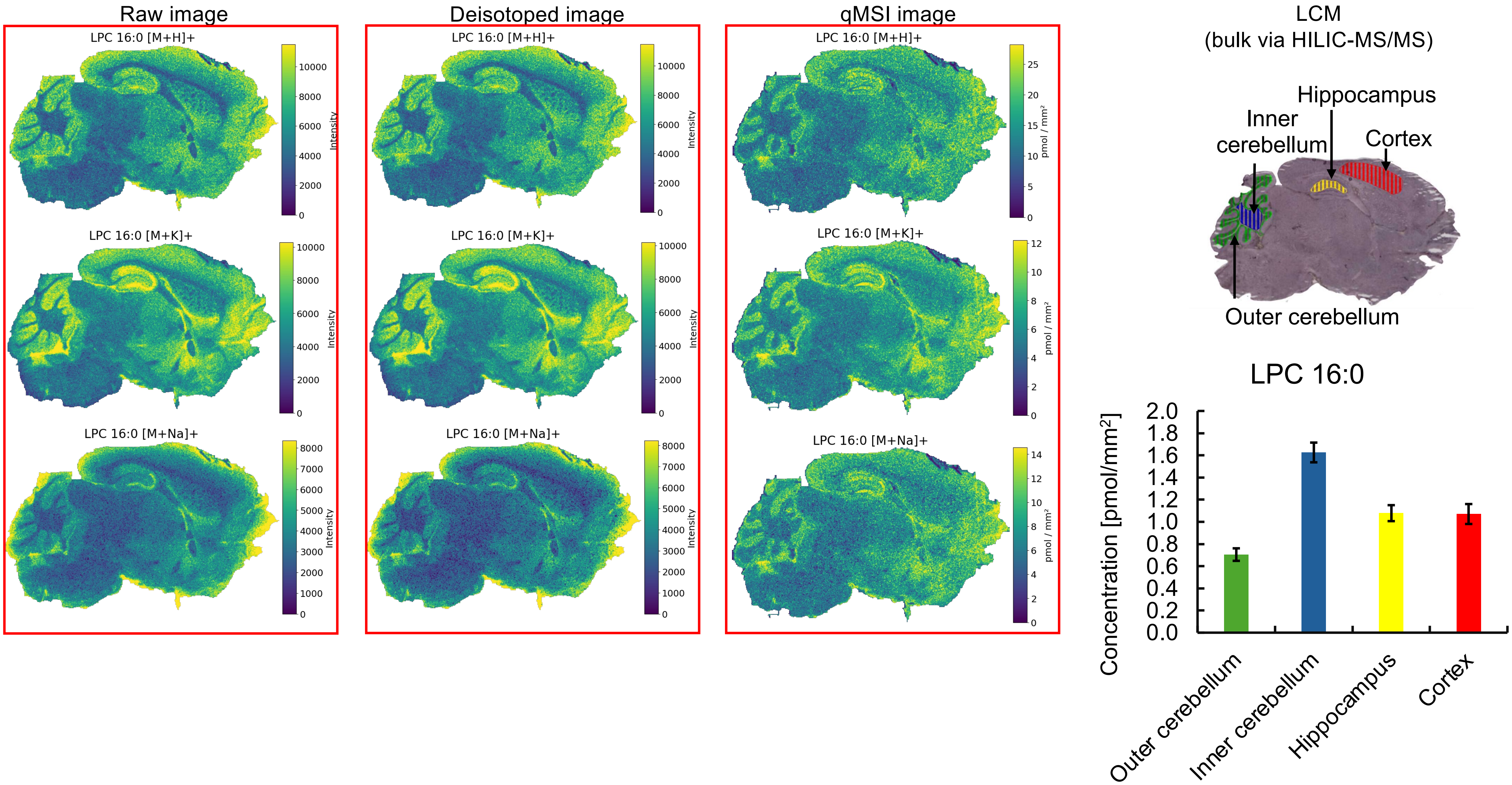
Raw image
Deisotoped image
qMSI image
LPC 16:0 [M+H]+
LPC 16:0 [M+K]+
LPC 16:0 [M+Na]+
Intensity
pmol / mm²
LCM
(bulk via HILIC-MS/MS)
Hippocampus
Inner cerebellum
Cortex
Outer cerebellum
LPC 16:0
Concentration [pmol/mm²]
Outer cerebellum
Inner cerebellum
Hippocampus
Cortex

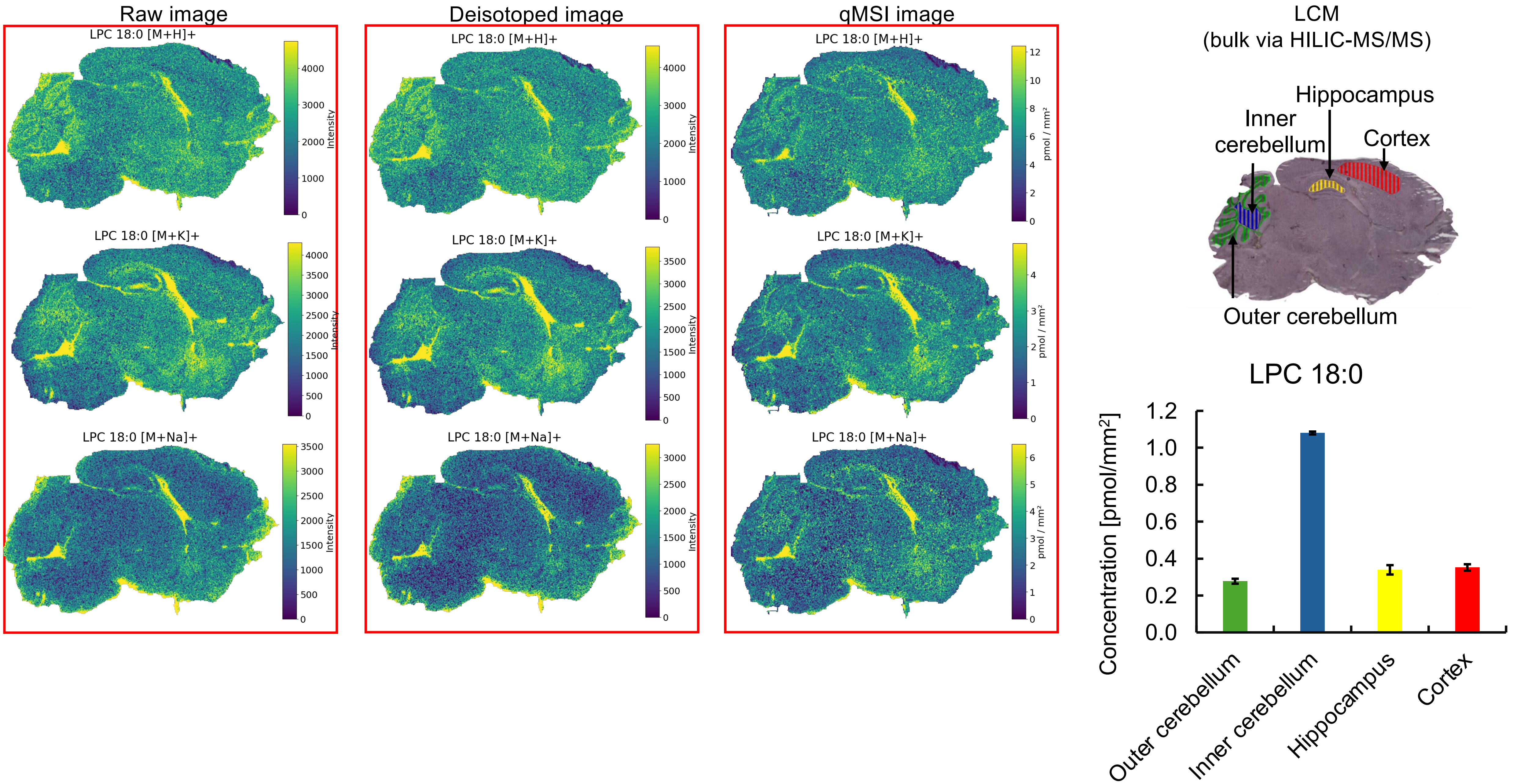

Raw image
Deisotoped image
qMSI image
LCM
(bulk via HILIC-MS/MS)
LPC 18:0 [M+H]+
LPC 18:0 [M+K]+
LPC 18:0 [M+Na]+
Intensity
pmol / mm²
Hippocampus
Inner cerebellum
Cortex
Outer cerebellum
LPC 18:0
Concentration [pmol/mm²]
Outer cerebellum
Inner cerebellum
Hippocampus
Cortex

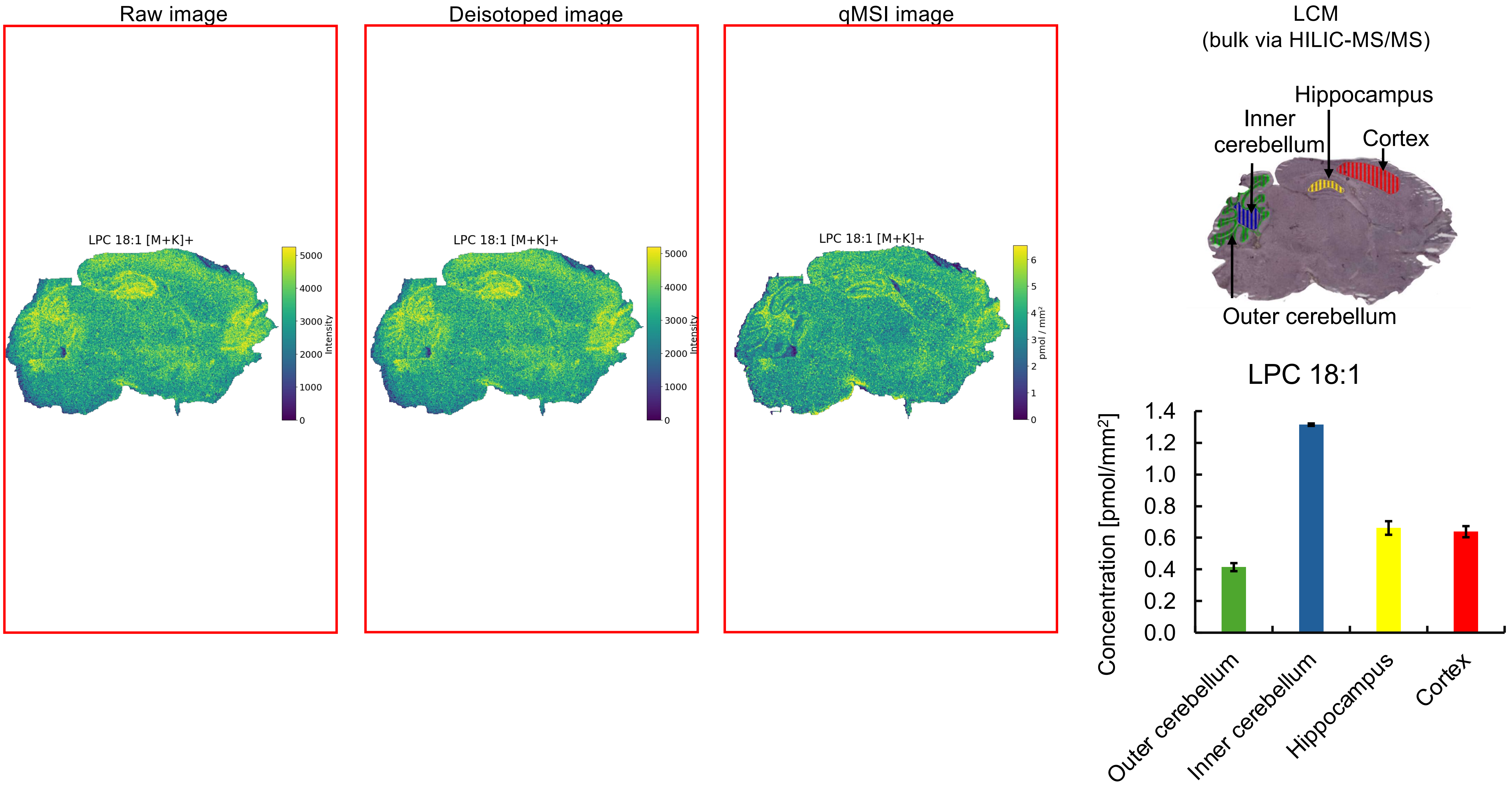

Raw image
LPC 18:1 [M+K]+
Intensity
Deisotoped image
LPC 18:1 [M+K]+
Intensity
qMSI image
LPC 18:1 [M+K]+
pmol / mm²
LCM
(bulk via HILIC-MS/MS)
Hippocampus
Inner cerebellum
Cortex
Outer cerebellum
LPC 18:1
Concentration [pmol/mm²]
Outer cerebellum
Inner cerebellum
Hippocampus
Cortex

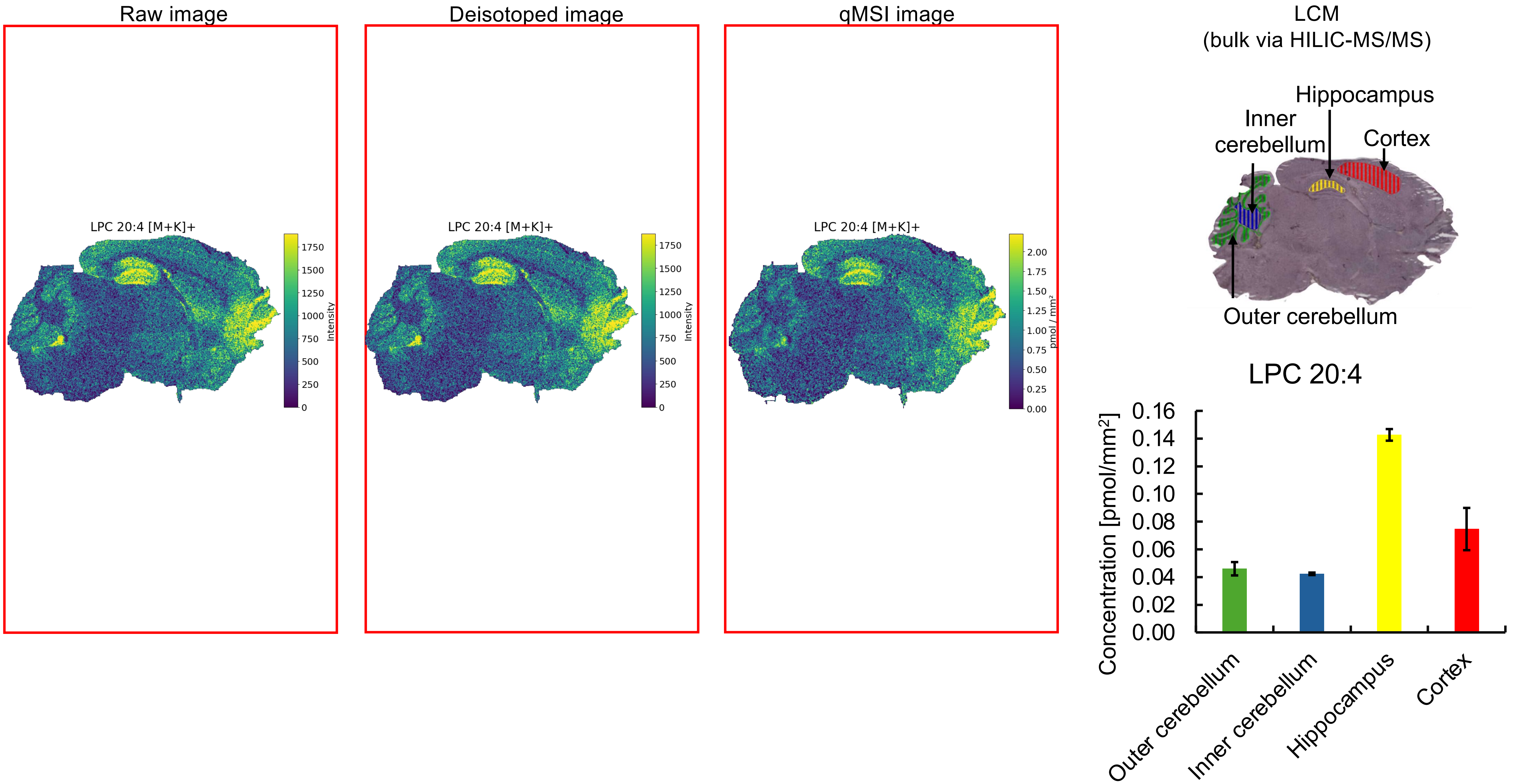
Raw image
LPC 20:4 [M+K]+
Intensity
Deisotoped image
LPC 20:4 [M+K]+
Intensity
qMSI image
LPC 20:4 [M+K]+
pmol / mm²
LCM
(bulk via HILIC-MS/MS)
Hippocampus
Inner cerebellum
Cortex
Outer cerebellum
LPC 20:4
Concentration [pmol/mm²]
Outer cerebellum
Inner cerebellum
Hippocampus
Cortex

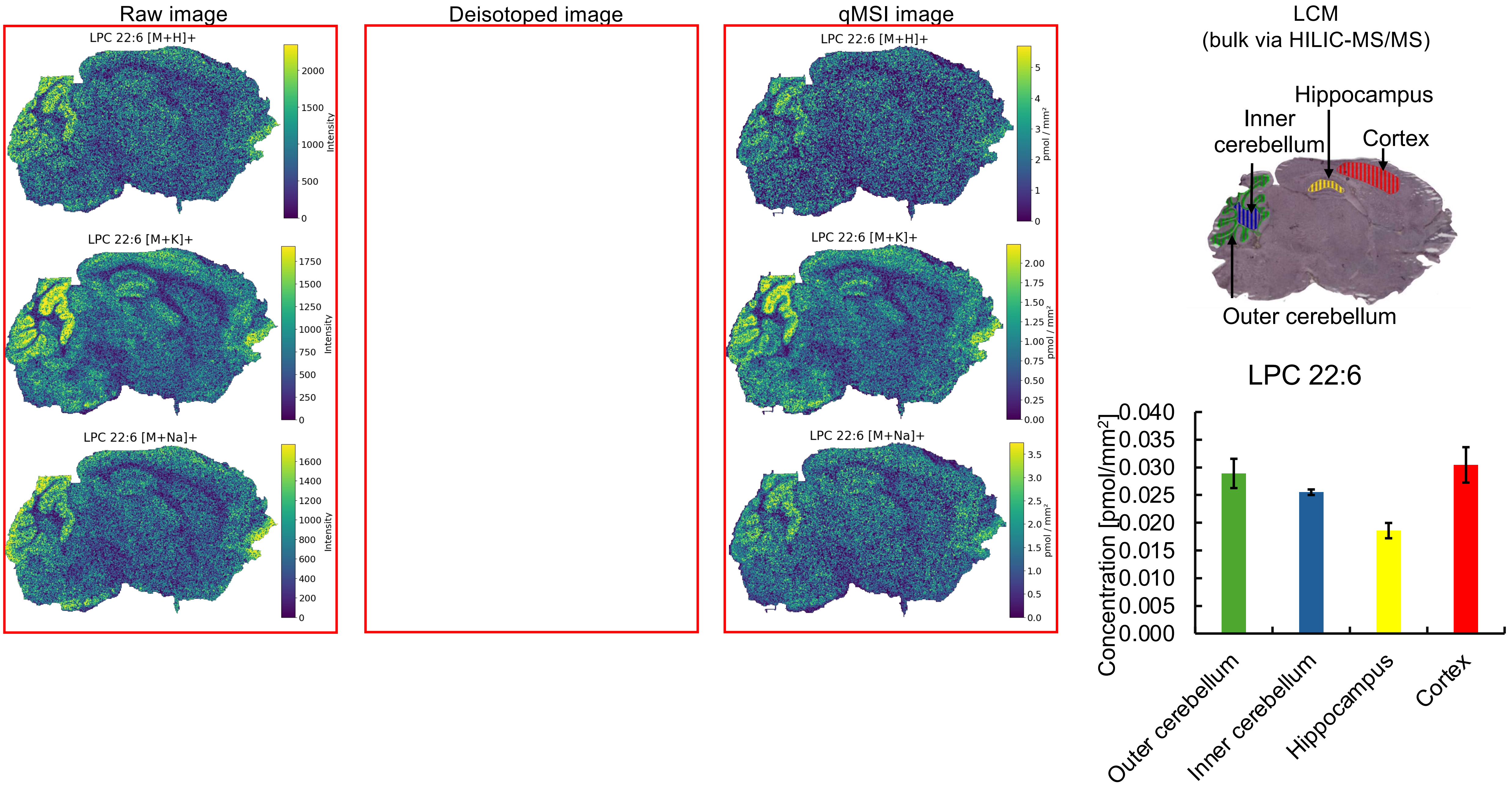

Raw image
Deisotoped image
qMSI image
LCM
(bulk via HILIC-MS/MS)
LPC 22:6 [M+H]+
Intensity
LPC 22:6 [M+K]+
Intensity
LPC 22:6 [M+Na]+
Intensity
LPC 22:6 [M+H]+
pmol / mm²
LPC 22:6 [M+K]+
pmol / mm²
LPC 22:6 [M+Na]+
pmol / mm²
Hippocampus
Inner cerebellum
Cortex
Outer cerebellum
LPC 22:6
Concentration [pmol/mm²]
0.040
0.035
0.030
0.025
0.020
0.015
0.010
0.005
0.000
Outer cerebellum
Inner cerebellum
Hippocampus
Cortex

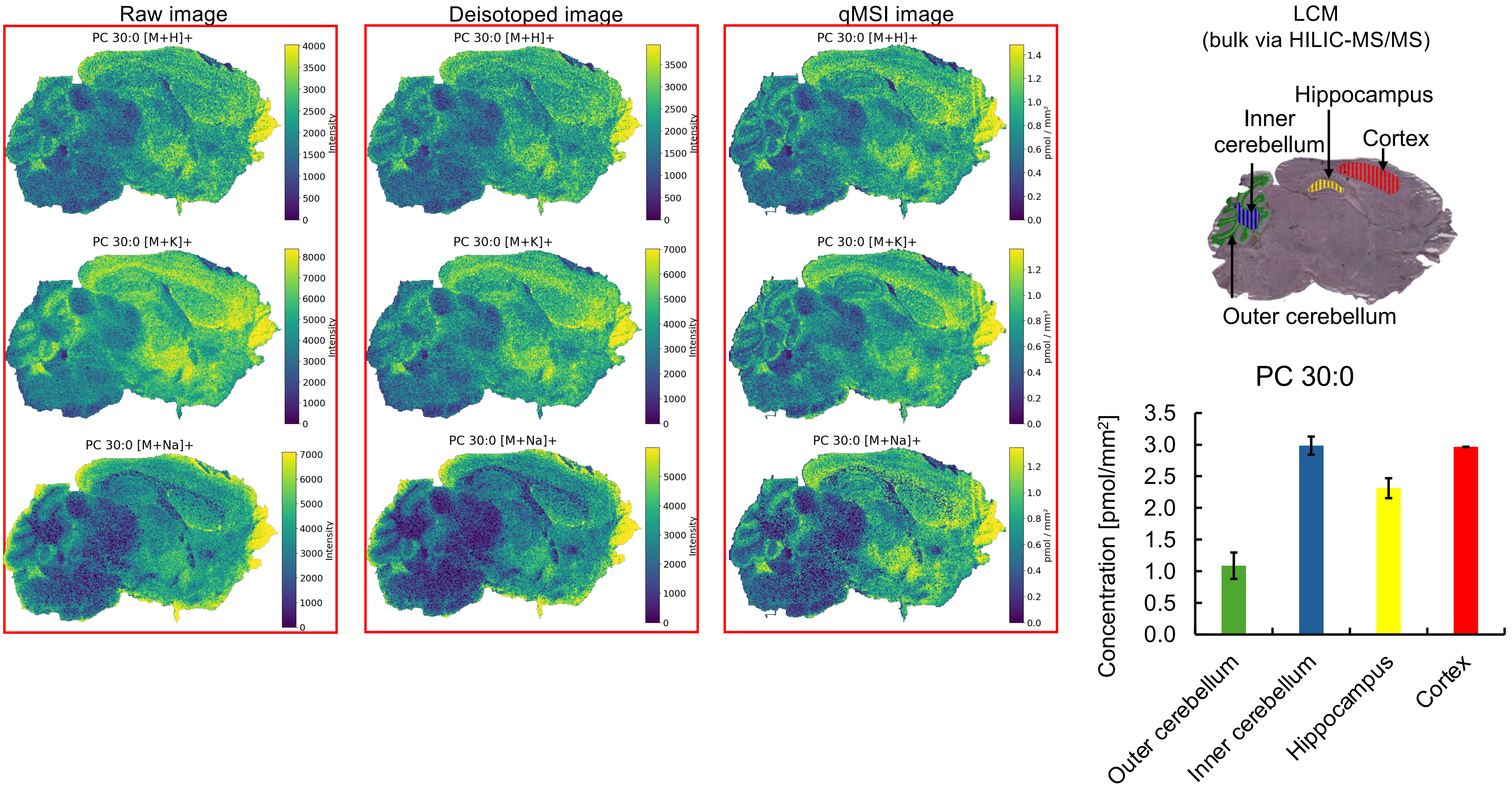


*Missing pixels in **$[M+Na]^+$ adduct** (profile to centroid transformation)

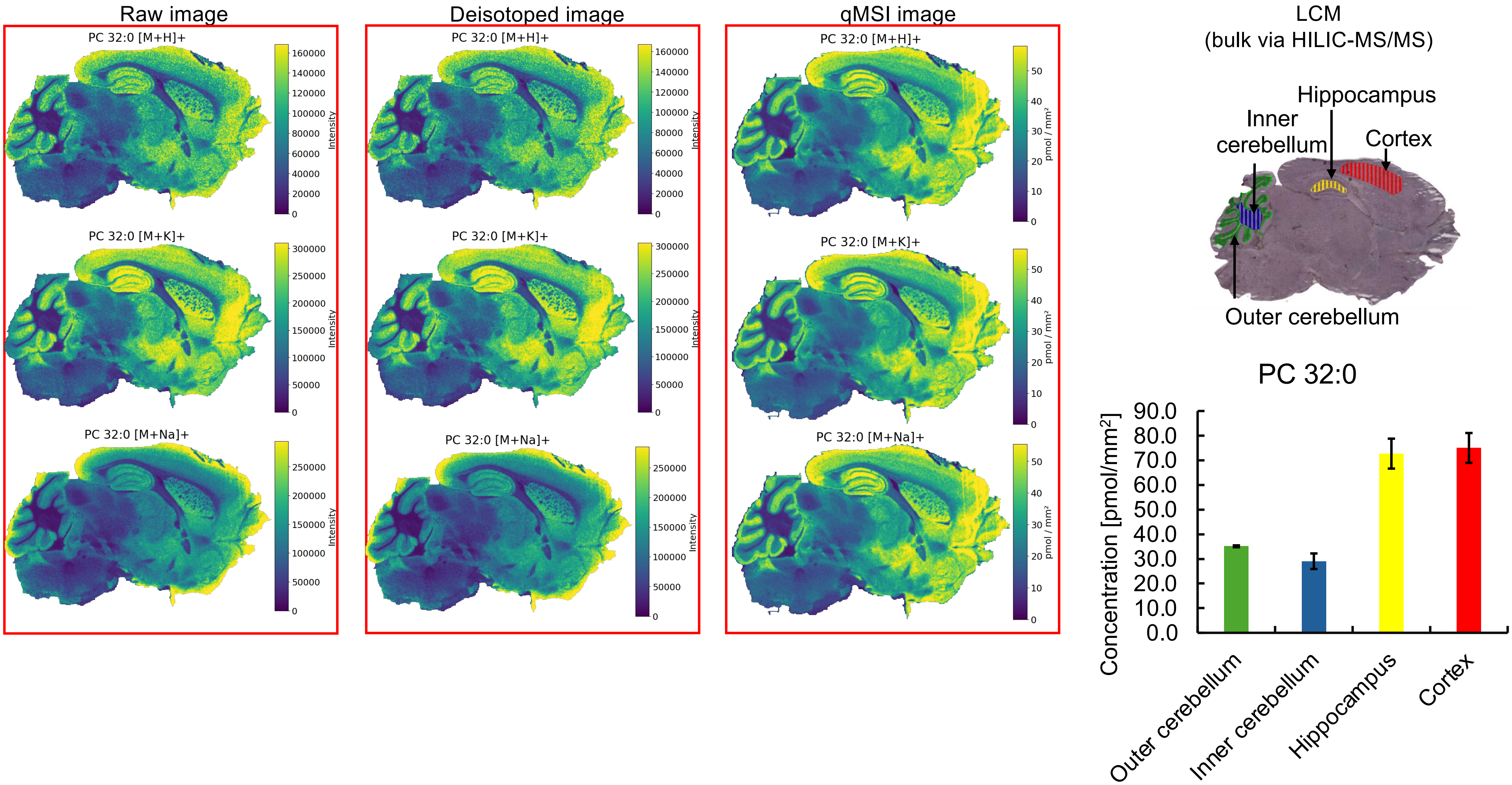

Raw image
Deisotoped image
qMSI image
PC 32:0 [M+H]+
PC 32:0 [M+K]+
PC 32:0 [M+Na]+
Intensity
pmol / mm²
LCM
(bulk via HILIC-MS/MS)
Hippocampus
Inner cerebellum
Cortex
Outer cerebellum
PC 32:0
Concentration [pmol/mm²]
Outer cerebellum
Inner cerebellum
Hippocampus
Cortex

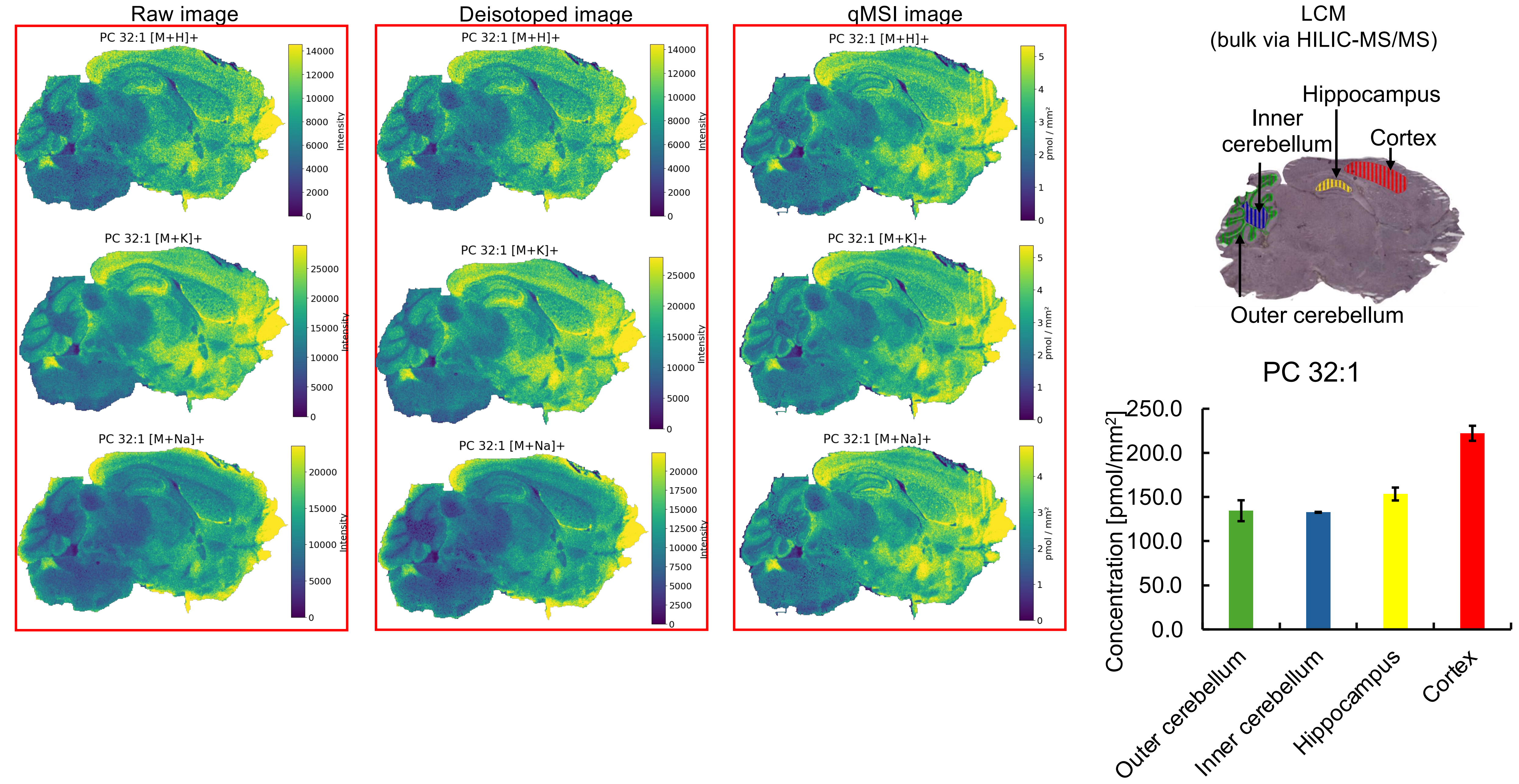


* Missing pixels for **[M+H]⁺** and **[M+Na]⁺** adducts (profile to centroid transformation).

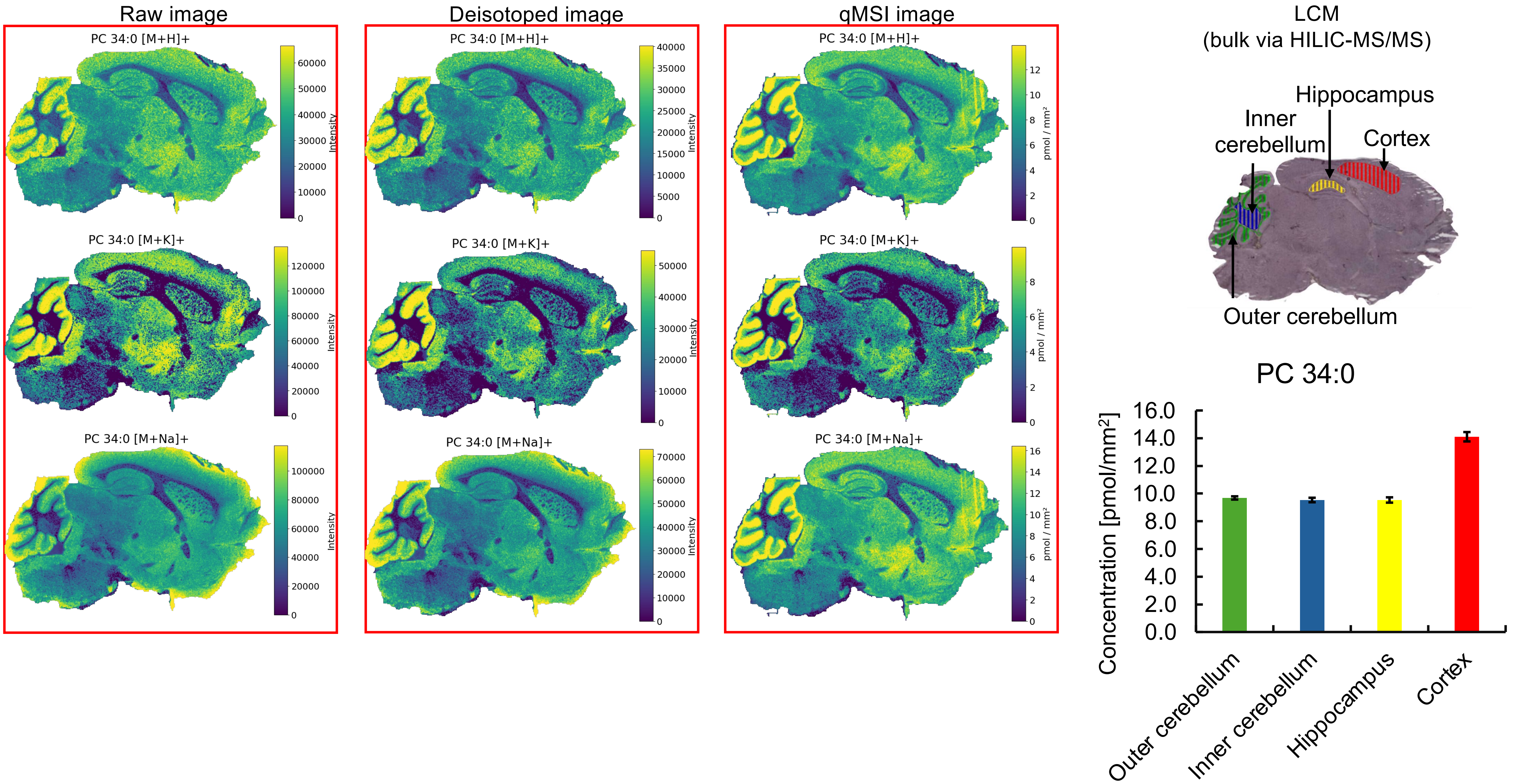


* Missing pixels for **[M+H]⁺, [M+K]⁺, [M+Na]⁺** adducts (profile to centroid transformation).

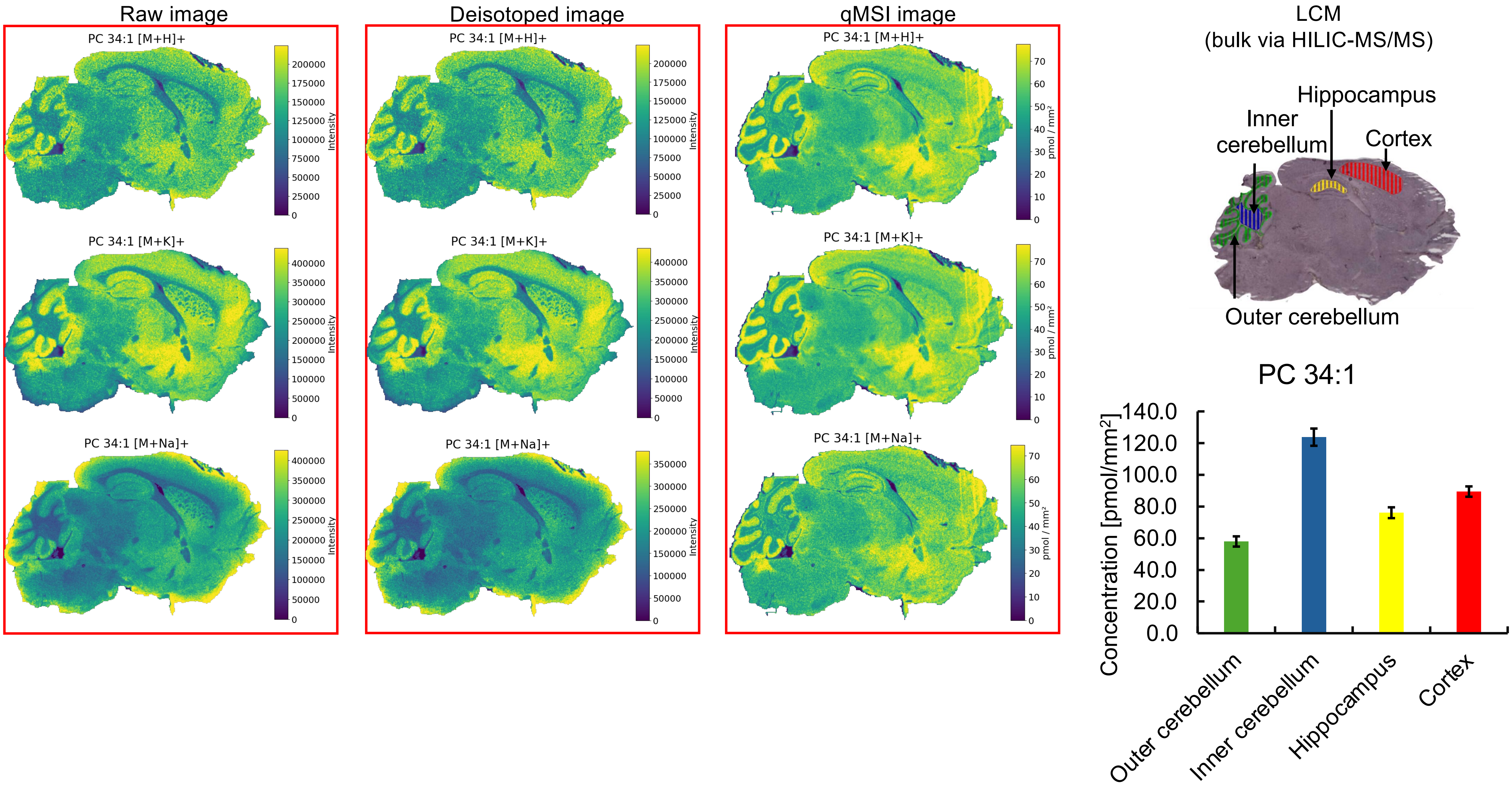

Raw image
PC 34:1 [M+H]+
PC 34:1 [M+K]+
PC 34:1 [M+Na]+
Intensity
Deisotoped image
PC 34:1 [M+H]+
PC 34:1 [M+K]+
PC 34:1 [M+Na]+
Intensity
qMSI image
PC 34:1 [M+H]+
PC 34:1 [M+K]+
PC 34:1 [M+Na]+
pmol / mm²
LCM
(bulk via HILIC-MS/MS)
Hippocampus
Inner cerebellum
Cortex
Outer cerebellum
PC 34:1
Concentration [pmol/mm²]
Outer cerebellum
Inner cerebellum
Hippocampus
Cortex

Raw image

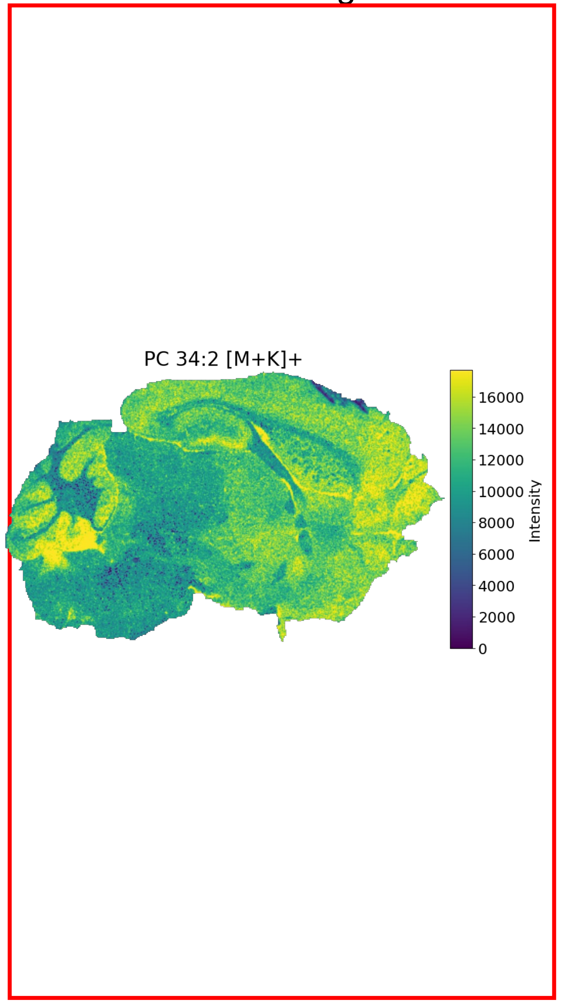


Deisotoped image

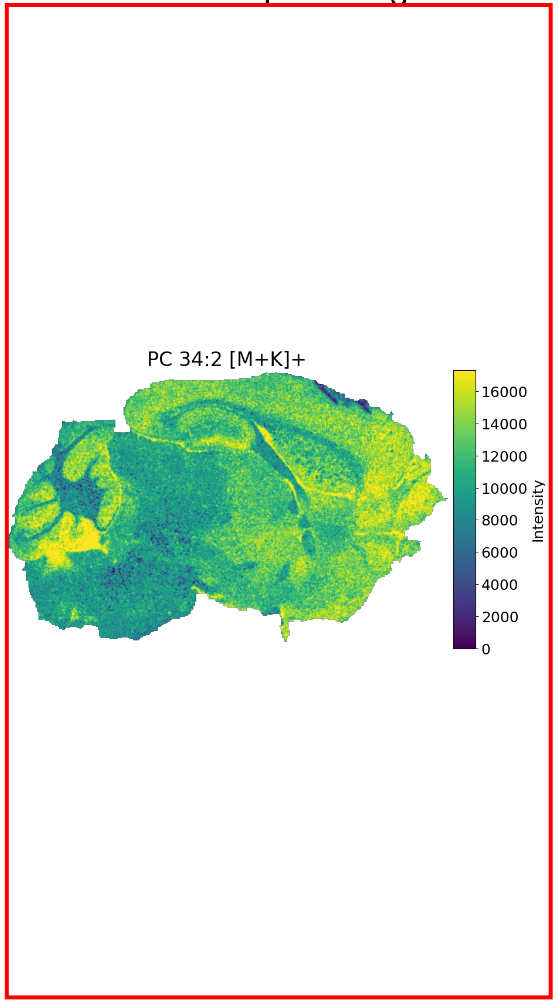


qMSI image

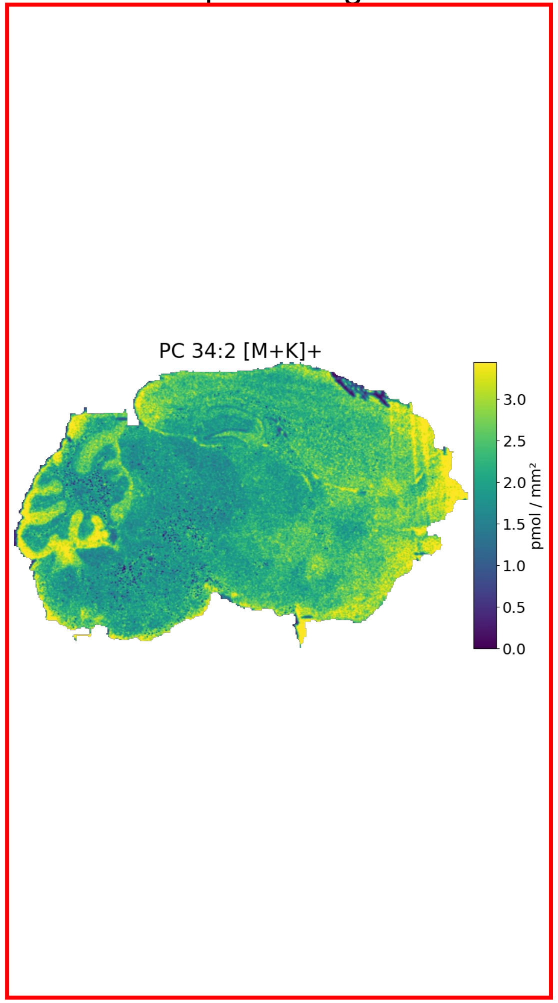


LCM
(bulk via HILIC-MS/MS)

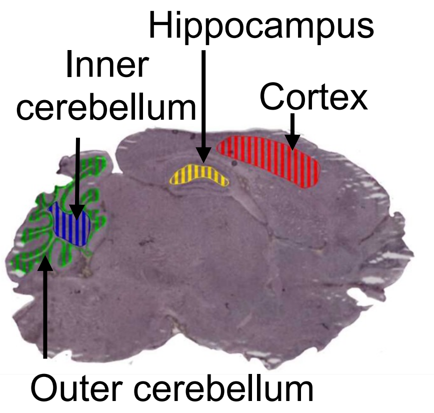


PC 34:2

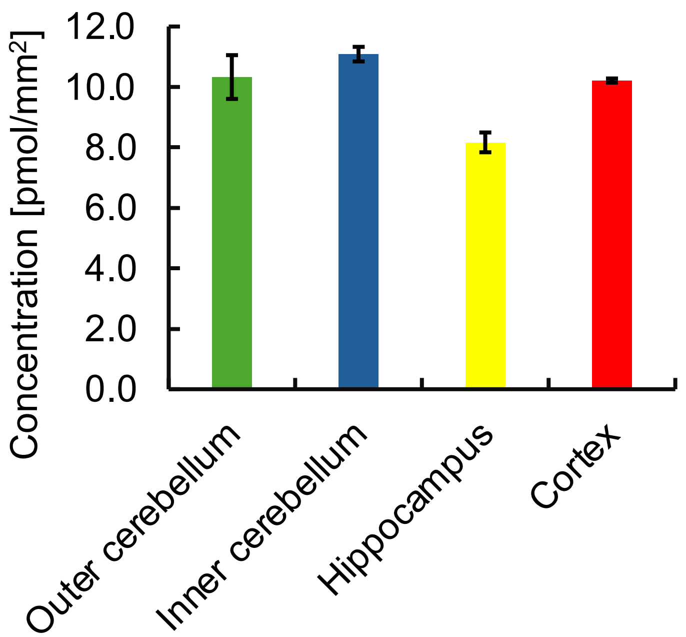


* Other adducts = too low abundant. Missing pixels for **[M+K]⁺** adduct (profile to centroid transformation).

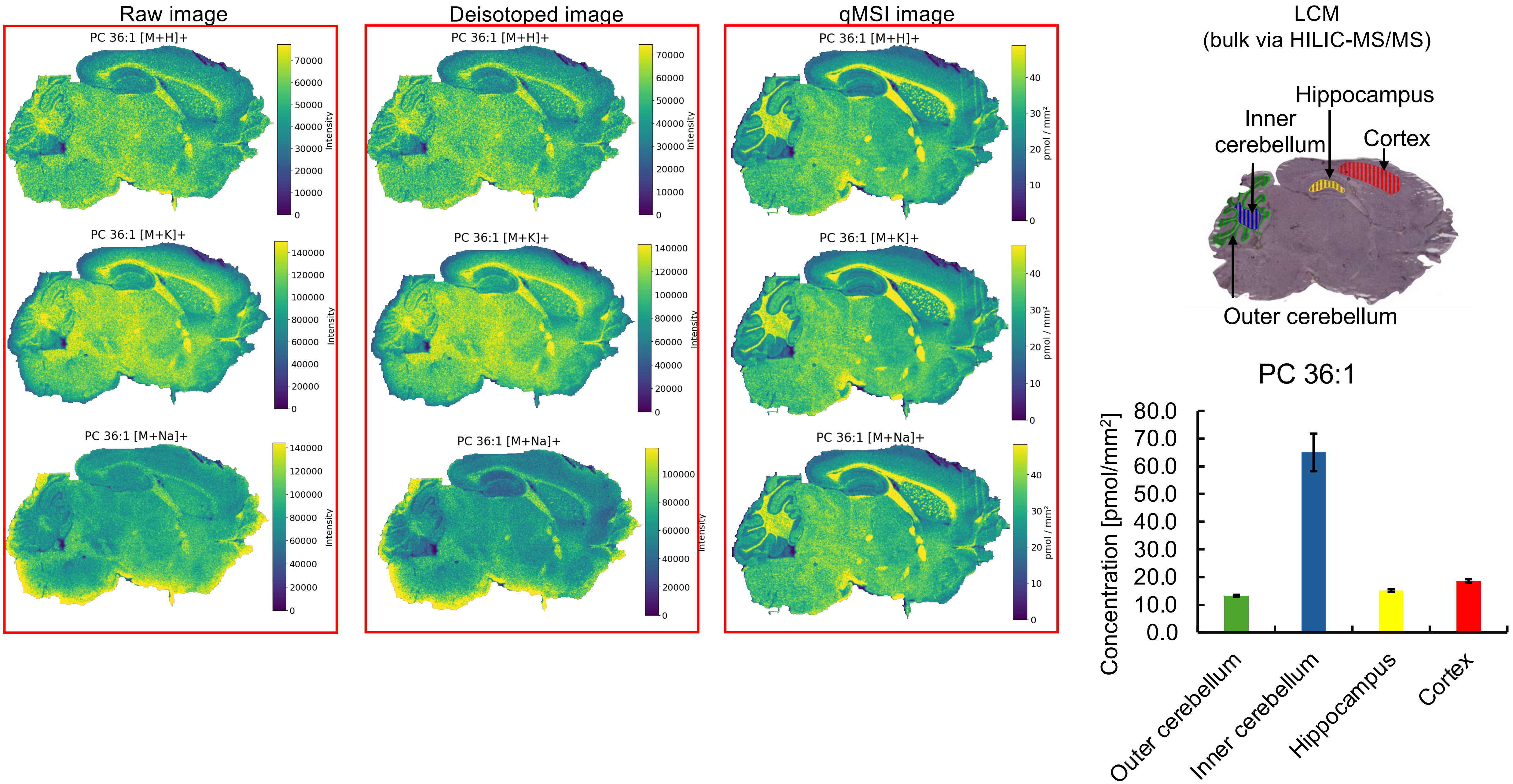

Raw image
Deisotoped image
qMSI image
PC 36:1 [M+H]+
PC 36:1 [M+K]+
PC 36:1 [M+Na]+
Intensity
pmol / mm²
LCM
(bulk via HILIC-MS/MS)
Hippocampus
Inner cerebellum
Cortex
Outer cerebellum
PC 36:1
Concentration [pmol/mm²]
80.0
70.0
60.0
50.0
40.0
30.0
20.0
10.0
0.0
Outer cerebellum
Inner cerebellum
Hippocampus
Cortex

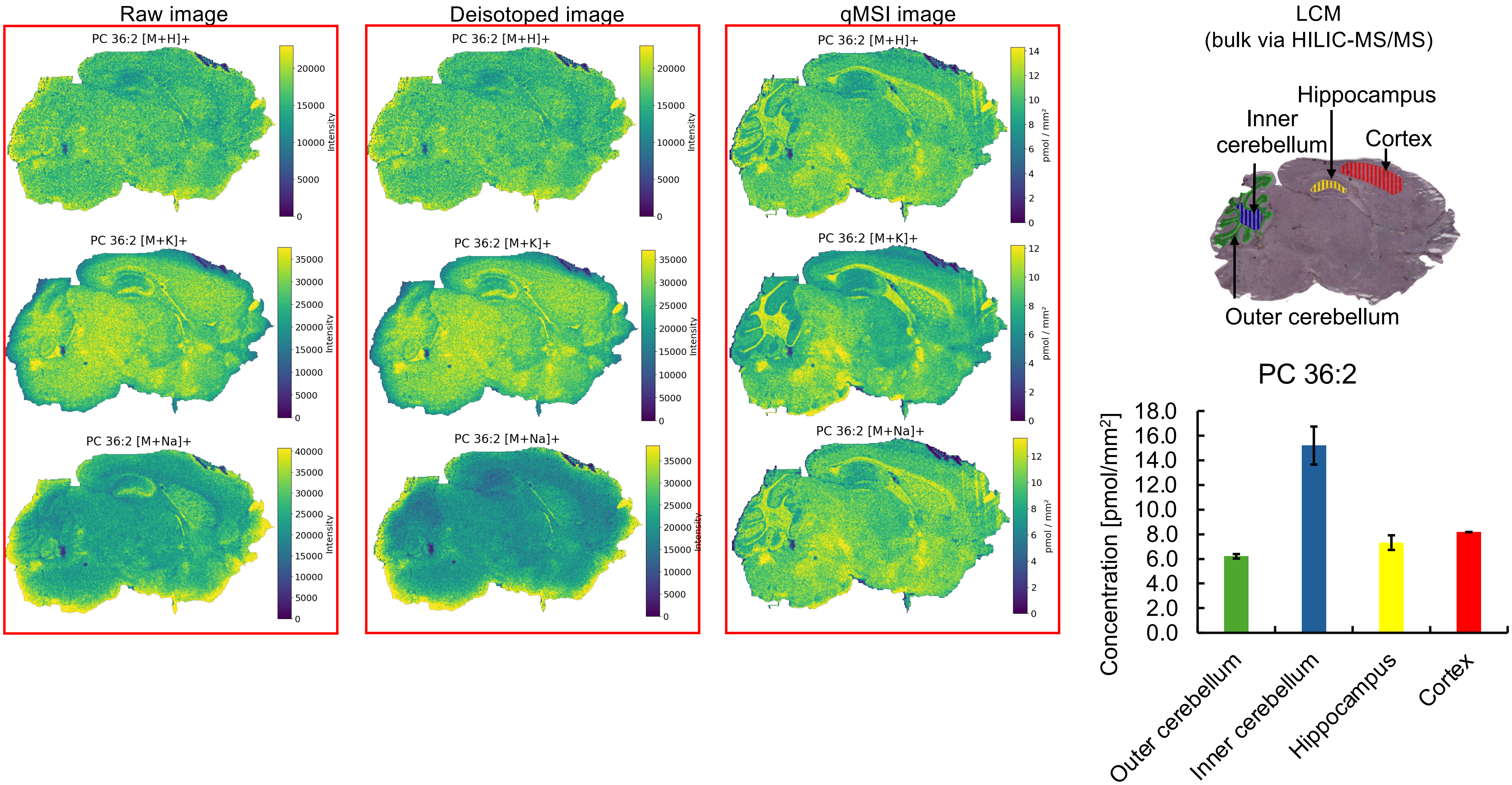
Raw image
Deisotoped image
qMSI image
LCM
(bulk via HILIC-MS/MS)
PC 36:2 [M+H]+
PC 36:2 [M+K]+
PC 36:2 [M+Na]+
Intensity
pmol / mm²
Hippocampus
Inner cerebellum
Cortex
Outer cerebellum
PC 36:2
Concentration [pmol/mm²]
Outer cerebellum
Inner cerebellum
Hippocampus
Cortex

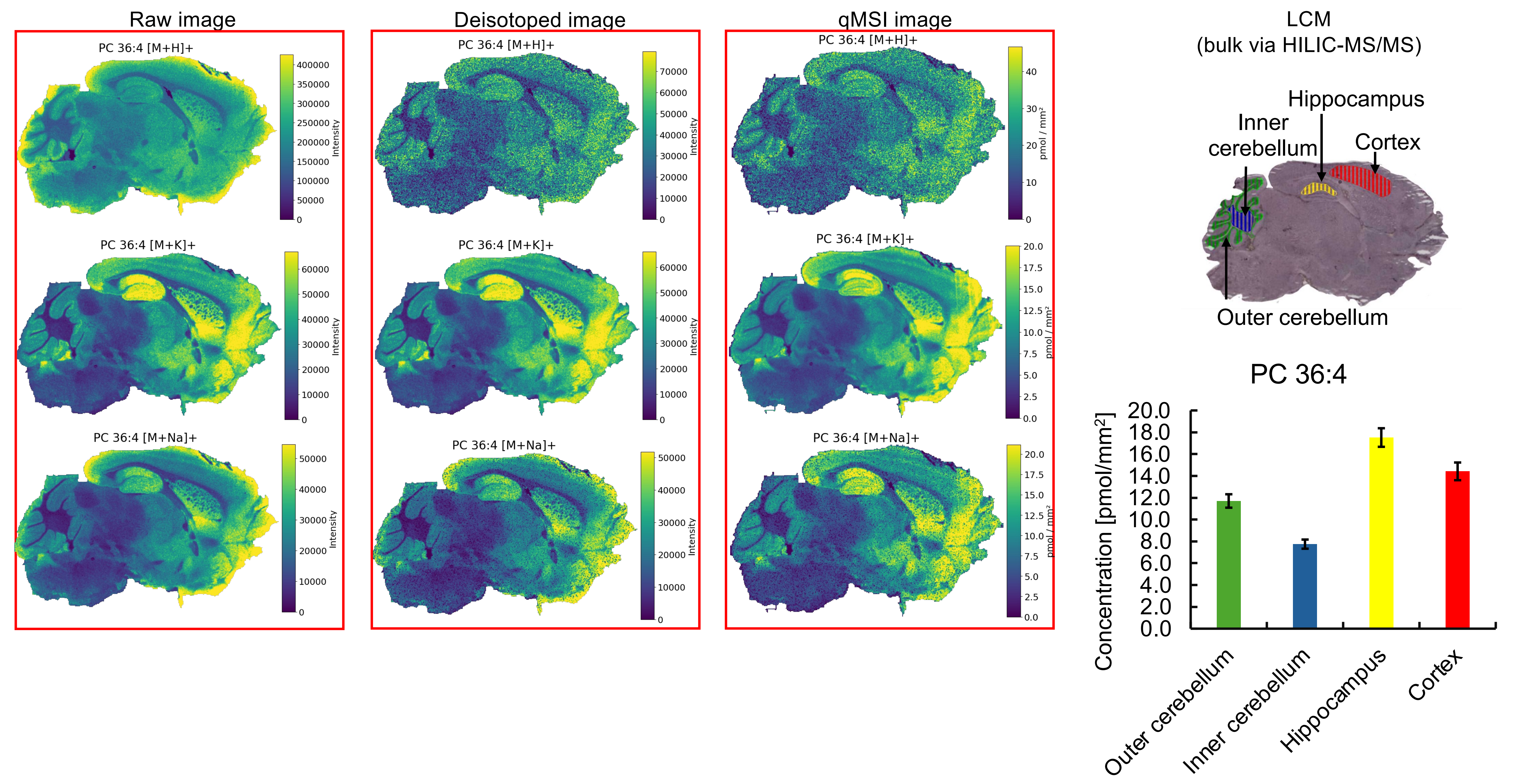

Raw image
Deisotoped image
qMSI image
PC 36:4 [M+H]+
PC 36:4 [M+K]+
PC 36:4 [M+Na]+
Intensity
pmol / mm²
LCM
(bulk via HILIC-MS/MS)
Hippocampus
Inner cerebellum
Cortex
Outer cerebellum
PC 36:4
Concentration [pmol/mm²]
Outer cerebellum
Inner cerebellum
Hippocampus
Cortex

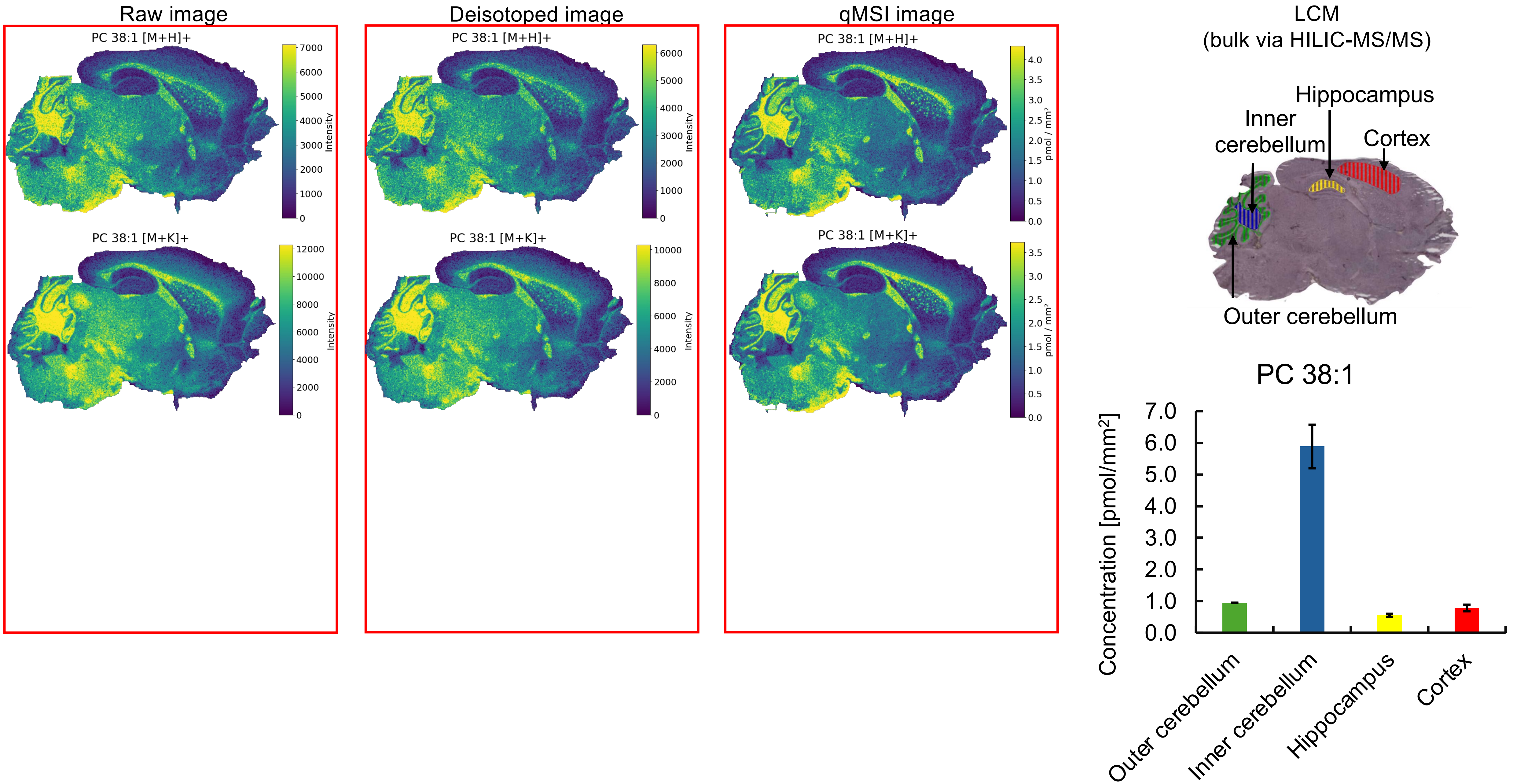

Raw image
PC 38:1 [M+H]+
Intensity
PC 38:1 [M+K]+
Intensity
Deisotoped image
PC 38:1 [M+H]+
Intensity
PC 38:1 [M+K]+
Intensity
qMSI image
PC 38:1 [M+H]+
pmol / mm²
PC 38:1 [M+K]+
pmol / mm²
LCM
(bulk via HILIC-MS/MS)
Hippocampus
Inner cerebellum
Cortex
Outer cerebellum
PC 38:1
Concentration [pmol/mm²]
Outer cerebellum
Inner cerebellum
Hippocampus
Cortex

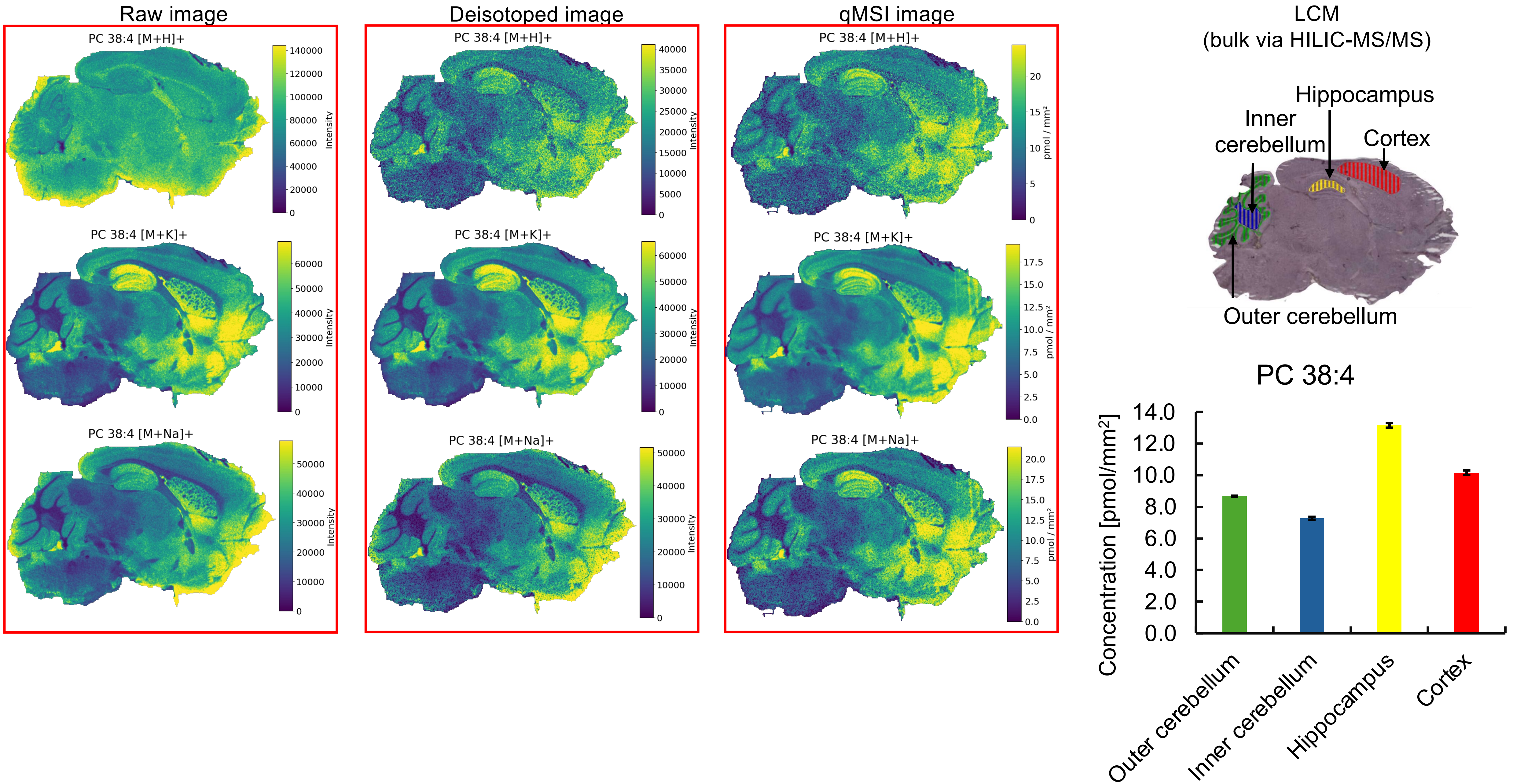

Raw image
Deisotoped image
qMSI image
LCM
(bulk via HILIC-MS/MS)
PC 38:4 [M+H]+
PC 38:4 [M+K]+
PC 38:4 [M+Na]+
Intensity
pmol / mm²
Hippocampus
Inner cerebellum
Cortex
Outer cerebellum
PC 38:4
Concentration [pmol/mm²]
14.0
12.0
10.0
8.0
6.0
4.0
2.0
0.0
Outer cerebellum
Inner cerebellum
Hippocampus
Cortex

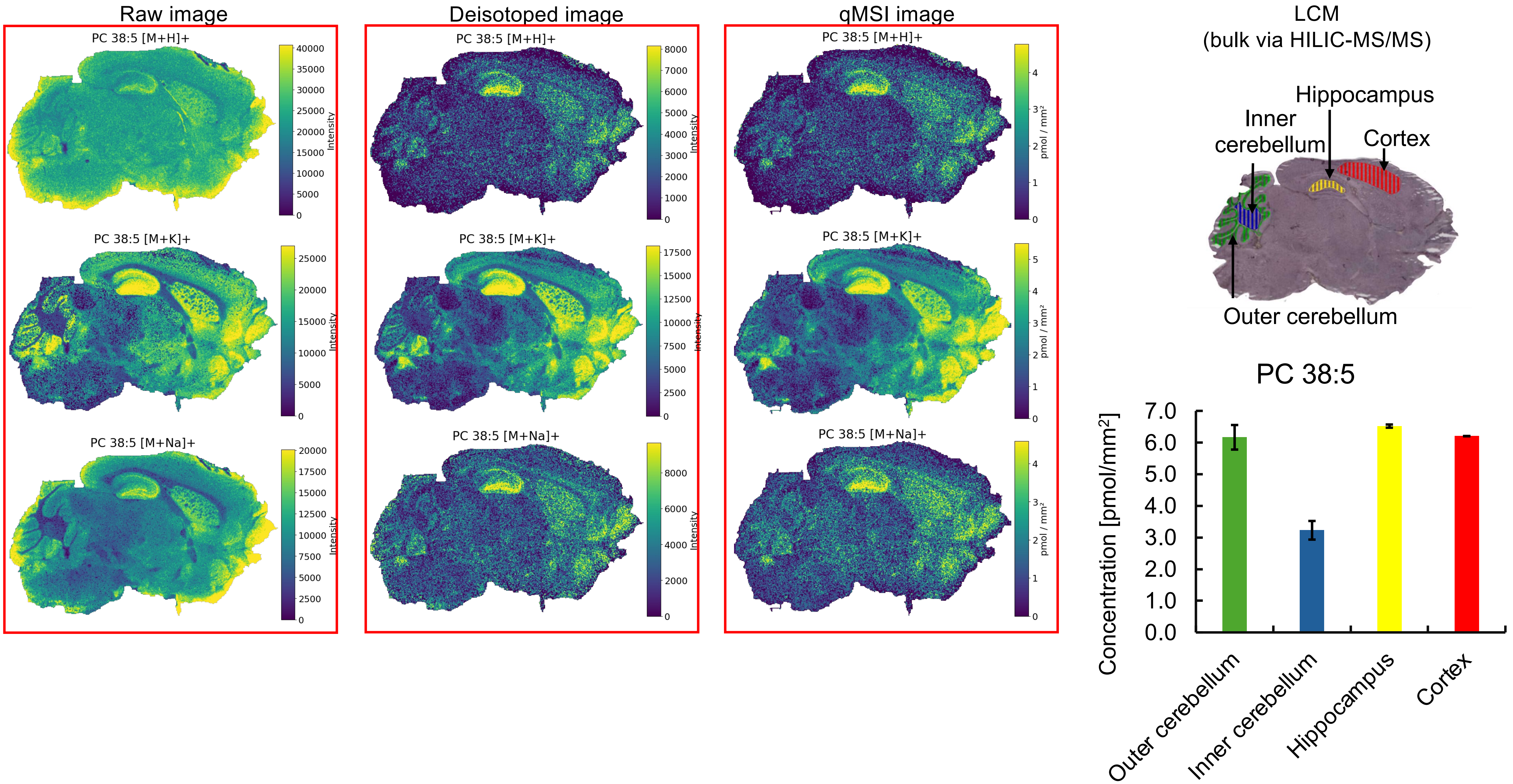

Raw image
Deisotoped image
qMSI image
LCM
(bulk via HILIC-MS/MS)
PC 38:5 [M+H]+
PC 38:5 [M+K]+
PC 38:5 [M+Na]+
Intensity
pmol / mm²
Hippocampus
Inner cerebellum
Cortex
Outer cerebellum
PC 38:5
Concentration [pmol/mm²]
Outer cerebellum
Inner cerebellum
Hippocampus
Cortex

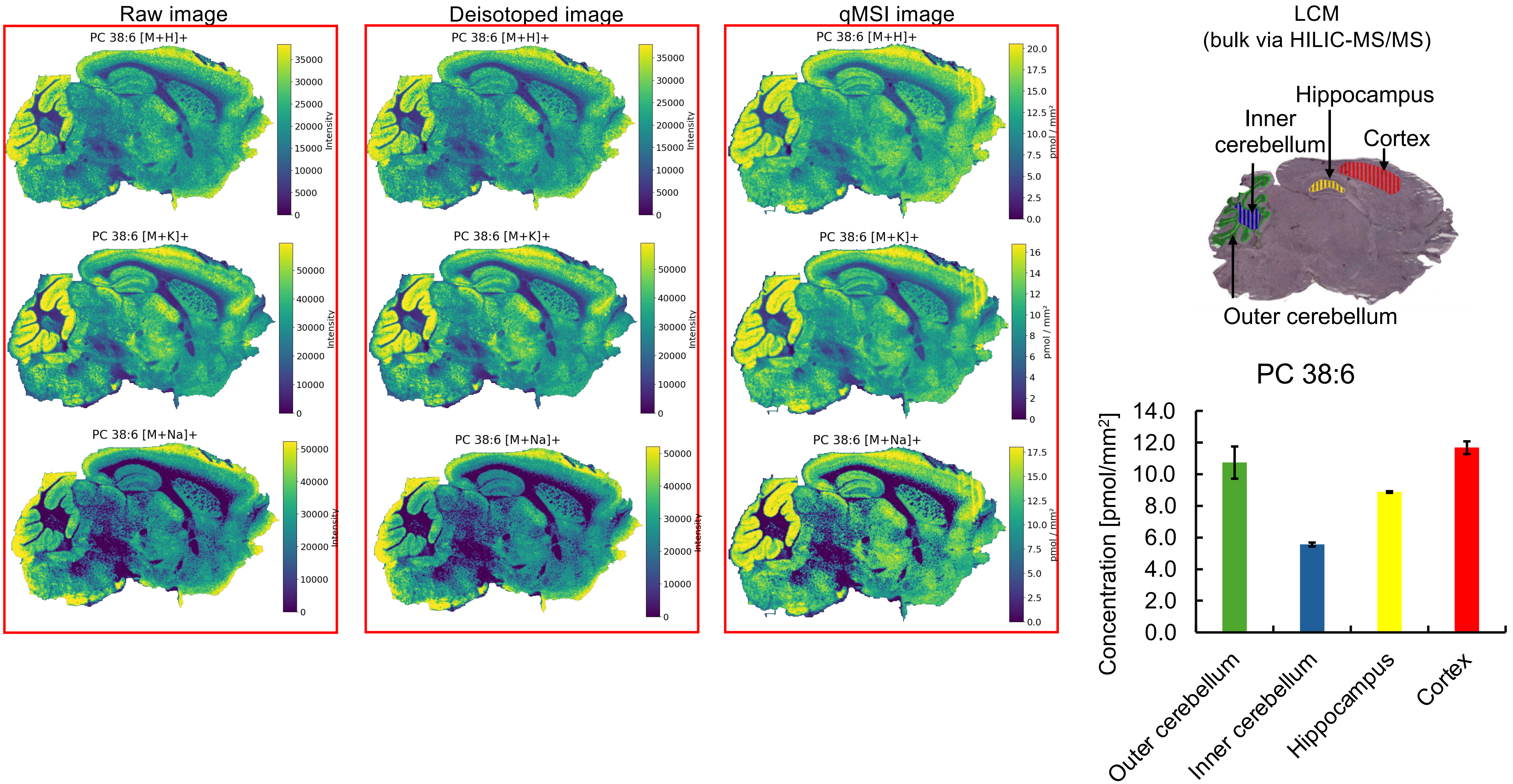

Raw image
Deisotoped image
qMSI image
PC 38:6 [M+H]+
PC 38:6 [M+K]+
PC 38:6 [M+Na]+
Intensity
pmol / mm²
LCM
(bulk via HILIC-MS/MS)
Hippocampus
Inner cerebellum
Cortex
Outer cerebellum
PC 38:6
Concentration [pmol/mm²]
14.0
12.0
10.0
8.0
6.0
4.0
2.0
0.0
Outer cerebellum
Inner cerebellum
Hippocampus
Cortex

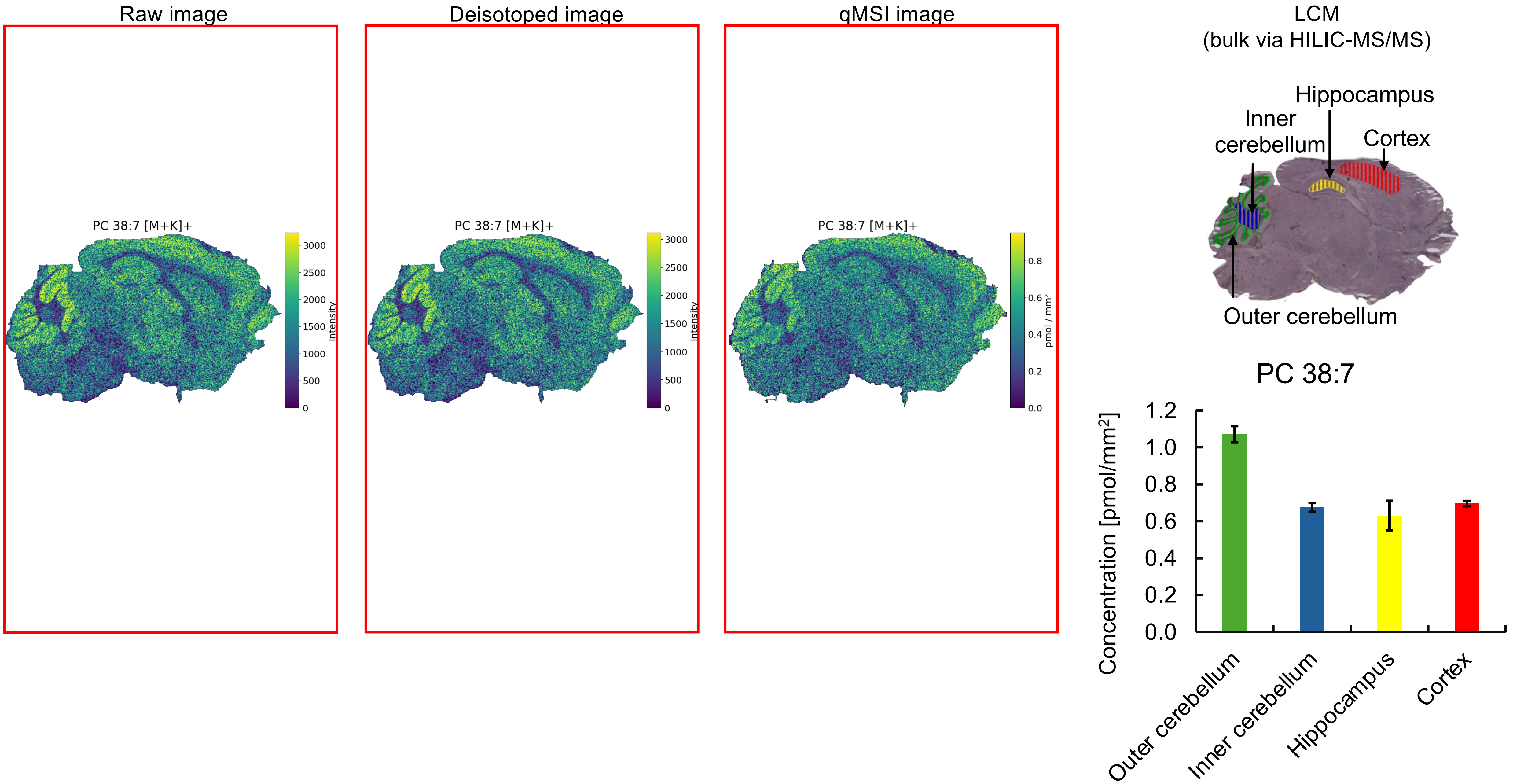

Raw image
Deisotoped image
qMSI image
PC 38:7 [M+K]+
Intensity
pmol / mm²
LCM
(bulk via HILIC-MS/MS)
Hippocampus
Inner cerebellum
Cortex
Outer cerebellum
PC 38:7
Concentration [pmol/mm²]
Outer cerebellum
Inner cerebellum
Hippocampus
Cortex

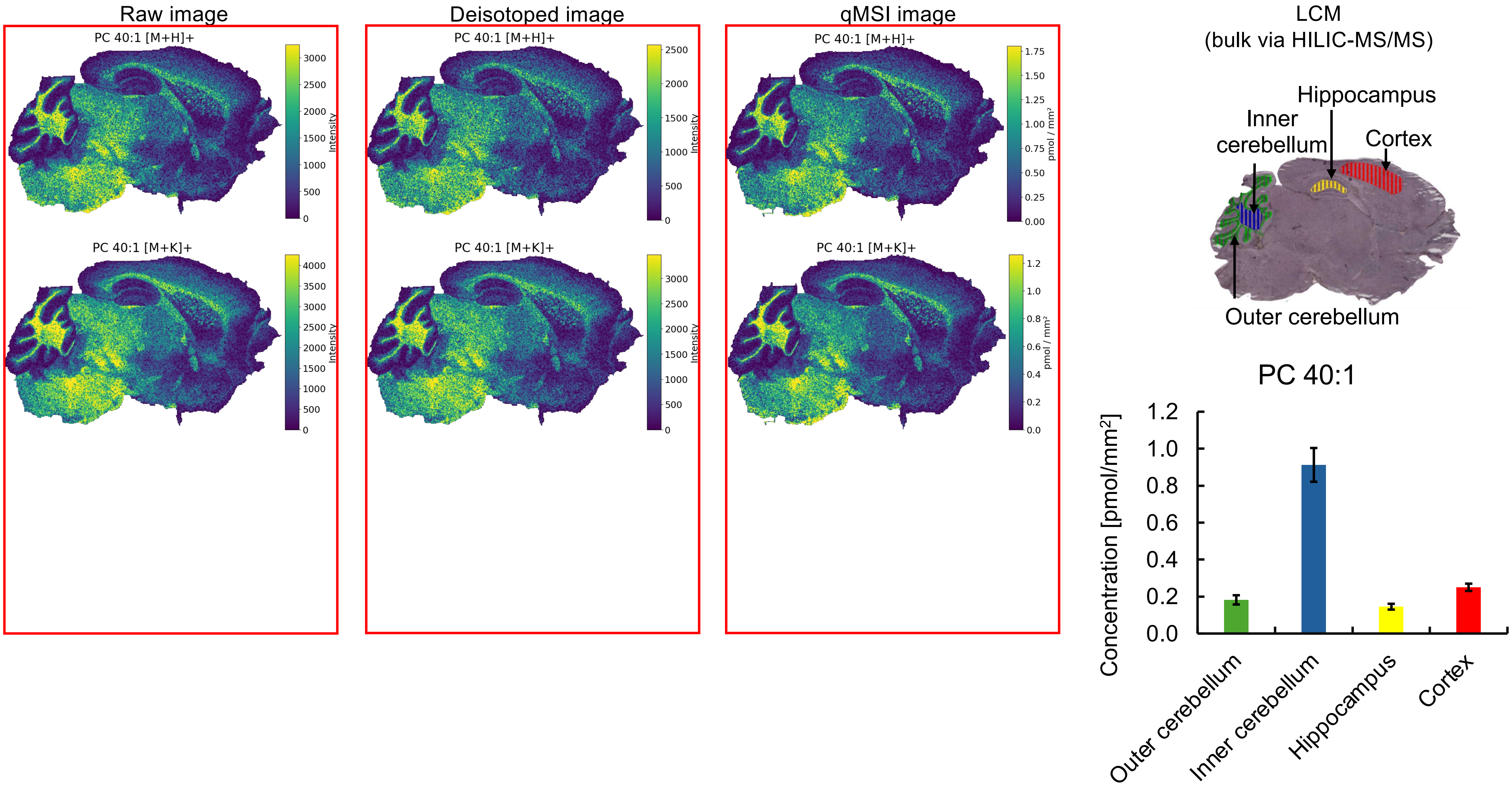

Raw image
PC 40:1 [M+H]+
3000
2500
2000
1500
1000
500
0
Intensity
PC 40:1 [M+K]+
4000
3500
3000
2500
2000
1500
1000
500
0
Intensity
Deisotoped image
PC 40:1 [M+H]+
2500
2000
1500
1000
500
0
Intensity
PC 40:1 [M+K]+
3000
2500
2000
1500
1000
500
0
Intensity
qMSI image
PC 40:1 [M+H]+
1.75
1.50
1.25
1.00
0.75
0.50
0.25
0.00
pmol / mm²
PC 40:1 [M+K]+
1.2
1.0
0.8
0.6
0.4
0.2
0.0
pmol / mm²
LCM
(bulk via HILIC-MS/MS)
Hippocampus
Inner cerebellum
Cortex
Outer cerebellum
PC 40:1
Concentration [pmol/mm²]
1.2
1.0
0.8
0.6
0.4
0.2
0.0
Outer cerebellum
Inner cerebellum
Hippocampus
Cortex

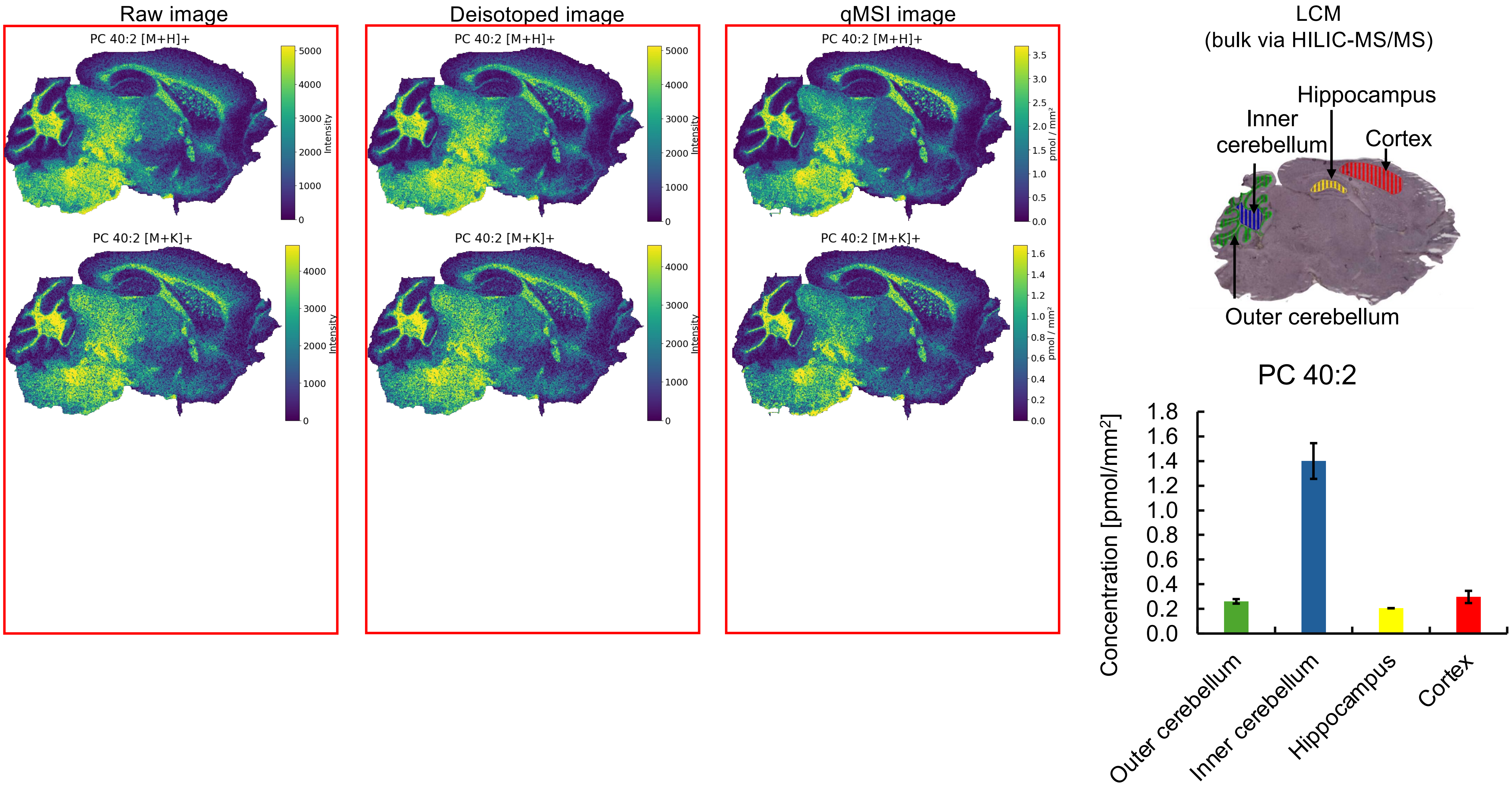
Raw image
Deisotoped image
qMSI image
PC 40:2 [M+H]+
PC 40:2 [M+K]+
Intensity
pmol / mm²
LCM
(bulk via HILIC-MS/MS)
Hippocampus
Inner cerebellum
Cortex
Outer cerebellum
PC 40:2
Concentration [pmol/mm²]
Outer cerebellum
Inner cerebellum
Hippocampus
Cortex

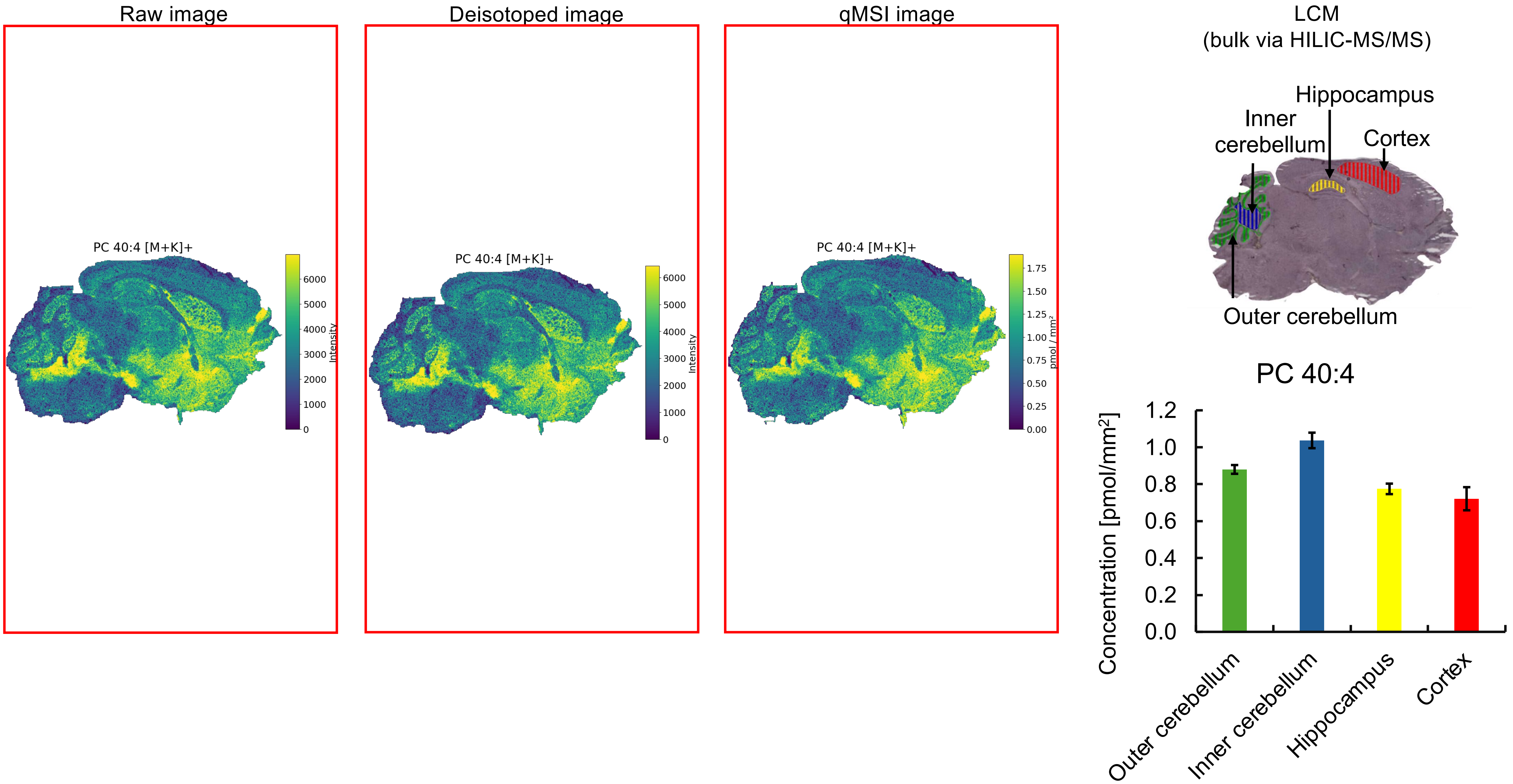
Raw image
Deisotoped image
qMSI image
PC 40:4 [M+K]+
Intensity
PC 40:4 [M+K]+
Intensity
PC 40:4 [M+K]+
pmol / mm²
LCM
(bulk via HILIC-MS/MS)
Hippocampus
Inner cerebellum
Cortex
Outer cerebellum
PC 40:4
Concentration [pmol/mm²]
Outer cerebellum
Inner cerebellum
Hippocampus
Cortex

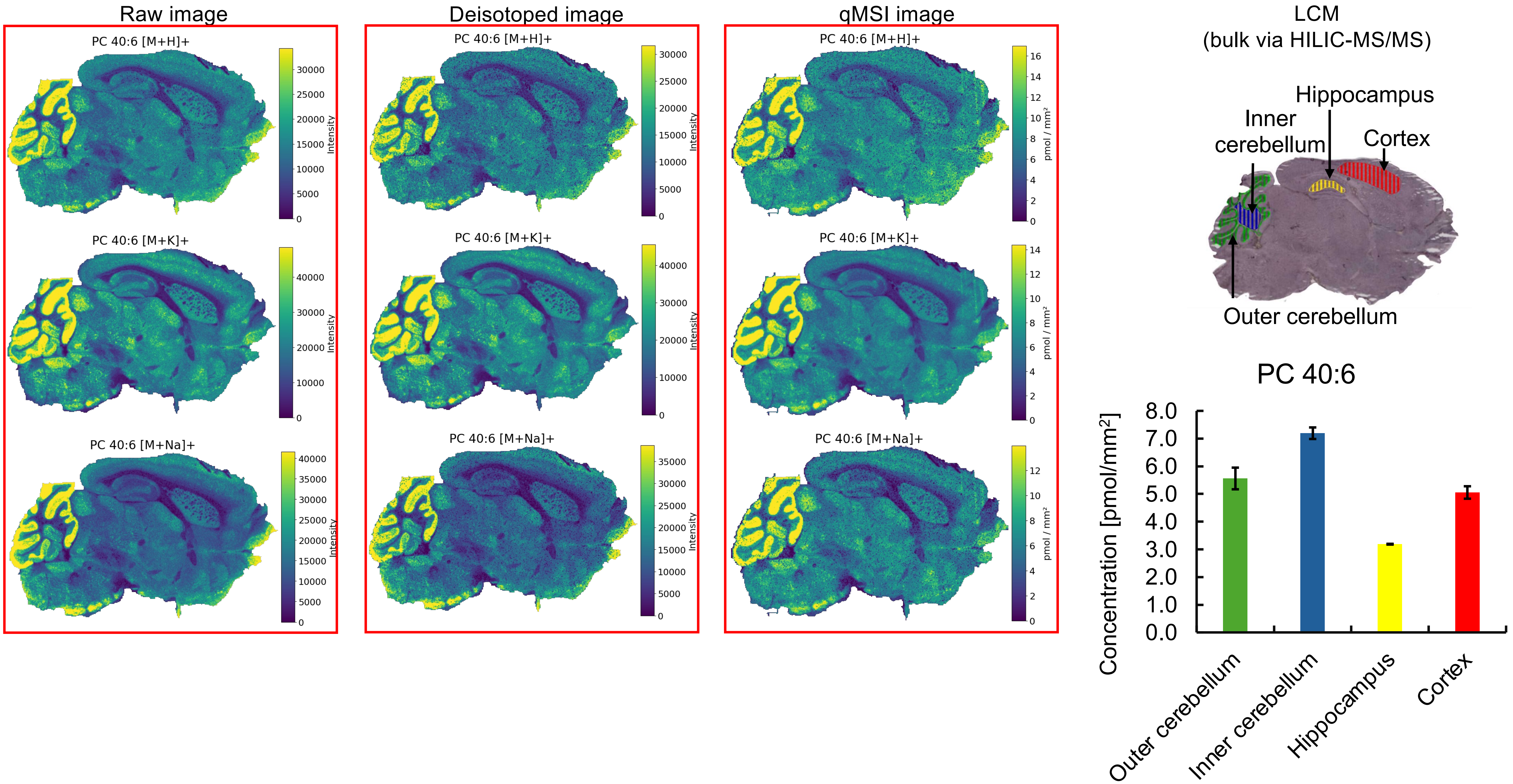

Raw image
Deisotoped image
qMSI image
LCM
(bulk via HILIC-MS/MS)
PC 40:6 [M+H]+
PC 40:6 [M+K]+
PC 40:6 [M+Na]+
Intensity
pmol / mm²
Hippocampus
Inner cerebellum
Cortex
Outer cerebellum
PC 40:6
Concentration [pmol/mm²]
Outer cerebellum
Inner cerebellum
Hippocampus
Cortex

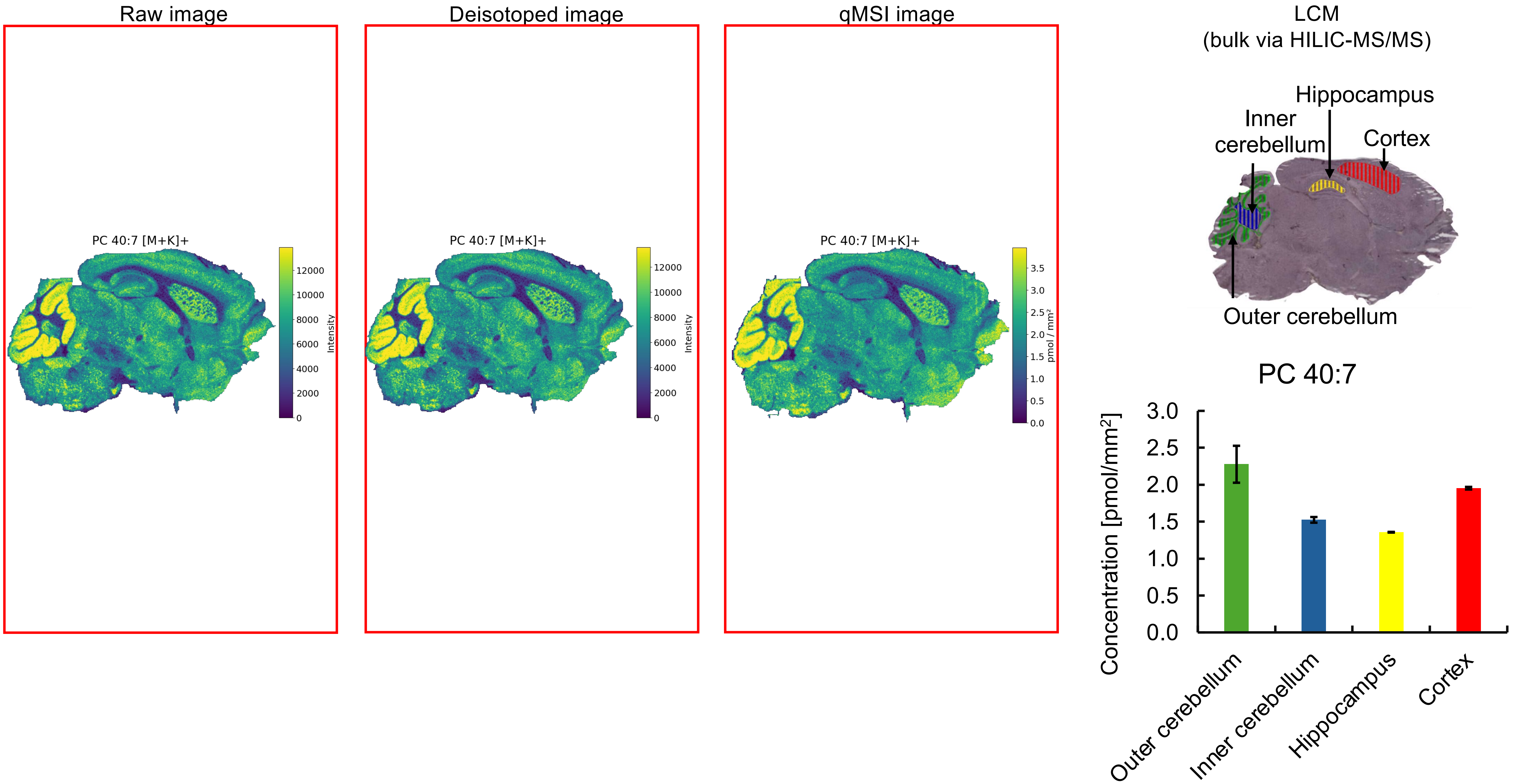

Raw image
Deisotoped image
qMSI image
LCM
(bulk via HILIC-MS/MS)
PC 40:7 [M+K]+
Intensity
pmol / mm²
Hippocampus
Inner cerebellum
Cortex
Outer cerebellum
PC 40:7
Concentration [pmol/mm²]
3.0
2.5
2.0
1.5
1.0
0.5
0.0
Outer cerebellum
Inner cerebellum
Hippocampus
Cortex

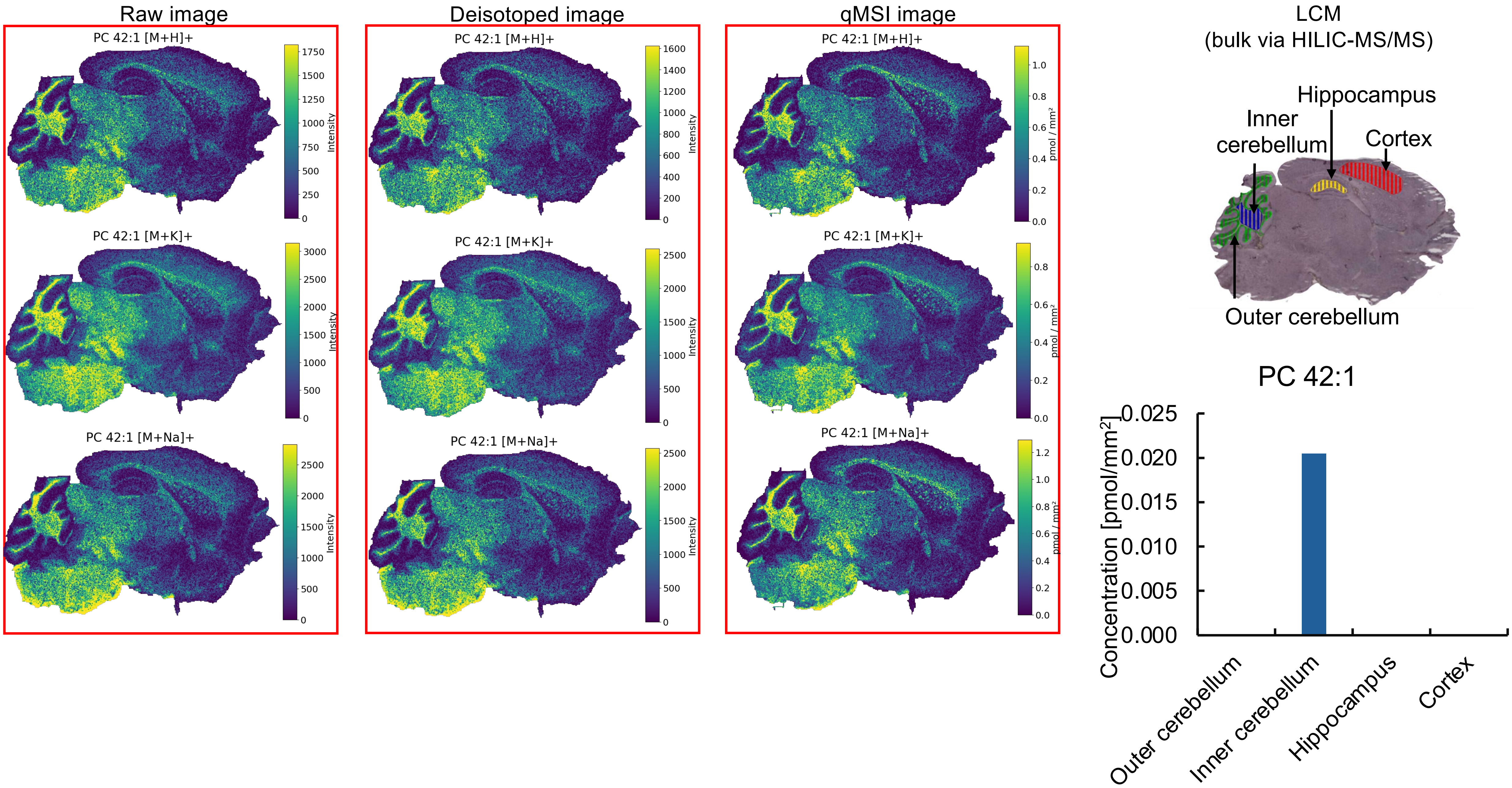

Raw image
PC 42:1 [M+H]+
Intensity
PC 42:1 [M+K]+
Intensity
PC 42:1 [M+Na]+
Intensity
Deisotoped image
PC 42:1 [M+H]+
Intensity
PC 42:1 [M+K]+
Intensity
PC 42:1 [M+Na]+
Intensity
qMSI image
PC 42:1 [M+H]+
pmol / mm²
PC 42:1 [M+K]+
pmol / mm²
PC 42:1 [M+Na]+
pmol / mm²
LCM
(bulk via HILIC-MS/MS)
Hippocampus
Inner cerebellum
Cortex
Outer cerebellum
PC 42:1
Concentration [pmol/mm²]
0.025
0.020
0.015
0.010
0.005
0.000
Outer cerebellum
Inner cerebellum
Hippocampus
Cortex

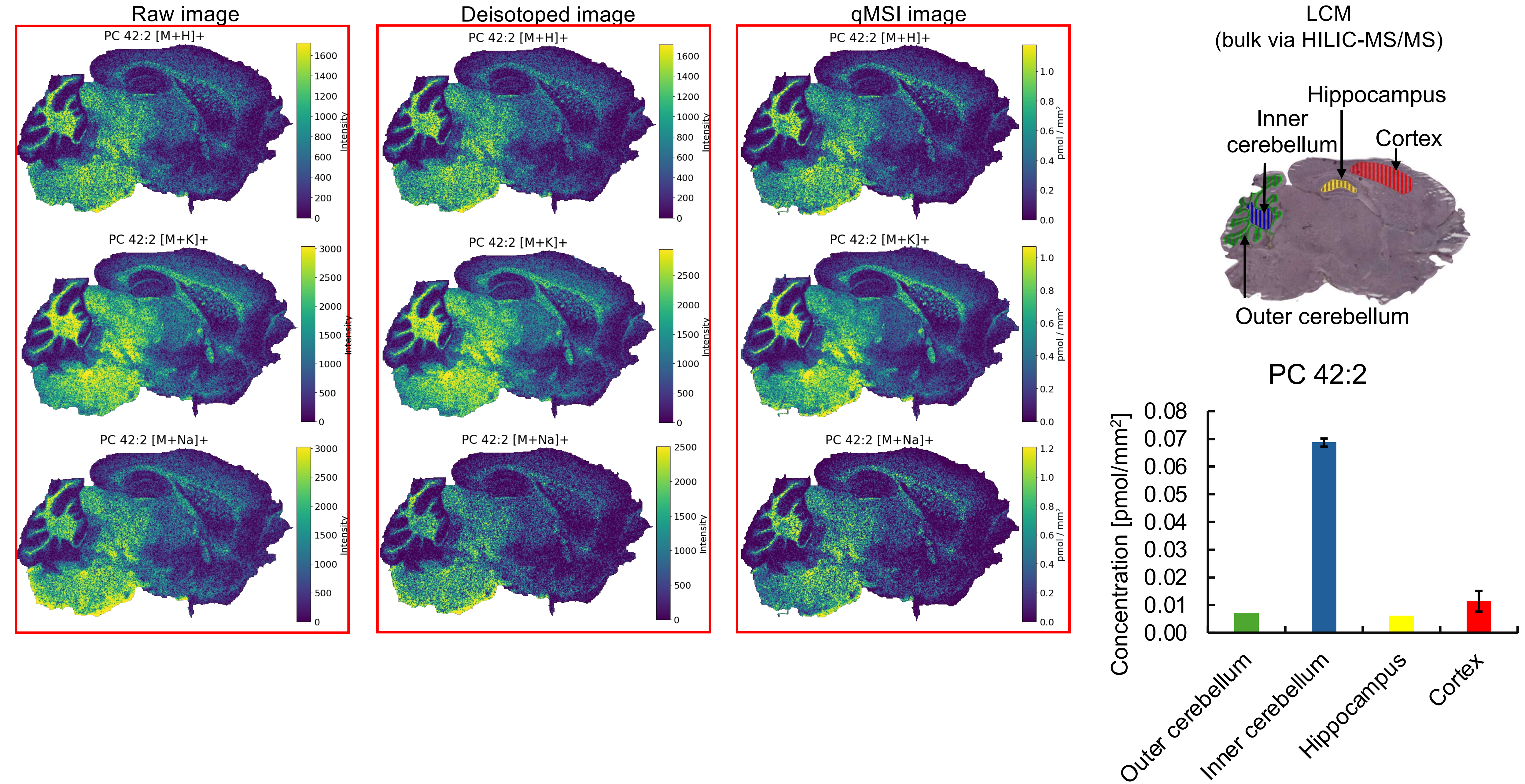

Raw image
Deisotoped image
qMSI image
LCM
(bulk via HILIC-MS/MS)
PC 42:2 [M+H]+
PC 42:2 [M+K]+
PC 42:2 [M+Na]+
Intensity
pmol / mm²
Hippocampus
Inner cerebellum
Cortex
Outer cerebellum
PC 42:2
Concentration [pmol/mm²]
Outer cerebellum
Inner cerebellum
Hippocampus
Cortex

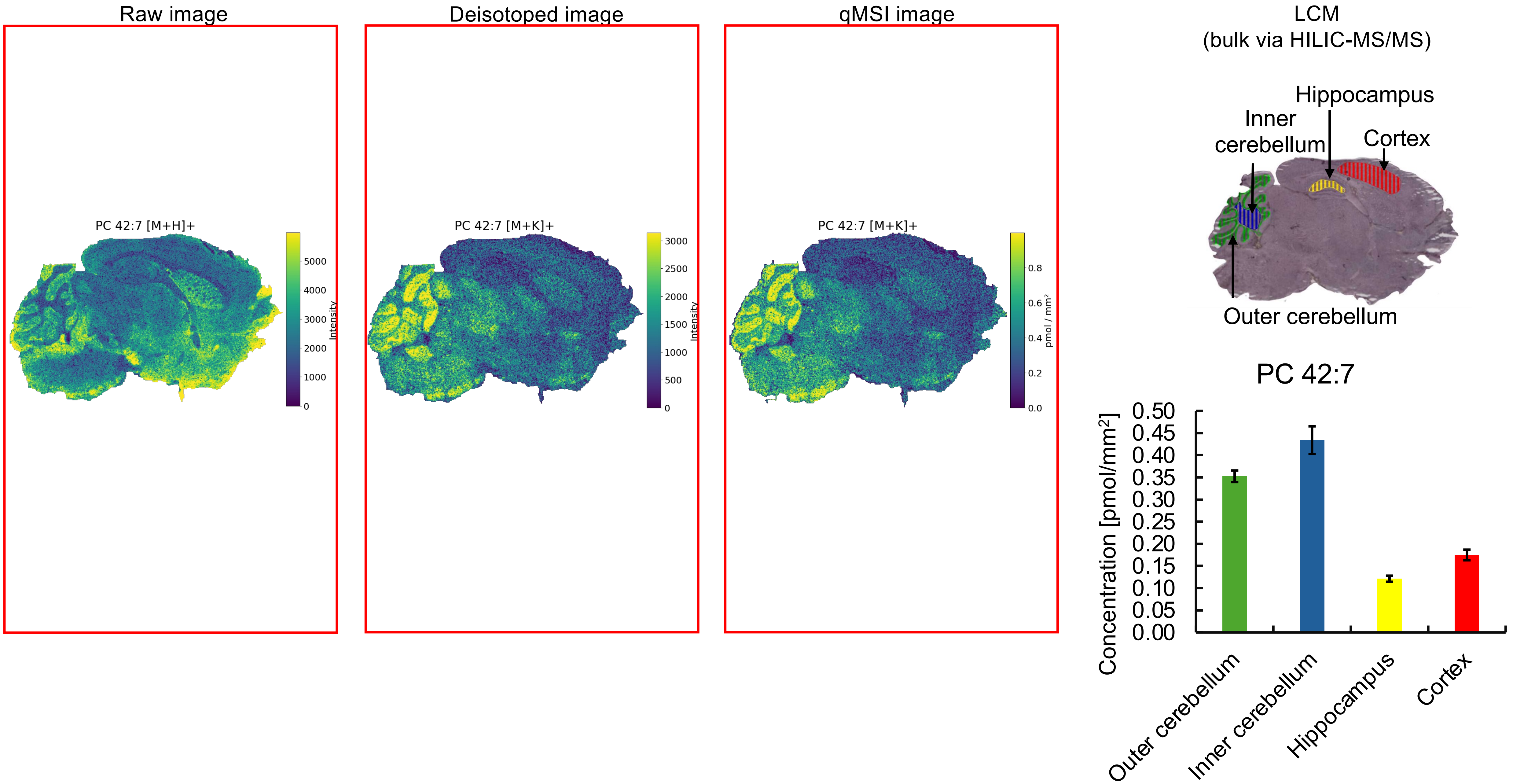

Raw image
Deisotoped image
qMSI image
LCM
(bulk via HILIC-MS/MS)
PC 42:7 [M+H]+
PC 42:7 [M+K]+
PC 42:7 [M+K]+
Intensity
Intensity
pmol / mm²
Hippocampus
Inner cerebellum
Cortex
Outer cerebellum
PC 42:7
Concentration [pmol/mm²]
Outer cerebellum
Inner cerebellum
Hippocampus
Cortex

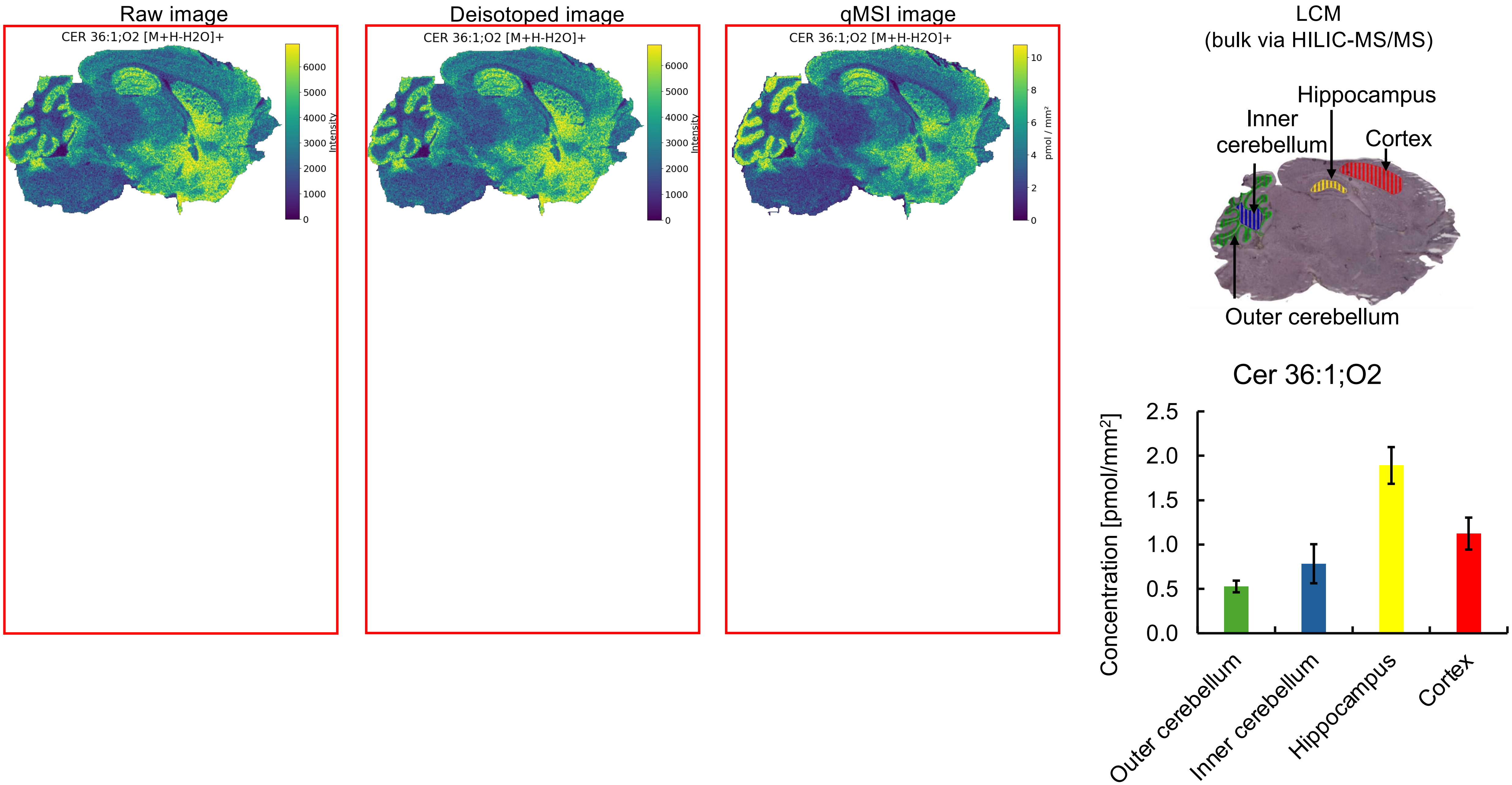
Raw image
CER 36:1;O2 [M+H-H2O]+
Intensity
0
1000
2000
3000
4000
5000
6000
Deisotoped image
CER 36:1;O2 [M+H-H2O]+
Intensity
0
1000
2000
3000
4000
5000
6000
qMSI image
CER 36:1;O2 [M+H-H2O]+
pmol / mm²
0
2
4
6
8
10
LCM
(bulk via HILIC-MS/MS)
Hippocampus
Inner cerebellum
Cortex
Outer cerebellum
Cer 36:1;O2
Concentration [pmol/mm²]
0.0
0.5
1.0
1.5
2.0
2.5
Outer cerebellum
Inner cerebellum
Hippocampus
Cortex

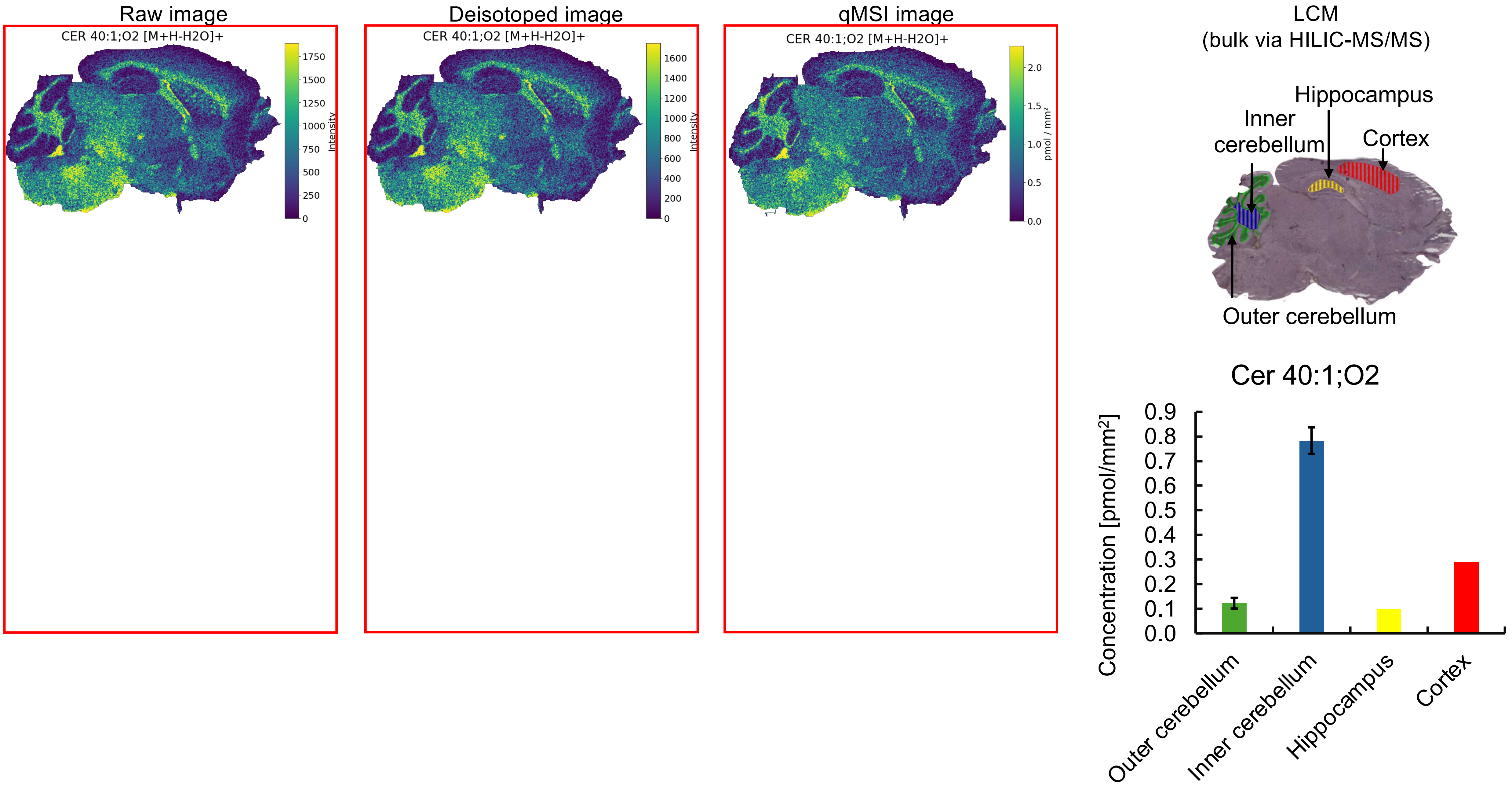

Raw image
CER 40:1;O2 [M+H-H2O]+
Intensity
0
250
500
750
1000
1250
1500
1750
Deisotoped image
CER 40:1;O2 [M+H-H2O]+
Intensity
0
200
400
600
800
1000
1200
1400
1600
qMSI image
CER 40:1;O2 [M+H-H2O]+
pmol / mm²
0.0
0.5
1.0
1.5
2.0
LCM
(bulk via HILIC-MS/MS)
Hippocampus
Inner cerebellum
Cortex
Outer cerebellum
Cer 40:1;O2
Concentration [pmol/mm²]
0.0
0.1
0.2
0.3
0.4
0.5
0.6
0.7
0.8
0.9
Outer cerebellum
Inner cerebellum
Hippocampus
Cortex

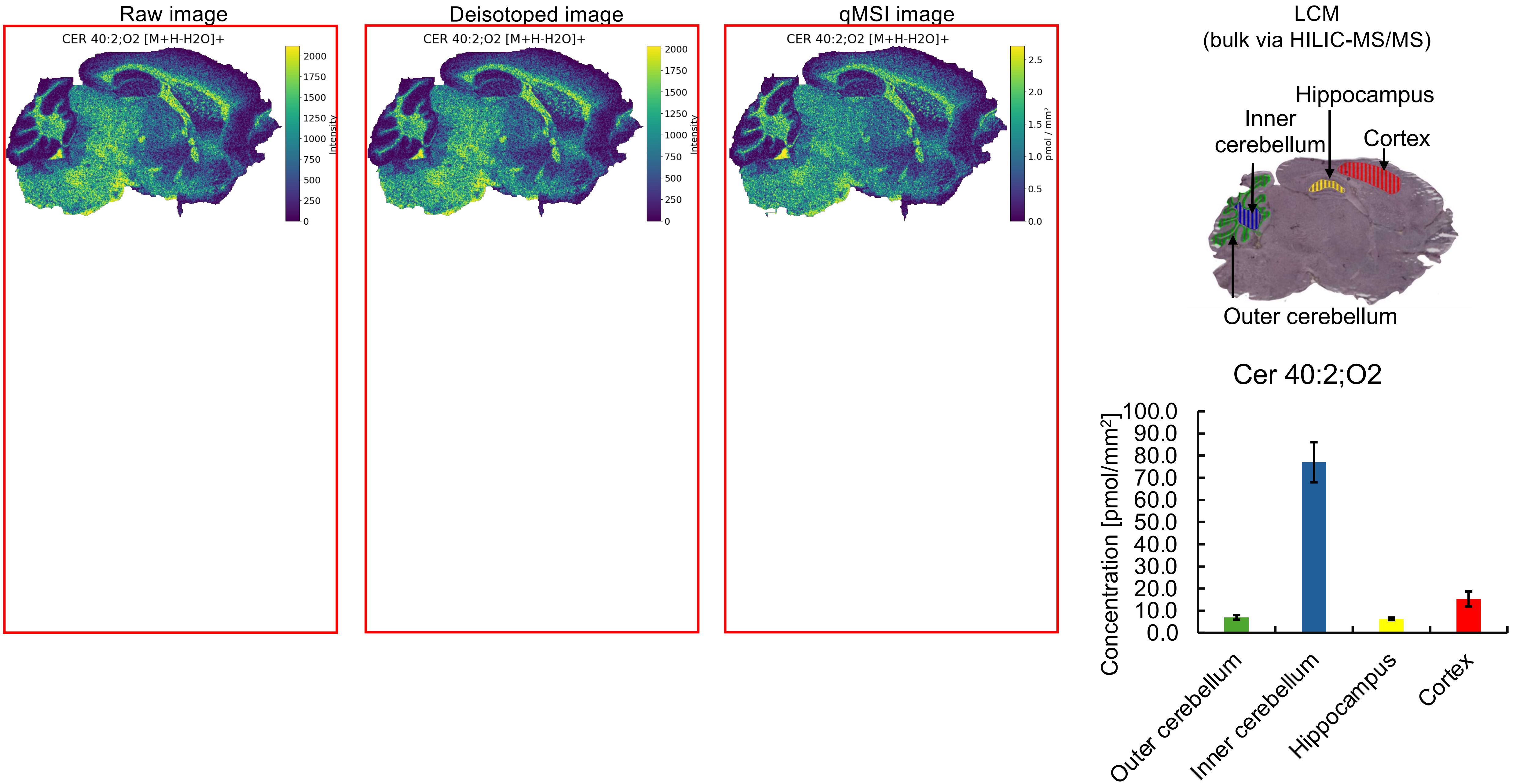
Raw image
CER 40:2;O2 [M+H-H2O]+
Intensity
Deisotoped image
CER 40:2;O2 [M+H-H2O]+
Intensity
qMSI image
CER 40:2;O2 [M+H-H2O]+
pmol / mm²
LCM
(bulk via HILIC-MS/MS)
Hippocampus
Inner cerebellum
Cortex
Outer cerebellum
Cer 40:2;O2
Concentration [pmol/mm²]
Outer cerebellum
Inner cerebellum
Hippocampus
Cortex

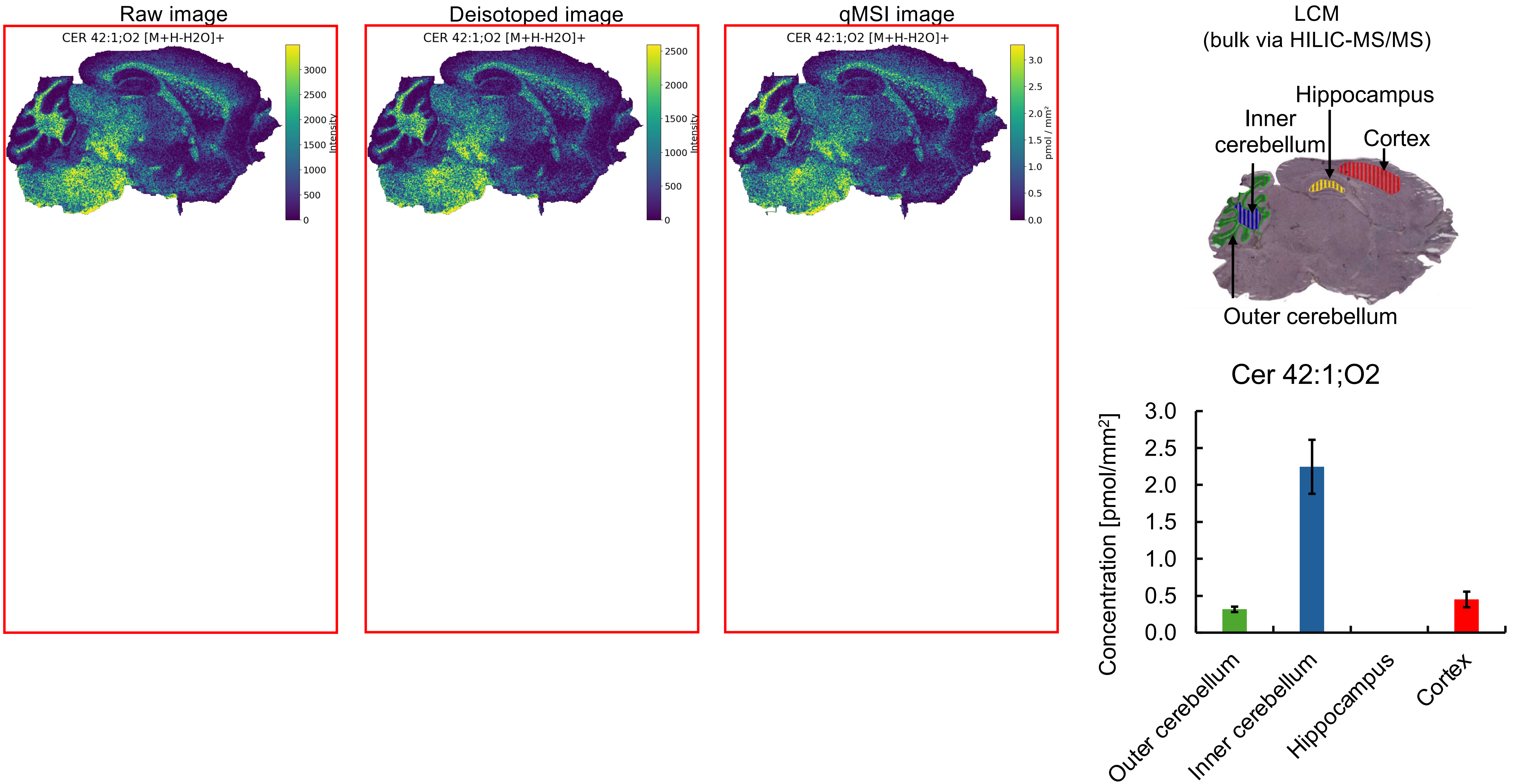
Raw image
CER 42:1;O2 [M+H-H2O]+
Intensity
Deisotoped image
CER 42:1;O2 [M+H-H2O]+
Intensity
qMSI image
CER 42:1;O2 [M+H-H2O]+
pmol / mm²
LCM
(bulk via HILIC-MS/MS)
Hippocampus
Inner cerebellum
Cortex
Outer cerebellum
Cer 42:1;O2
Concentration [pmol/mm²]
Outer cerebellum
Inner cerebellum
Hippocampus
Cortex

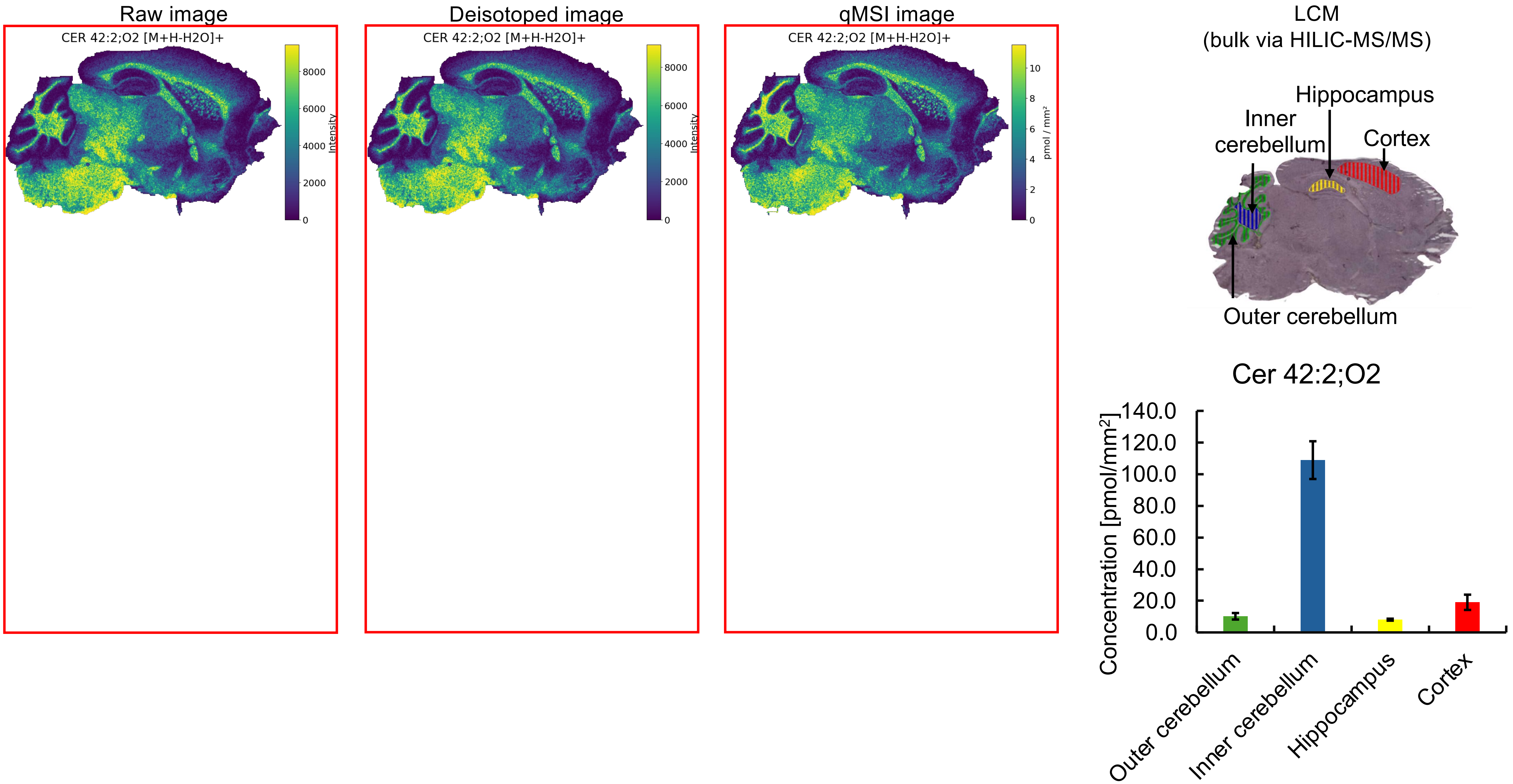
Raw image
CER 42:2;O2 [M+H-H2O]+
Intensity
8000
6000
4000
2000
0
Deisotoped image
CER 42:2;O2 [M+H-H2O]+
Intensity
8000
6000
4000
2000
0
qMSI image
CER 42:2;O2 [M+H-H2O]+
pmol / mm²
10
8
6
4
2
0
LCM
(bulk via HILIC-MS/MS)
Hippocampus
Inner cerebellum
Cortex
Outer cerebellum
Cer 42:2;O2
Concentration [pmol/mm²]
140.0
120.0
100.0
80.0
60.0
40.0
20.0
0.0
Outer cerebellum
Inner cerebellum
Hippocampus
Cortex

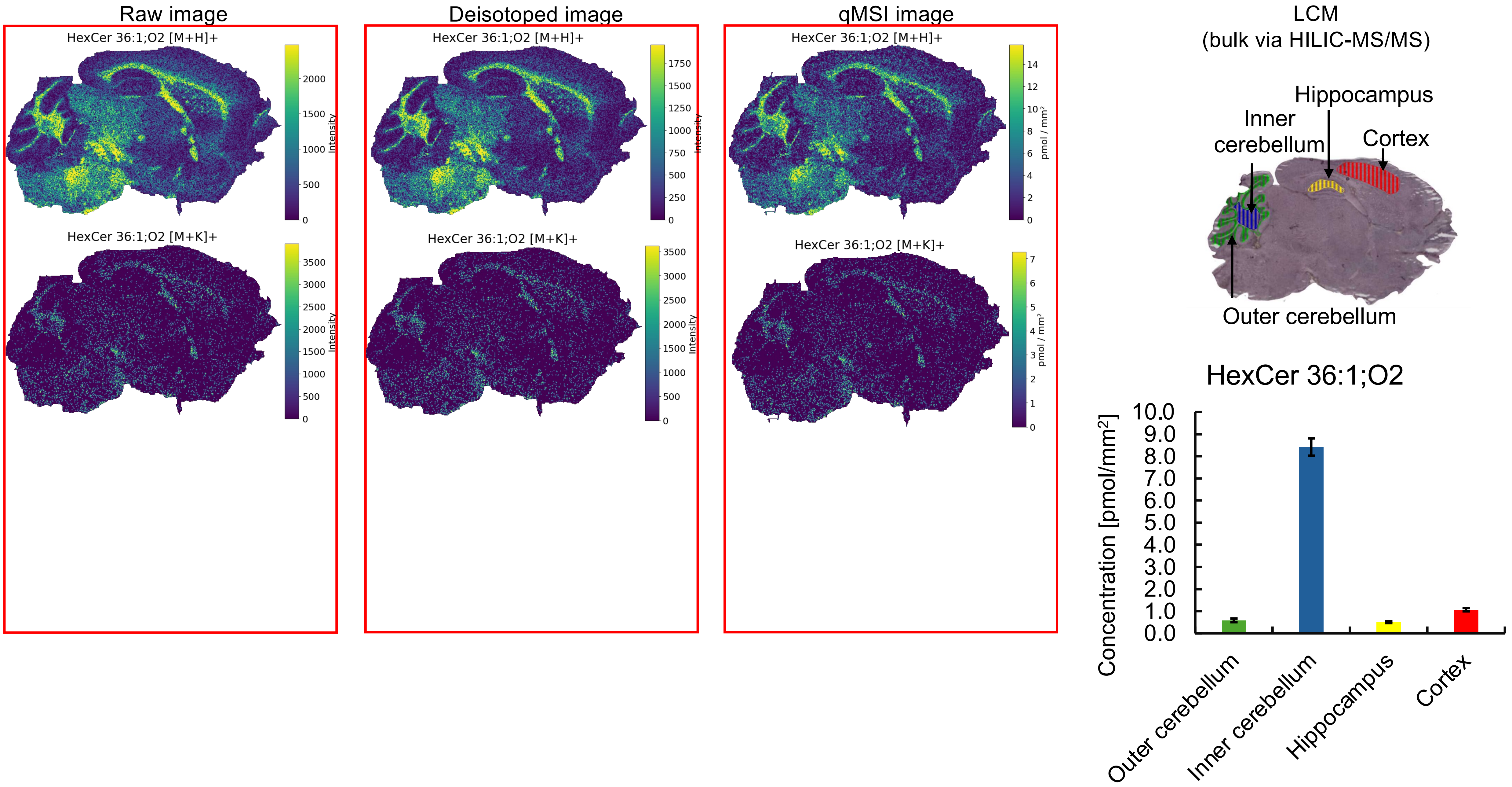


*Images normalized to Cer 30:1;O2 $[M\text{-}H_2O+H]^+$ (Insufficient correction for different adduct response)

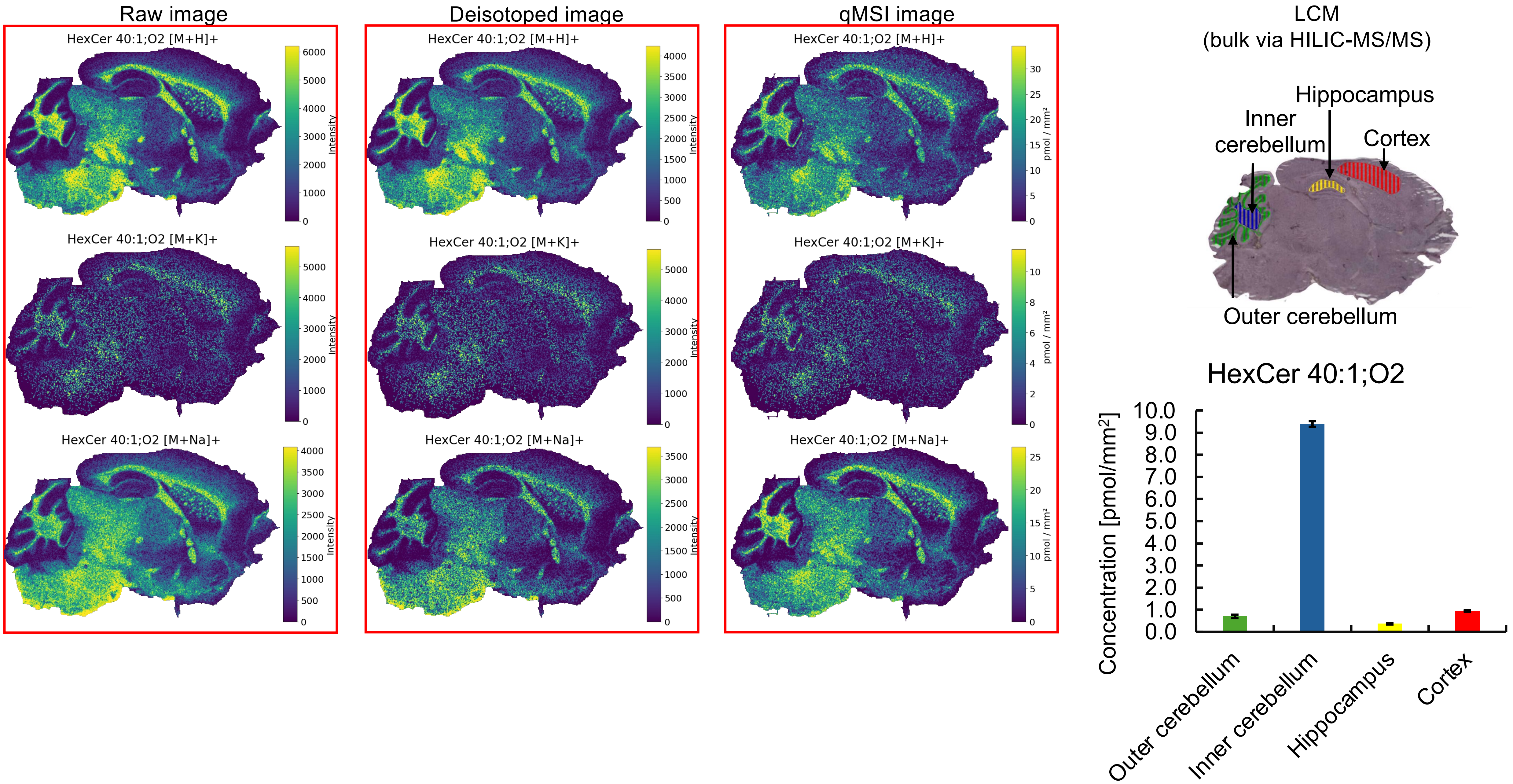


*Images normalized to Cer 30:1;O2 [M-$H_2O$+H]$^+$ (Insufficient correction for different adduct response)

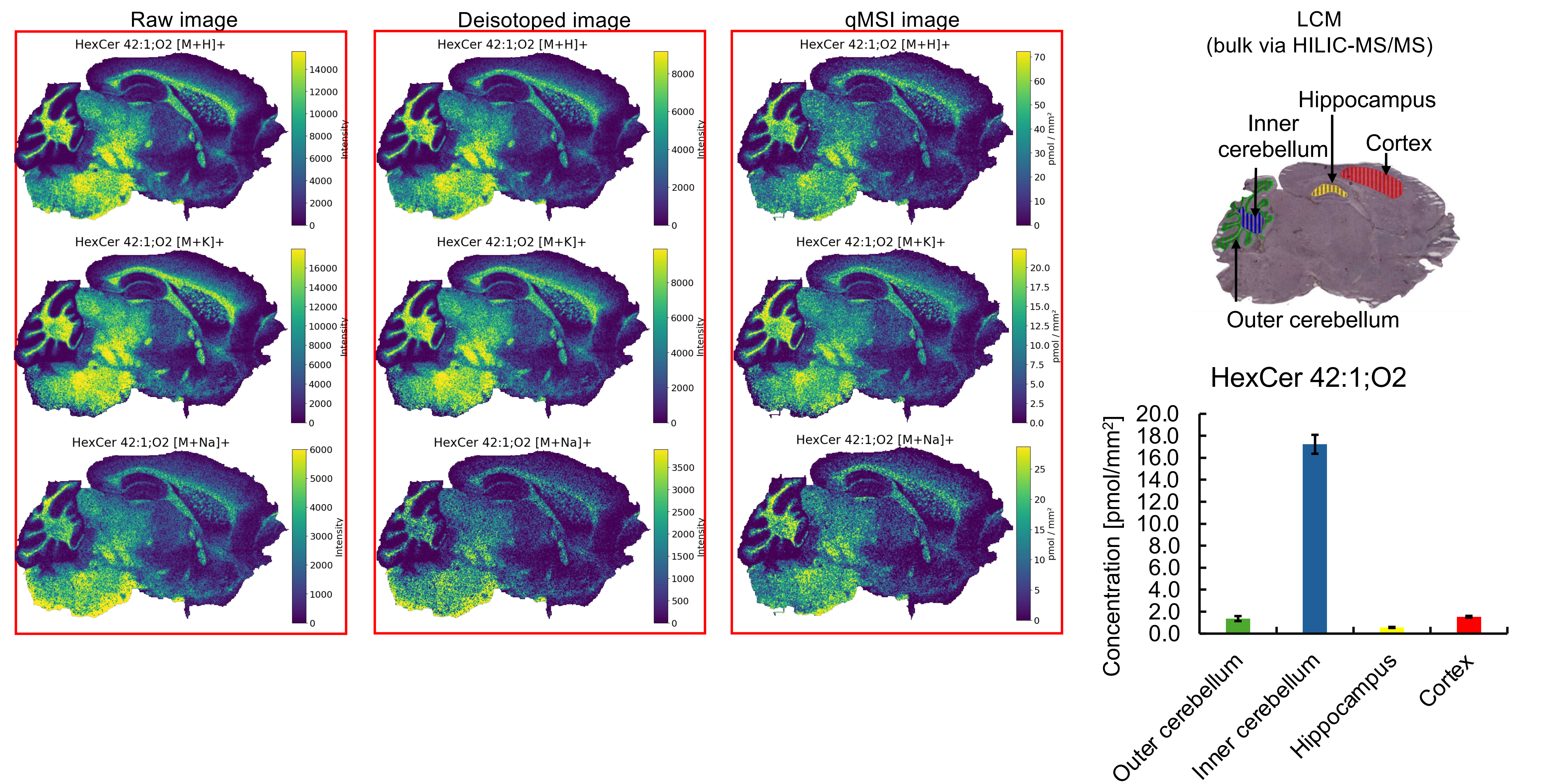


*Images normalized to Cer 30:1;O2 $[M-H_2O+H]^+$ (Insufficient correction for different adduct response)

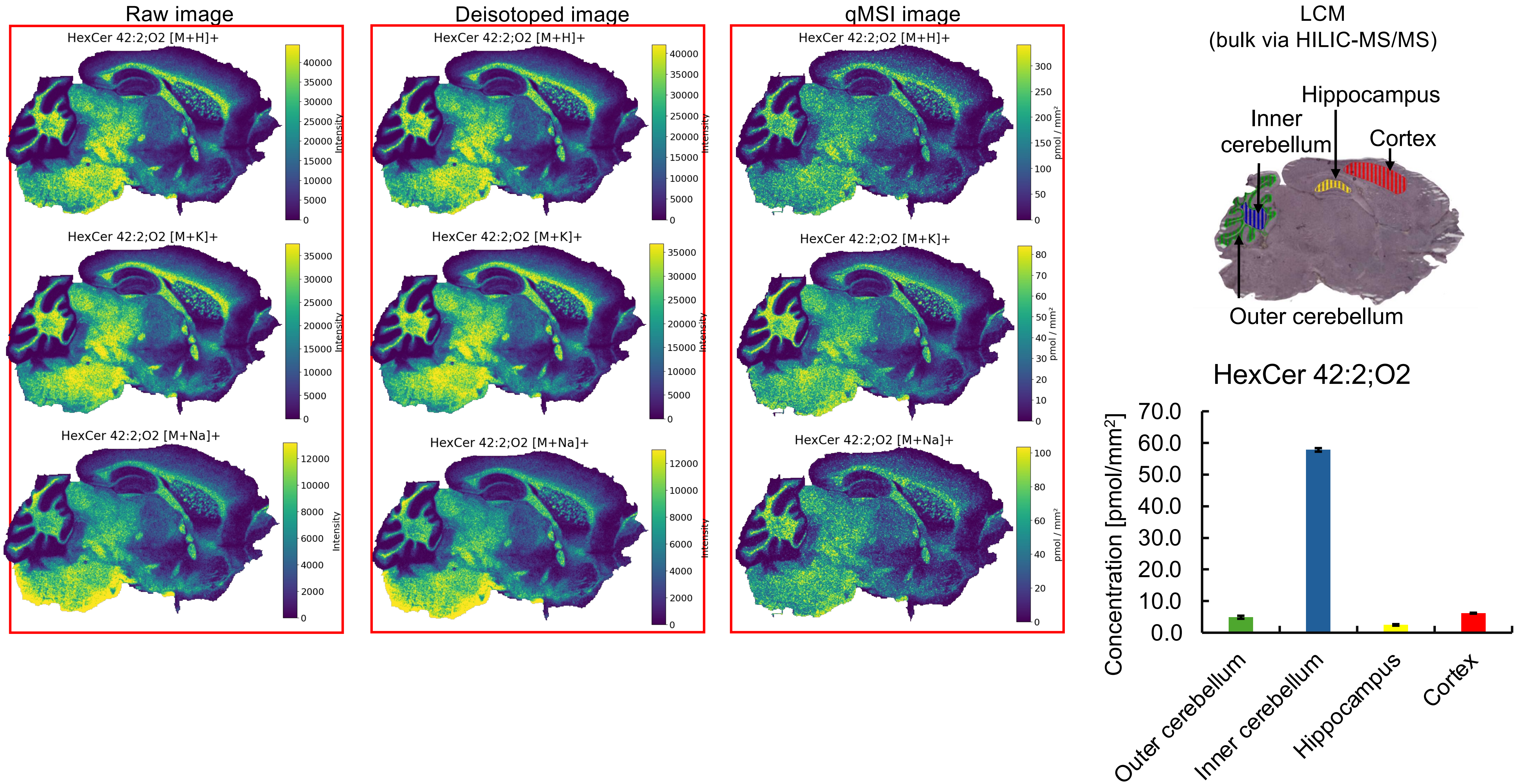


*Images normalized to Cer 30:1;O2 [M-$H_2O$+H]$^+$ (Insufficient correction for different adduct response)
